# Rewiring Oncogenic Transcriptional Complexes with Domain-ALTerAtion Chimeras (DALTACs) in Prostate Cancer

**DOI:** 10.64898/2026.04.24.720638

**Authors:** Jie Luo, Jianzhang Yang, Jean Ching-Yi Tien, Mi Wang, Sumit Das, Xiaoju Wang, Eleanor Young, Jelena Tošović, Rahul Mannan, Weiguo Xiang, Xia Jiang, Jocelyn Cai, Yihan Liu, Kenneth Gu, Somnath Mahapatra, Shiting Li, Yitong Yin, Sanjana Eyunni, Abigail J. Todd, Shicheng Jin, Xuhong Cao, Fengyun Su, Stephanie J. Miner, Ranga Sudharshan, Arvind Rao, Abhijit Parolia, Yuanyuan Qiao, Shaomeng Wang, Arul M. Chinnaiyan

**Affiliations:** Michigan Center for Translational Pathology, University of Michigan, Ann Arbor, MI, USA; Department of Pathology, University of Michigan, Ann Arbor, MI, USA; Division of Hematology-Oncology, Department of Internal Medicine, University of Michigan, Ann Arbor, MI, USA; Rogel Cancer Center, University of Michigan, Ann Arbor, MI, USA; Cancer Biology Program, University of Michigan, Ann Arbor, MI, USA; Howard Hughes Medical Institute, University of Michigan, Ann Arbor, MI, USA; Department of Computational Medicine and Bioinformatics, University of Michigan, Ann Arbor, MI, USA; Department of Urology, University of Michigan, Ann Arbor, MI, USA; Department of Pharmacology, University of Michigan, Ann Arbor, MI, USA; Department of Medicinal Chemistry, University of Michigan, Ann Arbor, MI, USA

**Keywords:** androgen receptor, p300, DALTAC, chemical-induced proximity, neo-enhanceosome, prostate cancer

## Abstract

Transcriptional addiction to the androgen receptor (AR) underlies metastatic castration-resistant prostate cancer (mCRPC)^1^, where AR maintains oncogenic programs through domain-specific interactions with the acetyltransferases p300/CBP^2^. Here, we describe a distinct therapeutic modality, Domain-ALTerAtion Chimeras (DALTACs), that rewire endogenous protein complexes by enforcing non-native domain–domain interactions. Our first-in-class AR–p300/CBP DALTAC-1 induces synthetic proximity between the AR ligand-binding domain and p300/CBP-bromodomain, misconfiguring their native interface and locking the complex in a non-productive state. DALTAC-1 triggers a “super-inhibitory” effect, suppressing AR-driven transcription and proliferation more potently than combined AR and p300/CBP inhibition. Mechanistically, DALTAC-1 reprograms p300/CBP substrate engagement, abolishing the histone mark H2B N-terminal acetylation (H2BNTac) while inducing neomorphic acetylation of AR and coactivators, ultimately collapsing the AR neo-enhanceosome. Chromatin profiling revealed redistribution of AR and p300/CBP toward palindromic AREs, accompanied by disruption of cofactor and transcriptional machinery recruitment at oncogenic AR/ERG neo-enhancers. DALTAC-1 exhibits exquisite lineage selectivity, displaying potent activity in AR-p300/CBP dual-dependent cells while sparing AR-negative cells. In multiple in vivo models, DALTAC-1 induces deep and durable tumor regressions. Together, these findings establish DALTACs as a broadly applicable strategy to rewire disease-defining complexes through domain-topology alteration, expanding induced-proximity therapeutics, while DALTAC-1’s precise lineage-selectivity supports its translational potential in AR-driven prostate cancer.

## Introduction

Aberrant transcriptional programs are a hallmark of cancer, driving malignant transformation, particularly in enhancer-driven cancers such as mCRPC^3,4^. AR is the central driver of mCRPC, undergoing androgen-induced activation and dimerization before binding androgen-response-element (ARE)-enriched enhancers. AR recruits coregulators to assemble cancer-specific neo-enhanceosome complexes^1^, promoting oncogenic transcriptional programs^4–6^.

Enhancer activation requires specific histone modifications, mediated by key coactivators such as the paralogous lysine acetyltransferases p300 and CBP^7^. Our previous work identified p300 as the predominant paralog supporting AR-positive (AR+) prostate cancer (PCa) and showed that p300/CBP, the p300/CBP-dependent enhancer mark H2B N-terminal acetylation (H2BNTac), and AR–p300 co-bound enhancer activity are elevated in prostate tumors^8^. Although inhibitors and degraders targeting p300/CBP have been developed for enhancer-driven malignancies, including CRPC^9^, global p300/CBP inhibition may cause toxicity in normal tissues^10^, highlighting the need for more selective therapeutic strategies.

Protein–protein interactions (PPIs) are tightly regulated by domain compatibility, which determines interaction specificity and function^11^. In transcriptional regulation, the enhancer complexes rely on precise domain–domain interactions for cofactor recruitment, enzymatic activation, and chromatin engagement^12^. For instance, upon chromatin binding and dimerization, the AR N-terminal domain (NTD) interacts with p300^2^. This interaction stimulates the histone acetyltransferase (HAT) activity of p300, leading to enhancer remodeling and transcriptional activation by establishing permissive chromatin^13,14^. Disrupting AR-p300/CBP interactions therefore represents an intriguing strategy to selectively target p300/CBP in AR-positive prostate cancer cells.

Chemical-induced proximity (CIP) strategies have revolutionized therapeutic design by reprogramming protein–protein interactions to manipulate cellular function^15^. Traditional CIP modalities include proteolysis-targeting chimeras (PROTACs), which promote target protein degradation via E3 ligase recruitment^16^, and more recent approaches such as regulated induced proximity targeting chimeras (RIPTACs) and transcriptional/epigenetic chemical inducers of proximity (TCIPs). RIPTACs are heterobifunctional molecules that couple a target protein to a pan-essential effector, causing effector inactivation and selective cell death^17^. TCIPs, by contrast, redirect transcriptional coactivators to activate pro-apoptotic genes normally repressed by BCL6^18–20^. Another emerging class, termed targeted relocalization-activating molecules (TRAMs), relocalizes endogenous proteins, displacing them from their native compartments and disrupting their functions^21^. Despite their mechanistic differences, current CIP approaches predominantly function by inducing proximity of non-native interacting proteins.

Here, we introduce a mechanistically distinct modality, Domain-ALTerAtion Chimeras (DALTACs), that selectively reprogram endogenous protein complexes by enforcing neomorphic domain–domain interactions between physiological partners. In contrast to CIP strategies that induce proteolysis or repurpose coactivators, DALTACs function as chemical dominant-negatives that lock transcriptional regulators into inert states. As a proof of concept, we developed DALTAC-1 (JZY3032), an AR–p300/CBP DALTAC that shows greater activity than combined AR and p300/CBP inhibition and selectively targets AR–p300/CBP-dependent cancer cells. DALTAC-1 enforces AR ligand-binding domain (LBD)–p300/CBP bromodomain (BRD) interactions, reprograms p300/CBP substrate engagement, eliminates H2BNTac and disrupts the oncogenic AR–p300/CBP transcriptional program. In multiple preclinical models, DALTAC-1 inhibits tumor growth or induces regression without overt toxicity. These findings establish DALTACs as a strategy to selectively disable native oncogenic protein complexes.

## Results

### Development of AR-p300/CBP DALTACs

Previous studies established that AR physically interacts with the coactivator p300 through its N-terminal domain (NTD)^2^, and that dimerized transcription factors can stimulate p300 histone acetyltransferase activity^13^. To reprogram this native interaction, we designed heterobifunctional molecules that tether an AR ligand-binding domain (LBD) inhibitor to a p300/CBP bromodomain (BRD) inhibitor through an appropriate linker. We reasoned that these molecules would induce non-native AR–p300/CBP interactions and thereby inhibit the growth of AR-positive cancer cells by concurrently inhibiting both AR and p300/CBP functions (**Fig. 1a**).

**Figure 1.**
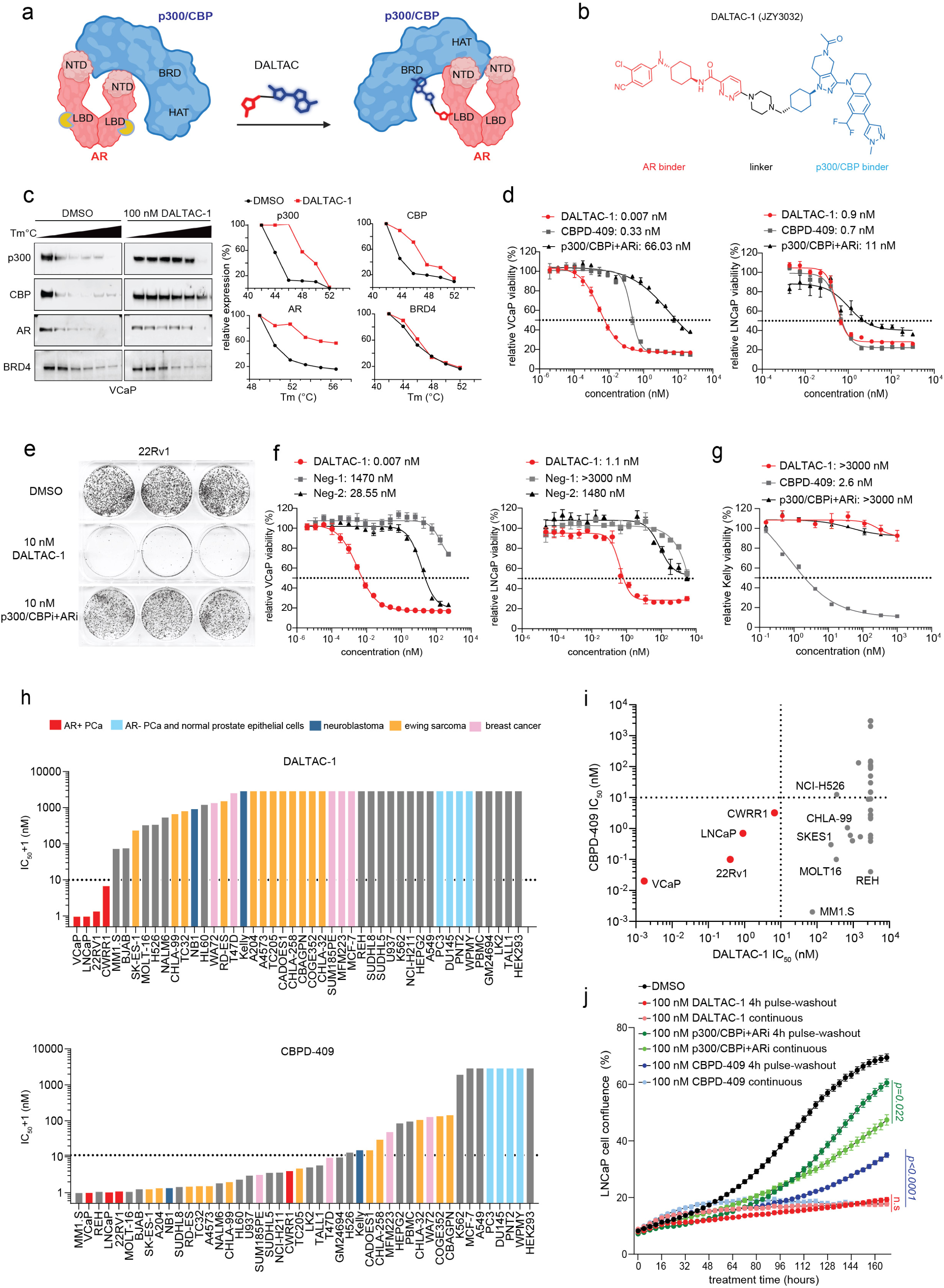
AR-p300/CBP DALTAC-1 selectively suppresses the growth of AR-positive prostate cancer cells. a. Schematic showing AR-p300/CBP DALTAC-1 rewires domain-domain interactions of the native AR-p300/CBP complex. NTD, AR N-terminal domain; CTD, AR C-terminal domain; BRD, p300 bromodomain; HAT, p300 histone acetyltransferase. b. Structure of DALTAC-1 (JZY3032), comprising AR and p300/CBP ligands connected by a linker. c. Cellular thermal shift assay of VCaP cells treated with DMSO or DALTAC-1 (100 nM, 4 h). Left, immunoblots showing stabilization of p300, CBP, and AR, but not BRD4; right, protein levels normalized to the lowest-temperature condition (three independent experiments). d. Dose-response curves and IC_50_ values of AR-positive VCaP and LNCaP cells treated with DALTAC-1, CBPD-409, or p300i+ARi for 5 days. (mean ± s.d.; n=3; three independent experiments). e. Representative colony-formation images of 22Rv1 cells treated for 14 days with DMSO, DALTAC-1, or p300/CBPi+ARi (10 nM each; n=3; three independent experiments).. f. Dose-response curves and IC_50_ values of AR-positive VCaP and LNCaP cells treated with DALTAC-1, Neg-1 (Negative control-1), or Neg-2 (Negative control-2) for 5 days (mean ± s.d.; n=3; three independent experiments). g. Dose-response curves and IC_50_ values of Kelly cells treated with DALTAC-1, CBPD-409, or p300i+ARi for 5 days (mean ± s.d.; n=3; three independent experiments). h. Ranked IC_50_ values for DALTAC-1 and CBPD-409 across 46 human cell lines after 5 days; colors indicate cell type. i. Comparison of DALTAC-1 and CBPD-409 IC_50_ values from Fig. 1h. AR-positive prostate cancer lines are red; dotted lines indicate 10 nM. j. Growth curves of LNCaP cells treated with 100 nM DALTAC-1, p300/CBPi + ARi, or CBPD-409 under 4-h pulse–washout or continuous-treatment conditions, monitored using Incucyte Live-Cell Analysis (mean ± s.d.; n=6; three independent experiments). Two-way ANOVA with no correction was used. Exact P values are shown; n.s., not significant.

To test this premise, we designed and synthesized a series of heterobifunctional molecules using a potent p300/CBP BRD inhibitor (GNE-049^22^) and a selective AR LBD binder^23^ tethered through a linker (**Extended Data Fig. 1a**). We selected the tethering site in the p300/CBP BRD inhibitor based upon the co-crystal structure of GNE-049 and its analogues, which showed that our selected tethering site was exposed to the solvent environment^24^. Similarly, we chose the tethering site in the AR inhibitor based upon our modeled complex structure with AR and extensive structure-activity relationships reported for this class of AR inhibitors^23,25^. We evaluated our synthesized compounds with proliferation assays in AR-positive LNCaP and 22Rv1 cells, which were shown to be responsive to p300/CBP BRD inhibitors and PROTAC degraders^9,26^. GNE-049, the AR binder (JZY3221), and their combination were included as controls. The individual treatment of GNE-049 and JZY3221 or their combination displayed limited inhibitory activity, particularly in 22Rv1 cells (**Extended Data Fig. 1b**).

To facilitate the synthesis of our designed heterobifunctional compounds, we replaced the tetrahydropyran group in GNE-049 with a piperidinyl group, which was shown to not affect the binding to p300^24^. Directly connecting the AR inhibitor through its pyridazine to the nitrogen atom of the piperidinyl group in the p300/CBP inhibitor yielded heterobifunctional compound **1**, which showed an improved cell growth inhibition activity in both LNCaP and 22Rv1 cells over the combination of GNE-049 and JZY3221 (**Extended Data Fig. 1b**). Inserting a short and rigid 4-membered azetidine between the pyridazine of the AR inhibitor and the nitrogen atom of the piperidinyl group in the p300/CBP inhibitor as the linker in compound **1** generated compound **2**, which had IC_50_ values of 33.2 nM and 16.4 nM, respectively, in the LNCaP and 22Rv1 cells, with Imax = 69-70% (**Extended Data Fig. 1b**). Replacing the 4-membered azetidine linker in compound **2** with a 6-membered piperidine linker generated compound **3**, which was equally potent and effective compared to compound **2** in 22Rv1 cells but was more potent than compound 2 in LNCaP cells (**Extended Data Fig. 1b**). Extending the linker by one methylene group in compound **2** led to compound **4**, which was equally potent and effective as compared to compound **3** in both cell lines (**Extended Data Fig. 1b**). Changing the 4-membered azetidine in compound **2** with a [6,6] spiro ring system resulted in compound **5**, which was also similarly potent and effective as compared to compound **2** in 22Rv1 cells but more potent than compound **2** in LNCaP cells (**Extended Data Fig. 1b**). Inserting a methylene between the [6,6] spiro ring and the piperidinyl group in compound **5** generated compound **6**, which was similarly potent and effective as compared to compound **5** in both cell lines (**Extended Data Fig. 1b**). We next synthesized compound **7** using a methyl-piperazinyl group as the linker but changing the piperidinyl group in the p300/CBP inhibitor with a trans-cyclohexyl group. Compound **7** (JZY3031) achieved IC_50_ = 1.9 nM and 2.6 nM in LNCaP and 22Rv1 cells, respectively, and I_max_ = 70-72%. Hence, compound **7** was >300-times more potent than the combination of the AR inhibitor and GNE-049 based upon their IC_50_ values and more effective based upon their Imax values (**Extended Data Fig. 1b**).

Pharmacokinetic (PK) data in mice showed that compound **7** had a slow clearance, a moderate volume of distribution, and excellent exposure with intravenous administration. Surprisingly, despite its high molecular weight (MW = 962), compound **7** achieved appreciable oral exposure with C_max_ = 269 ng/mL and AUC_0-t_ = 798 h* ng/mL at 5.0 mg/kg oral dose and an oral bioavailability of 6.5% (**Extended Data Fig. 1c**).

Our previous study showed that replacement of the oxygen atom with an *N*-methyl group in the AR inhibitor significantly improved oral bioavailability^25^. Accordingly, we replaced the oxygen atom with an *N*-methyl group in the AR inhibitor portion in compound **7**, which yielded JZY3032 (**Fig. 1b**). JZY3032 attained IC_50_ values of 0.26 nM and 1.0 nM in LNCaP and 22Rv1 cells, respectively, and I_max_ = 66-69% and was therefore more potent than compound **7** (**Extended Data Fig. 1b**). A PK study in mice showed that JZY3032 had a slower clearance, a higher volume of distribution, and a higher exposure than compound **7** with intravenous administration and achieved good oral exposure (C_max_ = 284 ng/mL and AUC_0-t_ = 2952 h* ng/mL at 5.0 mg/kg dose) and an oral bioavailability of 18.5% (**Extended Data Fig. 1c**). Hence, JZY3032 had a significantly improved PK profile over compound **7**.

We evaluated our synthesized bifunctional molecules for their ability to induce formation of an AR-p300 complex using an ELISA assay and identified that compound **7** and JZY3032 were the most potent and effective compounds in inducing the AR–p300 interaction (**Extended Data Fig. 1b**). In addition, we evaluated the effects of the indicated compounds on AR transactivation using HEK293 ARE-GFP reporter cells. Among all compounds tested, DALTAC-1 showed the strongest inhibition of AR transcriptional activity, which is consistent with its potent cell growth inhibition activity (**Extended Data Fig. 1b**).

To determine whether an alternative clinically established AR LBD inhibitor could further improve the activity, we designed and synthesized three enzalutamide-based DALTACs and benchmarked them against JZY3032 (**Extended Data Fig. 1d**). In CellTiter-Glo proliferation assays, while the three enzalutamide-based compounds exhibited significant activity in LNCaP cells, they were much weaker than JZY3032 (**Extended Data Fig. 1e**). Furthermore, these enzalutamide-based compounds showed weak or no activity in 22Rv1 cells. These data highlighted the importance of the AR ligand for the development of highly potent and effective DALTAC compounds.

Based upon our extensive mechanistic data discussed later in the manuscript, we named JZY3032 as DALTAC-1 (Domain-ALTerAtion Chimera 1). For mechanistic investigations, two control compounds, Negative control-1 (Neg-1) and Negative control-2 (Neg-2), were designed and synthesized (**Extended Data Fig. 1f**). Neg-1 was designed by reducing the binding with p300/CBP^24^ and Neg-2 was designed by reducing its binding to AR^27^ (**Extended Data Fig. 1f**). Fluorescence polarization (FP) assays confirmed that DALTAC-1 bound both p300/CBP and AR with comparable affinities to their respective parent inhibitors (**Extended Data Fig. 1g-i**). Furthermore, while Neg-1 displayed a similar binding affinity to AR compared to DALTAC-1, it had a much weaker affinity to p300 and CBP than DALTAC-1 (**Extended Data Fig. 1g-i**). In comparison, while Neg-2 had a similar binding affinity to p300 and CBP compared to DALTAC-1, it showed much weaker affinity to AR than DALTAC-1 (**Extended Data Fig. 1g-i**). Cellular thermal shift assays (CETSA) revealed a pronounced increase in the thermal stability of AR, p300, and CBP, but not BRD4, upon DALTAC-1 treatment, supporting strong and specific engagement of DALTAC-1 with its intended targets (**Fig. 1c** and **Extended Data Fig. 1j**).

Collectively, by tethering an AR LBD inhibitor and a p300/CBP BRD domain inhibitor with appropriate linkers, we have identified a series of heterobifunctional molecules. A number of them achieved superior cell growth inhibitory activity compared to the combination of the parent AR inhibitor and the corresponding p300/CBP BRD inhibitor. In particular, DALTAC-1 was >1,000 times more potent than the combination of the AR inhibitor and the p300/CBP BRD inhibitor in cell growth inhibition in the LNCaP and 22Rv1 cell lines. Despite its high MW of 975, DALTAC-1 achieved a good oral bioavailability and an excellent overall PK profile in mice.

### DALTAC-1 selectively inhibits AR+ PCa

To further assess the activity and selectivity of AR–p300 DALTAC-1, we compared its cell growth inhibitory effects with those of the potent and efficacious p300/CBP dual degrader CBPD-409^8^ and the combination of p300/CBP BRD inhibitor GNE-049 (p300i) and AR-LBD binder JZY3221 (ARi) in various AR-positive prostate cancer cell lines. DALTAC-1 treatment resulted in potent cytotoxicity across all tested AR-positive prostate cancer cell lines (VCaP, LNCaP, 22Rv1, and CWRR1), similar to or better than that observed with CBPD-409 (**Fig. 1d** and **Extended Data Fig. 2a**). In each model, DALTAC-1 demonstrated superior activity relative to the combined treatment with AR-LBD and p300/CBP-BRD inhibitors in all tested AR-positive prostate cancer cells, indicating a gain-of-function mechanism arising from enforced miswiring of the AR–p300 domain interactions (**Fig. 1d** and **Extended Data Fig. 2a**). Notably, DALTAC-1 also retained potent activity in the AR-V7–high, p300/CBP-dependent 22Rv1 and CWR-R1 cell lines, demonstrating efficacy in prostate cancer models expressing AR splice variants (**Extended Data Fig. 2a**). Moreover, DALTAC-1 exhibited greater antiproliferative activity than several clinically relevant AR-directed therapies, including the standard-of-care antiandrogens enzalutamide and darolutamide^28,29^, the AR PROTAC degrader ARV-766^30^, and the recently reported AR-BRD4 RIPTAC-1^31^ (**Extended Data Fig. 2b**). Consistently, colony formation assays revealed that DALTAC-1 abolished the clonogenic potential of AR-positive prostate cancer cells, outperforming the combination of AR-LBD and p300/CBP-BRD inhibitors (**Fig. 1e** and **Extended Data Fig. 2c**). In contrast, the inactive analogs Neg-1 and Neg-2 exhibited dramatically reduced cytotoxic effects (**Fig. 1f** and **Extended Data Fig. 2d**). Furthermore, the addition of p300/CBPi or ARi in molar excess competitively blocked DALTAC-1 binding to p300/CBP or AR and significantly attenuated its activity, indicating that engagement of both AR and p300/CBP was essential for the superior activity of DALTAC-1 (**Extended Data Fig. 2e**).

Compared with CBPD-409, DALTAC-1 exhibited markedly reduced cytotoxicity in AR-negative yet p300-dependent cancer cells (Kelly and NCI-H211)^8^ (**Fig. 1g** and **Extended Data Fig. 2f**). Consistent with its requirement for dual AR and p300/CBP dependency, DALTAC-1 was highly active in AR-positive MDA-MB-453 cells^32^, which are sensitive to both CBPD-409 and the AR degrader ARV-766 (**Extended Data Fig. 2g**). In contrast, DALTAC-1 showed minimal activity in AR-positive U-87 MG cells^33^, which are modestly sensitive to CBPD-409 but resistant to ARV-766 (**Extended Data Fig. 2g**). These findings support the selectivity of DALTAC-1 for tumors that are dependent on both AR and p300/CBP. To further validate this selectivity, we systematically compared the anti-proliferative effects (IC_50_ values) of DALTAC-1 and CBPD-409 across a broad panel of cancer cell lines. In contrast to the broad anti-proliferative activity of CBPD-409 across diverse cancer lineages, DALTAC-1 exhibited remarkable lineage selectivity, effectively inhibiting proliferation exclusively in AR-positive prostate cancer cells (**Fig. 1h, i**). Notably, DALTAC-1 also exhibited prolonged cytotoxic effects. A 4-hour pulse treatment of DALTAC-1 followed by compound washout resulted in comparable inhibition to continuous exposure, whereas washout of CBPD-409 or the p300i + ARi combination led to a marked reduction in anti-proliferative activity (**Fig. 1j** and **Extended Data Fig. 2h, i**). These results highlight the durable and sustaining nature of DALTAC-1 activity. Collectively, these findings establish DALTAC-1 as a potent and selective heterobifunctional compound with strong inhibitory activity in AR-positive prostate cancer cells.

### DALTAC-1 induces AR-p300 ternary complex

Ternary complex formation between target proteins and a small molecule is a central mechanism underlying the CIP strategy^34^. To determine whether DALTAC-1 directly promotes AR–p300 association independently of other cellular proteins, we performed an in vitro interaction assay using purified full-length HA-AR and Flag-p300 proteins. DALTAC-1 induced AR–p300 complex formation in a concentration-dependent manner (**Fig. 2a**). In contrast, neither the combination of the individual AR and p300/CBP inhibitors nor the p300/CBP-binding-deficient Neg-1 or AR-binding-deficient Neg-2 promoted AR–p300 association (**Extended Data Fig. 3a**), demonstrating that complex formation requires the simultaneous engagement of both proteins by intact DALTAC-1. We next obtained orthogonal biophysical evidence by biolayer interferometry (BLI) using flag-tagged full-length p300 and purified GST–AR-LBD proteins. In the presence of 10 μM DALTAC-1, increasing concentrations of GST–AR-LBD produced a concentration-dependent binding response, yielding an apparent K_d_ of 333.1 nM under these assay conditions (**Fig. 2b**). In contrast, matched titration of GST–AR-LBD in the absence of DALTAC-1 resulted in little to no detectable binding to p300, indicating that DALTAC-1 promotes assembly of an AR–p300 complex that is not detectable under the same conditions in its absence (**Fig. 2b**). Reciprocally, at a fixed concentration of GST–AR-LBD protein, increasing concentrations of DALTAC-1 progressively enhanced the binding response, yielding an apparent K_d_ of approximately 4.3 μM (**Extended Data Fig. 3b**). As a specificity control, the assay was performed using immobilized Flag–FOXA1 (residues 1-268), which is not directly engaged by DALTAC-1. Neither DALTAC-1 nor GST-AR-LBD alone, nor their combination, produced a significant binding response on the FOXA1 control surface (**Extended Data Fig**. **3c**), excluding nonspecific association of either the compound or GST-AR-LBD to the immobilized protein or the sensor. Notably, neither the AR LBD inhibitor nor the p300/CBP BRD inhibitor, alone or in combination, induced ternary-complex formation (**Extended Data Fig. 3d**). Conversely, an excess concentration of the p300/CBP BRD inhibitor markedly attenuated the DALTAC-1-induced ternary-complex BLI signal (**Extended Data Fig. 3e**). Together, these reciprocal titration and negative-control experiments demonstrate that DALTAC-1 specifically recruits AR-LBD to p300 and promotes concentration-dependent formation of the AR–DALTAC-1–p300 ternary complex.

**Figure 2.**
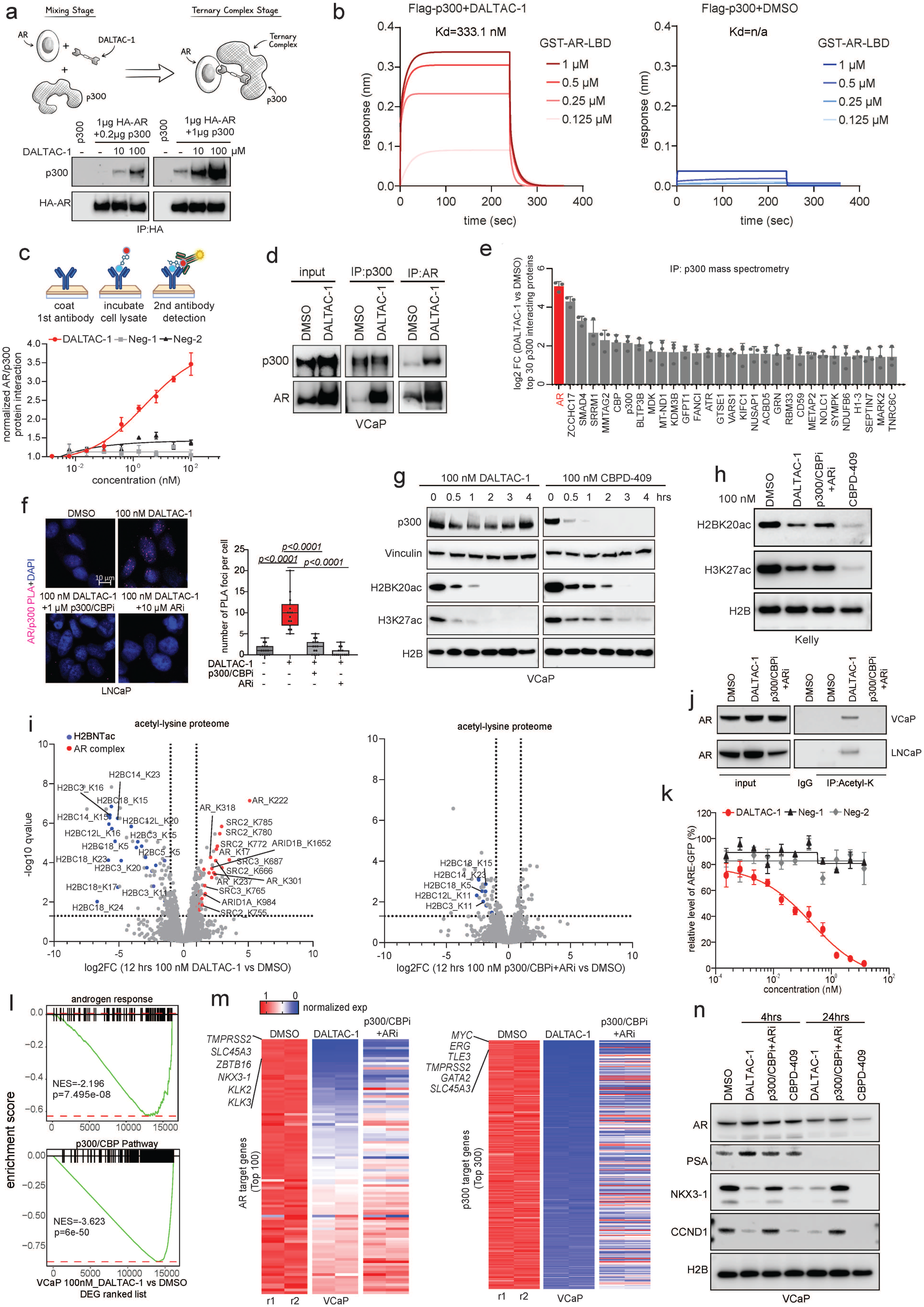
DALTAC-1-induced proximity of AR and p300/CBP proteins inhibits their oncogenic transcriptional output in AR-positive prostate cancer cells. a. Schematic and immunoblot showing dose-dependent DALTAC-1-induced association of purified AR-p300 proteins (three independent experiments). b. BLI sensorgrams showing GST–AR-LBD binding to p300 with 10 μM DALTAC-1 (apparent Kd, 333.1 nM) but not with DMSO (three independent experiments). c. Schematic and dose–response curves of AR–p300 ternary-complex formation induced by indicated compounds (mean ± s.d.; n=4; three independent experiments). d. Co-IP showing increased AR–p300 interaction in VCaP cells treated with DALTAC-1 (100 nM, 4 h, three independent experiments). e. Top 30 DALTAC-1-enriched p300 interactors identified by IP–MS in VCaP cells (AR, red; n □= □3). Two-sided moderated t-test with Benjamini–Hochberg correction. f. Left, representative AR–p300 PLA images in LNCaP cells treated as indicated for 4 □ h. Scale bar, 10 µm. Right, PLA foci per cell (n □ = □15 cells; three independent experiments). Boxes show median, interquartile range and min–max whiskers. Two-sided t-test. g. Immunoblot analysis of indicated proteins and histone marks in VCaP cells treated with 100 nM indicated compounds for noted durations (three independent experiments). Vinculin, same-gel p300 loading control; H2B, separate-gel control from the same samples. h. Immunoblot analysis of H2BK20ac and H3K27ac in Kelly cells treated with 100 nM indicated compounds for 4 hours (three independent experiments). H2B, separate-gel control from the same samples. i. Volcano plots of lysine acetylation changes following DALTAC-1 or p300i+ARi treatment (100 nM, 12 h; n=3, two-sided unpaired t-tests with Benjamini–Hochberg correction). j. Immunoblot analysis showing AR acetylation in VCaP and LNCaP cells treated with DALTAC-1 or p300i+ARi (100 nM, 4 h, three independent experiments). k. GFP dose–response curves in LNCaP-ARE-GFP cells treated for 48 h. Data are mean ± SD; n=3 (three independent experiments). l. GSEA of AR- and p300-pathway genes ranked by fold change in VCaP cells treated with 100 nM DALTAC-1 for 6 h; adj P, adjusted P value;. n = 2; significance by two-sided GSEA permutation test with multiple-comparison adjustment. m. RNA-seq heatmaps for AR and p300 target genes in VCaP cells treated with DALTAC-1 or p300i+ARi (100 nM, 4 hours). n=2. n. Immunoblot analysis of indicated proteins in VCaP cells treated with 100 nM indicated compounds. H2B, separate-gel control from the same samples (three independent experiments).

To assess intracellular ternary complex formation, we performed a ternary complex ELISA, which showed that DALTAC-1 markedly enhanced AR–p300 association in VCaP cells (**Fig. 2c**). In contrast, the inactive analogs Neg-1 and Neg-2, deficient in binding to p300 or AR, respectively, failed to promote this interaction (**Fig. 2c**). Co-immunoprecipitation (Co-IP) assays further demonstrated that DALTAC-1 robustly enhanced AR–p300/CBP interactions in AR-positive VCaP cells (**Fig. 2d and Extended Data Fig. 3f**). To characterize the DALTAC-1–induced complex, we performed IP–MS analyses in VCaP cells treated with DMSO or DALTAC-1. AR was strongly enriched in p300 immunoprecipitates following DALTAC-1 treatment, and a reciprocal AR IP–MS analysis identified p300 and its paralog CBP among the most enriched associated proteins (**Fig. 2e** and **Extended Data Fig. 3g, h**). A proximity ligation assay (PLA) further confirmed that DALTAC-1 treatment substantially increased the number of AR–p300 PLA foci, indicating enhanced spatial proximity between the two proteins (**Fig. 2f** and **Extended Data Fig. 3i**). This effect was competitively blocked by either a p300 inhibitor or an AR inhibitor, supporting the formation of a DALTAC-1-induced AR–p300 ternary complex (**Fig. 2f** and **Extended Data Fig. 3i**). Sequential co-IP of co-expressed Flag-AR, HA-AR, and V5-p300 showed that DALTAC-1 increased the recovery of HA-AR with Flag-AR from p300-containing complexes, indicating the incorporation of multiple AR molecules into the DALTAC-1-induced complex and supporting AR dimerization within the complex (**Extended Data Fig. 3j**). Consistent with an AR LBD-dependent mechanism, DALTAC-1 markedly enhanced p300 association with full-length AR in 22Rv1 cells but did not increase its association with the LBD-deficient AR-V7 splice variant (**Extended Data Fig. 3k**). DALTAC-1 also retained the ability to enhance p300 association with the DNA-binding-deficient AR-C619Y mutant^35^ (**Extended Data Fig. 3l**), indicating that DALTAC-1-induced complex formation did not require AR binding to DNA. Together, these findings support direct, LBD-dependent AR–p300 complex formation.

Domain-resolved interaction mapping was next performed in HEK293 cells to define the domain contacts underlying DALTAC-1-induced AR–p300 complex formation. Under basal conditions, little to no interaction was detected between AR and the p300 core region, which contains the bromodomain (BRD), RING/PHD domain, and histone acetyltransferase (HAT) domain (**Extended Data Fig. 3m**). In contrast, DALTAC-1 induced a strong interaction between full-length AR and the p300 core region (**Extended Data Fig. 3m**). Reciprocal mapping showed that DALTAC-1 specifically enhanced the interaction of the AR LBD with p300, without increasing interactions mediated by the AR NTD (**Extended Data Fig. 3n**). Direct co-expression of the AR LBD and p300 core region further demonstrated minimal detectable association in the absence of DALTAC-1 and marked complex formation following DALTAC-1 treatment (**Extended Data Fig. 3o**). Thus, DALTAC-1 establishes a non-native interaction between the AR LBD and the p300 core region, imposing a distinct domain-level configuration on the AR–p300 complex.

To gain structural insights into its mechanism of action, we conducted a computational modeling of the AR-LBD/DALTAC-1/p300-BRD ternary complex, as well as of AR-LBD/DALTAC-1 and p300-BRD/DALTAC-1 binary complexes. The top-scoring ternary complex and binary complexes were subjected to a 1 µs molecular dynamics (MD) simulation. In addition to those interactions observed for ARi and p300i for their respective proteins, the predicted AR-DALTAC-1-p300 ternary complex revealed a well-defined interface between AR and p300, stabilized by a network of polar and hydrophobic interactions (**Extended Data Fig. 3p**). Specifically, two strong salt bridges between p300 Asp1088 and AR Arg871, and between p300 Arg1133 and AR Glu872, complemented by a hydrogen bond between Arg871 (AR) and the backbone carbonyl of Gly1085 (p300). A second key contact involved a backbone–backbone hydrogen bond between Ile816 (AR) and Leu1083 (p300), further supported by hydrophobic packing between their side chains. Notably, a cation–π interaction was detected between His874 (AR) and the piperazine moiety of DALTAC-1 (**Extended Data Fig. 3p**). MM-GBSA analyses of the MD trajectories showed that the DALTAC-1-bound ternary complex had a more favorable calculated model-dependent interaction-energy estimate (−161.8 ± 0.6 kcal/mol) than the DALTAC-1/AR binary complex (−81.4 ± 0.6 kcal/mol) and DALTAC-1/p300 binary complex (−90.3 ± 0.4 kcal/mol) (**Extended Data Fig. 3q, r**). Furthermore, when DALTAC-1 was computationally removed from the ternary complex, the calculated MM-GBSA interaction-energy estimate for the resulting AR-p300 interphase was −53.5 ± 1.1 kcal/mol, consistent with favorable protein-protein complementarity within the predicted ternary-complex model (**Extended Data Fig. 3q, r**).

To experimentally validate the predicted model, we generated interface-disrupting AR-R871A/E872A and p300-D1088A/R1133A mutants. HA-tagged wild-type or mutant AR was co-expressed with Flag-tagged wild-type or mutant p300 in HEK293 cells, followed by DALTAC-1 treatment and co-immunoprecipitation analysis. DALTAC-1 markedly enhanced the interaction between wild-type AR and wild-type p300, whereas mutation of the predicted interface residues substantially impaired DALTAC-1-induced complex formation despite comparable expression of the wild-type and mutant proteins (**Extended Data Fig. 3s**). Taken together, our data establish that DALTAC-1 induces a stable AR–p300 ternary complex and rewires their domain-level interactions, providing the mechanistic foundation for its downstream epigenetic and transcriptional effects.

### DALTAC-1 reprograms AR+ PCa acetylome

In contrast to the p300/CBP dual degrader CBPD-409, DALTAC-1 did not reduce p300/CBP protein abundance; instead, it rapidly and robustly suppressed enhancer-associated histone acetylation marks, including H3K27ac and H2BNTac (H2BK20ac) (**Fig. 2g and Extended Data Fig. 4a-c**). Notably, H2BNTac is a key chromatin mark of active enhancers in prostate cancer cells and is exclusively catalyzed by p300/CBP through BRD-independent, HAT activities^8^. Our data demonstrate that DALTAC-1 exerts a *de novo* heterobifunctional effect by potently suppressing this critical histone mark without degrading p300/CBP. This level of H2BNTac suppression was not observed with the inactive analogs Neg-1 and Neg-2, nor with the combination of p300/CBP and AR inhibitors (**Extended Data Fig. 4a–c**). In AR-negative, p300-dependent Kelly neuroblastoma cells, DALTAC-1 only modestly reduced H2BK20ac levels, to an extent comparable to combined p300/CBPi and ARi but markedly less than CBPD-409, further supporting its AR-lineage selectivity (**Fig. 2h**). Furthermore, in HEK293 cells, the degree of H2BNTac reduction by DALTAC-1 scaled proportionally with the level of ectopically expressed AR, reinforcing that both AR and p300 are required to mediate DALTAC activity (**Extended Data Fig. 4d**). Consistent with its durable pharmacologic effect, a 2-hour pulse treatment of DALTAC-1 followed by compound washout resulted in robust inhibition of histone acetylation and MYC protein expression, whereas washout of the p300/CBP and AR inhibitors markedly attenuated their combination activity (**Extended Data Fig. 4e**).

Beyond histone substrates, p300/CBP acetylate a range of non-histone proteins that regulate transcriptional complexes and signal transduction^36^. Previous studies using p300/CBP HAT inhibitors or PROTAC degraders demonstrated that acute p300/CBP inhibition results in broad suppression of the cellular acetylome, reducing acetylation on both histone and non-histone targets^7,8^. Acetyl-lysine mass spectrometry (acetyl-K MS) profiling in VCaP cells revealed that DALTAC-1, unlike conventional p300/CBP inhibition, exerted a distinct acetylation signature. DALTAC-1 markedly suppressed global histone acetylation, most prominently the enhancer-associated H2BNTac mark, surpassing the effect of combined p300/CBP and AR inhibitors (**Fig. 2i** and **Extended Data Fig. 4f, g**). Consistent with this observation, immunoprecipitation-mass spectrometry analysis demonstrated that H2B was the most significantly displaced p300-interacting protein upon DALTAC-1 treatment (**Extended Data Fig. 3g**), indicating a selective disruption of p300’s association with its canonical histone substrate.

Unexpectedly, DALTAC-1 simultaneously induced specific non-histone acetylation events, particularly within components of the AR transcriptional complex. Prominent increases in acetylation were observed in AR as well as its cofactors SRC2, SRC3, ARID1A, and ARID1B^37^ (**Fig. 2i, j** and **Extended Data Fig. 4g, h**). These acetylation sites were located within the NTD of AR, suggesting that the robust ternary complex formation between AR and p300 redirects p300’s catalytic activity toward the AR-NTD and AR-associated cofactors. An androgen response element (ARE) reporter assay assessing the five SRC2 lysines most strongly acetylated following DALTAC-1 treatment showed that an acetylation-mimetic SRC2 mutant modestly enhanced AR transcriptional activity, whereas an acetylation-deficient mutant modestly reduced activity (**Extended Data Fig. 4i, j**). These findings indicate that DALTAC-1-induced SRC2 acetylation has only a limited effect on SRC2 coactivator function. Importantly, such acetylation increases were not observed in cells treated with the combination of the p300/CBP and AR inhibitors (**Fig. 2i, j** and **Extended Data Fig. 4h**). These combined data demonstrate that DALTAC-1 induces an unprecedented acetylation pattern in AR-positive prostate cancer cells, characterized by strong suppression of enhancer-linked histone acetylation and concurrent increases in acetylation of AR and its associated cofactors.

### DALTAC-1 blocks oncogenic transcription

Given the distinct reprogramming of p300 substrate engagement by DALTAC-1, we examined its impact on transcriptional output. An ARE reporter assay demonstrated that DALTAC-1 potently suppressed AR transcriptional activity in LNCaP cells, producing a markedly greater reduction in ARE–GFP reporter signal compared with Neg-1 and Neg-2 (**Fig. 2k** and **Extended Data Fig. 5a**). Consistently, qPCR analysis showed strong repression of canonical AR target genes by DALTAC-1, whereas the combination of p300/CBP and AR inhibitors elicited only moderate effects (**Extended Data Fig. 5b**). To further investigate the global transcriptional effects of DALTAC-1, we performed RNA sequencing (RNA-seq) in VCaP cells. Gene set enrichment analysis (GSEA) revealed that DALTAC-1 profoundly downregulated pathways associated with androgen response, p300 targets, MYC targets, and MTORC1 signaling (**Fig. 2l, m** and **Extended Data Fig. 5c-e**). Chromatin immunoprecipitation followed by sequencing (ChIP–seq) further demonstrated that DALTAC-1 rapidly displaced RNA polymerase II from AR and p300 target loci within hours of treatment, signifying prompt transcriptional shutdown (**Extended Data Fig. 5f**). These data demonstrate that, unlike the transcriptional activation mechanism of TCIPs^18,19^, DALTAC-1 preferentially represses oncogenic transcriptional programs driven by AR and p300/CBP.

At the protein level, both DALTAC-1 and the p300/CBP degrader CBPD-409 markedly reduced PSA, CCND1, and NKX3-1 expression, whereas the combination of p300/CBP and AR inhibitors produced only modest effects (**Fig. 2n**). Quantitative proteomic profiling further confirmed that DALTAC-1 selectively downregulated AR and MYC-regulated proteins without altering the abundance of AR or p300/CBP themselves (**Extended Data Fig. 5g**). Combined, these findings demonstrate that DALTAC-1 reprograms p300 substrate utilization and enforces broad suppression of AR-driven oncogenic transcriptional programs in prostate cancer cells.

We next examined whether the transcriptional inhibitory activity of AR–p300/CBP DALTACs requires an antagonist AR-binding moiety. To this end, we generated a new AR-p300/CBP DALTAC, JZY4055, by replacing the AR antagonist moiety in DALTAC-1 with an enobosarm^38^-derived non-steroidal AR agonist moiety (**Extended Data Fig. 5h**). Despite incorporating an agonist-derived AR ligand, JZY4055 potently inhibited the growth of AR-positive VCaP cells (**Extended Data Fig. 5i**). Whereas enobosarm activated canonical AR target gene expression, JZY4055 suppressed AR target genes and reduced p300/CBP-dependent histone acetylation marks, including H2BK20ac and H3K27ac, together with the AR-and p300-regulated proteins NKX3-1 and MYC (**Extended Data Fig. 5j, k**). These findings indicate that the transcriptional inhibitory activity of AR–p300/CBP DALTACs is not restricted to molecules containing an antagonist-derived AR ligand.

### DALTAC-1 reshapes AR–p300 enhanceosome

p300/CBP serve as central coactivators that bind to active AR enhancer sites and catalyze histone acetylation to sustain oncogenic transcriptional programs in prostate cancer cells^8^. To investigate how DALTAC-1 modulates p300 chromatin occupancy, we performed p300 ChIP–seq in VCaP cells. DALTAC-1 treatment led to a striking expansion of p300-binding events genome-wide (**Fig. 3a**). Genomic annotation revealed distinct redistribution patterns of p300 chromatin binding upon DALTAC-1 treatment. Among the DALTAC-1–depleted peaks, approximately 40% were located at promoter-proximal regions (**Fig. 3b**). In contrast, the DALTAC-1–gained peaks were predominantly mapped to intronic and distal intergenic regions, with only about 10% residing near promoters (**Fig. 3b**), indicating increased engagement of enhancer-associated chromatin. Since DALTAC-1 enforces AR–p300/CBP proximity, we next evaluated how p300 occupancy changed at AR-binding sites. At AR-binding enhancer sites identified under DMSO (control) conditions, DALTAC-1 induced a marked increase in AR occupancy, accompanied by a proportional enhancement in p300 occupancy (**Fig. 3c** and **Extended Data Fig. 6a**). CBP ChIP–seq similarly revealed a marked increase in CBP occupancy at AR-bound enhancers following DALTAC-1 treatment (**Extended Data Fig. 6b**). This reciprocal recruitment pattern suggests that DALTAC-1 acts as a non-degrader molecular glue, stabilizing chromatin-anchored AR–p300 complexes. Motif analysis revealed that DALTAC-1–induced p300 and CBP peaks were most strongly enriched for canonical palindromic AREs and closely related nuclear receptor response elements with near identical consensus sequences, including progesterone response elements (PREs) and glucocorticoid response elements (GREs)^39^ (**Fig. 3d** and **Extended Data Fig. 6c, d**). Together, these findings demonstrate that DALTAC-1 amplifies and redistributes both p300 and CBP enhancer occupancy through AR-dependent recruitment, consistent with DALTAC-1–induced alteration of domain–domain connectivity within AR and p300/CBP that stabilizes a neomorphic ternary complex on enhancer chromatin.

**Figure 3.**
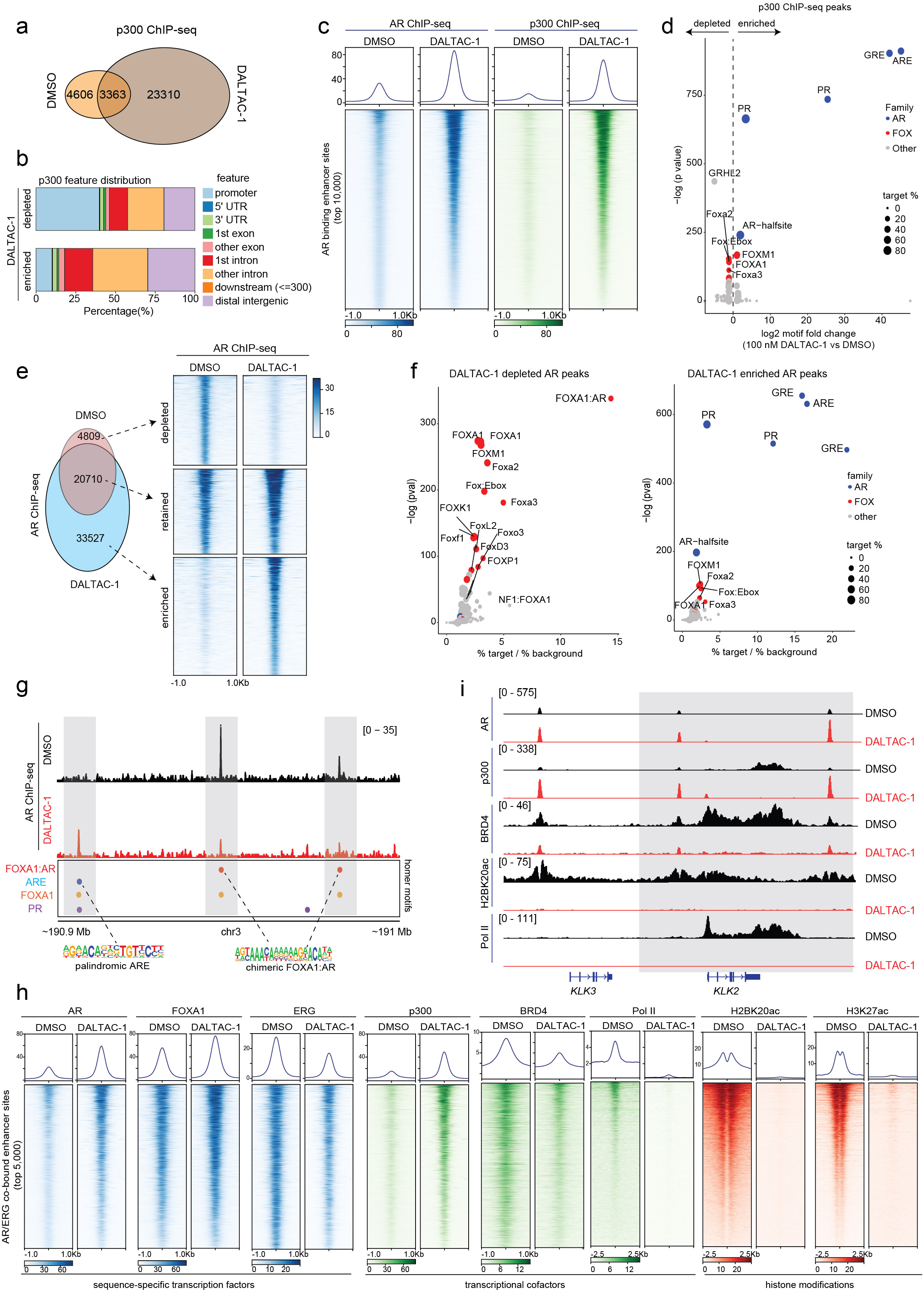
DALTAC-1 reshapes the AR–p300/CBP cistrome and remodels the oncogenic enhancer landscape. a. Venn diagram of genome-wide p300 ChIP-seq peaks in VCaP cells treated with DMSO or 100 nM DALTAC-1 for 3 hours. b. Genomic distribution of DALTAC-1-depleted and -enriched p300 peaks in VCaP cells treated with DMSO or 100 nM DALTAC-1 for 3 hours. c. ChIP-seq read-density heatmaps of AR and p300 at AR-bound enhancer sites in VCaP cells treated with DMSO or 100 nM DALTAC-1 for 3 hours (n=1). d. Fold change and significance of HOMER motifs at non-promoter p300-binding sites in VCaP cells treated with 100 nM DALTAC-1 versus DMSO for 3 h. AR, FOX, and other motifs are blue, red, and grey, respectively. P values were calculated using HOMER’s cumulative binomial test. e. Left, overlap of genome-wide AR ChIP–seq peaks in VCaP cells treated with DMSO or 100 nM DALTAC-1 for 3 h. Right, read-density heatmaps showing DALTAC-1-depleted, retained, and enriched AR-binding sites (n=1). f. HOMER motifs enriched at non-promoter AR sites depleted (left) or gained (right) after 3 h of 100 nM DALTAC-1 treatment in VCaP cells. Target/background frequencies and cumulative binomial P values are shown. AR-, FOX-, and other motifs are blue, red, and grey, respectively. g. AR ChIP–seq tracks on chromosome 3 (190.9–191.0 Mb) in VCaP cells treated with 100 nM DALTAC-1 or DMSO for 3 hours. ARE, FOXA1-AR, and FOXA1 motifs are highlighted as indicated. h. ChIP–seq heatmaps of the indicated factors and histone marks at the top 5,000 non-promoter AR–ERG co-bound sites in VCaP cells treated with DMSO or 100 nM DALTAC-1 for 3 h (n=1). i. ChIP–seq tracks of the indicated proteins and histone marks at the *KLK3*–*KLK2* locus in VCaP cells treated with DMSO or 100 nM DALTAC-1 for 3 h.

We next investigated how DALTAC-1 reshapes the AR cistrome. In accordance with the strengthened AR–p300 interaction, DALTAC-1 markedly increased AR occupancy at palindromic AREs (**Fig. 3e, f** and **Extended Data Fig. 6e, f**). Unexpectedly, DALTAC-1 simultaneously reduced AR binding at FOXA1-AR chimeric motifs, the non-canonical ARE configurations previously reported to be associated with the AR neo-enhanceosome^40^ (**Fig. 3f, g** and **Extended Data Fig. 6f, g**). Thus, instead of uniformly increasing AR binding, DALTAC-1 redirected AR away from chimeric oncogenic enhancers and concentrated it at canonical AREs and other canonical nuclear receptor response elements (e.g., PRE and GRE), effectively remodeling the AR enhancer network. Although ARE, PRE, and GRE motifs share substantial sequence similarity^39^, we sought to exclude the possibility that their enrichment reflected off-target activation of alternative steroid receptor pathways. We, therefore, profiled progesterone receptor (PR) chromatin occupancy as a representative example. PR ChIP–seq revealed that DALTAC-1 markedly reduced PR binding genome-wide, in contrast to the increased AR occupancy observed following treatment (**Extended Data Fig. 6h, i**). Notably, at regions occupied by PR under DMSO conditions, DALTAC-1 decreased PR binding while increasing AR occupancy (**Extended Data Fig. 6i**). These findings indicate that motifs annotated as PRE predominantly represent shared binding sites engaged by AR, rather than activation of PR-dependent chromatin binding. In line with this selective redistribution, DALTAC-1 led to a profound loss in coactivator (BRD4, ERG) recruitment, RNA polymerase II loading, and histone acetylation (H3K27ac and H2BK20ac) at AR/p300 co-bound enhancers (**Extended Data Fig. 6j, k**), demonstrating that DALTAC-1 functionally decommissioned the active AR transcriptional complex.

Oncogenic transcription factors such as ERG and FOXA1 mutants are known to hijack AR to non-canonical AREs, forming cancer-specific AR neo-enhanceosomes that drive lineage-dependent oncogene programs^8,41^. To evaluate how DALTAC-1 influences this architecture, we examined its effects on AR/ERG neo-enhanceosomes in *TMPRSS2–ERG* fusion positive VCaP cells^42^. DALTAC-1 globally suppressed ERG chromatin occupancy (**Extended Data Fig. 7a, b**). Interestingly, a subset of ERG-binding peaks emerged following treatment, predominantly within distal intergenic and intronic regions (**Extended Data Fig. 7b-d**). Furthermore, motif analysis revealed that DALTAC-1 depleted ERG binding at canonical ETS motifs while redirecting a fraction of ERG binding toward palindromic AR motifs (**Extended Data Fig. 7e**), suggesting that ERG can be shifted from its native ETS-binding sites and tethered to AR-dominant enhancers within the reconfigured complex.

We next analyzed the chromatin state at the AR/ERG co-bound non-promoter regions, which represent the AR/ERG neo-enhancers (**Extended Data Fig. 7f**). DALTAC-1 markedly increased AR and p300 occupancy at these loci (**Fig. 3h**), consistent with its ability to stabilize AR–p300/CBP interactions. We also examined whether other AR-associated cofactors were similarly altered. FOXA1 occupancy was largely maintained, with only a slight increase, whereas SRC2, identified by acetylome analysis as the p160 family member most prominently acetylated following DALTAC-1 treatment, showed a modest increase in occupancy at these regions (**Fig. 3h** and **Extended Data Fig. 7g**). In stark contrast, these same regions became transcriptionally inert: ERG and BRD4 binding were diminished, and, strikingly, RNA polymerase II occupancy and enhancer-associated histone acetylation marks were almost completely extinguished (**Fig. 3h**). The pronounced increase in AR/p300 occupancy alongside the sharp loss of ERG, BRD4, RNA polymerase II, and histone acetylation marks demonstrates that DALTAC-1 collapsed AR–ERG neo-enhancers into an inactive epigenetic state, thereby suppressing ERG engagement and dismantling the transcriptional output of the AR–ERG neo-enhanceosome (**Fig. 3i** and **Extended Data Fig. 7h**).

### DALTAC-1 suppresses PCa organoid growth

To assess the therapeutic efficacy of DALTAC-1, we interrogated distinct molecular subtypes of prostate cancer represented by organoids generated from genetically engineered mouse models (GEMMs). We treated organoids derived from the prostate tissues of wild-type, *Rosa26^ERG^/Pten^-/-^*, *Pten*^-/-^, *FOXA1 R265–71del^+/+^/Trp53*^-/-^ (class1 FOXA1 mutant), and *Trp53*^-/-^ transgenic animals with DALTAC-1^41,43^. Compared to wild-type, *Pten*^-/-^, and *Trp53*^-/-^ organoids, DALTAC-1 markedly suppressed the growth of *Rosa26^ERG^/Pten^-/-^*and *FOXA1 R265–71del^+/+^/Trp53*^-/-^ organoids, leading to a significant reduction in both organoid number and size (**Fig. 4a, b** and **Extended Data Fig. 8a**). Correspondingly, DALTAC-1 treatment potently downregulated key oncogenes, including MYC and CCND1, in these lineage-defined organoids relative to *Pten*^-/-^ and *Trp53*^-/-^ controls (**Extended Data Fig. 8b**). Given that ERG and mutant FOXA1 enhance AR activity and sustain the AR neo-enhanceosome^5,41,43,44^, these findings suggest that DALTAC-1 effectively suppresses the AR neo-enhanceosome driving prostate cancer.

**Figure 4.**
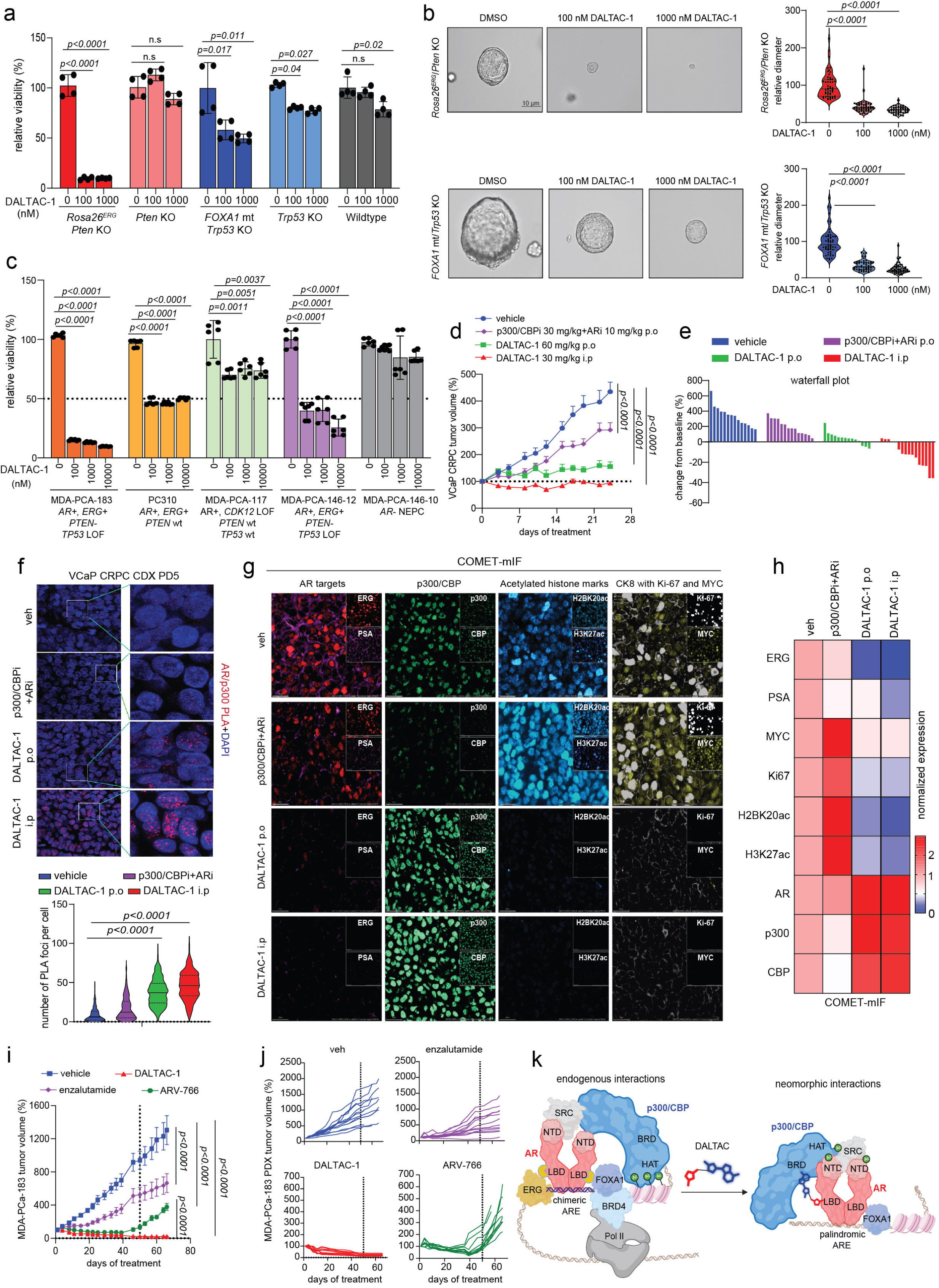
DALTAC-1 potently suppresses prostate cancer growth in preclinical models. a. Viability of GEMM-derived mouse organoids treated with DMSO or DALTAC-1 (100 or 1,000□nM) for 7□d (mean□±□s.d.; n□=□4; two-sided t-test performed separately for each model, three independent experiments). b. Representative images and relative diameters of the indicated organoids treated as in Fig. 4a (n□=□50; two-sided t-test, three independent experiments). Scale bar, 10□μm. c. Viability of PDX-derived human organoids treated with DMSO or DALTAC-1 (100, 1,000 or 10,000□nM) for 14□d (n□=□6; two-sided t-test performed separately for each model, three independent experiments). d. VCaP-CRPC tumor volumes after vehicle, DALTAC-1 (30□mg/kg i.p. or 60□mg/kg p.o.; 5× weekly in week 1, then 3× weekly for 3 weeks), or p300/CBPi□+□ARi (30□+□10□mg/kg p.o., 5× weekly). Data are mean□±□s.e.m.; n□=□14 mice for vehicle, oral DALTAC-1, and p300/CBPi□+□ARi, and n=16 for i.p. DALTAC-1. Endpoint volumes were analyzed by two-sided t-test. e. Individual tumor-volume changes after 24□d in Fig. 4d. f. Representative AR–p300 PLA images and foci per cell in day-5 tumors (n□=□200 cells from each group; two-sided t-test). Center line, median; dotted lines, quartiles; violin, kernel-density estimate; whiskers, range. Scale bar, 20□μm. g. Representative COMET-mIF merged images of indicated targets from PD5 tumors in the VCaP-CRPC study. Scale bar, 20 µm. h. Heatmap showing normalized median expression levels of the indicated targets from Fig. 4g. i. MDA-PCa-183 PDX growth during treatment with vehicle, DALTAC-1 (30□mg/kg i.p., 3× weekly), ARV-766 (10□mg/kg p.o., 5× weekly), or enzalutamide (10□mg/kg p.o., 5× weekly). Dosing stopped on day 50, followed by 16□d of monitoring (mean□±□s.e.m.; n□=□12 mice for vehicle, DALTAC-1, and enzalutamide and n□=□11 for ARV-766; two-sided t-test). j. Graph depicting the tumor volume curves of individual tumors from Fig. 4i. k. Model of DALTAC-1-mediated AR–p300/CBP rewiring, coactivator displacement, loss of histone acetylation, and AR neo-enhanceosome collapse.

To further evaluate DALTAC-1 activity in human prostate cancer models, we treated a panel of organoids derived from patient-derived xenograft (PDX) tumors with DALTAC-1^45,46^. Notably, DALTAC-1 exhibited robust growth inhibitory effects in AR-positive organoids, particularly *AR*⁺/*ERG*⁺, aligning with its potent blockade of the AR neo-enhanceosome (**Fig. 4c**). In contrast, AR-negative neuroendocrine prostate cancer (NEPC) organoids displayed minimal response, consistent with the AR-dependent mechanism of action of DALTAC-1 (**Fig. 4c**). These results establish DALTAC-1 as a potent suppressor of AR-driven prostate cancer growth across both murine and human organoid models, highlighting its broad yet lineage-selective anti-tumor activity.

### Potent in vivo efficacy of DALTAC-1 in PCa

To evaluate the anti-tumor activity of DALTAC-1, we employed a VCaP castration-resistant prostate cancer (VCaP-CRPC) xenograft model established by subcutaneous implantation of VCaP cells into castrated male SCID mice (**Extended Data Fig. 8c**). Once tumors reached ∼150 mm³, mice were randomized and treated with DALTAC-1 via intraperitoneal (ip) injection (30 mg/kg) or oral administration (po) (60 mg/kg). For comparison, a combination of a p300/CBP bromodomain inhibitor and an AR-LBD inhibitor was administered at equipotent doses. IP administration of DALTAC-1 as a single agent elicited markedly superior anti-tumor activity, inducing tumor regression in 75% (12 of 16) of treated tumors (**Fig. 4d, e** and **Extended Data Fig. 8d, e**). Oral dosing of DALTAC-1 also achieved robust tumor suppression and outperformed the combination of the p300/CBP and AR inhibitors, underscoring its favorable translational properties (**Fig. 4d, e** and **Extended Data Fig. 8d, e**). Mechanistically, in vivo PLA revealed a marked increase in AR–p300 proximity in DALTAC-1–treated tumors, supporting induced formation of the AR–p300–DALTAC-1 ternary complex in vivo (**Fig. 4f**). Consistently, COMET multiplex immunofluorescence (COMET-mIF) analyses of VCaP-CRPC tumors collected after five days of treatment of DALTAC-1 revealed marked reductions in ERG, PSA, MYC, Ki67, H3K27ac, and H2BK20ac signals, while AR, p300, and CBP protein levels did not decrease, consistent with its mechanism of action (**Fig. 4g, h** and **Extended Data Fig. 8f**).

To extend these findings to clinically relevant models, we assessed the efficacy of DALTAC-1 in two PDX models, MDA-PCa-146-12 and MDA-PCa-183^45^. Consistent with the lineage-selective activity observed in organoids, DALTAC-1 treatment significantly inhibited tumor growth in MDA-PCa-146-12 (**Extended Data Fig. 8g**). In an initial efficacy study, DALTAC-1 induced rapid tumor shrinkage and substantial regress on of all tumors in the MDA-PCa-183 model, which harbors a *TMPRSS2-ERG* fusion and *PTEN* loss^43,47^, with no progression observed during the 35-day treatment period (**Extended Data Fig. 8h-k**). Given this pronounced response, we next benchmarked DALTAC-1 against two distinct AR-directed therapies in MDA-PCa-183. DALTAC-1 induced sustained and complete tumor regression during the 50-day treatment period (**Fig. 4i, j**). In contrast, the clinical AR antagonist enzalutamide produced only modest tumor growth inhibition, whereas the AR degrader ARV-766 showed substantial anti-tumor activity but was less efficacious than DALTAC-1 (**Fig. 4i, j**). To assess response durability, treatment was discontinued on day 50 and tumors were monitored for an additional 16 days. No evident tumor recurrence was observed in the DALTAC-1-treated group, whereas tumors in the ARV-766-treated group showed marked regrowth following treatment withdrawal (**Fig. 4i, j**). Importantly, throughout all the studies, DALTAC-1 was well tolerated, with no significant changes in body weight, serum liver or kidney function, or hematologic parameters (**Extended Data Fig. 9a-e**). Together, these results demonstrate that DALTAC-1 is an effective and well-tolerated anti-tumor agent that exerts durable efficacy across multiple prostate cancer models through selective AR–p300 domain alterations and chromatin reprogramming, establishing it as a promising first-in-class therapeutic with strong potential for clinical translation.

## Discussion

Recent advances in chemical-induced proximity (CIP) have led to the development of a diverse array of compounds capable of modulating protein functions through enforced protein–protein interactions^15^. RIPTACs induce proximity between a target protein and a pan-essential effector that disrupts the essential effector’s function, leading to selective cancer cell death while sparing healthy cells^17^. In contrast, TCIPs recruit transcriptional coactivators to specific genomic loci, paradoxically activating the target pathway by enhancing transcription of pro-apoptotic genes normally repressed by corepressors such as BCL6^18,19^. The emerging clinical activity of daraxonrasib (RMC-6236), which forms a tri-complex between cyclophilin A and active RAS, further supports the translational potential of proximity-induced pharmacology^48^. Recent molecular glue degraders further demonstrate that small molecules can stabilize pre-existing or newly enabled protein interfaces and alter target organization to promote E3-ligase-dependent degradation^49^. Complementary studies targeting FOXA1, SWI/SNF, and p300 establish that DNA-binding specificity, ATPase cycling, and chromatin engagement are also pharmacologically actionable^50–52^. DALTACs build on these principles but produce a distinct pharmacological outcome: rather than recruiting an E3 ligase, degrading a complex component, or modulating an individual regulator, a DALTAC bifunctional molecule imposes an additional domain-selective interaction between two members within an existing transcriptional complex, converting the complex into a non-productive state. Our proof-of-concept compound, AR–p300/CBP DALTAC-1, targets the AR–p300/CBP complex by repurposing their natural interfaces to block activation of the AR oncogenic enhanceosome. Mechanistically, AR–p300/CBP DALTAC-1 potently suppresses AR transactivation by disrupting the chromatin landscape at AR-bound enhancer sites, particularly the AR/ERG co-bound neo-enhancer sites, as evidenced by reduced cofactor recruitment, loss of histone acetylation marks, and displacement of RNA polymerase II. Functionally, this shutdown of AR neo-enhanceosome activity translates into strong anti-tumor efficacy in AR-positive prostate cancer models, underscoring the translational potential of DALTAC-1 as a precision therapeutic.

The activity of CIP molecules is founded on formation of a ternary complex between the recruited proteins. Using purified full-length proteins, we showed that DALTAC-1 directly enhances AR–p300 complex formation in the absence of DNA or accessory factors. This association requires simultaneous engagement of both proteins by the intact bifunctional molecule and cannot be reproduced by unlinked ligands or control compounds lacking either AR- or p300-binding activity. The activities of multi-protein transcriptional complexes are governed by finely tuned protein–protein interactions (PPIs) at the sub-domain level^53^. AR normally recruits transcriptional coactivators primarily through its intrinsically disordered NTD, whereas transcription factors engage distinct regions of p300, including its N- and C-terminal regions, to support histone acetylation and transcriptional activation^2,13^. Our domain-mapping and interface-mutagenesis studies support a refined model in which DALTAC-1 does not necessarily abolish these native contacts. Rather, it imposes an additional, neomorphic interaction between the AR LBD and the p300 core region containing the BRD and HAT domain, thereby reorganizing the domain-level engagement within the retained AR–p300 complex. Although the atomic architecture of the ternary complex remains unresolved, owing in part to the large size, conformational flexibility, and extensive intrinsically disordered regions of full-length AR and p300, our orthogonal biochemical and cellular data demonstrate a DALTAC-induced binding pattern that is distinct from the basal complex and functionally alters AR–p300/CBP activity. This interaction reprogramming markedly suppresses histone acetylation, including H2BNTac, despite the inability of the p300-binding moiety alone to directly inhibit this modification^8^. In parallel, DALTAC-1 induces acetylation of the AR-NTD and core components of the AR complex, including SRC2/3 and ARID1A/B^4,37,54^. Because p300 has previously been shown to acetylate AR primarily within the hinge region under androgen stimulation^55^, the emergence of AR-NTD acetylation suggests that DALTAC-1 redirects p300 substrate engagement, potentially by bringing the AR NTD into altered proximity to the p300 HAT domain. This substrate reprogramming distinguishes DALTAC-1 from conventional p300 inhibitors and degraders: unlike BRD inhibitors, it extinguishes enhancer-associated H2BNT acetylation and broadly represses transcription, while unlike PROTAC degraders, it acts independently of ubiquitin–proteasome-mediated target depletion.

Because many oncogenes also play essential roles in normal physiology, their pan-inhibition often causes severe side effects. For example, systemic inhibition of p300/CBP disrupts hematopoiesis and leads to thrombocytopenia^10^. Thus, achieving lineage-selective targeting of oncogenic pathways is an urgent therapeutic need. Existing approaches such as antibody–drug conjugates (ADCs) deliver cytotoxic payloads selectively to cells expressing target surface antigens^56^. However, their broader applicability is limited by the restricted repertoire of tumor-specific surface markers and, in many cases, by suboptimal pharmacokinetic properties^57^,. By contrast, DALTACs offer a fundamentally distinct strategy, selectively rewiring lineage-specific protein complexes that drive oncogenic transcription. In this study, AR– p300/CBP DALTAC-1 specifically inhibits the growth of AR-positive prostate cancer cells with potency comparable to or better than the p300/CBP degrader CBPD-409, but shows dramatically reduced activity in p300/CBP-dependent, AR-negative cancers such as neuroblastoma, Ewing sarcoma, and hematologic malignancies^8^. Notably, in AR-negative cells, DALTAC-1 fails to suppress H2BNTac, underscoring that DALTAC-1–mediated inhibition of p300/CBP histone acetylation is context-dependent and conferred by domain reprogramming within AR-p300/CBP complexes.

In summary, we establish DALTACs as a new type of chemical proximity modulators that rewire domain-specific protein interactions to selectively deactivate oncogenic complexes. DALTAC-1 induces formation of an AR-p300-DALTAC-1 ternary complex and reprograms the substrate engagement of p300/CBP, markedly suppressing global histone acetylation while enhancing the acetylation of core components within the AR complex. Consequently, DALTAC-1 reinforces AR-p300/CBP co-occupancy at palindromic ARE loci but disrupts the higher-order assembly of the AR complex and erases histone activation marks at AR neo-enhanceosomes (**Fig. 4k**). This work expands the conceptual framework of induced-proximity therapeutics beyond degradation and coactivator hijacking, paving the way for the rational design of domain-alteration strategies for precise control of disease-relevant protein complexes.

## Acknowledgements

We thank Lanbo Xiao, Amanda Miller, Christine Caldwell-Smith, Shuqing Li, and Nidhi Mistry from the Michigan Center for Translational Pathology at the University of Michigan as well as V. Basrur and the Rogel Cancer Center Proteomics Shared Resource for providing technical assistance. This work was supported by the following mechanisms: Prostate Cancer Foundation 2023 Challenge Award (to A.M.C., S.W., and A.P.), National Cancer Institute (NCI) Specialized Programs of Research Excellence (P50-CA186786 to A.M.C.), NCI Outstanding Investigator Award (R35-CA231996 to A.M.C.), Prostate Cancer Foundation 2023 Young Investigator Award (to J.L.), Prostate Cancer Foundation 2025 Challenge Award (to A.M.C., S.W., and Y.Q.), Department of War Prostate Cancer Research Program (PCRP) 2025 Idea Development Award (HT9425-26-1-E093 to A.M.C.), NCI R01 Award (R01-CA289013 to S.W.), and a contract from Medsyn Biopharma (to S.W.). A.M.C. is a Howard Hughes Medical Institute Investigator, A. Alfred Taubman Scholar, and American Cancer Society Professor.

## Author contributions

J.L., J.Y., S.W., and A.M.C. conceived and designed the studies; J.L. performed all of the in vitro and functional genomics experiments with assistance from S.D., J.C., Y.L., X.J., X.W., K.G., S.E., and M.W.; Y.Q., J.C.T., and J.L. performed all of the organoid and animal efficacy studies with assistance from S.E., Y.Y., and A.J.T.; E.Y. and S.L. carried out all of the bioinformatics analyses with assistance from A.P.; J.T. and J.Y. performed the computational modeling of the AR–p300–DALTAC-1 complex; R.M. and S.M. carried out all of the histopathological evaluations of drug toxicity as well as quantified all of the histology-based data; R.M. and S.M. carried out all of the immunohistochemistry and immunofluorescence staining; R.S. and A.R. conducted quantification of Seq-mIF datasets; X.C. and F.S. performed sequencing; J.Y., W.X., M.W., S.J., and S.W. designed, synthesized, and performed initial evaluations of a large number of DALTAC molecules, leading to the discovery of DALTAC-1, as well as Neg-1 and Neg-2 control compounds; J.L., J.Y., S.J.M., J.C.T., S.W., and A.M.C. wrote the manuscript and organized the final figures.

## Data availability

All data are available in the manuscript and the supplementary information. Raw sequencing data have been deposited in the Gene Expression Omnibus (GEO) with the accession numbers GSE310948 and GSE310949.

## Code availability

No custom code or software was used in this study. All computational analyses were performed using publicly available software and code, which are described in the Methods section.

## Additional information

Supplementary Information is available for this paper. Correspondence and requests for materials should be addressed to Arul M. Chinnaiyan and Shaomeng Wang. Reprints and permissions information is available at www.nature.com/reprints.

## Methods

### Cell lines

All cell lines used in this study were originally sourced from ATCC, DSMZ, ECACC, Lonza, or maintained as authenticated internal stocks. The CWR-R1 prostate cancer line was kindly provided by Dr. David Vander Griend (University of Illinois at Chicago). Cell line authentication was performed semi-annually through genotyping at the University of Michigan DNA Sequencing Core, and routine Mycoplasma testing was carried out biweekly. LNCaP, 22Rv1, CWR-R1, and DU145 cells were cultured in RPMI-1640 medium supplemented with 10% fetal bovine serum (FBS; Thermo Fisher Scientific). VCaP cells were maintained in DMEM GlutaMAX supplemented with 10% FBS (Thermo Fisher Scientific).

### Chemistry

Detailed synthesis and chemical data for all designed DALTAC molecules and negative controls are provided in the **Supplementary Methods** (Supplementary Information file).

### Antibodies

For immunoblotting, the following antibodies were used: p300 (Invitrogen: MA1-16608); AR (Abcam: ab133273); H2BK20ac (Cell Signaling Technology: 34156S); H2BK16ac (Abcam: ab177427); H3K27ac (Cell Signaling Technology: 8173S); H2B (Active Motif: 39210); H3 (Cell Signaling Technology: 3638S); c-Myc (Cell Signaling Technology: 9402S); c-Myc/N-Myc (Cell Signaling Technology: 13987S); KLK3/PSA (Dako: A0562); NKX3-1 (Cell Signaling Technology:83700S); CCND1 (Abcam: ab16663); Vinculin (Cell Signaling Technology: 18799S); GAPDH (Santa Cruz Biotechnology: sc-47724); BRD4 (Bethyl Laboratories: A700-004CF), HA-Tag (Cell Signaling Technology: 3724S).

For IP and ChIP-seq, the following antibodies were used: H3K27ac (Diagenode: C15410196); p300 (active motif: 61401); H2BK20ac (Cell Signaling Technology: 34156S); AR (Abcam: ab108341); FOXA1 (Thermo Fisher Scientific: PA5-27157); BRD4 (Diagenode: C15410337); ERG (Cell Signaling Technology: 97249S); RNA Pol II (Diagenode: C15200253); acetylated-lysine (Cell Signaling Technology: 9441S); CBP (Abcam: ab253202); SRC2 (Cell Signaling Technology: 96687S); PR (Cell Signaling Technology: 8757S), AR-V7 (Cell Signaling Technology: 19672S)

For immunofluorescent staining (COMET-mIF) and PLA assay, the following antibodies were used: p300 (Invitrogen: MA1-16608); CBP (Invitrogen: PA5-27369); AR (Abcam: ab108341); H3K27ac (Cell Signaling Technology: 8173S); Ki67 (Abcam: ab243878); H2BK20ac (Abcam: ab177430); CK8 (Abcam, ab53280); ERG (Abcam, ab92513); PSA (Cell Marque, 324M-16); MYC (Abcam, ab32072).

For ELISA, the following antibodies were used: p300 (Cell Signaling Technology: 86377S); AR (Thermo Fisher Scientific: MA5-13426).

### Cell viability assay

VCaP cells were seeded at a density of 10,000 cells per well, and all other prostate cancer cell lines were seeded at 3,000 cells per well in poly-D-lysine–coated 96-well plates (Corning) and maintained at 37 °C with 5% CO₂. After 24 hours, cells were treated with a serial dilution of test compounds, with six replicate wells for each concentration. Following 120 hours of incubation, cell viability was determined using the CellTiter-Glo Luminescent Cell Viability Assay (Promega) according to the manufacturer’s instructions. Luminescence was measured on an Infinite M1000 Pro plate reader (Tecan), and data were analyzed using GraphPad Prism (GraphPad Software).

### Incucyte proliferation assays

A total of 4,000 VCaP cells or 1,500 LNCaP cells per well were seeded into poly-D-lysine–coated 96-well plates. After 24 hours of incubation, cells were treated with a fixed concentration (100 nM) or serial dilutions of compounds and incubated for 4 hours. The medium was then removed, and cells were washed three times with PBS before fresh medium containing either vehicle or compounds was added. Phase-contrast images were collected every 4 hours, and cell proliferation was quantified by measuring the percentage of confluence (phase object area).

### Immunoblotting

Cells were lysed in RIPA buffer (Thermo Fisher Scientific) supplemented with 1× Halt™ Protease Inhibitor Cocktail (Thermo Fisher Scientific) and denatured in NuPAGE™ 1× LDS sample buffer containing a reducing agent (Invitrogen) by incubation at 70 °C for 15 minutes. Protein concentrations were determined using the Pierce™ BCA Protein Assay Kit (Thermo Fisher Scientific). Equal amounts of protein (15–30 µg) were resolved on NuPAGE™ 3–8% Tris-Acetate or 4–12% Bis-Tris protein gels (Thermo Fisher Scientific) and transferred onto 0.45-µm nitrocellulose membranes (Thermo Fisher Scientific) using the iBlot™ 3 Dry Blotting System (Thermo Fisher Scientific). Membranes were blocked in TBS-T (Tris-buffered saline with 0.1% Tween-20) containing 5% non-fat dry milk for 1 hour at room temperature and incubated overnight at 4 °C with primary antibodies. After washing, membranes were incubated with HRP-conjugated secondary antibodies and imaged using an Odyssey CLx Imager (LI-COR Biosciences).

### Fluorescence polarization (FP) binding assays

The binding affinities of test compounds to the androgen receptor (AR) and CBP/p300 were measured using fluorescence polarization (FP)–based competitive binding assays. AR binding was assessed using the PolarScreen™ Androgen Receptor Competitor Assay Kit (Thermo Fisher Scientific, Cat. #A15880) following the manufacturer’s instructions.

CBP/p300 FP assays were performed in 96-well plates (MTP96VPWB-EP, Eppendorf) using a CLARIOstar microplate reader (BMG Labtech). Serial dilutions of test compounds were incubated with fluorescein-labeled tracer (2 nM final) and CBP/p300 protein (20 nM final) in assay buffer (PBS pH 7.4 supplemented with 0.01% BGG, 0.01% Tween-20, and 2 mM DTT) to a final reaction volume of 100 µL per well. Plates were mixed for 15 minutes and incubated at room temperature for 1 hour to reach equilibrium. Fluorescence polarization (mP units) was measured at 485 nm excitation and 530 nm emission.

For both AR and CBP/p300 assays, data analysis was performed in GraphPad Prism 10 using nonlinear regression. IC₅₀ values were determined by fitting competition curves, and K□values for competitive inhibitors were derived using nonlinear regression based on the tracer K□ and the concentrations of tracer and protein in the reaction. All FP experiments were conducted in duplicates across three independent biological replicates.

### Cellular thermal shift assay

VCaP or LNCaP cells were seeded in Poly-D-Lysine 100 mm culture dishes (Corning) and incubated overnight. Cells were incubated with the DMSO control or indicated compounds for 2 h at 37 °C before being collected and washed once with PBS. The cell suspension was then distributed into PCR tubes (50 µL per tube) and exposed to a predefined temperature gradient for 3 min using a Veriti Thermal Cycler (Applied Biosystems). Following equilibration to room temperature, cells were lysed by three freeze–thaw cycles in liquid nitrogen. Insoluble protein aggregates were pelleted by centrifugation at 20,000 × g for 20 min at 4 °C, and the resulting supernatants containing soluble proteins were recovered for Western blot analysis.

### Computational modeling

All modelling and simulations were performed with Schrödinger 2025-2 (Schrödinger, LLC). Because co-crystal structures of the androgen receptor (AR) ligand-binding domain (LBD) in an open antagonist conformation were not available, we used our previously reported pseudo-open model^58^. The bound AR ligand was modified to generate JZY3221, the AR ligand used in our design of DALTACs. To model the p300 bromodomain interaction with GNE-049, we used the CBP bromodomain GNE-781 co-crystal structure (PDB 5W0E) as the starting structure^59^. CBP and p300 are highly homologous proteins, especially in their bromodomains which share conserved sequence and structures. Both proteins were prepared for further modeling using the Protein Preparation Wizard^60^.

We employed the following structures as inputs to generate the initial AR:DALTAC-1:p300 complex: (i) the AR in complex with JZY3221, (ii) the p300 complex with GNE-049, and (iii) JZY3032 (DALTAC-1) prepared with LigPrep module^61^. These inputs were submitted to Generate Degrader Complex (default settings: MCMM 10 000 steps, maximum clashes 5, maximum poses 50) to construct the ternary AR:JZY3032:p300 complex. The top 20 predicted complexes were further refined using Prime^62^. The top-scoring ternary complex was then selected for molecular dynamics (MD) simulations.

To perform MD simulations, the predicted ternary complex was solvated in an explicit water box with periodic boundary conditions. A cubic box was constructed with a 10 Å buffer between solute and box edge and filled with TIP4P water^63^. The OPLS4 force field was used for all components. Production dynamics used a RESPA integrator with a 2 fs time step. Temperature was maintained at 300 K with a Nosé–Hoover chain thermostat (relaxation time 1.0 ps)^64^. Pressure was controlled at 1 atm with a Martyna–Tobias–Klein barostat (relaxation time 2.0 ps)^65^. Long-range electrostatics used a 9 Å real-space cutoff with particle–mesh Ewald for reciprocal space^66^. One continuous 1 µs simulation was performed without restraints.

Root-mean-square deviation (RMSD) analysis of protein backbones and ligand position indicated that the system equilibrated after approximately 750 ns. Accordingly, trajectory analysis focused on the last 250 ns. Protein–protein interface features were defined as minimum heavy-atom distances between AR and p300. Contacts shorter than 0.5 nm (5 Å) were retained. Principal component analysis (PCA) was applied to these interface distances, and components explaining 95% of the variance were kept. The reduced data were clustered with HDBSCAN (min_cluster_size 25, centroids enabled)^67^. For each cluster, the frame closest to the centroid in PCA space was saved as a representative PDB. Cluster occupancies are reported as percentages of frames per cluster. Protein–protein interaction fingerprints were computed with Schrödinger’s analyze_trajectory_ppi.py. Clustering in this reduced dimensional space revealed two dominant populations, with the most prevalent state comprising nearly 80% of the ensemble. The interaction analysis was performed on snapshots representative of this dominant state.

### MM-GBSA calculations

MM-GBSA binding energies were estimated using Prime MM-GBSA (Schrödinger Release 2025-2; thermal_mmgbsa protocol) with the OPLS4 force field and the VSGB 2.0 implicit-solvent model^68–70^. Snapshots were sampled evenly from the final 250 ns of each trajectory, after which water molecules and ions were removed and each complex, receptor, and ligand were locally energy-minimized. The MM-GBSA binding energy was calculated as D*G*_bind_ = *E*(complex) - *E*(receptor) - *E*(ligand). For compound binding, DALTAC-1 was defined as the ligand and AR, p300, or AR + p300 as the receptor. For p300 association with the ternary complex, p300 (residues 1049-1160) was defined as the ligand and AR + DALTAC-1 as the receptor. To isolate the direct AR-p300 contribution in the induced geometry, DALTAC-1 was deleted from the same ternary snapshots, with p300 defined as the ligand and AR as the receptor. Snapshots were sampled evenly over the final 250 ns. MM-GBSA values were used as model-dependent interaction-energy estimates to compare the relative stability of the modeled complexes and were not interpreted as experimental binding affinities.

### Full-length AR protein expression and purification

AR protein was expressed according to the manufacturer’s protocol for the Expi293™ PRO Expression System Kit (Thermo Fisher Scientific, Cat. No. A40003643). Briefly, Expi293™ PRO cells were transfected with the HA-AR expression plasmid and cultured for 24 h before the addition of 1 μM R1881. The cultures were then incubated for an additional 24 h, and cells were harvested 48 h post-transfection. Cell pellets were washed and collected by centrifugation at 5,000 rpm for 10 min at 4°C. The cells were resuspended in lysis buffer containing 50 mM Tris-HCl (pH 8.0), 150 mM NaCl, 100 nM R1881, and 0.5% NP-40, supplemented with cOmplete™ EDTA-free Protease Inhibitor Cocktail (Roche, Cat. No. 4693159001). Cells were lysed by sonication, and the lysates were clarified by centrifugation at 15,000 rpm for 40 min at 4°C. The clarified lysate was incubated with anti-HA magnetic beads (Thermo Scientific, Cat. No. 88837) for 3 h at 4°C. After binding, the beads were washed three times with wash buffer containing 50 mM Tris-HCl (pH 8.0), 150 mM NaCl, and 0.05% NP-40. Bound AR protein was eluted twice using elution buffer (50 mM Tris-HCl, pH 8.0, 150 mM NaCl, 100 nM R1881, and 0.05% NP-40) supplemented with 2 mg/mL HA peptide (Thermo Scientific, Cat. No. 26184). The eluted protein was further purified by size-exclusion chromatography using a Superdex 200 Increase 10/300 GL column (GE Healthcare) pre-equilibrated with gel filtration buffer containing 50 mM Tris-HCl (pH 8.0), 150 mM NaCl, 100 nM R1881, and 0.05% NP-40. Peak fractions containing purified AR protein were pooled to a final concentration of approximately 0.5 mg/mL and used for all subsequent studies.

### Purified-protein AR–p300 immunoprecipitation assay

For each reaction, 15 μL of anti-HA magnetic beads (Thermo Fisher Scientific, 88837) was blocked with 0.3% BSA in PBS for 1 h at 4□°C and washed three times with AR protein buffer (50 mM Tris-HCl, pH8.0; 150 mM NaCl; 0.05% NP-40) for 5 min per wash. Purified HA-tagged full-length AR (1 μg) was incubated with the blocked beads for at least 4 h at 4□°C. The beads were then washed three times with p300 protein buffer (25 mM HEPES-NaOH pH 7.5, 300 mM NaCl, 10% glycerol, 0.04% Triton X-100 and 0.5 mM TCEP) for 10 min per wash and incubated overnight at 4□°C with purified Flag-tagged p300 (Active motif, 81858) (200 ng or 1 μg) in 350 μL of p300 protein buffer containing the indicated compounds or an equivalent volume of DMSO. Beads were subsequently washed at least three times with p300 protein buffer containing the corresponding compounds for 10 min per wash. Bound proteins were eluted in SDS–PAGE loading buffer and analyzed by immunoblotting.

### Ternary complex formation by biolayer interferometry (BLI)

Anti-FLAG IgG biosensors (Sartorius, Cat. No. 18-5187) were pre-equilibrated in assay buffer containing 25 mM HEPES-NaOH (pH 7.5), 300 mM NaCl, 10% glycerol, 0.04% Triton X-100, and 0.5 mM TCEP. N-terminal FLAG-tagged full-length recombinant p300 protein (1 μg; Active Motif, Cat. No. 81858) was immobilized onto the biosensors at 25°C until the binding signal reached a plateau. After immobilization, the biosensors were washed in assay buffer to remove unbound protein and minimize nonspecific binding.

The biosensors were then incubated with recombinant GST-His-tagged AR-LBD protein (Invitrogen, Cat. No. A15675) at varying concentrations in assay buffer containing either 10 μM DALTAC-1 or an equivalent volume of DMSO. The association phase was monitored, followed by transfer of the biosensors into assay buffer to measure dissociation. Binding data were analyzed using Octet® Analysis Studio 13.0 (Sartorius). Sensorgrams obtained in the presence of DMSO were used as reference controls and subtracted from those obtained with DALTAC-1. The reference-subtracted data were globally fitted to a 1:1 binding model to determine the equilibrium dissociation constant (K_d_).

### Proximity ligation assay (PLA)

PLA was performed according to the manufacturer’s instructions (Navinci Diagnostics) using the Naveni™ TriFlex Cell kit for cultured cells (AR–p300) and the NaveniFlex™ Tissue Red kit for VCaP-CRPC tumor sections.

Cultured cells (Naveni TriFlex Cell): Cells were fixed, permeabilized, and blocked per kit protocol instructions. Primary antibodies against AR (1:400) and p300 (1:200) were applied in antibody diluent and incubated as recommended. After washes, Naveni proximity probes were added, followed by ligation and rolling-circle amplification steps according to the vendor’s workflow. Slides were counterstained with DAPI and mounted in antifade medium. PLA signals were visualized on a Zeiss LSM900 fluorescence microscope under identical exposure settings across conditions.

FFPE tumor sections (NaveniFlex Tissue Red): Sections were deparaffinized, rehydrated, and subjected to heat-induced epitope retrieval as specified by the kit. After blocking, primary antibodies against AR (1:1000) and p300 (1:100) were applied in antibody diluent and incubated per protocol. Following washes, NaveniFlex proximity probes, ligation, and amplification were performed according to the manufacturer’s instructions. Nuclei were counterstained with DAPI, slides were mounted, and images were acquired on a Zeiss LSM900 microscope using matched acquisition parameters.

Where indicated, PLA puncta were quantified as foci per nucleus using standardized thresholds across groups.

### Ternary complex ELISA assay

VCaP cells were plated in 96-well plates (50,000 cells per well) and cultured for 3 days in medium containing 5% charcoal-stripped FBS. Cells were treated with a serial dilution of DALTAC-1, Neg-1, or Neg-2 compounds for 3 hours. Following treatment, cells were lysed in 1× AlphaLISA SureFire Ultra Lysis Buffer (PerkinElmer) for 30 minutes at 4 °C with gentle mixing.

For detection, high-binding MaxiSorp 96-well plates (Thermo Fisher Scientific) were coated overnight at 4 °C with 50 µL of anti-AR antibody (clone AR441, Thermo Fisher Scientific) diluted 1:50 in PBS. After washing with TBST and blocking with 5% BSA for 2 hours at room temperature, 50 µL of cell lysate was added to each well and incubated for 1 hour at room temperature. Plates were washed three times with TBST and then incubated overnight at 4 °C with rabbit anti-p300 (D8Z4E, #86377, Cell Signaling Technology) antibody diluted 1:5000 in blocking buffer.

After washing, HRP-conjugated goat anti-rabbit secondary antibody (Invitrogen, #32260; 1:100,000 dilution) was added and incubated for 1 hour at room temperature. Plates were washed 7–8 times with TBST, and chemifluorescent signals were developed using QuantaRed™ Enhanced Chemifluorescent HRP Substrate (Thermo Fisher Scientific). Fluorescence (Ex = 570 nm, Em = 585 nm) was immediately measured using a Tecan Infinite M1000 Pro plate reader, with readings taken at 5 and 15 minutes.

### Nuclear immunoprecipitation

Cells (50 million) were washed once with ice-cold phosphate-buffered saline (PBS), scraped, and pelleted at 500 g for 5 minutes at 4 °C. The cell pellet was resuspended in 2 mL hypotonic buffer (20 mM HEPES-KOH pH 7.5, 10 mM KCl, 1 mM MgCl₂, 0.1% Triton X-100, 15 mM β-mercaptoethanol) supplemented with protease and phosphatase inhibitors and rotated for 1 hour at 4 °C. Nuclei were collected by centrifugation at 2,000 g for 5 minutes at 4 °C, the supernatant (cytoplasmic fraction) was discarded, and nuclei were washed once with hypotonic buffer.

The nuclear pellet was lysed in 1 mL nuclear lysis buffer (20 mM HEPES-KOH pH 7.5, 150 mM NaCl, 10 mM KCl, 1 mM MgCl₂, 0.5% Triton X-100, 15 mM β-mercaptoethanol) containing protease and phosphatase inhibitors and rotated for 1 hour at 4 °C. Benzonase® (500 U mL⁻¹; Sigma-Aldrich #E1014) was added during lysis to digest nucleic acids. Lysates were clarified by centrifugation at 16,000 g for 20 minutes at 4 °C, and the supernatant was collected and precleared with 50 µL Dynabeads™ Protein G magnetic beads (Thermo Fisher #10004D). Protein concentration was determined using the Pierce™ 660 nm Protein Assay (Thermo Fisher #22660).

For each immunoprecipitation, 400 µg of precleared nuclear lysate was incubated overnight at 4 °C with either anti-androgen receptor antibody (Abcam #ab108341) or anti-p300 antibody (Active Motif #61401). Antigen–antibody complexes were captured with 30 µL Dynabeads™ Protein G for 2 hours at 4 °C. Beads were washed three times with wash buffer (20 mM HEPES-KOH pH 7.5, 150 mM NaCl, 10 mM KCl, 1 mM MgCl₂).

For immunoblotting, bound proteins were eluted in 1× NuPAGE™ LDS Sample Buffer (Thermo Fisher #NP0007) supplemented with 50 mM DTT and subjected to SDS–PAGE. For mass spectrometry, washed beads were flash-frozen and stored at −80 °C for subsequent on-bead trypsin digestion.

### RNA extraction and quantitative polymerase chain reaction

Total RNA was isolated using QIAzol™ Lysis Reagent (QIAGEN). Briefly, 1 × 10□ cells were lysed in 700 µL QIAzol and incubated for 5 minutes at room temperature to allow for complete dissociation of nucleoprotein complexes. Following the addition of 140 µL chloroform, samples were vigorously mixed and centrifuged at 12,000 × g for 15 minutes at 4 °C. The aqueous phase was collected, mixed with 1.5 volumes of ethanol, and applied to a RNeasy Mini spin column (QIAGEN) for purification according to the miRNeasy Mini Kit protocol. RNA was eluted in RNase-free water, and the concentration and purity were assessed using a NanoDrop spectrophotometer (Thermo Fisher Scientific).

For quantitative PCR analysis, cDNA was synthesized from purified RNA using the Maxima™ First Strand cDNA Synthesis Kit (Thermo Fisher Scientific) according to the manufacturer’s instructions. qPCR reactions were performed with SYBR™ Green PCR Master Mix (Applied Biosystems) on a QuantStudio™ 7 Real-Time PCR System (Applied Biosystems). Primers used in this study are listed below: *GAPDH* forward (F): ACAACTTTGGTATCGTGGAAGG and reverse (R): GCCATCACGCCACAGTTTC; *KLK3/PSA* F: GACCAAGTTCATGCTGTGTGC and R: CCACTCACCTTTCCCCTCAAG; *TMPRSS2*F: GTCCCCACTGTCTACGAGGT and R: CAGACGACGGGGTTGGAAG; *NKX3-1* F: CCCACACTCAGGTGATCGAG and R: GAGCTGCTTTCGCTTAGTCTT; *MYC*F: CCTGGTGCTCCATGAGGAGAC and R: CAGACTCTGACCTTTTGCCAGG; *CCND1* F: GCTGCGAAGTGGAAACCATC and R: CCTCCTTCTGCACACATTTGAA.

### Tandem mass tag (TMT) proteomic analysis

Tandem mass tag (TMT) proteomic analysis was performed as previously described8, with minor modifications. VCaP cells were treated with DMSO or DALTAC-1 (JZY3032) in three independent biological replicates. Cell lysates were prepared, and protein concentrations were determined. For each sample, 75□µg of protein was reduced with 5 mM dithiothreitol, alkylated with 15 mM 2-chloroacetamide, precipitated with ice-cold acetone, and digested overnight with a trypsin/Lys-C mixture at a 1:50 protease-to-protein ratio. Peptides were labelled with TMTpro 16-plex reagents (Thermo Fisher Scientific, A44521), with DMSO replicates assigned to channels 126, 127N and 128N and JZY3032 replicates assigned to channels 130C, 131C and 132C. Labelled peptides were pooled and separated into 12 fractions by high-pH reversed-phase chromatography.

Fractions were analyzed at the University of Michigan Proteomics Resource Facility using an Orbitrap Ascend Tribrid mass spectrometer equipped with a FAIMS source and coupled to a Vanquish Neo UHPLC system. Peptides were separated on an Easy-Spray PepMap Neo C18 column, and data were acquired using two FAIMS compensation voltages (−45 and −65 V) and a MultiNotch MS3 method to minimize reporter-ion ratio distortion. Data were processed using Proteome Discoverer v3.0 and searched against the Swiss-Prot Homo sapiens protein database (TaxID 9606; version 5 February 2025). Peptide-spectrum matches and proteins were filtered to a false-discovery rate of ≤1%. TMT quantification was based on high-quality MS3 spectra with an average signal-to-noise ratio of ≥10 and isolation interference of <50%.

### Acetyl (lysine)-proteomics analysis

VCaP cells were treated with DMSO, 100 nM DALTAC-1 for 4 or 12 hours, or a combination of p300/CBPi and ARi for 12 hours, each in three biological replicates. The cells were then subjected to acetyl-lysine proteomics analysis as previously described^8^. In brief, total proteins were extracted, digested with trypsin, and acetylated peptides were enriched using anti–acetyl-lysine antibody–conjugated beads. Enriched peptides were analyzed by nanoLC-MS/MS on a Q Exactive™ HF mass spectrometer (Thermo Fisher Scientific), and raw data were processed using MaxQuant (v2.3.0.0) with acetyl-lysine and protein N-terminal acetylation set as variable modifications.

### Protein Identification by LC–tandem mass spectrometry

In-solution digestion: VCaP cells were treated with 100 nM DALTAC-1 (JZY3032) or DMSO for 4 hours. Co-IP assays were performed as described above. The beads were resuspended in 50 µl of 0.1M ammonium bicarbonate buffer (pH∼8). Cysteines were reduced by adding 50 µl of 10 mM DTT and incubating at 45 °C for 30 min. Samples were cooled to room temperature and alkylation of cysteines was achieved by incubating with 65 mM 2-chloroacetamide, under darkness, for 30 min at room temperature. An overnight (∼16 h) digestion with 1 ug sequencing grade, modified trypsin was carried out at 37 °C with constant shaking in a Thermomixer. Digestion was stopped by acidification and peptides were desalted using SepPak C18 cartridges using the manufacturer’s protocol (Waters). Samples were completely dried using a vacufuge. The resulting peptides were dissolved in 8 µl of 0.1% formic acid/2% acetonitrile solution, and 2 µl of the peptide solution was resolved on a nano-capillary reverse phase column (Acclaim PepMap C18, 2 micron, 50 cm, ThermoScientific) using a 0.1% formic acid/2% acetonitrile (Buffer A) and 0.1% formic acid/95% acetonitrile (Buffer B) gradient at 300 nl/min over a period of 180 min (2-22% buffer B in 110 min, 22-40% in 25 min, 40-90% in 5 min followed by holding at 90% buffer B for 5 min and requilibration with Buffer A for 25 min). The eluent was directly introduced into the Orbitrap Fusion tribrid mass spectrometer (Thermo Scientific, San Jose CA) using an EasySpray source. MS1 scans were acquired at 120K resolution (AGC target=1×106; max IT=50 ms). Data-dependent collision induced dissociation MS/MS spectra were acquired using the Top speed method (3 seconds) following each MS1 scan (NCE ∼32%; AGC target 1×105; max IT 45 ms).

Proteins were identified by searching the MS/MS data against the UniProt human database (sp_canonical TaxID=9606, v2025-02-05) using Proteome Discoverer (v3, Thermo Scientific). Search parameters included MS1 mass tolerance of 10 ppm and fragment tolerance of 0.2 Da; two missed cleavages were allowed. Carbamidomethylation of cysteine was considered a fixed modification and oxidation of methionine, deamidation of asparagine, and glutamine were considered as potential modifications. The false discovery rate (FDR) was determined using Percolator, and proteins/peptides with a FDR of ≤1% were retained for further analysis.

### RNA-seq and data analysis

VCaP cells were treated with 100 nM DALTAC-1 or a combination of 100 nM p300i and ARi for 4, 12, or 24 hours, each in two biological replicates. The cells were then subjected to RNA sequencing as previously described^7^. In brief, total RNA was extracted, and ribosomal RNA was removed using the RiboErase module of the KAPA RNA HyperPrep Kit (Roche). rRNA-depleted RNA was used for library preparation following the manufacturer’s protocol, and libraries were quality-checked on an Agilent 2100 Bioanalyzer before sequencing on an Illumina HiSeq 2500 platform (paired-end, 2 × 100 bp). Sequencing reads were quantified with Kallisto (v0.46.1), normalized with edgeR (v3.39.6), and analyzed for differential expression using limma-voom (v3.53.10). Gene set enrichment analysis was performed using fgsea (v1.24.0), and data visualization employed tidyverse, gplots, ggplot2, and EnhancedVolcano (v1.15.0) packages in R.

### ChIP-seq and data analysis

Chromatin immunoprecipitation followed by sequencing (ChIP-seq) was performed as previously described^7^. In brief, cells were crosslinked in 1% formaldehyde, quenched with glycine, and lysed for chromatin extraction. Chromatin was sheared to 200–600 bp fragments using a Bioruptor (Diagenode) and subjected to immunoprecipitation with antibodies against transcription factors (AR, FOXA1, p300, ERG, and BRD4) or histone marks (H3K27ac, H2BK20ac, and RNA Pol II) using the iDeal ChIP-seq Kit (Diagenode). Drosophila spike-in chromatin (Active Motif, 53083) and spike-in antibody (Active Motif, 61686) were added during the overnight incubation. After reverse crosslinking and DNA purification, sequencing libraries were prepared and sequenced on an Illumina HiSeq 2500 platform. For PR, SRC2, and CBP ChIP-seq, we performed double-cross-linking. Briefly, 10 million cells were cross-linked with 250 μL of 200 mM DSP (Thermo Fisher Scientific, 22585) for 30 min at room temperature. PBS was then added to a final volume of 9 mL, followed by 40 μL of 0.5 M EDTA. Formaldehyde (250 μL, 37%) was added, and the cells were incubated for an additional 10 min at room temperature. Cross-linking was quenched by adding 1 mL of 1.25 M glycine and 200 μL of 10% Tween-20, followed by incubation for 5 min. Samples were then processed according to the ChIP–seq protocol described previously.

ChIP-seq reads were trimmed using Trimmomatic (v0.39). Alignment was then performed separately to both the human genome (hg38) and the drosophila genome (dm6) using BWA (v0.7.17). Low-quality and duplicate reads were filtered with SAMtools and Picard, respectively. Normalization ratios were calculated based on the lowest amount of uniquely aligned reads per ChIP target to each genome as recommended on the vendor’s website. Reads were then downsampled per the normalization ratios. Peak calling was then performed with MACS2 using narrow or broad settings depending on the target, and blacklisted genomic regions were removed using BEDTools with the ENCODE exclusion list. Processed data were converted to BigWig format using UCSC wigToBigWig for visualization.

### Enrichment heatmaps and profile plots

Read density heatmaps and average enrichment profiles were generated using deepTools. For histone marks, signals were plotted within ±2.5 kb of the reference point, and for transcriptome factors within ±1 kb. The parameters --skipZeros, --averageType mean, and --plotType se were applied. ENCODE blacklist regions (ENCFF356LFX) were excluded from analysis. Non-promoter regions were defined using the ChIPseeker R package tool annotatePeak excluding ±1 kb windows surrounding annotated gene TSS regions.

### Prostate organoid culture

Mouse prostate organoids and patient-derived xenograft organoids were established and maintained following previously described protocols^41,71^. Briefly, organoids were cultured in 50 µL Matrigel domes and overlaid with prostate organoid medium, with media changed every 2–3 days and passaging performed weekly. For growth and drug-response assays, organoid fragments were dissociated into single cells and used to generate new organoids (1,000 cells per dome for standard viability assays) in complete organoid medium. Cell viability was measured using the CellTiter-Glo 3D assay (Promega) according to the manufacturer’s instructions.

### Human prostate tumor xenograft models

Male SCID mice (6–8 weeks old) were obtained from the breeding colony at the University of Michigan and maintained under specific pathogen–free conditions. All animal procedures were approved by the University of Michigan Institutional Animal Care and Use Committee (IACUC). Subcutaneous xenograft tumors were established by injecting 3 × 10□ VCaP cells suspended in serum-free medium mixed 1:1 with Matrigel (BD Biosciences) into both dorsal flanks. Tumor growth was monitored weekly using digital calipers, and volumes were calculated as (π/6) × (length × width²). At study endpoints, mice were euthanized, and tumors were excised and weighed.

The castration-resistant VCaP model was generated as previously described^7^. Briefly, mice were castrated once tumors reached 150–200 mm³, and treatment with DALTAC-1 (30 mg/kg i.p. or 60 mg/kg p.o.), combination of p300i (30 mg/kg p.o.) and ARi (10 mg/kg p.o.), or vehicle was initiated after tumor regrowth to baseline size for 24 days.

### Tissue microarray generation

A tissue microarray (TMA) was created according to a design involving 8 unique cases from 4 different subgroups: vehicle (n=2), p300i+ARi (n=2), DALTAC-1 ip (n=2), and DALTAC-1 po (n=2). Two independent samples were selected from each subgroup. To ensure data reliability and representativeness, a replicate core from each sample was included in the TMA, resulting in a focused 16-core TMA. Technical details are as follows: Briefly, FFPE donor blocks for the selected samples were chosen by two study pathologists (R.M. and S.M.) after reviewing the hematoxylin & eosin (H&E) staining. The pathologists then annotated the H&E slides to identify the most representative tumor area of interest (AOI) on the surface of each donor block. The Semiautomatic tissue arrayer from Beecher Instruments (Sun Prairie, Wisconsin, United States) was used to punch 1 mm-diameter cores from each marked region. These cores were carefully transferred into a recipient acrylic block in a grid-like pattern. Once transferred, the recipient block was incubated at 37 degrees Celsius for 30 minutes to ensure proper fusion of the cores within the paraffin. The prepared TMA block was then sectioned at 4-micron thickness and stained with H&E to assess the quality of the final TMA.

### Sequential multiplex immunofluorescence (Seq-mIF) by COMET analysis

Four-micron-thick sections from the PDX TMA, mounted on positively charged slides, were cut and prepared according to the guidelines and recommendations. Subsequently, the slides were deparaffinized using a standard xylene and graded ethanol series, followed by heat-induced epitope retrieval (HIER) as per the Lunaphore COMET protocol to unmask target epitopes^72^. Automated sequential immunofluorescence (seqIF™) staining was performed on the COMET™ instrument (Lunaphore) to enable highly multiplexed protein detection with the target primary antibodies. This process involved cycles of staining, imaging, and elution managed by the COMET Control software through controlled temperature and microfluidic reagent delivery. Each cycle included incubation with unconjugated primary antibodies, followed by species-specific fluorescently conjugated secondary antibodies (e.g., Alexa Fluor 555 and 647), automated image acquisition including DAPI counterstaining, and complete antibody removal via elution. After the final staining cycle, the COMET Control software automatically stitched, aligned, and stacked images to produce a single OME-TIFF file for each sample. During post-processing, autofluorescence background was systematically subtracted using reference images acquired from unstained tissue sections at the beginning of the protocol.

### Statistics used for Seq-mIF

The raw prostate TMA data included single-cell measurements across multiple markers along with their spatial locations. Markers were grouped into two panels based on biological relevance, as shown in the plots. To normalize marker expression and reduce skewness, a log1p transformation (log1p(Expression)) was applied. Outliers were identified and removed separately for each marker within each group. Outliers were defined as values exceeding ±5 median absolute deviations (MAD) from the group-specific median. The Median Absolute Deviation (MAD) is a robust measure of statistical dispersion that calculates the median of the absolute deviations from the median, making it less sensitive to extreme outliers than standard deviation. This approach preserves most of the data while minimizing the impact of extreme values.

### Prostate patient-derived xenograft (PDX) models

Prostate cancer PDX models MDA-PCa-146-12 and MDA-PCa-183 were established and maintained as previously described^7^. Briefly, tumor fragments (∼2 mm³) were implanted subcutaneously into male SCID mice and propagated through serial passage. Once tumors reached approximately 200 mm³, mice were randomized into treatment groups and administered vehicle, DALTAC-1 (30 mg/kg, i.p., 3x/week), ARV-766 (10 mg/kg, p.o, 5x/week), or enzalutamide (10 mg/kg, p.o 5x/week). Tumor volumes were measured weekly, and all procedures were conducted in accordance with protocols approved by the University of Michigan Institutional Animal Care and Use Committee (IACUC).

### Statistics and reproducibility

All experimental procedures and statistical analyses were performed as previously described^7^. In brief, key experiments were independently repeated at least three times, with most quantitative assays carried out in biological replicates as noted. Representative images and immunoblots reflect reproducible results, and the number of biological replicates is provided in the figure legends. No statistical method was used to predetermine sample size, no data were excluded, and experiments were neither randomized nor blinded. Two-sided statistical tests were used unless otherwise stated. Comparisons between two groups used unpaired t-tests or Wilcoxon rank-sum tests, and multiple comparisons were corrected using the Benjamini– Hochberg method. Exact p-values are reported in the figure legends.

## Extended Data Figure Legends

**Extended Data Figure 1. Development of AR-p300/CBP DALTAC-1.**

a. Chemical structure of the AR ligand-binding domain binder JZY3221 (ARi) and p300/CBP bromodomain inhibitor GNE-049 (p300i) and the workflow of AR-p300 DALTAC compound design.

b. Structure–activity relationship (SAR) table summarizing linker structures, cellular AR–p300 ternary-complex induction in VCaP cells, antiproliferative IC50 and maximum inhibition (Imax) values in LNCaP and 22Rv1 cells and inhibition of ARE-GFP expression in HEK293-ARE-GFP cells for the indicated compounds. Maximum ternary-complex induction was measured at 300 nM, Imax was determined at 3 μM, HEK293-ARE-GFP cells were treated with 10 nM compounds for 48 h.

c. Pharmacokinetic parameters of compound 7 (JZY3031) and DALTAC-1 (JZY3032) in mice following intravenous (i.v.; 2.0 mg kg−1) or oral (p.o.; 5.0 mg kg−1) administration. The table reports terminal half-life (t1/2), maximum plasma concentration (Cmax), area under the concentration–time curve from 0 to 24 h (AUC0–24), steady-state volume of distribution (Vss), clearance (CL) and oral bioavailability (F).

d. Chemical structures of the indicated enzalutamide-based AR–p300/CBP DALTAC compounds.

e. Dose–response curves of VCaP, LNCaP, and 22Rv1 cells treated with the indicated compounds for 5 days. Data represent mean ± SD; n=3, three independent experiments.

f. Chemical modifications of DALTAC-1 (JZY3032) used to generate the inactive analogues JZY3137 (Neg-1; impaired p300/CBP binding with retained AR binding) and JZY3138 (Neg-2; impaired AR binding with retained p300/CBP binding).

g. Dose–response curves from fluorescence polarization (FP) assays measuring p300 binding for Neg-1, Neg-2, GNE-049 (p300 bromodomain inhibitor), JZY3221, JZY3222, and DALTAC-1. Data represent mean ± SD; n = 4, three independent experiments.

h. Dose–response curves from FP assays measuring CBP binding for Neg-1, Neg-2, GNE-049 (p300 bromodomain inhibitor), JZY3221, JZY3222, and DALTAC-1. Data represent mean ± SD; n=4, three independent experiments.

i. Dose–response curves from FP assays measuring AR binding for Neg-1, Neg-2, R1881, JZY3221, JZY3222, and DALTAC-1. Data represent mean ± SD; n=4, three independent experiments.

j. Immunoblot analysis demonstrating stabilization of p300 and AR in a cellular thermal shift assay. LNCaP cells were treated with DMSO or 100 nM DALTAC-1 for 4 h and heated at the indicated temperatures, three independent experiments.

**Extended Data Figure 2. AR-p300/CBP DALTAC-1 specifically inhibits AR-positive prostate cancer growth.** a. Dose-response curves and IC_50_ values of AR-positive CWRR1 and 22Rv1 cells treated with DALTAC-1, CBPD-409, p300/CBPi+ARi, or CCS1477 (p300/CBP BRD inhibitor) for 5 days. Data represent mean ± SD; n=3, three independent experiments.

b. Dose-response curves of VCaP and LNCaP cells treated with DALTAC-1, enzalutamide, darolutamide, ARV-766, or AR-BRD4 RIPTAC-1 for 5 days. Data represent mean ± SD; n=3, three independent experiments.

c. Representative images of colonies of DMSO, 10 nM DALTAC-1, or p300/CBPi+ARi treated LNCaP cells for 14 days. n=3, three independent experiments.

d. Dose-response curves and IC_50_ values of AR-positive 22Rv1 cells treated with DALTAC-1, Neg-1, or Neg-2 for 5 days. Data represent mean ± SD; n=3.

e. Relative viability of LNCaP cells treated with 20 nM DALTAC-1 together with increasing concentrations of p300/CBPi or ARi for 5 days. Data represent mean ± SD; n=6, three independent experiments.

f. Dose-response curves and IC_50_ value of NCI-H211 cells treated with DALTAC-1, CBPD-409, or p300i+ARi for 5 days. Data represent mean ± SD; n=3, three independent experiments.

g. Dose-response curves of MDA-MB-453 and U-87 MG cells treated with DALTAC-1, CBPD-409, or ARV-766 for 5 days. Data represent mean ± SD; n=3, three independent experiments.

h. Growth curves of VCaP cells treated with 100 nM DALTAC-1, p300/CBPi + ARi or CBPD-409 under 4-h pulse–washout or continuous-treatment conditions, monitored by Incucyte Live-Cell Analysis. For pulse–washout treatment, compounds were removed after 4 h and replaced with compound-free medium. Data represent mean ± SD; n=6, three independent experiments. Significance at endpoints was assessed using two-sided t-test.

i. Dose-response curves of VCaP cells treated with DALTAC-1 or ARi + p300/CBPi under washout or continuous treatment conditions. Data represent mean ± SD. n=6, three independent experiments. Significance at endpoints was assessed using two-sided t-test.

**Extended Data Figure 3. DALTAC-1 induces formation of an AR-p300/CBP-DALTAC-1 ternary complex.**

a. Immunoblot analysis of the interaction between purified full-length HA–AR and p300 proteins following incubation with DMSO, 10 μM DALTAC-1, ARi + p300/CBPi, Neg-1, or Neg-2, followed by HA immunoprecipitation. Three independent experiments.

b. BLI sensorgrams showing dose-dependent binding of GST–AR-LBD to immobilized p300 with increasing concentrations of DALTAC-1, yielding an apparent Kd of 4.3 μM, whereas minimal binding was detected in the DMSO control. Three independent experiments.

c. BLI sensorgrams using immobilized Flag–FOXA1 (residues 1–268) as a negative control showed no detectable binding of GST–AR-LBD alone, DALTAC-1 alone, or their combination. Three independent experiments

d. Representative BLI sensorgrams of GST–AR-LBD binding to immobilized Flag–p300 in the presence of 10 μM ARi, p300/CBPi, or their combination. Three independent experiments.

e. Representative BLI sensorgrams showing DALTAC-1-induced binding of GST–AR-LBD to immobilized Flag–p300 in the absence or presence of excess p300/CBPi (100 μM). Three independent experiments.

f. Immunoblot analysis of a co-immunoprecipitation assay showing increased interaction between AR and CBP in VCaP cells treated with 100 nM DALTAC-1 for 4 h. Three independent experiments.

g. Volcano plot showing changes in p300-interacting proteins in VCaP cells treated with 100 nM DALTAC-1 versus DMSO for 4 hours by IP-mass spectrometry. Histone proteins are highlighted in blue and AR in red. Dotted lines indicate log2 FC=1 and p=0.05. Data represent n = 3. p values were calculated using a two-sided t-test.

h. Exploratory reciprocal AR IP–mass spectrometry analysis showing proteins with the largest increases in abundance in AR immunoprecipitates from DALTAC-1-treated relative to DMSO-treated VCaP cells. p300 and CBP are highlighted in blue. n=1.

i. Representative images of AR–p300 PLA foci in 22Rv1 cells treated with indicated compounds for 4 hours, scale bar = 10 µm. Three independent experiments.

j. Left: schematic illustrating the workflow of the two-step sequential co-immunoprecipitation assay. Right: immunoblot analysis showing the co-occurrence of Flag-AR and HA-AR within the V5 antibody-enriched DALTAC-1-induced AR–V5-p300 complex in HEK293 cells treated with 1 μM DALTAC-1 for 4 h.

k. Immunoblot analysis showing that treatment of 22Rv1 cells with 100 nM DALTAC-1 for 4 hours increased the interaction of p300 with full-length AR, but not with AR-V7. Three independent experiments.

l. Immunoblot analysis showing increased association of AR-C619Y with p300 in HEK293 following 1 μM DALTAC-1 treatment for 4 hours. Three independent experiments.

m. Immunoblot analysis of the interactions between HA-tagged full-length AR and the Flag-tagged p300 core domain in HEK293 cells treated with DMSO or 1 μM DALTAC-1 for 4 hours. Three independent experiments.

n. Immunoblot analysis of co-IP assay showing the interaction of HA-tagged AR-NTD-DBD or AR-DBD-LBD domains with Flag-tagged full-length p300 in HEK293 cells treated with DMSO or 1 μM DALTAC-1 for 4 hours. Three independent experiments.

o. Immunoblot analysis of co-IP assay showing the interaction of HA-tagged AR-DBD-LBD with Flag-tagged p300 core domain in HEK293 cells treated with DMSO or 1 μM DALTAC-1 for 4 hours. Three independent experiments.

p. Computational modeling of the AR-LBD/DALTAC-1/p300-BRD ternary complex. DALTAC-1 (gray sticks), AR-LBD (pink, PDB: 2AXA), and p300-BRD (cyan, PDB: 5W0E). Protein–protein interactions are shown as gray dashed lines, protein–ligand interactions as yellow double-dashed lines, and π–cation interactions as blue dashed lines. Nitrogen, oxygen, fluorine, and chlorine atoms are colored blue, red, light blue, and green, respectively.

q. Framewise MM-GBSA calculations of DALTAC-1:AR binary, DALTAC-1-p300 binary, and DALTAC-1-AR-p300 ternary complexes calculated from 101 snapshots sampled over the final 250 ns of the molecular-dynamics trajectories. The dashed line denotes the mean, and the shaded region denotes ±1 s.d. across sampled conformations.

r. Mean and SD values of MM-GBSA binding free-energy calculations for binary and ternary complexes depicted in Extended Data Fig 3q.

s. Immunoblot analysis showing that treatment with 1 μM DALTAC-1 for 4 hours increased the interaction between wild-type AR and p300, whereas the indicated AR and p300 mutations markedly impaired the DALTAC-1-induced association. Three independent experiments.

**Extended Data Figure 4. DALTAC-1 reprograms the substrate engagement of p300/CBP in an AR-dependent manner.** a. Immunoblot analysis of indicated histone marks in VCaP cells treated with DMSO, 10 nM DALTAC-1, Neg-1, Neg-2, p300i + ARi, or CBPD-409 for 4 hours. H2B, separate-gel control from the same samples. Three independent experiments.

b. Immunoblot analysis of indicated histone marks and proteins in LNCaP cells treated with DMSO, 100 nM DALTAC-1, p300/CBPi + ARi, or CBPD-409 for 4 hours. H2B, separate-gel control from the same samples. Three independent experiments.

c. Immunoblot analysis of indicated histone marks and proteins in 22Rv1 cells treated with DMSO, 100 nM DALTAC-1, p300/CBPi + ARi, or CBPD-409 for 4 hours. GAPDH, same-gel loading control. Three independent experiments.

d. Left: Immunoblot analysis of indicated histone marks and proteins in HEK293 cells expressing varying levels of ectopic AR and treated with 10 nM DALTAC-1 or CBPD-409 for 4 hours. H2B, separate-gel control from the same samples. Three independent experiments. Right: Bar charts showing the relative expression of the indicated histone marks in HEK293 cells.

e. Immunoblot analysis of indicated histone marks and proteins in VCaP cells pre-treated with 100 nM DALTAC-1 or a combination of p300i and ARi for 2 hours, followed by washout and culture for the indicated durations. H2B, separate-gel control from the same samples. Three independent experiments.

f. Acetyl-lysine mass spectrometry heatmap showing H2BNTac levels in VCaP cells treated with DMSO, 100 nM DALTAC-1 or combined p300/CBPi + ARi for 12 h. Values are shown as z scores; n= 3.

g. Volcano plots showing altered lysine acetylation levels in VCaP cells treated with 100 nM DALTAC-1 versus DMSO for 4 hours. H2BNTac sites are highlighted in blue, and components of the AR complex are highlighted in red. Statistical tests were two-tailed t-tests, n=3.

h. Acetyl-lysine mass spectrometry heatmap showing the top 50 acetyl-lysine sites most increased or decreased in VCaP cells treated with DMSO, 100 nM DALTAC-1, or ARi + p300i for 3 hours. Data represent n = 3.

i. Relative ARE–GFP reporter activity in HEK293-ARE-GFP cells ectopically expressing AR alone or together with wild-type SRC2, the multi-lysine-to-alanine SRC2 mutant (Multi K-A), or the multi-lysine-to-glutamine SRC2 mutant (Multi K-Q). Data represent mean ± SD; n = 3, three independent experiments. Significance was assessed by two-tailed t-tests. P values are shown.

j. Immunoblot analysis confirming expression of the indicated proteins in the corresponding samples shown in panel i. GAPDH, same-gel loading control. Three independent experiments.

**Extended Data Figure 5. DALTAC-1 inhibits oncogenic transcriptional programs in prostate cancer cells.**

a. Representative fluorescence microscopy images of LNCaP-ARE-GFP cells treated with DMSO, 100 nM DALTAC-1, Neg-1, or Neg-2 for 48 hours. Scale bar, 50 µm. Three independent experiments.

b. Relative mRNA expression of the indicated genes in VCaP cells treated with DMSO, 100 nM DALTAC-1, CBPD-409, or p300/CBPi + ARi for 4 h, as determined by quantitative PCR. Data represent mean ± SD; n = 3. Target gene expressions were normalized to *GAPDH* expression. Three independent experiments.

c. GSEA plots for MYC- and mTORC1-signalling gene sets using the fold-change-ranked gene signature from DALTAC-1-treated versus DMSO-treated VCaP cells. Normalized enrichment scores (NES) and nominal p values are shown; n = 2. Significance was assessed by GSEA using gene-set permutations.

d. Normalized enrichment scores for Hallmark gene sets in VCaP cells treated with 100 nM DALTAC-1 or combined p300/CBPi + ARi versus DMSO for 4 h.

e. RNA-seq heatmaps for MYC target genes in VCaP cells treated with 100 nM DALTAC-1 or p300/CBPi = ARi for 4 hours. n=2.

f. RNA polymerase II ChIP–seq signal intensities within ±2 kb of transcription start sites (TSS) of AR- and p300-regulated genes in VCaP cells treated with 100 nM DALTAC-1 for 1 hour and 3 hours. n = 1.

g. Volcano plot showing changes in protein abundance in VCaP cells treated with 100 nM DALTAC-1 vs DMSO for 12 h, as determined by tandem mass tag (TMT)-based mass spectrometry. AR- and MYC-regulated proteins, together with AR and p300, are highlighted. Data are from n = 3. Significance was assessed using two-sided *t*-tests followed by adjusted correction for multiple comparisons. The axes show log2 fold change and −log10(*q* value).

h. Chemical structure of JZY4055, a AR–p300/CBP DALTAC using enobosarm and GNE-049 as binding moieties.

i. Dose-response curves and IC_50_ values of VCaP cells treated with JZY4055, DALTAC-1, or enobosarm for 5 days. Data represent mean ± SD; n = 3. Three independent experiments.

j. Relative mRNA expression of the indicated genes in VCaP cells treated with indicated compounds for 6 h. Data represent mean ± SD; n = 3. Target gene expressions were normalized to *GAPDH* expression. Three independent experiments.

k. Immunoblot analysis of the indicated histone acetylation marks and proteins in VCaP cells treated with DMSO or 1 μM JZY4055 for 6 h. GAPDH, same-gel as loading control. Three independent experiments.

**Extended Data Figure 6. DALTAC-1 remodels the chromatin landscape of the AR neo-enhanceosome.**

a. ChIP-seq read-density heatmaps of p300 and AR at DALTAC-1 enriched AR binding sites in VCaP cells treated with DMSO or 100 nM DALTAC-1 for 3 hours, n = 1.

b. ChIP–seq read-density heatmaps showing CBP occupancy at the top 10,000 AR-bound enhancer sites in VCaP cells treated with DMSO or 100 nM DALTAC-1 for 3 h, n = 1.

c. Target and background frequencies and statistical significance of HOMER motifs enriched at non-promoter p300-binding sites depleted (left) or enriched (right) following treatment of VCaP cells with 100 nM DALTAC-1 for 3 h. AR-family motifs are highlighted in blue, FOX-family motifs in red and other motifs in grey. P values were calculated using HOMER’s cumulative binomial test.

d. Target and background frequencies and statistical significance of HOMER motifs enriched at non-promoter CBP-binding sites following treatment of VCaP cells with 100 nM DALTAC-1 for 3 h. Motif families are highlighted as indicated. *P* values were calculated using HOMER’s cumulative binomial test.

e. Genomic distribution of DALTAC-1 depleted AR ChIP-seq peaks and DALTAC-1 enriched AR peaks in VCaP cells treated with 100 nM DALTAC-1 for 3 h.

f. Top five known HOMER motifs enriched within top 2000 DALTAC-1 depleted or enriched AR sites in VCaP cells. P-values were calculated using HOMER’s binomial test.

g. AR ChIP–seq tracks on *WDFY4* gene loci in VCaP cells treated with 100 nM DALTAC-1 or DMSO for 3 hours. Highlighted motifs are shown in different colors.

h. Venn diagram illustrating overlap of genome-wide progesterone receptor (PR) ChIP-seq peaks in VCaP cells treated with DMSO or 100 nM DALTAC-1 for 3 h.

i. ChIP-seq read-density heatmaps of PR and AR at PR binding enhancer sites in VCaP cells treated with DMSO or 100 nM DALTAC-1 for 3 h, n = 1.

j. Venn diagram illustrating overlap of genome-wide AR and p300 ChIP-seq peaks in VCaP cells treated with DMSO (top) or 100 nM DALTAC-1 (bottom) for 3 hours.

k. ChIP-seq read-density heatmaps of indicated transcription factors, transcriptional cofactors, and histone modifications at AR/p300 co-bound non-promoter sites in VCaP cells treated with DMSO or 100 nM DALTAC-1 for 3 hours, n = 1.

**Extended Data Figure 7. DALTAC-1 inhibits the activity of the AR/ERG neo-enhanceosome.**

a. ChIP-seq read-density heatmaps of ERG at top 10,000 ERG binding sites in VCaP cells treated with DMSO or 100 nM DALTAC-1 for 3 hours, n = 1.

b. Venn diagram illustrating overlap of genome-wide ERG ChIP-seq peaks in VCaP cells treated with DMSO or 100 nM DALTAC-1 for 3 hours.

c. ChIP-seq read-density heatmaps of ERG at indicated sites in VCaP cells treated with DMSO or 100 nM DALTAC-1 for 3 hours, n = 1.

d. Genomic distribution of DALTAC-1 depleted ERG ChIP-seq peaks and DALTAC-1 enriched ERG peaks in VCaP cells.

e. Target and background frequencies and statistical significance of HOMER motifs enriched at non-promoter ERG-binding sites depleted (left) or enriched (right) following treatment of VCaP cells with 100 nM DALTAC-1 for 3 h. AR-family motifs are highlighted in red, FOX-family motifs in blue and ETS-family motifs in green. *P* values were calculated using HOMER’s cumulative binomial test.

f. Venn diagram illustrating overlap of genome-wide AR and ERG ChIP-seq peaks in VCaP cells treated with DMSO.

g. ChIP–seq read-density heatmaps of SRC2 at the top 5,000 AR–ERG co-bound enhancer sites in VCaP cells treated with DMSO or 100 nM DALTAC-1 for 3 h, n = 1.

h. ChIP–seq tracks of indicated proteins and histone mark on *TMPRSS2* gene loci in VCaP cells treated with 100 nM DALTAC-1 or DMSO for 3 hours.

**Extended Data Figure 8. DALTAC-1 suppresses prostate cancer growth across multiple preclinical models.**

a. Representative images of *Rosa26^ERG^/Pten-/-* and *FOXA1* mt*/Trp53-/-* organoids treated with DMSO, 100 nM DALTAC-1, or 1000 nM DALTAC-1 for 7 days. Scale bar, 200 µm. Three independent experiments.

b. Immunoblot analysis of indicated proteins in *Rosa26^ERG^/Pten-/-, Pten-/-, FOXA1* mt*/Trp53-/-* and *Trp53-/-* organoids treated with DMSO or 1000 nM DALTAC-1 for 24 h. GAPDH, same-gel loading control. Three independent experiments.

c. Schematic illustrating the design, treatment schedule, and pharmacodynamic and endpoint analyses of the DALTAC-1 efficacy study using the VCaP-CRPC xenograft model.

d. Endpoint tumor weights in the VCaP-CRPC model treated with vehicle, DALTAC-1 ip, DALTAC-1 po, or combination of p300i and ARi po. Data are mean□±□SEM; n□=□16 per group. Two-sided t-test.

e. Tumor volume curves of individual tumors from the VCaP-CRPC study shown in **Fig. 4d**.

f. Violin plots showing the log10-transformed expression levels of individual targets across single cells based on Seq-mIF staining from PD5 tumors in the VCaP-CRPC study (n > 30,000 cells). The violin outline represents the kernel density estimate; grey dots indicate individual cells; and the central white line denotes the median.

g. Graph depicting the tumor volume curves of MDA-PCa-146-12 CRPC PDX tumors treated with vehicle or 30 mg/kg DALTAC-1 ip. Data are presented as mean +/− SD (vehicle: n□=□16, DALTAC-1 ip: n=7). P values were calculated using a two-sided t-test.

h. Tumor volume curves in the MDA-PCa-183 PDX model treated with vehicle or DALTAC-1 (30□mg/kg, ip, 5×/week during the first week, then 2×/week for an additional 4 weeks). Data are mean□±□SEM; n□=□8 in vehicle group and n=7 in DALTAC-1 treated group. P values were calculated using a two-sided t-test.

i. Tumor volume curves of individual tumors from the MDA-PCa-183 PDX study shown in **Extended Data Fig. 8h.**

j. Waterfall plot showing the percentage change in individual tumor volumes from baseline after 35 days of treatment in the MDA-PCa-183 PDX study.

k. Kaplan–Meier survival curve of MDA-PCa-183 PDX–bearing mice, with survival defined as the time for tumors to reach threefold the initial volume. Statistical significance was assessed using a two-sided log-rank (Mantel–Cox) test.

**Extended Data Figure 9. DALTAC-1 exhibits no apparent toxicity in tumor-bearing mice.** a. Mouse body weight (%) measurements throughout the treatment period from VCaP-CRPC study (two-sided t-test). Data are presented as mean +/− SEM (vehicle: n□=□7, DALTAC-1 i.p.: n=8, DALTAC-1 p.o.: n=7, p300i+ARi: n=7).

b. Liver and kidney function tests from VCaP-CRPC mouse study. Data are presented as mean +/− SD (For ALT, BUN and CREA, vehicle: n□=□7, DALTAC-1 IP: n=8, DALTAC-1 PO: n=7, p300i+ARi: n=7; for AST, vehicle: n□=□7, DALTAC-1 IP: n=8, DALTAC-1 PO: n=6, p300i+ARi: n=7).

c. Complete blood count from VCaP-CRPC mice. Data are presented as mean +/− SD (vehicle: n□=□7, DALTAC-1 IP: n=8, DALTAC-1 PO: n=7, p300i+ARi: n=7).

d. Mouse body weight (%) measurements throughout the treatment period from MDA-PCa-146-12 CRPC study, Data are presented as mean +/− SEM (vehicle: n□=□9, DALTAC-1 ip: n=4).

e. Mouse body weight (%) measurements throughout the treatment period from MDA-PCa-183 study. Data are presented as mean +/− SEM (vehicle: n□=4, DALTAC-1 ip: n=4).

