## Supplemental Information for "Rewiring Oncogenic Transcriptional Complexes with Domain-ALTerAtion Chimeras (DALTACs) in Prostate Cancer"

*(SI Guide)*

### General Synthesis Information

All commercially available reagents and solvents were used without further purification. All chemical reactions were monitored by thin-layer chromatography (TLC) plates with visualization under UV light (254 or 365 nm). $^{1}H$ NMR spectra were performed with Bruker AV-400 spectrometer, $^{13}C$ NMR spectra were recorded on Bruker AV-400 spectrometer at 101 MHz, internal reference was either TMS or deuterated NMR solvent. In reported spectral data, the format (δ) chemical shift (multiplicity, *J* values in Hz, integration) was used with the following abbreviations: s = singlet, d = doublet, t = triplet, q = quartet, hept = heptet, dd = doublet of doublets, and m = multiplet. Low-resolution mass spectrometric (MS) analysis was carried out with a Waters UPLC ACQUITY QDa mass spectrometer. Flash column chromatography was performed by Teledyne CombiFlash RF+ using RediSep Rf silica gel flash column. The final compounds were all purified by a C18 reverse phase preparative HPLC column (SunFire Prep C18 OBD 5 μm, 50 mm × 100 mm) with solvent A (0.1% TFA in $H_2O$) and solvent B (0.1% TFA in MeCN) as eluents at 60 mL/min flow rate. The purity of all the final compounds was measured and confirmed to be >95% by rapid ultra-performance liquid chromatography mass spectrometry (UPLCMS) analysis (10-100% MeCN in $H_2O$ containing 0.1% formic acid in 5 min, 1.0 mL/min flow rate) with a C18 column (ACQUITY UPLC BEH C18 1.7 μm, 2.1 mm × 50 mm).

### Scheme 1: Synthesis of Compounds 1-6.

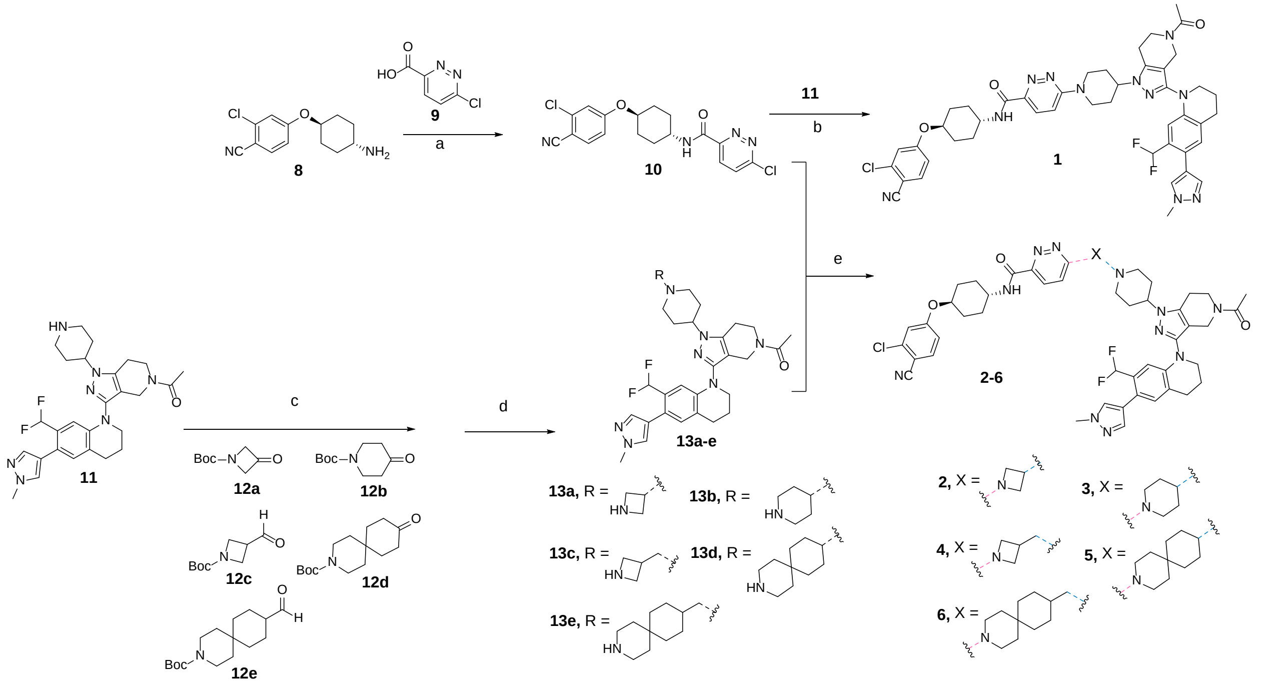

***Reagents and conditions*****:** (a) HATU, DMF, $Et_3N$, rt, 15 min; (b) $K_2CO_3$, DMF, 65 $^oC$, 3 h. (c) $Et_3N$, Sodium triacetoxyborohydride, DCM, rt, 3 h; (d) DCM, TFA, 40 $^oC$, 3 h; (e) $K_2CO_3$, DMF, 65 $^oC$, 3 h.

Intermediate **11** was prepared according to the reported procedures.[1]

***6-chloro-N-((1r,4r)-4-(3-chloro-4-cyanophenoxy)cyclohexyl)pyridazine-3-carboxamide***

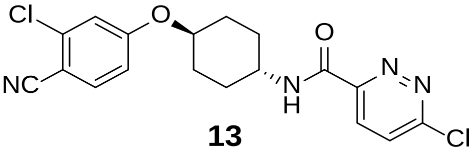

**13**

Step a: 6-chloropyridazine-3-carboxylic acid **8** (5.0 g, 31.54 mmol) was dissolved in DMF (100 mL), HATU (12.0 g, 31.54) and $Et_3N$ (4.8 g, 47.31 mmol) was added. The mixture was stirred at room temperature for 2 min, then intermediate **9** (7.9 g, 31.54 mmol) was added in to the mixture. The result mixture was stirred at rt for additional 13 min, after that the mixture was diluted with water and extracted with DCM. The organic phases were combined and concentrated under reduced pressure. The residue was purified by flash column chromatography (0–50% EtOAc/*n*-hexane) to give intermediate **10** (5.5 g, yield: 45%) as yellow solid. $^1$H NMR (400 MHz, DMSO-$d_6$) δ 9.17 (d, *J* = 8.1 Hz, 1H), 8.23 (d, *J* = 8.8 Hz, 1H), 8.11 (d, *J* = 8.8 Hz, 1H), 7.86 (d, *J* = 8.8 Hz, 1H), 7.40 (d, *J* = 2.4 Hz, 1H), 7.15 (dd, *J* = 8.8, 2.4 Hz, 1H), 4.54 (tt, *J* = 10.4, 4.2 Hz, 1H), 3.92 (tdq, *J* = 11.7, 8.0, 3.9 Hz, 1H), 2.12 (d, *J* = 13.8 Hz, 2H), 1.90 (d, *J* = 14.2 Hz, 2H), 1.71 (q, *J* = 13.2 Hz, 2H), 1.52 (q, *J* = 12.9 Hz, 2H). UPLC-MS (ESI) *m*/*z* $[M+H]^+$ 391.11.

***6-(4-(5-acetyl-3-(7-(difluoromethyl)-6-(1-methyl-1H-pyrazol-4-yl)-3,4-dihydroquinolin-1(2H)-yl)-4,5,6,7-tetrahydro-1H-pyrazolo[4,3-c]pyridin-1-yl)piperidin-1-yl)-N-((1r,4r)-4-(3-chloro-4-cyanophenoxy)cyclohexyl)pyridazine-3-carboxamide (1)***

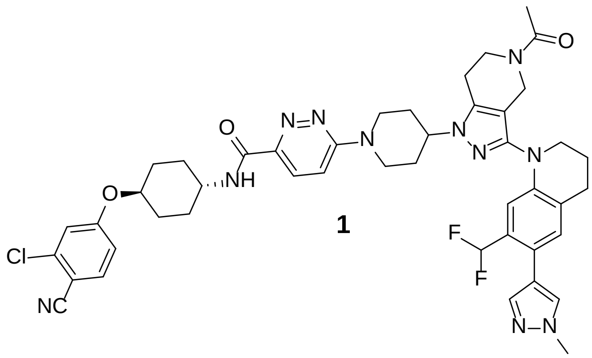

**1**

Step b: To a solution of intermediate **10** (46.9 mg, 0.12 mmol) and **11** (51.0 mg, 0.1 mmol) in DMF (5 mL), was added $K_2CO_3$ (27.6 mg, 0.2 mmol). The mixture was stirred at 65 $^oC$ for 3 h. Monitor the reaction through LC/MS until the intermediate **11** was completely consumed and then the mixture was concentrated under reduced pressure. The crude residue was purified by pre-HPLC (25−100% MeCN/$H_2O$ in 70 min, the desired product started eluting at ~ 46% MeCN) to give the compound **1** (64.8 mg, yield: 75%) as yellow solid. $^1$H NMR (400 MHz, DMSO-$d_6$) δ 8.63 (d, *J* = 8.2 Hz, 1H), 7.89 – 7.81 (m, 2H), 7.75 (s, 1H), 7.49 (s, 1H), 7.46 (d, *J* = 9.9 Hz, 1H), 7.40 (d, *J* = 2.4 Hz, 1H), 7.14 (dd, *J* = 8.7, 2.5 Hz, 1H), 7.09 (s, 1H), 6.78 (td, *J* = 51.6, 6.8 Hz, 1H), 6.77 (s,1H), 4.61 (d, *J* = 13.1 Hz, 2H), 4.57 – 4.44 (m, 2H), 4.39 – 4.22 (m, 1H), 4.15 (d, *J* = 20.4 Hz, 2H), 3.86 (s, 3H), 3.80 – 3.69 (m, 2H), 3.55 (q, *J* = 5.8 Hz, 2H), 3.24 (t, *J* = 25.7 Hz, 2H), 2.95 – 2.74

(m, 4H), 2.17 – 2.04 (m, 4H), 2.04 – 1.81 (m, 9H), 1.64 (q, $J$ = 12.3 Hz, 2H), 1.51 (q, $J$ = 11.7 Hz, 2H). Purity >95.0%. UPLC-MS (ESI) $m/z$ $[M+H]^+$ 864.28.

***1-(1-(1-(azetidin-3-yl)piperidin-4-yl)-3-(7-(difluoromethyl)-6-(1-methyl-1H-pyrazol-4-yl)-3,4-dihydroquinolin-1(2H)-yl)-1,4,6,7-tetrahydro-5H-pyrazolo[4,3-c]pyridin-5-yl)ethan-1-one (13a)***

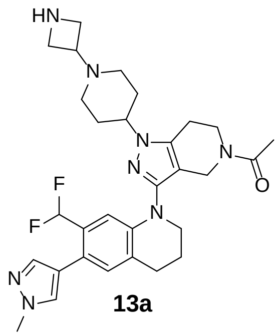

**13a**

Step c: To a solution of intermediate **11** (51.0 mg, 0.1 mmol) and **12a** (34.2 mg, 0.2 mmol) in DCM at rt, was added $Et_3N$ (20.2 mg, 0.2 mmol). The mixture was stirred at rt for 30 min. Then sodium triacetoxyborohydride (31.8 mg, 0.15 mmol) was added to this reaction, and the mixture was stirred at rt for 3 h. Monitor the reaction through LC/MS until the intermediate **11** was completely consumed and then the mixture was concentrated under reduced pressure. The crude residue was purified by pre-HPLC (25−100% MeCN/$H_2O$ in 70 min, the desired product started eluting at ~ 38% MeCN).

Step d: The product obtained above was dissolved in 6 mL of DCM: TFA (5:1) and stirred at 40 $^oC$ for an additional 3 h. Monitor the reaction through LC/MS until the reaction was completed. Then the mixture was concentrated, and purified by pre-HPLC (10−100% MeCN/$H_2O$ in 90 min, the desired product started eluting at ~ 20% MeCN), to give intermediate **13a** (34 mg, yield: 60%, two steps) as a clear, gel-like solid. $^1H$ NMR (400 MHz, DMSO-$d_6$) δ 7.77 (s, 1H), 7.51 (d, $J$ = 0.8 Hz, 1H), 7.13 (s, 1H), 6.80 (td, $J$ = 55.2, 6.8 Hz, 1H), 6.79 (d, $J$ = 3.0 Hz, 1H), 4.44 – 4.24 (m, 3H), 4.23 – 4.03 (m, 5H), 3.87 (s, 3H), 3.79 – 3.67 (m, 2H), 3.59 (t, $J$ = 6.0 Hz, 2H), 3.51 – 3.36 (m ,2H), 3.03 – 2.79 (m, 5H), 2.76 (s, 1H), 2.28 – 2.02 (m, 6H), 2.01 – 1.92 (m, 3H). UPLC-MS (ESI) $m/z$ $[M+H]^+$ 565.35.

***1-(1-([1,4'-bipiperidin]-4-yl)-3-(7-(difluoromethyl)-6-(1-methyl-1H-pyrazol-4-yl)-3,4-dihydroquinolin-1(2H)-yl)-1,4,6,7-tetrahydro-5H-pyrazolo[4,3-c]pyridin-5-yl)ethan-1-one (13b)***

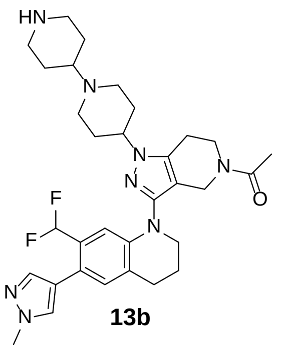

**13b**

Compound **13b** was synthesized from intermediates **11** and **12b** following a similar procedure to that of **13a**. $^1$H NMR (400 MHz, DMSO-$d_6$) δ 7.77 (s, 1H), 7.51 (s, 1H), 7.13 (s, 1H), 6.81 (td, *J* = 55.2, 7.0 Hz, 1H), 6.77 (d, *J* = 3.0 Hz, 1H), 4.51 – 4.39 (s, 1H), 4.14 (d, *J* = 23.0 Hz, 2H), 3.87 (s, 3H), 3.80 – 3.69 (m, 2H), 3.67 – 3.53 m, 4H), 3.53 – 3.37 (m, 6H), 3.22 (q, *J* = 11.0 Hz, 2H), 2.95 (q, *J* = 11.9 Hz, 2H), 2.89 – 2.81 (d, *J* = 5.8 Hz, 3H), 2.79 – 2.72 (m, 1H) , 2.37 – 2.13 (m, 6H), 2.08 (d, *J* = 2.5 Hz, 2H), 1.91 – 1.78 (m, 2H). UPLC-MS (ESI) *m*/*z* $[M+H]^+$ 593.41.

***1-(1-(1-(azetidin-3-ylmethyl)piperidin-4-yl)-3-(7-(difluoromethyl)-6-(1-methyl-1H-pyrazol-4-yl)-3,4-dihydroquinolin-1(2H)-yl)-1,4,6,7-tetrahydro-5H-pyrazolo[4,3-c]pyridin-5-yl)ethan-1-one (13c)***

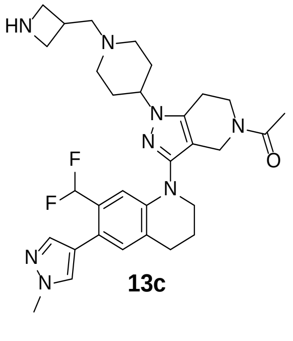

**13c**

Compound **13c** was synthesized from intermediates **11** and **12c** following a similar procedure to that of **13a**. $^1$H NMR (400 MHz, DMSO-$d_6$) δ 7.76 (s, 1H), 7.51 (s, 1H), 7.13 (s, 1H), 6.80 (td, *J* = 55.2, 6.8 Hz, 1H), 6.77 (d, *J* = 3.8 Hz, 1H), 4.44 – 4.33 (m, 1H), 4.14 (d, *J* = 22.3 Hz, 2H), 4.09 – 3.97 (m, 2H), 3.94 – 3.80 (s, 5H), 3.81 – 3.67 (m, 2H), 3.66 –

3.49 (m, 4H), 3.41 (s, 2H), 3.36 – 3.26 (m, 1H), 3.12 (s, 2H), 2.92 – 2.79 (m, 3H), 2.74 (t, *J* = 6.9 Hz, 1H), 2.32 – 2.17 (m, 2H), 2.17 – 2.05 (m, 4H), 2.04 – 1.90 (m, 3H). UPLC-MS (ESI) *m*/*z* $[M+H]^+$ 578.30.

***1-(1-(1-(3-azaspiro[5.5]undecan-9-yl)piperidin-4-yl)-3-(7-(difluoromethyl)-6-(1-methyl-1H-pyrazol-4-yl)-3,4-dihydroquinolin-1(2H)-yl)-1,4,6,7-tetrahydro-5H-pyrazolo[4,3-c]pyridin-5-yl)ethan-1-one (13d)***

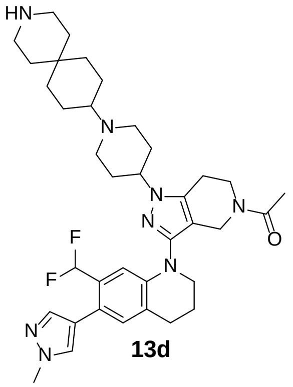

**13d**

Compound **13d** was synthesized from intermediates **11** and **12d** following a similar procedure to that of **13a**. $^1$H NMR (400 MHz, $CD_3OD$) δ 7.65 (s, 1H), 7.52 (s, 1H), 7.11 (d, *J* = 8.5 Hz, 1H), 6.86 – 6.37 (m, 2H), 4.68 – 4.40 (m, 1H), 4.25 (dd, *J* = 23.5, 7.2 Hz, 2H), 3.93 (d, *J* = 1.0 Hz, 3H), 3.86 (dt, *J* = 25.9, 5.8 Hz, 2H), 3.73 – 3.62 (m, 4H), 3.49 – 3.23 (m, 3H), 3.23 – 3.10 (m, 4H), 2.92 (d, *J* = 13.3 Hz, 3H), 2.87 – 2.78 (m, 1H), 2.52 – 2.15 (m, 6H), 2.12 – 1.90 (m, 7H), 1.88 – 1.66 (m, 4H), 1.65 – 1.54 (m, 2H), 1.46 – 1.27 (m, 2H). UPLC-MS (ESI) *m*/*z* $[M+H]^+$ 661.29

***1-(1-(1-((3-azaspiro[5.5]undecan-9-yl)methyl)piperidin-4-yl)-3-(7-(difluoromethyl)-6-(1-methyl-1H-pyrazol-4-yl)-3,4-dihydroquinolin-1(2H)-yl)-1,4,6,7-tetrahydro-5H-pyrazolo[4,3-c]pyridin-5-yl)ethan-1-one (13e)***

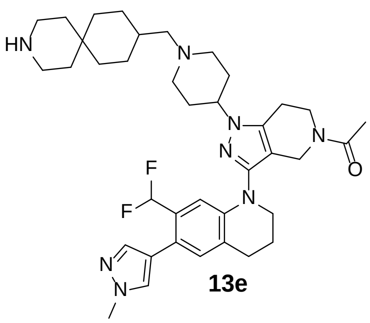

**13e**

Compound **13e** was synthesized from intermediates **11** and **12e** following a similar procedure to that of **13a**.[1]H NMR (400 MHz, $CD_3OD$) δ 7.65 (s, 1H), 7.51 (d, *J* = 0.8 Hz, 1H), 7.12 (d, *J* = 8.6 Hz, 1H), 6.88 – 6.35 (m, 2H), 4.66 – 4.38 (m, 1H), 4.24 (dd, *J* = 20.8, 7.3 Hz, 2H), 3.93 (d, *J* = 1.3 Hz, 3H), 3.86 (dt, *J* = 25.7, 5.8 Hz, 2H), 3.80 – 3.61 (m, 4H), 3.26 – 3.04 (m, 8H), 2.99 – 2.79 (m, 4H), 2.56 – 2.15 (m, 6H), 2.13 – 2.00 (m, 3H), 1.96 – 1.65 (m, 7H), 1.64 – 1.54 (m, 2H), 1.39 – 1.19 (m, 4H). UPLC-MS (ESI) *m*/*z* $[M+H]^+$ 675.33.

***6-(3-(4-(5-acetyl-3-(7-(difluoromethyl)-6-(1-methyl-1H-pyrazol-4-yl)-3,4-dihydroquinolin-1(2H)-yl)-4,5,6,7-tetrahydro-1H-pyrazolo[4,3-c]pyridin-1-yl)piperidin-1-yl)azetidin-1-yl)-N-((1r,4r)-4-(3-chloro-4-cyanophenoxy)cyclohexyl)pyridazine-3-carboxamide (2)***

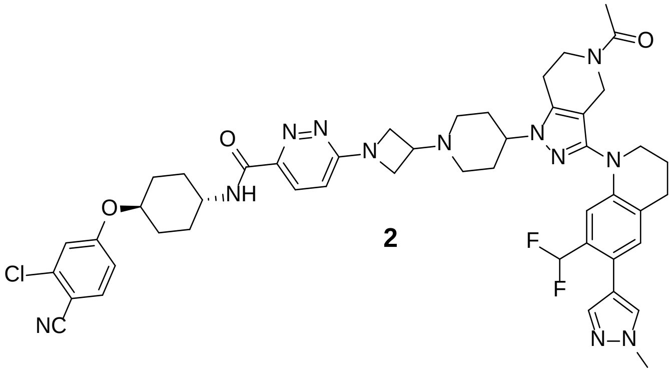

**2**

Step e: To a solution of intermediate **13a** (28.3 mg, 0.05 mmol) and **10** (23.4 mg, 0.06 mmol) in DMF (5 mL), was added $K_2CO_3$ (13.8 mg, 0.1 mmol). The mixture was stirred at 65 °C for 3 h. Monitor the reaction through LC/MS until the intermediate **13a** was completely consumed and then the mixture was concentrated under reduced pressure. The crude residue was purified by pre-HPLC (25−100% MeCN/$H_2O$ in 70 min, the desired product started eluting at ~ 40% MeCN) to give the compound **2** (36.5 mg, yield: 80%) as yellow solid. [1]H NMR (400 MHz, DMSO-*d*6) δ 8.65 (d, *J* = 8.2 Hz, 1H), 7.95 (d, *J* = 9.2 Hz, 1H), 7.87 (d, *J* = 8.8 Hz, 1H), 7.76 (s, 1H), 7.50 (s, 1H), 7.39 (d, *J* = 2.4 Hz, 1H), 7.14 (dd, *J* = 8.8, 2.4 Hz, 2H), 7.00 (d, *J* = 9.2 Hz, 1H), 6.81 (td, *J* = 51.6, 6.8 Hz, 1H), 6.79 (d, *J* = 3.7 Hz, 1H), 4.61 – 4.25 (m, 7H), 4.15 (d, *J* = 21.6 Hz, 2H), 3.93 – 3.80 (m, 4H), 3.81 – 3.40 (m, 6H), 3.23 – 3.05 (m, 2H), 2.95 – 2.73 (m, 4H), 2.34 – 2.15 (m,

4H), 2.15 – 2.05 (m, 4H), 2.02 – 1.93 (m, 3H), 1.90 (d, *J* = 13.1 Hz, 2H), 1.65 (q, *J* = 12.2 Hz, 2H), 1.52 (q, *J* = 11.5 Hz, 2H). Purity >95.0%, UPLC-MS (ESI) *m*/*z* $[M+H]^+$ 919.36.

***6-(4-(5-acetyl-3-(7-(difluoromethyl)-6-(1-methyl-1H-pyrazol-4-yl)-3,4-dihydroquinolin-1(2H)-yl)-4,5,6,7-tetrahydro-1H-pyrazolo[4,3-c]pyridin-1-yl)-[1,4'-bipiperidin]-1'-yl)-N-((1r,4r)-4-(3-chloro-4-cyanophenoxy)cyclohexyl)pyridazine-3-carboxamide (3)***

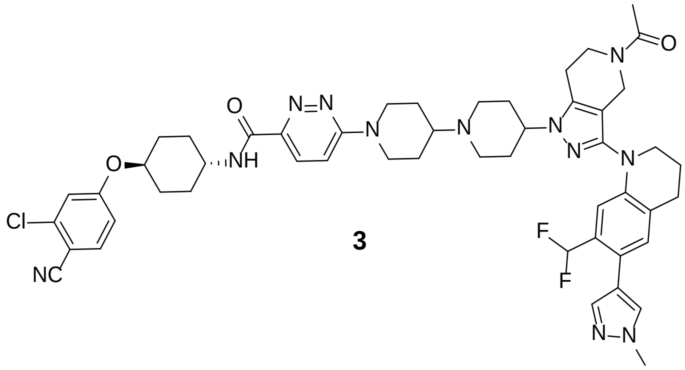

**3**

Compound **3** was synthesized from intermediates **13b** and **10** following a similar procedure to that of **2**. $^1H$ NMR (400 MHz, DMSO-$d_6$) δ 8.65 (d, *J* = 8.2 Hz, 1H), 7.88 (dd, *J* = 9.1, 5.8 Hz, 2H), 7.76 (s, 1H), 7.51 (s, 1H), 7.46 (d, *J* = 9.6 Hz, 1H), 7.40 (d, *J* = 2.4 Hz, 1H), 7.17 – 7.10 (m, 2H), 6.81 (td, *J* = 51.6, 7.2 Hz, 1H), 6.77 (d, *J* = 3.7 Hz, 1H), 4.68 (d, *J* = 13.1 Hz, 2H), 4.54 (q, *J* = 9.5, 7.6 Hz, 1H), 4.49 – 4.39 (m, 1H), 4.14 (d, *J* = 22.3 Hz, 2H), 3.93 – 3.81 (m, 4H), 3.80 – 3.68 (m, 2H), 3.59 (q, *J* = 6.2, 5.6 Hz, 5H), 3.21(q, *J* = 10.3 Hz, 2H), 3.04 (t, *J* = 12.7 Hz, 2H), 2.92 – 2.71 (m, 4H), 2.30 (d, *J* = 12.8 Hz, 2H), 2.23 – 2.05 (m, 8H), 2.05 – 1.84 (m, 5H), 1.65 (q, *J* = 12.3, 11.5 Hz, 4H), 1.52 (q, *J* = 11.6 Hz, 2H). Purity >95.0%. UPLC-MS (ESI) *m*/*z* $[M+H]^+$ 947.51.

***6-(3-((4-(5-acetyl-3-(7-(difluoromethyl)-6-(1-methyl-1H-pyrazol-4-yl)-3,4-dihydroquinolin-1(2H)-yl)-4,5,6,7-tetrahydro-1H-pyrazolo[4,3-c]pyridin-1-yl)piperidin-1-yl)methyl)azetidin-1-yl)-N-((1r,4r)-4-(3-chloro-4-cyanophenoxy)cyclohexyl)pyridazine-3-carboxamide (4)***

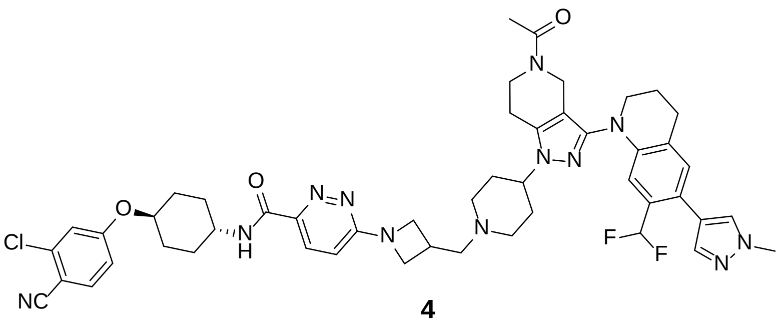

**4**

Compound **4** was synthesized from intermediates **13c** and **10** following a similar procedure to that of **2**. $^1$H NMR (400 MHz, DMSO-$d_6$) δ 8.59 (d, *J* = 7.4 Hz, 1H), 7.88 (dd, *J* = 9.1, 7.7 Hz, 2H), 7.77 (s, 1H), 7.51 (s, 1H), 7.39 (d, *J* = 2.4 Hz, 1H), 7.17 – 7.10 (m, 2H), 6.93 (d, *J* = 9.7 Hz, 1H), 6.82 (td, *J* = 51.6, 7.2 Hz, 1H), 6.82 – 6.77 (m, 1H), 4.59 – 4.49 (m, 1H), 4.45 – 4.36 (m, 1H), 4.32 (t, *J* = 8.5 Hz, 2H), 4.16 (d, *J* = 21.8 Hz, 2H), 3.97 (t, *J* = 7.3 Hz, 2H), 3.91 – 3.81 (m, 4H), 3.75 (d, *J* = 17.6 Hz, 2H), 3.67 – 3.56 (m, 4H), 3.56 – 3.41 (m, 2H), 3.39 – 3.27 (m,1H), 3.17 (q, *J* = 11.7 Hz, 2H), 2.92 – 2.73 (m, 4H), 2.30 (q, *J* = 13.5 Hz, 2H), 2.15 (d, *J* = 15.9 Hz, 3H), 2.09 (s, 3H), 2.02 – 1.93 (m, 2H), 1.90 (d, *J* = 12.1 Hz, 2H), 1.65 (q, *J* = 12.2 Hz, 2H), 1.52 (q, *J* = 12.1, 11.7 Hz, 2H). Purity >95.0%. Purity >95.0%. UPLC-MS (ESI) *m/z* $[M+H]^+$ 933.38.

***6-(9-(4-(5-acetyl-3-(7-(difluoromethyl)-6-(1-methyl-1H-pyrazol-4-yl)-3,4-dihydroquinolin-1(2H)-yl)-4,5,6,7-tetrahydro-1H-pyrazolo[4,3-c]pyridin-1-yl)piperidin-1-yl)-3-azaspiro[5.5]undecan-3-yl)-N-((1r,4r)-4-(3-chloro-4-cyanophenoxy)cyclohexyl)pyridazine-3-carboxamide (5)***

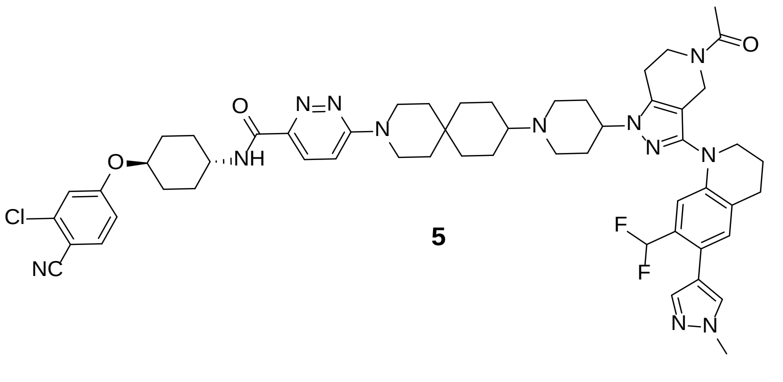

**5**

Compound **5** was synthesized from intermediates **13d** and **10** following a similar procedure to that of **2**. $^1$H NMR (400 MHz, DMSO-$d_6$) δ 8.60 (d, *J* = 8.1 Hz, 1H), 7.87 (d, *J* = 8.8 Hz, 1H), 7.83 (d, *J* = 9.5 Hz, 1H), 7.77 (s, 1H), 7.51 (s, 1H), 7.40 (d, *J* = 2.4 Hz,

1H), 7.37 (d, *J* = 9.6 Hz, 1H), 7.14 (dd, *J* = 8.8, 2.4 Hz, 2H), 6.82 (td, *J* = 51.6, 7.2 Hz, 1H), 6.81 – 6.76 (m, 1H), 4.54 (p, *J* = 5.6 Hz, 1H), 4.49 – 4.39 (m, 1H), 4.15 (d, *J* = 21.8 Hz, 2H), 3.92 – 3.82 (m, 4H), 3.80 – 3.65 (m, 6H), 3.60 (q, *J* = 5.1 Hz, 4H), 3.30 – 3.13 (m, 3H), 2.92 – 2.72 (m, 4H), 2.32 (d, *J* = 14.4 Hz, 2H), 2.21 – 2.05 (m, 6H), 2.02 – 1.94 (m, 3H), 1.94 – 1.78 (m, 6H), 1.73 – 1.57 (m, 6H), 1.53 (t, *J* = 11.4 Hz, 2H), 1.41 (s, 2H), 1.29 – 1.13 (m, 2H). Purity >95.0%. UPLC-MS (ESI) *m*/*z* $[M+H]^+$ 1015.59.

***6-(9-((4-(5-acetyl-3-(7-(difluoromethyl)-6-(1-methyl-1H-pyrazol-4-yl)-3,4-dihydroquinolin-1(2H)-yl)-4,5,6,7-tetrahydro-1H-pyrazolo[4,3-c]pyridin-1-yl)piperidin-1-yl)methyl)-3-azaspiro[5.5]undecan-3-yl)-N-((1r,4r)-4-(3-chloro-4-cyanophenoxy)cyclohexyl)pyridazine-3-carboxamide (6)***

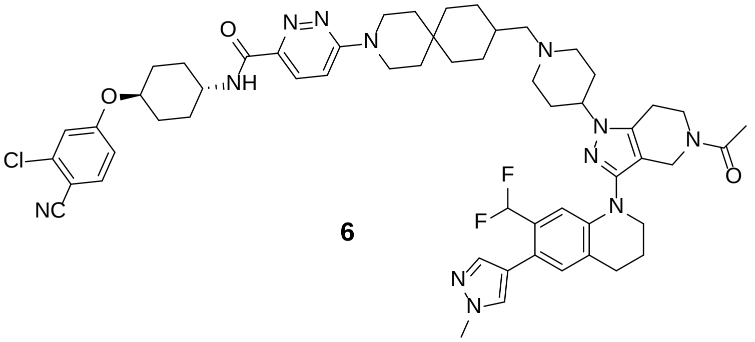

**6**

Compound **6** was synthesized from intermediates **13e** and **10** following a similar procedure to that of **2**. $^1$H NMR (400 MHz, DMSO-$d_6$) δ 8.58 (d, *J* = 8.2 Hz, 1H), 7.86 (d, *J* = 8.8 Hz, 1H), 7.83 (d, *J* = 9.6 Hz, 1H), 7.76 (s, 1H), 7.51 (s, 1H), 7.39 (d, *J* = 2.4 Hz, 1H), 7.36 (d, *J* = 9.7 Hz, 1H), 7.14 (dd, *J* = 8.8, 2.4 Hz, 2H), 6.81 (td, *J* = 51.6, 6.6 Hz, 1H), 6.78 (d, *J* = 2.6 Hz, 1H), 4.60 – 4.49 (m, 1H), 4.44 – 4.33 (s, 1H), 4.15 (d, *J* = 20.6 Hz, 2H), 3.92 – 3.80 (s, 4H), 3.79 – 3.39 (m, 10H), 3.20 – 3.04 (m, 2H), 3.01 (s, 2H), 2.93 – 2.72 (m, 4H), 2.36 (q, *J* = 14.4 Hz, 2H), 2.23 – 2.04 (m, 6H), 2.02 – 1.94 (m, 2H), 1.90 (d, *J* = 11.9 Hz, 2H), 1.85 – 1.70 (m, 3H), 1.71 – 1.45 (m, 8H), 1.41 (s, 2H), 1.27 – 1.10 (m, 4H). Purity >95.0%. UPLC-MS (ESI) *m*/*z* $[M+H]^+$ 1029.58.

### Scheme 2: Synthesis of Compound 7 and DALTAC-1 (JZY3032).

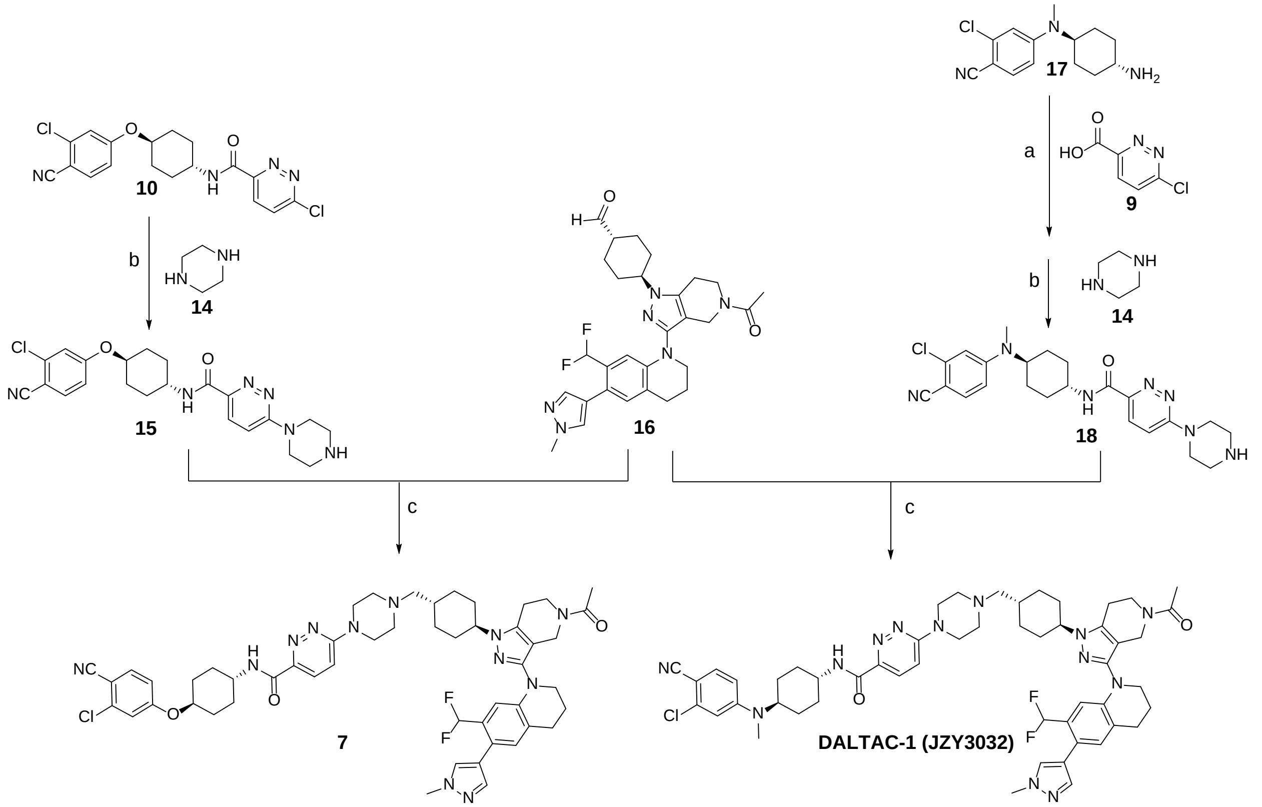

***Reagents and conditions*:** (a) HATU, DMF, $Et_3N$, rt, 15 min; (b) DIPEA, DMF, 55 °C, 30 min; (c) $Et_3N$, Sodium triacetoxyborohydride, DCM, rt, 3 h.

Intermediate **16** was prepared according to the reported procedures.[1]

Intermediate **17** was prepared according to the reported procedures.[2]

***N-((1r,4r)-4-((3-chloro-4-cyanophenyl)(methyl)amino)cyclohexyl)-6-(piperazin-1-yl)pyridazine-3-carboxamide (18)***

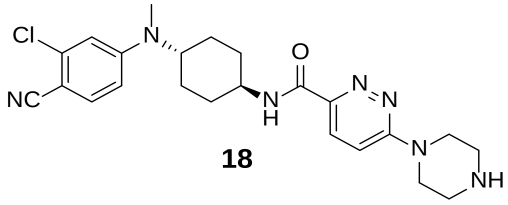

**18**

Step a: 6-chloropyridazine-3-carboxylic acid **9** (5.0 g, 31.54 mmol) was dissolved in DMF (100 mL), HATU (12.0 g, 31.54) and $Et_3N$ (4.8 g, 47.31 mmol) was added. The mixture was stirred at room temperature for 2 min, then intermediate **17** (8.32 g, 31.54 mmol) was added in to the mixture. The result mixture was stirred at rt for additional 13 min, after that the mixture was diluted with water and extracted with DCM. The organic phases were combined and concentrated under reduced pressure. The crude residue was used directly in the next step without further purification.

Step b: The residue obtained above was redissolved in DMF (80 mL), and then piperazine **14** (9.5 g, 110.39 mmol) was added. The mixture was stirred at 55 °C for 30 min. After that all volatile materials were removed in a rotary evaporator and purified by silica gel column chromatography (0–20% MeOH/DCM) to afford intermediate **18** (8.6 g, yield: 60%, two steps). $^1$H NMR (400 MHz, DMSO-$d_6$) δ 8.59 (d, *J* = 8.3 Hz, 1H), 7.94 (d, *J* = 9.5 Hz, 1H), 7.62 (d, *J* = 9.0 Hz, 1H), 7.46 (d, *J* = 9.6 Hz, 1H), 6.95 (d, *J* = 2.5 Hz, 1H), 6.83 (dd, *J* = 9.1, 2.6 Hz, 1H), 3.96 (t, *J* = 5.3 Hz, 4H), 3.91 – 3.84 (m, 1H), 3.83 – 3.71 (m, 1H), 3.60 (q, *J* = 4.7 Hz, 1H), 3.23 (t, *J* = 5.3 Hz, 4H), 2.86 (s, 3H), 1.92 (d, *J* = 10.6 Hz, 2H), 1.82 – 1.63 (m, 6H).

***N-((1r,4r)-4-(3-chloro-4-cyanophenoxy)cyclohexyl)-6-(piperazin-1-yl)pyridazine-3-carboxamide (15)***

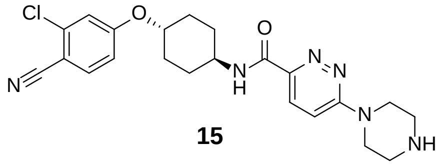

**15**

Intermediate **15** was synthesized with a similar procedure to that of intermediate **18**. $^1$H NMR (400 MHz, DMSO-$d_6$) δ 8.68 (d, *J* = 8.3 Hz, 1H), 7.89 (dd, *J* = 17.4, 9.1 Hz, 2H), 7.43 (d, *J* = 9.6 Hz, 1H), 7.40 (d, *J* = 2.4 Hz, 1H), 7.14 (dd, *J* = 8.8, 2.5 Hz, 1H), 4.53 (td, *J* = 10.1, 5.0 Hz, 1H), 3.86 (t, *J* = 5.2 Hz, 4H), 3.74 (s, 1H), 3.21 – 3.09 (m, 3H), 2.18 – 2.06 (m, 2H), 1.95 – 1.85 (m, 2H), 1.72 – 1.59 (m, 2H), 1.59 – 1.45 (m, 2H).

***6-(4-(((1r,4r)-4-(5-acetyl-3-(7-(difluoromethyl)-6-(1-methyl-1H-pyrazol-4-yl)-3,4-dihydroquinolin-1(2H)-yl)-4,5,6,7-tetrahydro-1H-pyrazolo[4,3-c]pyridin-1-yl)cyclohexyl)methyl)piperazin-1-yl)-N-((1r,4r)-4-((3-chloro-4-cyanophenyl)(methyl)amino)cyclohexyl)pyridazine-3-carboxamide (DALTAC-1)***

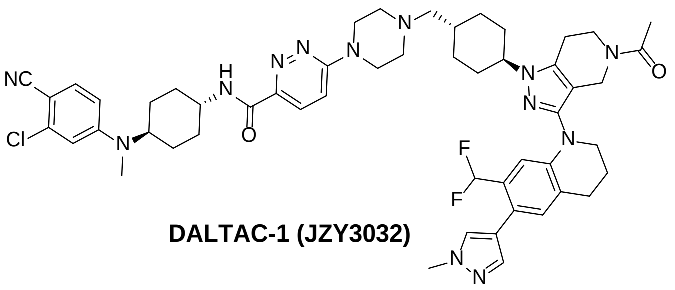

**DALTAC-1 (JZY3032)**

Step c: To a solution of intermediate **16** (54 mg, 0.1 mmol) and **18** (59 mg, 0.13 mmol) in DCM at rt, was added $Et_3N$ (20.2 mg, 0.2 mmol). The mixture was stirred at rt for 30 min. Then sodium triacetoxyborohydride (31.8 mg, 0.15 mmol) was added to this reaction, and the mixture was stirred at rt for 3 h. Monitor the reaction through LC/MS until the intermediate **16** was completely consumed and then the mixture was concentrated under reduced pressure. The crude residue was purified by pre-HPLC (20−100% MeCN/$H_2O$ in 60 min, the desired product started eluting at ~ 41% MeCN) to give **DALTAC-1** (63 mg, yield: 65%). $^1H$ NMR (400 MHz, DMSO-$d_6$) δ 8.50 (d, $J$ = 8.3 Hz, 1H), 7.84 (d, $J$ = 9.4 Hz, 1H), 7.76 (s, 1H), 7.61 (d, $J$ = 9.0 Hz, 1H), 7.50 (d, $J$ = 0.8 Hz, 1H), 7.35 (d, $J$ = 9.7 Hz, 1H), 7.11 (s, 1H), 6.95 (d, $J$ = 2.6 Hz, 1H), 6.83 (dd, $J$ = 9.1, 2.5 Hz, 1H), 6.81 – 6.79 (m, 1H), 6.78 (td, $J$ = 55.2, 6.8 Hz, 1H), 4.15 (d, $J$ = 17.2 Hz, 2H), 4.07 – 3.96 (m, 1H), 3.87 (s, 3H), 3.86 – 3.64 (m, 8H), 3.63 – 3.55 (m, 2H), 2.92 – 2.78 (m, 7H), 2.74 (s, 1H), 2.48 (s, 2H), 2.19 (s, 2H), 2.08 (s, 2H), 2.03 – 1.87 (m, 10H), 1.84 (d, $J$ = 11.8 Hz, 2H), 1.79 – 1.57 (m, 7H), 1.16 – 1.00 (m, 2H). $^{13}C$ NMR (101 MHz, DMSO-$d_6$) δ 169.27, 162.80, 160.56, 153.46, 148.01, 145.27, 142.36, 138.27, 137.69, 137.12, 135.18, 131.09, 129.77, 128.88 (t, J = 20.6 Hz, 1C), 126.81 (d, J = 63.9 Hz, 1C), 121.05, 118.95, 118.21, 114.25 (t, J = 235.9 Hz, 1C), 113.10, 112.33, 111.32, 110.29, 106.12, 105.87, 96.35, 64.74, 57.52, 56.24 (2C), 53.18, 49.57, 47.75 (2C), 44.81, 43.31, 39.03 (2C), 38.46, 32.22, 31.59 (2C), 31.45 (2C), 30.36, 28.20 (2C), 27.38, 22.47, 22.17, 22.12, 21.85. Purity >95.0%. UPLC-MS (ESI) $m/z$ $[M+H]^+$ 974.46.

***6-(4-(((1r,4r)-4-(5-acetyl-3-(7-(difluoromethyl)-6-(1-methyl-1H-pyrazol-4-yl)-3,4-dihydroquinolin-1(2H)-yl)-4,5,6,7-tetrahydro-1H-pyrazolo[4,3-c]pyridin-1-yl)cyclohexyl)methyl)piperazin-1-yl)-N-((1r,4r)-4-(3-chloro-4-cyanophenoxy)cyclohexyl)pyridazine-3-carboxamide (7)***

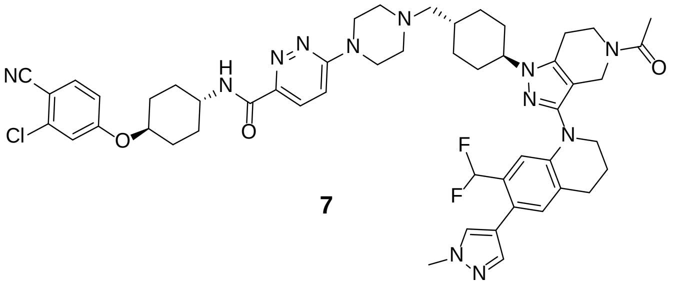

**7**

Compound **7** was synthesized from intermediates **15** and **16** following a similar procedure to that of **DALTAC-1**. $^1$H NMR (400 MHz, DMSO-d6) δ 8.70 (d, J = 8.1 Hz, 1H), 7.96 (d, J = 9.5 Hz, 1H), 7.87 (d, J = 8.8 Hz, 1H), 7.76 (s, 1H), 7.52 (d, J = 8.5 Hz, 2H), 7.40 (d, J = 2.4 Hz, 1H), 7.14 (dd, J = 8.8, 2.5 Hz, 1H), 7.11 (s, 1H), 6.81(td, J = 54.8, 7.1 Hz, 1H), 6.79 (d, J = 2.4 Hz, 1H), 4.63 – 4.44 (m, 4H), 4.16 (d, J = 17.9 Hz, 2H), 4.10 – 4.00 (m, 1H), 3.94 – 3.80 (m, 4H), 3.78 – 3.64 (m, 4H), 3.58 (t, J = 5.7 Hz, 2H), 3.45 (t, J = 12.7 Hz, 2H), 3.24 – 2.97 (m, 4H), 2.89 – 2.72 (m, 4H), 2.09 (s, 4H), 2.01 – 1.84 (m, 11H), 1.66 (q, J = 12.2, 11.8 Hz, 2H), 1.52 (q, J = 11.3 Hz, 2H), 1.29 – 1.13 (m, 2H). Purity >95.0%. UPLC-MS (ESI) *m/z* $[M+H]^+$ 961.37.

### Scheme 3: Synthesis of Neg-1.

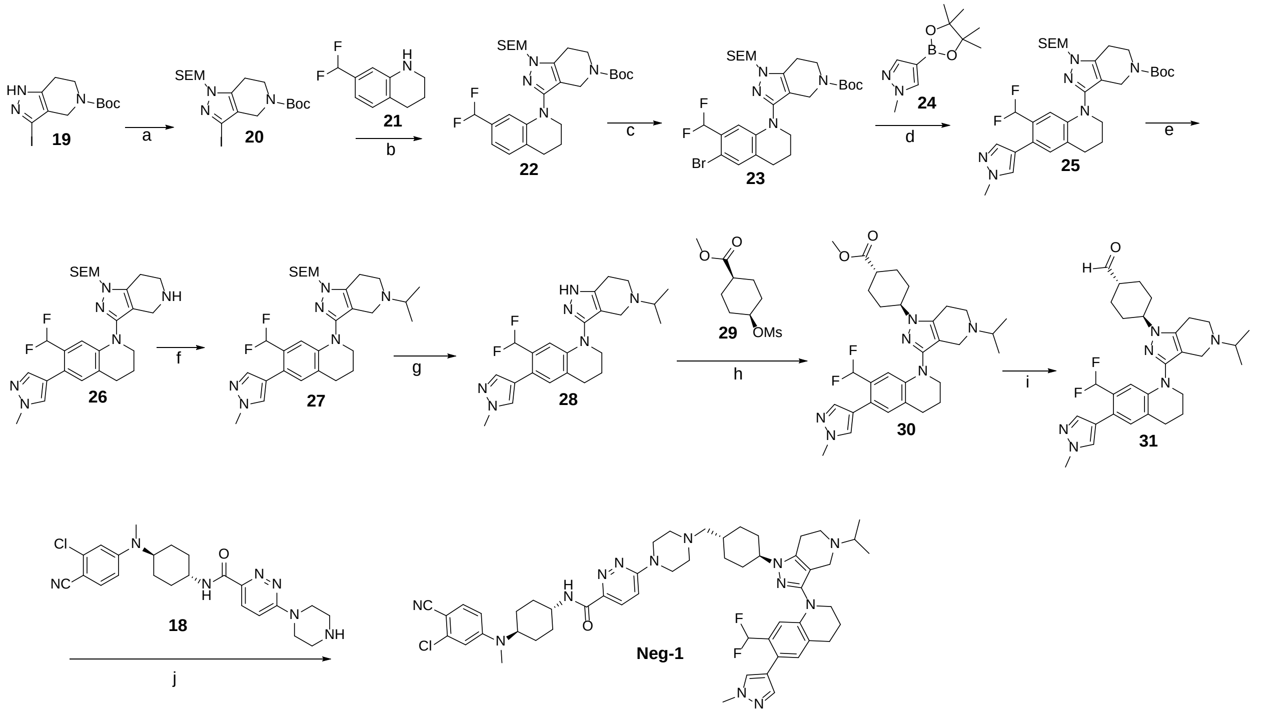

[a]***Reagents and conditions***: (a) 2-(Trimethylsilyl)ethoxymethyl chloride, NaH, DMF, 0 °C to rt, 3 h; (b) 1,4-Dioxane, Ruphos pd G2, Ruphos, Sodium *tert*-butoxide, 100 °C, $N_2$, overnight; (c) NBS, DCM, 0 °C, 30 min; (d) $Pd(OAc)_2$, $Pph_3$ ,$K_2CO_3$, 1,4-Dioxane : $H_2O$ (5:2), 100 °C, $N_2$, 3 h; (e) Formic acid, rt, 4 h; (f) Acetone, Sodium triacetoxyborohydride, DCM, rt, 30 min; (g) Tetrabutylammonium fluoride solution (1 M, in THF), THF, reflux, overnight; (h) $Cs_2CO_3$, DMF, 80 °C, overnight, (i) Diisobutylaluminium hydride (25% in toluene), DCM, −78 °C, 40% yield; (j) $Et_3N$, Sodium triacetoxyborohydride, DCM, rt, 2 h.

##### *tert-butyl 3-iodo-1-((2-(trimethylsilyl)ethoxy)methyl)-1,4,6,7-tetrahydro-5H-pyrazolo[4,3-c]pyridine-5-carboxylate (20)*

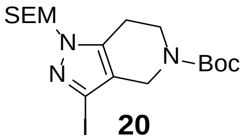

**20**

Step a: A solution of *tert*-butyl 3-iodo-1,4,6,7-tetrahydro-5*H*-pyrazolo[4,3-c]pyridine-5-carboxylate **19** (25.0 g, 71.6 mmol) in anhydrous DMF (300 mL) at 0 °C, was added NaH (3.4 g, 85.9 mmol) in portions. After stirred at 0 °C for 30 min, 2-(Trimethylsilyl)ethoxymethyl chloride (14.3 g, 85.9 mmol) was slowly added into the mixture. Afte that the mixture was warm to rt and stirred for additional 2.5 h. Monitor the reaction through LC/MS until the intermediate **19** was completely consumed and then the mixture was quenched with ice water and extracted with EtOAc. The organic phases were combined and concentrated under reduced pressure. The residue was purified by flash column chromatography (0–60% EtOAc/*n*-hexane) to give intermediate **20** (27.5 g, yield: 80%) as white solid. $^1$H NMR (400 MHz, DMSO-$d_6$) δ 5.35 (d, $J$ = 16.3 Hz, 2H), 4.16 (d, $J$ = 26.6 Hz, 2H), 3.61 (t, $J$ = 5.8 Hz, 2H), 3.58 – 3.46 (m, 2H), 2.65 (dt, $J$ = 37.0, 5.8 Hz, 2H), 1.42 (d, $J$ = 1.3 Hz, 9H), 0.89 – 0.75 (m, 2H), -0.04 (d, $J$ = 5.2 Hz, 9H).

##### *tert-butyl 3-(7-(difluoromethyl)-3,4-dihydroquinolin-1(2H)-yl)-1-((2-(trimethylsilyl)ethoxy)methyl)-1,4,6,7-tetrahydro-5H-pyrazolo[4,3-c]pyridine-5-carboxylate (22)*

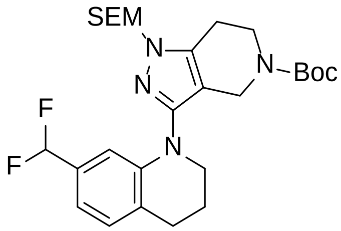

**22**

Step b: A mixture of Intermediate **20** (4.8 g, 10 mmol), 7-(difluoromethyl)-1,2,3,4-tetrahydroquinoline **21** (2.2 g, 12 mmol), RuPhos Pd G2 (777 mg, 1.0 mmol), RuPhos (447 mg, 1.0 mmol), and Sodium *tert*-butoxide (2.4 g, 25 mmol) in 1,4-dioxane (120 mL) was degassed and purged 3 times with $N_2$, and then the mixture was stirred at 100 °C for 12 h. After that the mixture was cooled, diluted with DCM, filtered through Celite, the filter cake was washed with DCM, and the filtrate was concentrated under reduced pressure. The residue was purified by flash column chromatography (0–5% MeOH/DCM) to give intermediate **22** (3.4 g, yield: 64%) as yellow solid. $^1$H NMR (400 MHz, DMSO-$d_6$) δ 7.14 (d, *J* = 7.7 Hz, 1H), 6.83 (d, *J* = 7.3 Hz, 2H), 6.80 (td, *J* = 55.2, 6.8 Hz, 2H), 6.60 (s, 1H), 5.30 (s, 2H), 3.98 (s, 2H), 3.65 – 3.52 (m, 6H), 2.79 (dt, *J* = 28.3, 6.1 Hz, 4H), 1.95 (p, *J* = 6.2 Hz, 2H), 1.36 (s, 9H), 0.83 (t, *J* = 7.2 Hz, 2H), -0.06 (s, 9H).

***tert-butyl 3-(6-bromo-7-(difluoromethyl)-3,4-dihydroquinolin-1(2H)-yl)-1-((2-(trimethylsilyl)ethoxy)methyl)-1,4,6,7-tetrahydro-5H-pyrazolo[4,3-c]pyridine-5-carboxylate (23)***

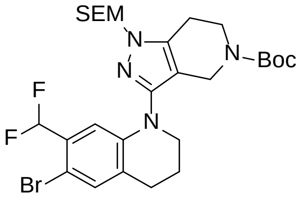

**23**

Step c: To a solution of intermediate **22** (3.0 g, 5.6 mmol) in DCM (80 mL) was added NBS (997 mg, 5.6 mmol) in portions under ice-bath condition. After 30 min, the reaction mixture was diluted with DCM, washed with aqueous $Na_2S_2O_3$ followed by brine, the organic layers were concentrated under reduced pressure. The resulting residue was purified by flash column chromatography (0−5% MeOH/DCM) to give intermediate **23** (2.8 g, yield: 82%) as a yellow solid. $^1$H NMR (400 MHz, DMSO-$d_6$) δ 7.37 (s, 1H), 6.92 (t, *J* =

54.2 Hz, 1H), 6.73 (s, 1H), 5.31 (s, 2H), 4.02 (s, 2H), 3.64 – 3.53 (m, 6H), 2.87 – 2.73 (dt, *J* = 28.3, 6.1 Hz, 4H), 1.94 (p, *J* = 6.2 Hz, 2H), 1.37 (s, 9H), 0.82 (t, *J* = 7.2, 2H), -0.06 (s, 9H).

***tert-butyl 3-(7-(difluoromethyl)-6-(1-methyl-1H-pyrazol-4-yl)-3,4-dihydroquinolin-1(2H)-yl)-1-((2-(trimethylsilyl)ethoxy)methyl)-1,4,6,7-tetrahydro-5H-pyrazolo[4,3-c]pyridine-5-carboxylate (25)***

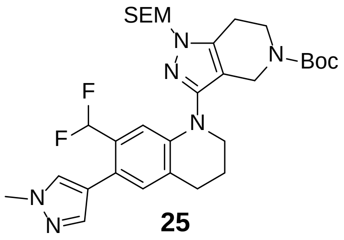

**25**

Step d: A mixture of intermediate **23** (1.4 g, 2.28 mmol), 1-methyl-4-(4,4,5,5-tetramethyl-1,3,2-dioxaborolan-2-yl)-1H-pyrazole **24** (570.0 mg, 2.74 mmol), $Pd(OAc)_2$ (51.1 mg, 0.23 mmol), $Pph_3$ (119.5 mg, 0.46) and $K_2CO_3$ (629.3 mg, 4.56 mmol) were added into 50 mL of 1,4-Dioxane : $H_2O$ (5:2). The mixture was degassed and purged 3 times with $N_2$, then stirred at 100 °C for 3 h. After that the mixture was cooled, filtered through Celite, the filter cake was washed with DCM, and the filtrate was concentrated under reduced pressure. The residue was purified by flash column chromatography (0−5% MeOH/DCM) to give intermediate **25** as white solid (1.1 g, yield: 80%)**.** $^1$H NMR (400 MHz, DMSO-$d_6$) δ 9.06 (s, 1H), 7.79 – 7.74 (m, 1H), 7.53 – 7.48 (m, 1H), 7.17 – 7.11 (m, 1H), 6.83 – 6.78 (m, 1H), 6.76 (td, *J* = 55.2, 6.8 Hz, 1H) 5.34 (d, *J* = 24.6 Hz, 2H), 4.02 (s, 2H), 3.88 (d, *J* = 2.1 Hz, 3H), 3.64 – 3.53 (m, 6H), 2.81 (dt, *J* = 30.9, 6.1 Hz, 4H), 1.97 (t, *J* = 5.8 Hz, 2H), 1.36 (s, 9H), 0.90 – 0.79 (m, 2H), -0.01 – -0.06 (m, 9H).

***7-(difluoromethyl)-6-(1-methyl-1H-pyrazol-4-yl)-1-(1-((2-(trimethylsilyl)ethoxy)methyl)-4,5,6,7-tetrahydro-1H-pyrazolo[4,3-c]pyridin-3-yl)-1,2,3,4-tetrahydroquinoline (26)***

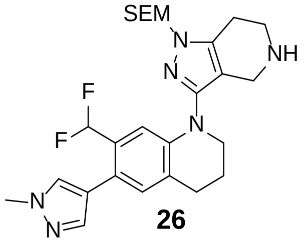

**26**

Step e: Intermediate **25** (1.0 g, 1.63 mmol) was added into formic acid (8 mL). The mixture was stirred at rt for 4 h. The reaction was monitored by LC-MS until completion. The mixture was concentrated and purified by pre-HPLC to give intermediate **26** (587 mg, yield: 70%). $^{1}$H NMR (400 MHz, DMSO-$d_6$) δ 7.78 (s, 1H), 7.51 (t, $J$ = 1.7 Hz, 1H), 7.16 (s, 1H), 6.96 – 6.66 (m, 2H), 5.37 (s, 2H), 3.84 (s, 2H), 3.63 – 3.56 (m, 4H), 3.47 – 3.37 (m, 2H), 3.03 (t, $J$ = 6.0 Hz, 2H), 2.85 (t, $J$ = 6.4 Hz, 2H), 1.98 (q, $J$ = 5.9 Hz, 2H), 0.87 (t, $J$ = 8.6, 2H), -0.015 (s, 9H).

***7-(difluoromethyl)-1-(5-isopropyl-1-((2-(trimethylsilyl)ethoxy)methyl)-4,5,6,7-tetrahydro-1H-pyrazolo[4,3-c]pyridin-3-yl)-6-(1-methyl-1H-pyrazol-4-yl)-1,2,3,4-tetrahydroquinoline (27)***

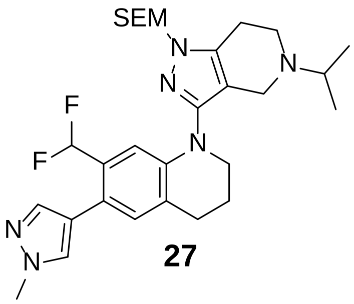

**27**

Step f: A solution of intermediate **26** (514 mg, 1.0 mmol) in DCM (5 mL) at rt, was added acetone (580 mg, 10.0 mmol). The mixture was stirred at rt for 30 min. Then sodium triacetoxyborohydride (318 mg, 1.5 mmol) was added to this reaction, and the mixture was stirred at rt for 3 h. Monitor the reaction through LC/MS until the intermediate **26** was completely consumed and then the mixture was concentrated under reduced pressure. The crude residue was purified by pre-HPLC to give intermediate **27** (456 mg, yield: 82%). $^{1}$H NMR (400 MHz, DMSO-$d_6$) δ 7.78 (s, 1H), 7.52 (d, $J$ = 0.8 Hz, 1H), 7.16 (s, 1H), 6.87 (s, 1H), 6.82 (td, $J$ = 55.1 Hz, 6.8 Hz, 1H), 5.46 – 5.31 (m, 2H), 4.01 – 3.95 (m, 2H), 3.94 – 3.89 (m, 1H), 3.88 (s, 3H), 3.81 – 3.73 (m, 1H), 3.72 – 3.52 (m, 4H), 3.40 – 3.28 (m, 1H), 3.23 – 3.15 (m, 1H), 3.12 – 3.00 (m, 1H), 2.86 (t, $J$ = 6.4 Hz, 2H), 1.99 (p, $J$ = 6.1 Hz, 2H), 1.27 (dd, $J$ = 10.6, 6.5 Hz, 6H), 0.88 (dt, $J$ = 8.9, 6.6 Hz, 2H), -0.06 (s,9H).

***7-(difluoromethyl)-1-(5-isopropyl-4,5,6,7-tetrahydro-1H-pyrazolo[4,3-c]pyridin-3-yl)-6-(1-methyl-1H-pyrazol-4-yl)-1,2,3,4-tetrahydroquinoline (28)***

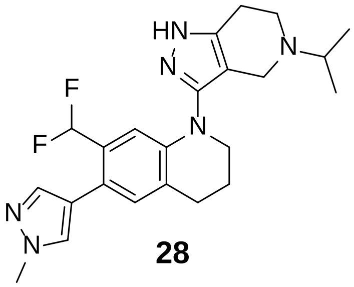

**28**

Step g: Intermediate **27** (400 mg, 0.72 mmol) was dissolved in THF (8 mL). Followed by 1 mL of tetrabutylammonium fluoride solution (1 M, in THF) was added into the mixture. The mixture was stirred and refluxed at 66 °C for overnight. Monitor the reaction through LC/MS until the intermediate **27** was completely consumed and then the mixture was concentrated under reduced pressure. The crude residue was purified by pre-HPLC to give intermediate **28** (159 mg, yield: 52%) as white solid. $^{1}$H NMR (400 MHz, DMSO-$d_6$) δ 7.77 (d, $J$ = 0.8 Hz, 1H), 7.51 (d, $J$ = 0.8 Hz, 1H), 7.14 (s, 1H), 6.87 (s, 1H), 6.81 (td, $J$ = 55.2 Hz, 6.8 Hz, 1H), 3.95 – 3.90 (m, 2H), 3.87 (s, 3H), 3.70 – 3.53 (m, 4H), 3.37 – 3.25 (m, 1H), 3.21 – 3.09 (m, 1H), 3.01 (dd, $J$ = 16.9, 4.4 Hz, 1H), 2.85 (t, $J$ = 6.4 Hz, 2H), 1.99 (p, $J$ = 6.5 Hz, 2H), 1.29 (dd, $J$ = 8.0, 6.6 Hz, 6H).

***methyl (1r,4r)-4-(3-(7-(difluoromethyl)-6-(1-methyl-1H-pyrazol-4-yl)-3,4-dihydroquinolin-1(2H)-yl)-5-isopropyl-4,5,6,7-tetrahydro-1H-pyrazolo[4,3-c]pyridin-1-yl)cyclohexane-1-carboxylate (30)***

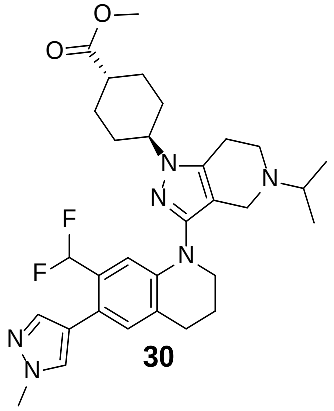

**30**

Step h: A solution of Intermediate **28** (150 mg, 0.35 mmol), $Cs_2CO_3$ (287.8 mg, 0.88 mmol) and methyl (1s,4s)-4-((methylsulfonyl)oxy)cyclohexane-1-carboxylate **29** (181.7 mg, 0.77 mmol) in DMF (5 mL). The mixture was stirred at 80 °C for overnight. Monitor the reaction through LC/MS until the intermediate **28** was completely consumed. Then the mixture was concentrated under reduced pressure. The crude residue was purified by pre-HPLC to give intermediate **30** (69 mg, yield: 35%) as yellow solid. (Two isomers are generated by substitution at the two nitrogen atoms of the pyrazole. In C18 reversed-phase chromatography, the desired compound has a slightly longer retention time than the other isomer). $^{1}$H NMR (400 MHz, DMSO-$d_6$) δ 7.75 (d, *J* = 0.8 Hz, 1H), 7.49 (d, *J* = 0.8 Hz, 1H), 7.07 (s, 1H), 6.78 (s, 1H), 6.78 (td, *J* = 55.2 Hz, 6.8 Hz, 1H), 4.05 – 3.95 (m, 1H), 3.87 (s, 3H), 3.61 (s, 3H), 3.59 – 3.53 (m, 2H), 3.16 (s, 2H), 2.88 – 2.78 (m, 3H), 2.72 (s, 4H), 2.38 (tt, *J* = 12.4, 8.8, 3.6 Hz, 1H), 2.06 – 1.76 (m, 8H), 1.53 (qd, *J* = 13.0, 3.8 Hz, 2H), 0.97 (d, *J* = 6.5 Hz, 6H).

***(1r,4r)-4-(3-(7-(difluoromethyl)-6-(1-methyl-1H-pyrazol-4-yl)-3,4-dihydroquinolin-1(2H)-yl)-5-isopropyl-4,5,6,7-tetrahydro-1H-pyrazolo[4,3-c]pyridin-1-yl)cyclohexane-1-carbaldehyde (31)***

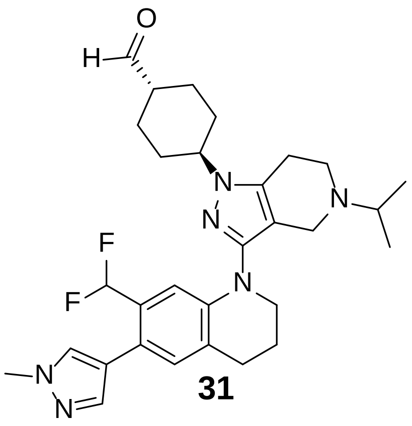

**31**

Step i: Intermediate **30** (60 mg, 0.10 mmol) was dissolved in anhydrous DCM (15 mL) and the solution was degassed and charged with $N_2$ 3 time. The mixture was stirred at -78 $^{o}$C for 5 min followed by diisobutylaluminum hydride (25% in toluene, 0.24 mL, 0.42 mmol) was added dropwise into the mixture. The reaction mixture was stirred at -78 $^{o}$C for additional 1.5 h. The reaction was then quenched with aqueous potassium sodium tartrate, after which the mixture was warmed to room temperature and stirred for an additional 30 minutes. The resulting mixture was diluted with DCM and filtered through Celite, and the filter cake was washed with DCM. The combined organic layers were concentrated under reduced pressure. The crude residue was used directly in the next step without further purification. UPLC-MS (ESI) *m*/*z* $[M+H]^+$ 537.30.

***N*-((1r,4r)-4-((3-chloro-4-cyanophenyl)(methyl)amino)cyclohexyl)-6-(4-(((1r,4r)-4-(3-(7-(difluoromethyl)-6-(1-methyl-1H-pyrazol-4-yl)-3,4-dihydroquinolin-1(2H)-yl)-5-isopropyl-4,5,6,7-tetrahydro-1H-pyrazolo[4,3-c]pyridin-1-yl)cyclohexyl)methyl)piperazin-1-yl)pyridazine-3-carboxamide (Neg-1)***

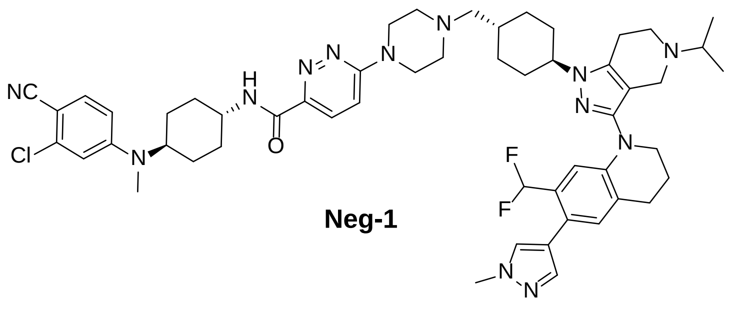

**Neg-1**

Step j: Compound **Neg-1** was synthesized from intermediate **31** and intermediate **18** following a similar procedure to that of **DALTAC-1** (yield 30%, two steps). $^{1}$H NMR (400 MHz, DMSO-$d_6$) δ 8.50 (d, $J$ = 8.4 Hz, 1H), 7.85 (d, $J$ = 9.3 Hz, 1H), 7.76 (s, 1H), 7.61 (d, $J$ = 8.9 Hz, 1H), 7.51 (s, 1H), 7.36 (d, $J$ = 9.7 Hz, 1H), 7.12 (s, 1H), 6.99 – 6.58 (m, 4H), 4.20 – 3.47 (m, 14H), 3.27 – 3.10 (m, 1H), 3.10 – 2.97 (m, 1H), 2.86 (s, 5H), 2.76 – 2.65 (m, 1H), 2.60 – 2.53 (m, 1H), 2.19 (s, 2H), 2.05 – 1.53 (m, 18H), 1.52 – 0.68 (m, 12H). Purity >95.0%. UPLC-MS (ESI) $m/z$ [M+H]$^+$ 974.52.

### Scheme 4: Synthesis of Neg-2.

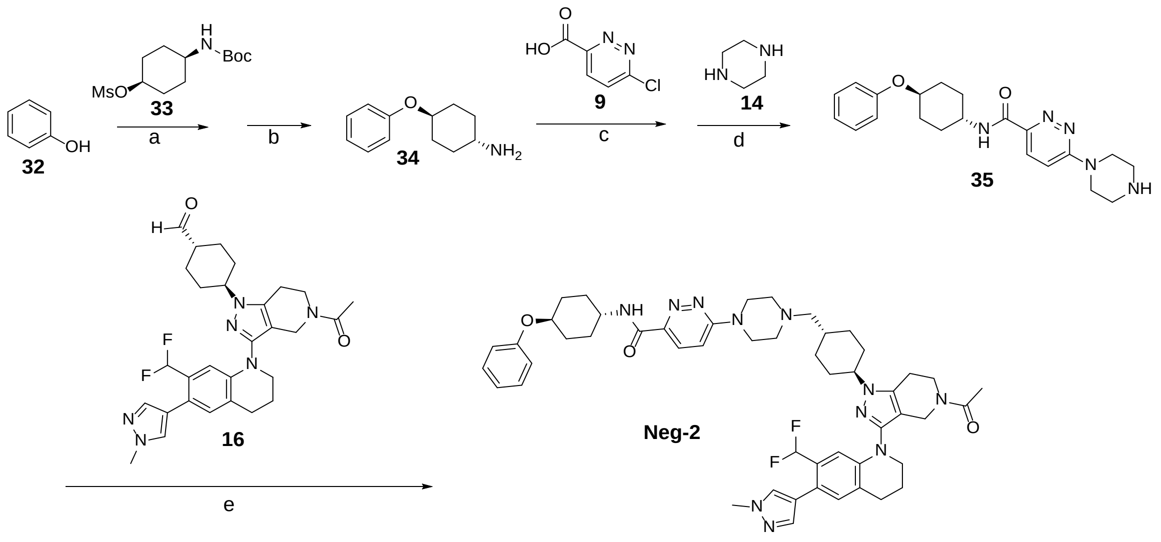

**_Reagents and conditions_**: (a) $K_2CO_3$, DMF, 70°C, 4 h; (b) DCM, TFA, reflex; (c) HATU, DMF, $Et_3N$, rt; (d) DIPEA, DMF, 55°C; (e) $Et_3N$, Sodium triacetoxyborohydride, DCM, rt.

***(1r,4r)-4-phenoxycyclohexan-1-amine (34)***

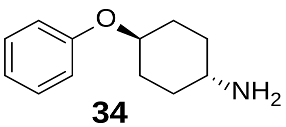

**34**

Step a: A solution of phenol **32** (420.0 mg, 4.4 mmol), $K_2CO_3$ (924.9 mg, 6.7 mmol) and compound **33** (943.0 mg, 4.4 mmol) in DMF (12 mL). The mixture was stirred at 70 °C for 4 h. After complete conversion, the mixture was diluted with $H_2O$, extracted with EtOAc, dried, and concentrated.

Step b: The above crude product was dissolved in 10 mL of DCM : TFA (4:1) and stirred at rt for an additional 4 h. Monitor the reaction through LC/MS until the reaction was completed. Then the mixture was concentrated, and purified by flash column chromatography (0–20% MeOH/DCM) to give intermediate **34** (337 mg, yield: 40%) as white solid. $^1$H NMR (400 MHz, DMSO-$d_6$) δ 7.31 – 7.23 (m, 2H), 6.99 – 6.88 (m, 3H), 4.34 – 4.12 (m, 1H), 3.16 – 3.00 (m, 1H), 2.15 – 2.04 (m, 2H), 2.04 – 1.94 (m, 2H), 1.56 – 1.35 (m, 4H).

***N-((1r,4r)-4-phenoxycyclohexyl)-6-(piperazin-1-yl)pyridazine-3-carboxamide (35)***

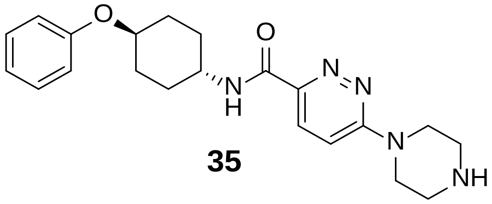

**35**

Step c and d: Compound **35** was synthesized from intermediate **34** following a similar procedure to that of intermediate **18** (yield 30%, two steps). $^1$H NMR (400 MHz, DMSO-$d_6$) δ 8.61 (d, $J$ = 8.3 Hz, 1H), 7.91 (d, $J$ = 9.5 Hz, 1H), 7.42 (d, $J$ = 9.6 Hz, 1H), 7.33 – 7.22 (m, 2H), 6.97 – 6.93 (m, 2H), 6.93 – 6.89 (m, 1H), 4.29 (tt, $J$ = 10.2, 4.1 Hz, 1H), 3.94 – 3.80 (m, 5H), 3.14 (t, $J$ = 5.1 Hz, 4H), 2.09 (d, $J$ = 5.9 Hz, 2H), 1.91 (d, $J$ = 13.0 Hz, 2H), 1.62 (q, $J$ = 12.9 Hz, 2H), 1.49 (q, $J$ = 10.9, 9.3 Hz, 2H).

***6-(4-(((1r,4r)-4-(5-acetyl-3-(7-(difluoromethyl)-6-(1-methyl-1H-pyrazol-4-yl)-3,4-dihydroquinolin-1(2H)-yl)-4,5,6,7-tetrahydro-1H-pyrazolo[4,3-c]pyridin-1-yl)cyclohexyl)methyl)piperazin-1-yl)-N-((1r,4r)-4-phenoxycyclohexyl)pyridazine-3-carboxamide (Neg-2)***

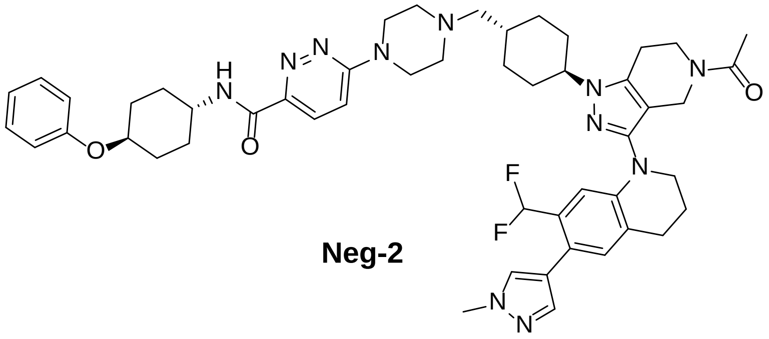

**Neg-2**

Step e: Compound **Neg-2** was synthesized from intermediate **35** and intermediate **16** following a similar procedure to that of compound **DALTAC-1** (yield 65%). $^{1}$H NMR (400 MHz, DMSO-$d_6$) δ 8.56 (dd, *J* = 8.3, 2.3 Hz, 1H), 7.83 (dd, *J* = 9.5, 1.9 Hz, 1H), 7.76 (s, 1H), 7.50 (s, 1H), 7.35 (dd, *J* = 9.7, 3.6 Hz, 1H), 7.31 – 7.24 (m, 2H), 7.11 (d, *J* = 3.0 Hz, 1H), 6.98 – 6.88 (m, 4H), 6.82 – 6.77 (m, 1H), 4.29 (tt, *J* = 10.4, 4.2 Hz, 1H), 4.23 – 4.07 (m, 2H), 4.07 – 3.95 (m, 1H), 3.87 (s, 4H), 3.79 – 3.63 (m, 5H), 3.63 – 3.55 (m, 2H), 2.89 – 2.81 (m, 3H), 2.49 – 2.44 (m, 2H), 2.34 (d, *J* = 7.6 Hz, 1H), 2.18 (dd, *J* = 7.4, 3.8 Hz, 1H), 2.15 – 2.05 (m, 4H), 2.01 – 1.86 (m, 8H), 1.86 – 1.69 (m, 3H), 1.62 (q, *J* = 12.7, 3H), 1.54 – 1.42 (m, 3H), 1.33 – 1.18 (m, 5H). Purity >95.0%. UPLC-MS (ESI) *m*/*z* [M+H]$^+$ 902.51.

### Scheme 5: Synthesis of Compounds p300i and ARi.

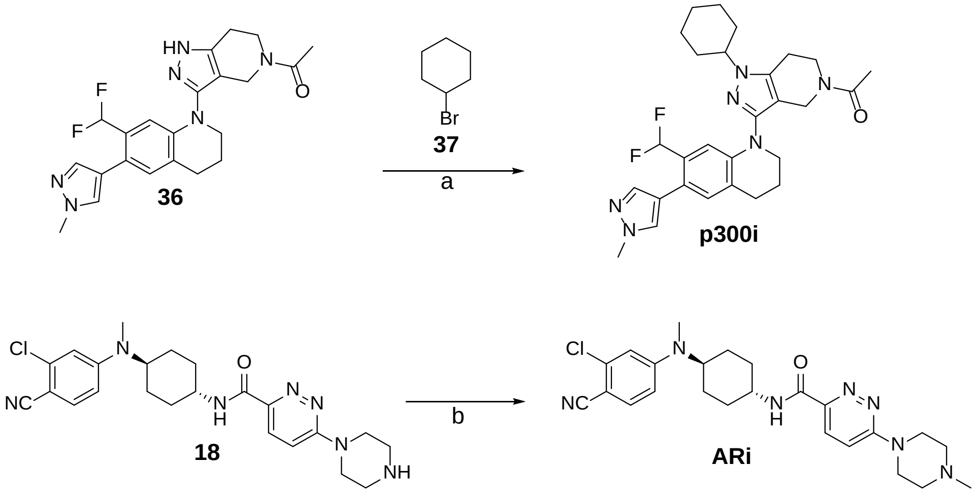

***Reagents and conditions*****:** (a) $Cs_2CO_3$, DMF, 80 °C, 8h; (b) Formaldehyde solution (37 wt. % in $H_2O$), MeOH, Sodium triacetoxyborohydride, rt, 30 min.

Intermediate **36** was prepared according to the reported procedures.[1]

***1-(1-cyclohexyl-3-(7-(difluoromethyl)-6-(1-methyl-1H-pyrazol-4-yl)-3,4-dihydroquinolin-1(2H)-yl)-1,4,6,7-tetrahydro-5H-pyrazolo[4,3-c]pyridin-5-yl)ethan-1-one (p300i)***

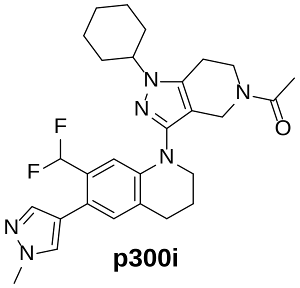

**p300i**

Step a: To a solution of intermediate **36** (43 mg, 0.1 mmol), $Cs_2CO_3$ (81.5 mg, 0.25 mmol) in DMF (5 mL) was added bromocyclohexane **37** (48.9 mg, 0.3 mmol). The mixture was stirred at 75 $^{o}$C for 7 h. After that the mixture was cooled, filtered through Celite, the filter cake was washed with DCM, and the filtrate was concentrated under reduced pressure. The residue was purified by pre-HPLC: $CH_3CN/H_2O$ from 35% to 100% in 60 min, flow rate (60 ml/min). The desired product started eluting at ~ 54% MeCN (Two isomers are generated by substitution at the two nitrogen atoms of the pyrazole. In C18 reversed-phase chromatography, the desired compound has a slightly longer retention time than the other isomer). $^1$H NMR (400 MHz, DMSO-$d_6$) δ 7.76 (s, 1H), 7.50 (s, 1H), 7.10 (s, 1H), 6.81 (d, *J* = 4.2 Hz, 1H), 6.78 (td, *J* = 55.2 Hz, 6.8 Hz, 1H), 4.14 (d, *J* = 20 Hz, 2H) 4.05 – 3.94 (m, 1H), 3.87 (s, 3H), 3.77 – 3.65 (m, 2H), 3.62 – 3.54 (m, 2H), 2.84 (q, *J* = 5.3 Hz, 3.26H), 2.72 (t, *J* = 5.6 Hz, 0.74H), 2.08 (s, 2H), 2.01 – 1.93 (m, 3H), 1.92 – 1.61 (m, 7H), 1.39 (q, *J* = 12.6, 12.2 Hz, 2H), 1.25 – 1.12 (m, 1H). Purity >95.0%. UPLC-MS (ESI) *m*/*z* $[M+H]^+$ 509.20.

***N-((1r,4r)-4-((3-chloro-4-cyanophenyl)(methyl)amino)cyclohexyl)-6-(4-methylpiperazin-1-yl)pyridazine-3-carboxamide (ARi)***

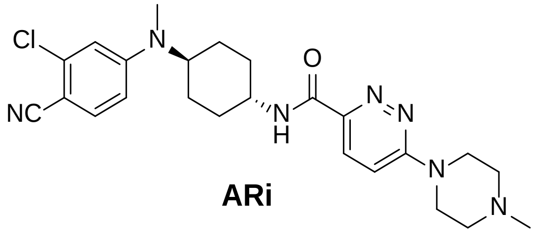

**ARi**

Step b: To a solution of intermediate **18** (45 mg, 0.1 mmol) in MeOH (2 mL), was added formaldehyde solution (37 wt. % in $H_2O$) (0.2 mL). The mixture was stirred at rt for 1 h. Then sodium triacetoxyborohydride (43 mg, 0.2 mmol) was added to this reaction. After complete conversion, the reaction mixture was concentrated under reduced pressure, and purified by flash column chromatography (0–15% MeOH/DCM) to give compound **ARi** (38 mg, yield: 82%). $^1$H NMR (400 MHz, DMSO-$d_6$) δ 8.58 (d, *J* = 8.3 Hz, 1H), 7.96 (d, *J* = 9.5 Hz, 1H), 7.61 (d, *J* = 9.0 Hz, 1H), 7.49 (d, *J* = 9.6 Hz, 1H), 6.95 (d, *J* = 2.6 Hz, 1H), 6.83 (dd, *J* = 9.1, 2.6 Hz, 1H), 4.62 (d, *J* = 13.8 Hz, 3H), 3.93 – 3.83 (m, 1H), 3.83 – 3.73 (m, 1H), 3.56 (d, *J* = 12.0 Hz, 2H), 3.35 (t, *J* = 12.5 Hz, 2H), 3.14 (s, 2H), 2.86 (s, 6H), 1.94 (d, *J* = 9.5 Hz, 2H), 1.83 – 1.60 (m, 6H). Purity >95.0%. UPLC-MS (ESI) *m*/*z* $[M+H]^+$ 468.20.

### Scheme 6: Synthesis of Compounds JZY3287, JZY3286, and JZY3288.

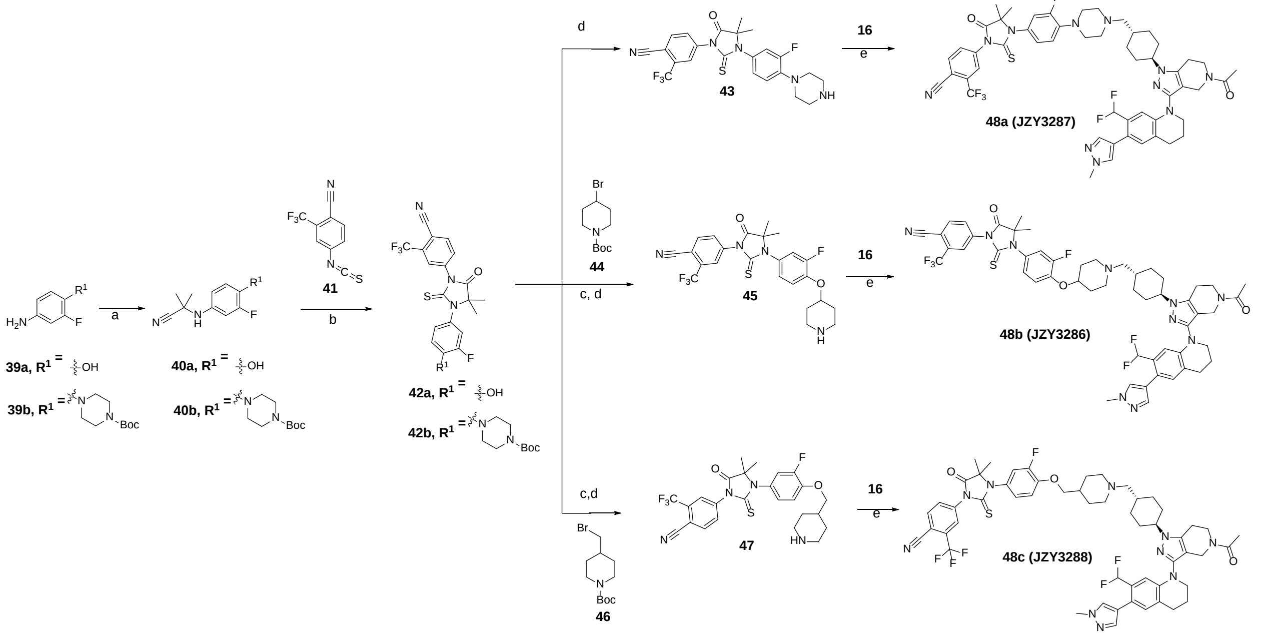

***Reagents and conditions***: (a) trimethylsilylformonitrile, trimethylsilyl trifluoromethanesulfonate, acetone: DCM (1:2), 20 °C,16 h; (b) i) *N,N*-dimethylacetamide, 20 °C, 2 h, ii) 2M HCl (aq.): MeOH (1:1), 0 °C to 70 °C for 2 h.1 (c) $Cs_2CO_3$, DMF, 65 °C, 5 h, (d) TFA, DCM, rt, 4 h, (e) $Et_3N$, Sodium triacetoxyborohydride, DCM, rt, 3 h.

***4-(3-(3-fluoro-4-hydroxyphenyl)-4,4-dimethyl-5-oxo-2-thioxoimidazolidin-1-yl)-2-(trifluoromethyl)benzonitrile (42a)***

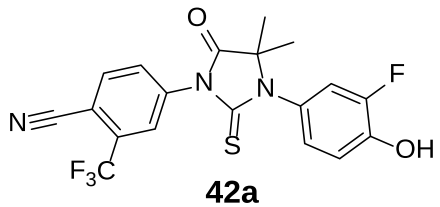

**42a**

Intermediate **42a** was prepared according to the reported procedures.[3]

$^1$H NMR (400 MHz, DMSO-$d_6$) δ 10.37 (s, 1H), 8.39 (d, *J* = 8.3 Hz, 1H), 8.28 (d, *J* = 1.9 Hz, 1H), 8.07 (dd, *J* = 8.2, 2.0 Hz, 1H), 7.23 (dd, *J* = 11.7, 2.4 Hz, 1H), 7.13 – 7.06 (m, 1H), 7.02 (ddd, *J* = 8.6, 2.4, 0.9 Hz, 1H), 1.50 (s, 6H).

***tert-butyl 4-(4-(3-(4-cyano-3-(trifluoromethyl)phenyl)-5,5-dimethyl-4-oxo-2-thioxoimidazolidin-1-yl)-2-fluorophenyl)piperazine-1-carboxylate (42b)***

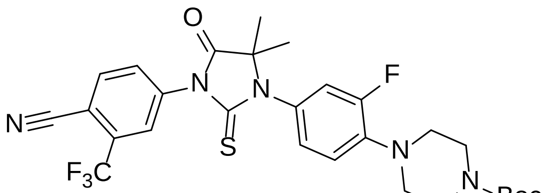

**42b**

Compound **42b** was synthesized following a similar procedure to that of **42a**. UPLC-MS (ESI) *m*/*z* $[M+H]^+$ 592.21.

***4-(3-(3-fluoro-4-(piperazin-1-yl)phenyl)-4,4-dimethyl-5-oxo-2-thioxoimidazolidin-1-yl)-2-(trifluoromethyl)benzonitrile (43)***

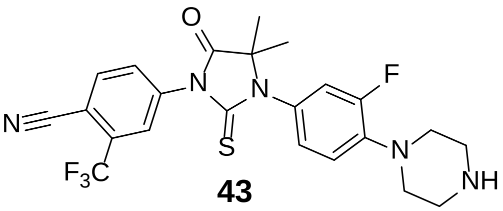

**43**

The intermediate **42b (**300 mg, 0.51 mmol) was dissolved in DCM (20 mL) and TFA (4 mL), the mixture was stirred at 40 °C for an additional 3 h. Monitor the reaction through LC/MS until the reaction was completed. Then the mixture was concentrated, and purified by pre-HPLC (15−100% MeCN/$H_2O$ in 85 min, the desired product started eluting at ~ 30% MeCN), to give intermediate **43** (200 mg, yield: 82%) as a clear solid.$^1$H NMR (400 MHz, DMSO-$d_6$) δ 8.40 (d, *J* = 8.2 Hz, 1H), 8.28 (d, *J* = 1.9 Hz, 1H), 8.07 (dd, *J* = 8.2, 2.0 Hz, 1H), 7.33 – 7.23 (m, 2H), 7.18 (dd, *J* = 8.6, 2.3 Hz, 1H), 3.32 (m, 8H), 1.52 (s, 6H).

***4-(3-(3-fluoro-4-(piperidin-4-yloxy)phenyl)-4,4-dimethyl-5-oxo-2-thioxoimidazolidin-1-yl)-2-(trifluoromethyl)benzonitrile (45)***

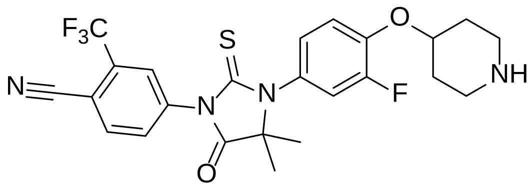

**45**

A mixture of intermediate **42a** (420 mg, 1.0 mmol), $Cs_2CO_3$ (809 mg, 2.5 mmol), and compound **44** (789 mg, 3.0 mmol) in DMF (25 mL) was stirred at 65 °C for 5 h. The reaction progress was monitored by LC/MS until complete consumption of intermediate **42a**. The reaction mixture was then quenched with ice water and extracted with EtOAc. The combined organic extracts were concentrated under reduced pressure to afford a crude residue, which was subsequently dissolved in DCM/TFA (3:1). The resulting mixture was stirred at room temperature for 4 h, monitored by LC/MS until completion. The solvent was then removed under reduced pressure, and the residue was purified by pre-HPLC to afford intermediate **45** (283 mg, 56% yield, two steps).$^1$H NMR (400 MHz, DMSO-$d_6$) δ 8.40 (d, $J$ = 8.2 Hz, 1H), 8.28 (d, $J$ = 1.9 Hz, 1H), 8.07 (dd, $J$ = 8.2, 2.0 Hz, 1H), 7.46 (t, $J$ = 9.1 Hz, 1H), 7.36 (dd, $J$ = 11.6, 2.5 Hz, 1H), 7.19 (ddd, $J$ = 8.7, 2.5, 1.4 Hz, 1H), 4.81 – 4.73 (m, 1H), 3.28 (s, 2H), 3.13 (s, 2H), 2.21 – 2.10 (m, 2H), 1.95 – 1.83 (m, 2H), 1.52 (s, 6H).

***4-(3-(3-fluoro-4-(piperidin-4-ylmethoxy)phenyl)-4,4-dimethyl-5-oxo-2-thioxoimidazolidin-1-yl)-2-(trifluoromethyl)benzonitrile (47)***

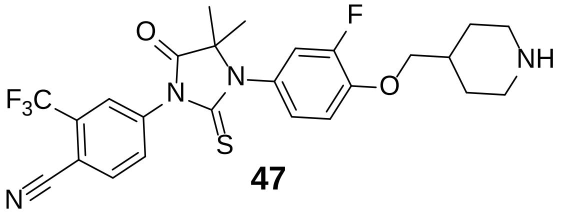

**47**

Compound **47** was synthesized from intermediate **42a** and intermediate **46** following a similar procedure to that of **45**. $^1$H NMR (400 MHz, DMSO-$d_6$) δ 8.40 (d, $J$ = 8.3 Hz, 1H), 8.28 (d, $J$ = 1.9 Hz, 1H), 8.07 (dd, $J$ = 8.2, 2.0 Hz, 1H), 7.39 – 7.31 (m, 2H), 7.18 (ddd, $J$ = 8.7, 2.5, 1.3 Hz, 1H), 4.04 (d, $J$ = 6.1 Hz, 2H), 3.34 (d, $J$ = 12.5 Hz, 2H), 2.95 (q, $J$ = 12.0 Hz, 2H), 2.20 – 2.08 (m, 1H), 1.95 (d, $J$ = 12.7 Hz, 2H), 1.59 – 1.44 (m, 8H).

***4-(3-(4-(4-(((1r,4r)-4-(5-acetyl-3-(7-(difluoromethyl)-6-(1-methyl-1H-pyrazol-4-yl)-3,4-dihydroquinolin-1(2H)-yl)-4,5,6,7-tetrahydro-1H-pyrazolo[4,3-c]pyridin-1-yl)cyclohexyl)methyl)piperazin-1-yl)-3-fluorophenyl)-4,4-dimethyl-5-oxo-2-thioxoimidazolidin-1-yl)-2-(trifluoromethyl)benzonitrile (JZY3287)***

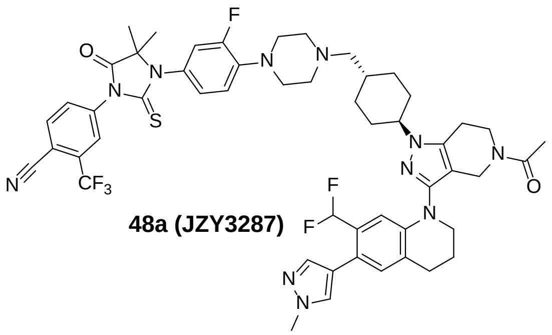

**48a (JZY3287)**

Compound **JZY3287** was synthesized from intermediates **43** and **16** following a similar procedure to that of **DALTAC-1**. $^{1}$H NMR (400 MHz, DMSO-$d_6$) δ 8.39 (d, *J* = 8.3 Hz, 1H), 8.28 (d, *J* = 1.9 Hz, 1H), 8.09 – 8.05 (m, 1H), 7.75 (s, 1H), 7.50 (s, 1H), 7.17 (dd, *J* = 38.6, 14.1 Hz, 4H), 6.80 (d, *J* = 2.8 Hz, 1H), 6.78 (td, J = 54.8, 7.1 Hz, 1H), 4.15 (d, *J* = 17.3 Hz, 2H), 4.06 – 3.96 (m, 1H), 3.87 (s, 3H), 3.77 – 3.66 (m, 2H), 3.59 (q, *J* = 5.5 Hz, 2H), 3.12 (s, 4H), 2.85 (d, *J* = 5.8 Hz, 3H), 2.54 (s, 4H), 2.20 (d, *J* = 6.9 Hz, 1H), 2.08 (d, *J* = 1.6 Hz, 2H), 2.01 - 1.87 (m, 7H), 1.83 (d, *J* = 11.8 Hz, 1H), 1.66 – 1.55 (m, 1H), 1.51 (s, 6H), 1.25 (s, 3H), 1.08 (q, *J* = 12.6 Hz, 2H). Purity >95.0%. UPLC-MS (ESI) *m/z* $[M+H]^+$ 1012.41.

***4-(3-(4-((1-(((1r,4r)-4-(5-acetyl-3-(7-(difluoromethyl)-6-(1-methyl-1H-pyrazol-4-yl)-3,4-dihydroquinolin-1(2H)-yl)-4,5,6,7-tetrahydro-1H-pyrazolo[4,3-c]pyridin-1-yl)cyclohexyl)methyl)piperidin-4-yl)oxy)-3-fluorophenyl)-4,4-dimethyl-5-oxo-2-thioxoimidazolidin-1-yl)-2-(trifluoromethyl)benzonitrile (JZY3286)***

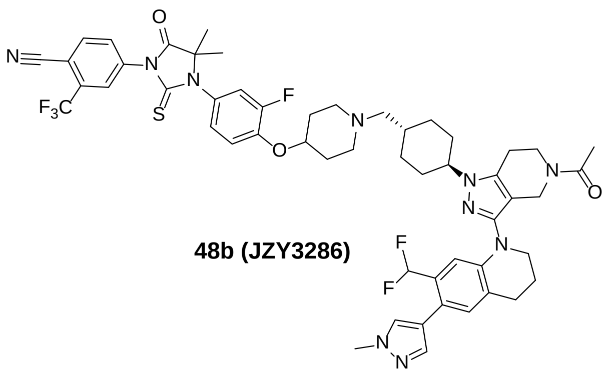

**48b (JZY3286)**

Compound **JZY3286** was synthesized from intermediates **45** and **16** following a similar procedure to that of **DALTAC-1**.$^{1}$H NMR (400 MHz, DMSO-$d_6$) δ 8.40 (d, *J* = 8.3 Hz, 1H), 8.28 (d, *J* = 1.9 Hz, 1H), 8.07 (dd, *J* = 8.2, 2.0 Hz, 1H), 7.75 (s, 1H), 7.50 (d, *J* = 0.9 Hz, 1H), 7.40 (t, *J* = 9.1 Hz, 1H), 7.31 (d, *J* = 11.6 Hz, 1H), 7.14 (d, *J* = 8.9 Hz, 1H), 7.10 (s, 1H), 6.79 (d, *J* = 2.6 Hz, 1H), 4.51 (s, 1H), 4.15 (d, *J* = 17.4 Hz, 2H), 4.00 (s, 1H), 3.87 (d, *J* = 1.3 Hz, 3H), 3.78 – 3.66 (m, 2H), 3.58 (t, *J* = 5.8 Hz, 2H), 2.85 (d, *J* = 5.8 Hz, 3H), 2.72 (d, *J* = 9.2 Hz, 3H), 2.23 (s, 2H), 2.15 (s, 2H), 2.08 (d, *J* = 1.6 Hz, 2H), 1.97 (d, *J* = 5.0 Hz, 5H), 1.82 (d, *J* = 12.8 Hz, 1H), 1.71 (s, 2H), 1.51 (s, 6H), 1.25 (s, 2H), 1.05 (d, *J* = 12.7 Hz, 2H). Purity >95.0%. Purity >95.0%. UPLC-MS (ESI) *m*/*z* $[M+H]^+$ 1027.42.

***4-(3-(4-((1-(((1r,4r)-4-(5-acetyl-3-(7-(difluoromethyl)-6-(1-methyl-1H-pyrazol-4-yl)-3,4-dihydroquinolin-1(2H)-yl)-4,5,6,7-tetrahydro-1H-pyrazolo[4,3-c]pyridin-1-yl)cyclohexyl)methyl)piperidin-4-yl)methoxy)-3-fluorophenyl)-4,4-dimethyl-5-oxo-2-thioxoimidazolidin-1-yl)-2-(trifluoromethyl)benzonitrile (JZY3288)***

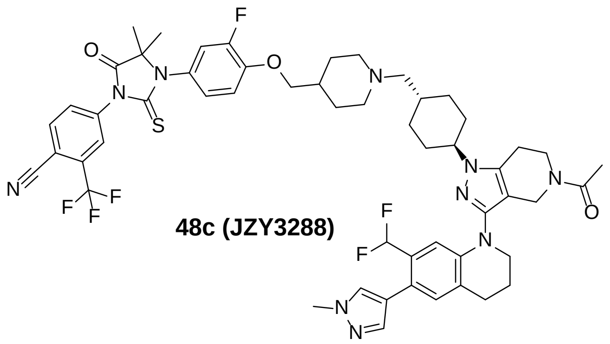

**48c (JZY3288)**

Compound **JZY3288** was synthesized from intermediates **47** and **16** following a similar procedure to that of **DALTAC-1**.$^{1}$H NMR (400 MHz, DMSO-$d_6$) δ 8.40 (d, *J* = 8.3 Hz, 1H), 8.28 (d, *J* = 1.9 Hz, 1H), 8.07 (dd, *J* = 8.3, 1.9 Hz, 1H), 7.75 (s, 1H), 7.50 (d, *J* = 0.9 Hz, 1H), 7.38 – 7.27 (m, 2H), 7.16 (d, *J* = 8.9 Hz, 1H), 7.10 (s, 1H), 6.80 - 6.78 (m, 1H), 6.78 (td, J = 54.8, 7.1 Hz, 1H), 4.14 (d, *J* = 16.7 Hz, 2H), 4.02 - 3.95 (m, 2H), 3.87 (s, 3H), 3.72 (d, *J* = 22.0 Hz, 2H), 3.58 (d, *J* = 5.5 Hz, 2H), 2.92 – 2.69 (m, 6H), 2.12 (s, 1H), 2.08 (s, 2H), 2.03 – 1.94 (m, 4H), 1.90 (s, 5H), 1.78 (s, 5H), 1.51 (s, 6H), 1.40 – 1.29 (m, 2H), 1.25 (s, 4H), 1.05 (s, 1H). Purity >95.0%. UPLC-MS (ESI) *m*/*z* $[M+H]^+$ 1041.45.

### Scheme 7: Synthesis of JZY4055.

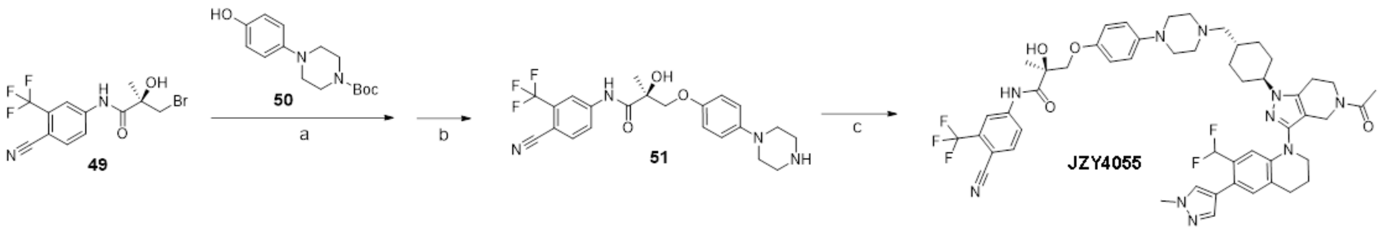

***Reagents and conditions*****:** (a) $Cs_2CO_3$, DMF, 65 °C, overnight, (b) TFA, DCM, rt, 4 h, (c) $Et_3N$, Sodium triacetoxyborohydride, DCM, rt, 3 h.

***(S)-N-(4-cyano-3-(trifluoromethyl)phenyl)-2-hydroxy-2-methyl-3-(4-(piperazin-1-yl)phenoxy)propenamide (51)***

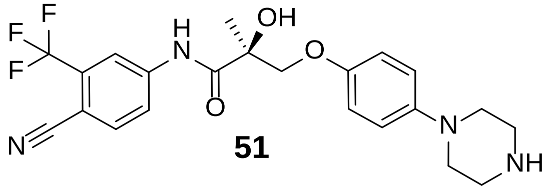

**51**

Compound **51** was synthesized from intermediates **49** and **50** following a similar procedure to that of intermediates **45**. $^1$H NMR (400 MHz, DMSO-$d_6$) δ 10.57 (s, 1H), 8.57 (d, *J* = 2.1 Hz, 1H), 8.30 (dd, *J* = 8.6, 2.1 Hz, 1H), 8.11 (d, *J* = 8.6 Hz, 1H), 6.94 – 6.87 (m, 2H), 6.87 – 6.82 (m, 2H), 4.16 (d, *J* = 9.6 Hz, 1H), 3.93 (d, *J* = 9.6 Hz, 1H), 3.27 – 3.16 (m, 8H), 1.43 (s, 3H).

***(S)-3-(4-(4-(((1r,4S)-4-(5-acetyl-3-(7-(difluoromethyl)-6-(1-methyl-1H-pyrazol-4-yl)-3,4-dihydroquinolin-1(2H)-yl)-4,5,6,7-tetrahydro-1H-pyrazolo[4,3-c]pyridin-1-yl)cyclohexyl)methyl)piperazin-1-yl)phenoxy)-N-(4-cyano-3-(trifluoromethyl)phenyl)-2-hydroxy-2-methylpropanamide (JZY4055)***

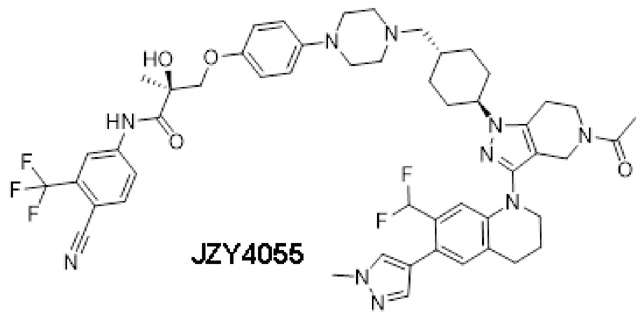

**JZY4055**

Compound **JZY4055** was synthesized from intermediates **51** and **16** following a similar procedure to that of **DALTAC-1**.$^{1}$H NMR (400 MHz, DMSO-$d_6$) δ 10.55 (s, 1H), 8.56 (d, *J* = 2.1 Hz, 1H), 8.31 (dd, *J* = 8.7, 2.1 Hz, 1H), 8.10 (d, *J* = 8.6 Hz, 1H), 7.75 (s, 1H), 7.50 (d, *J* = 0.9 Hz, 1H), 7.10 (s, 1H), 6.86 – 6.77 (m, 5H), 6.20 (s, 1H), 4.16 (d, *J* = 5.0 Hz, 1H), 4.13 - 4.10 (m, 1H), 4.00 (s, 1H), 3.91 (d, *J* = 9.6 Hz, 1H), 3.87 (s, 3H), 3.77 – 3.65 (m, 2H), 3.58 (q, *J* = 6.1 Hz, 2H), 2.99 (s, 4H), 2.85 (s, 3H), 2.47 (s, 4H), 2.16 (d, *J* = 6.8 Hz, 2H), 2.08 (s, 2H), 2.0 - 1.94 (m, 3H), 1.94 – 1.85 (m, 4H),1.82 (q, *J* = 12.6 Hz, 1H), 1.62 - 1.52 (m, 2H), 1.42 (s, 3H), 1.25 (s, 2H), 1.06 (d, *J* = 12.2 Hz, 2H). Purity >95.0%. UPLC-MS (ESI) *m*/*z* $[M+H]^+$ 969.43.

#### The $^{1}H$ NMR, $^{13}C$ NMR, and UPLC-MS traces of Compound DALTAC-1 (JZY3032)

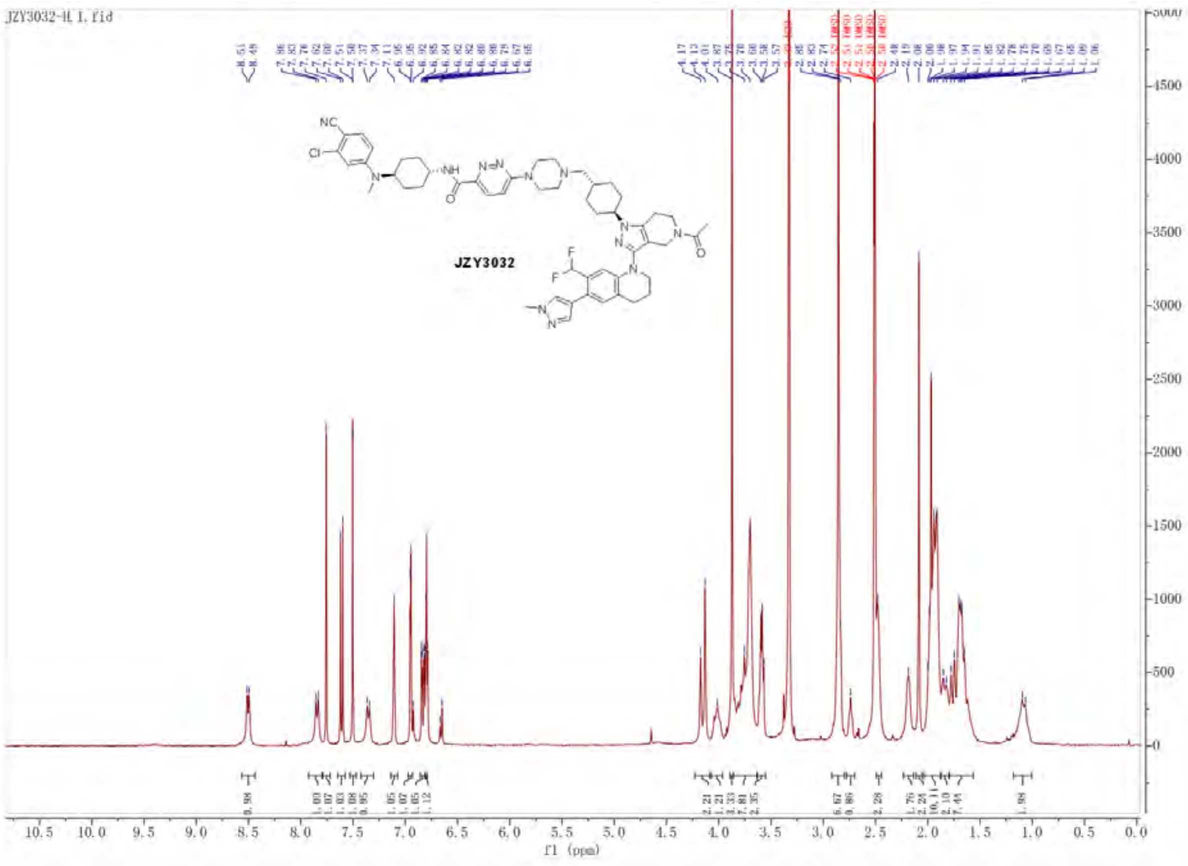

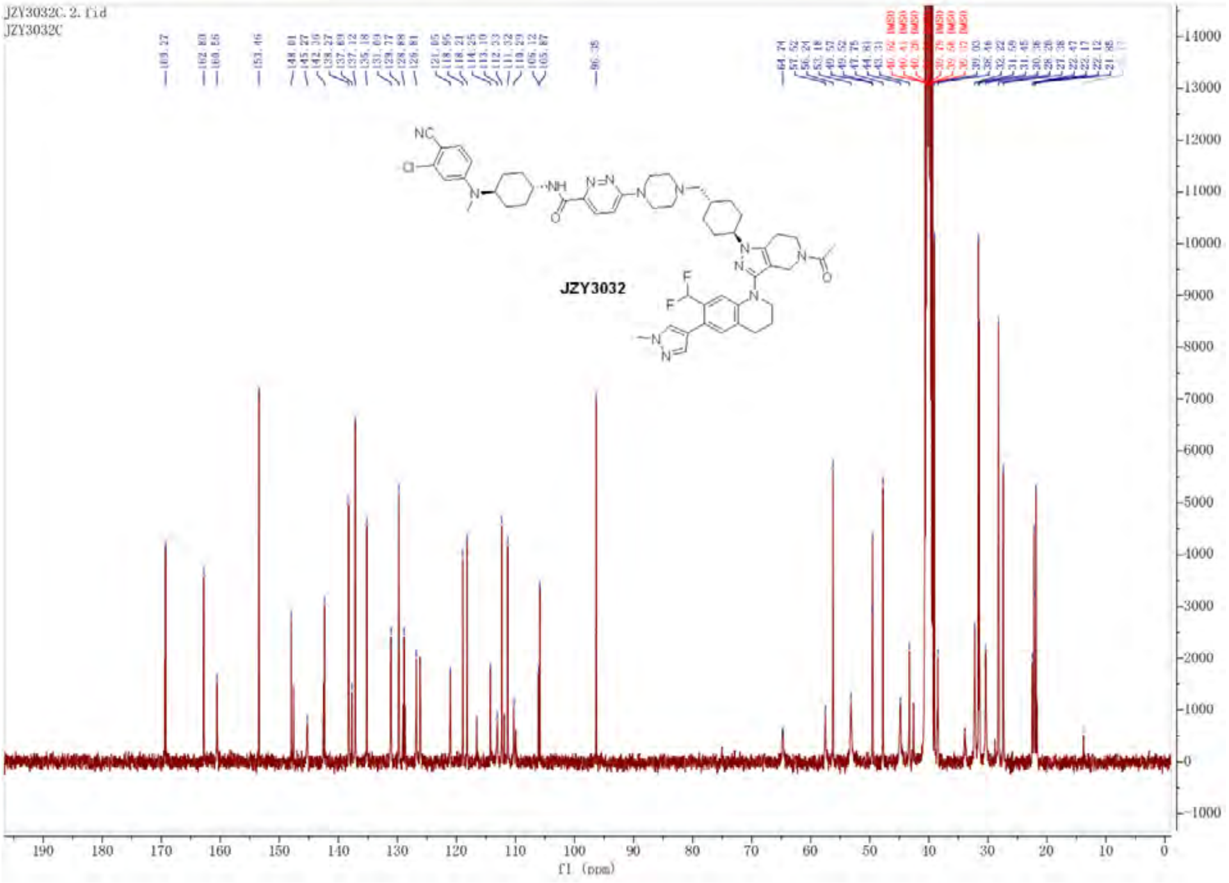
JZY3032C.2.fid
JZY3032C
JZY3032
f1 (ppm)

### Untitled

#### SAMPLE INFORMATION

| | | | |
|---|---|---|---|
| Sample Name: | jzy3032 | Acquired By: | System |
| Sample Type: | Unknown | Sample Set Name: | YQY |
| Vial: | 1:F,4 | Acq. Method Set: | 10to100Bin10min_5minDelay |
| Injection #: | 1 | Processing Method: | LC PQ ApexTrack |
| Injection Volume: | 8.00 ul | Channel Name: | 254.0nm |
| Run Time: | 10.0 Minutes | Proc. Chnl. Descr.: | PDA Spectrum PDA 254.0 nm |
| Date Acquired: | 10/8/2025 8:46:21 PM EDT | | |
| Date Processed: | 10/8/2025 9:17:01 PM EDT | | |

**Auto-Scaled Chromatogram**

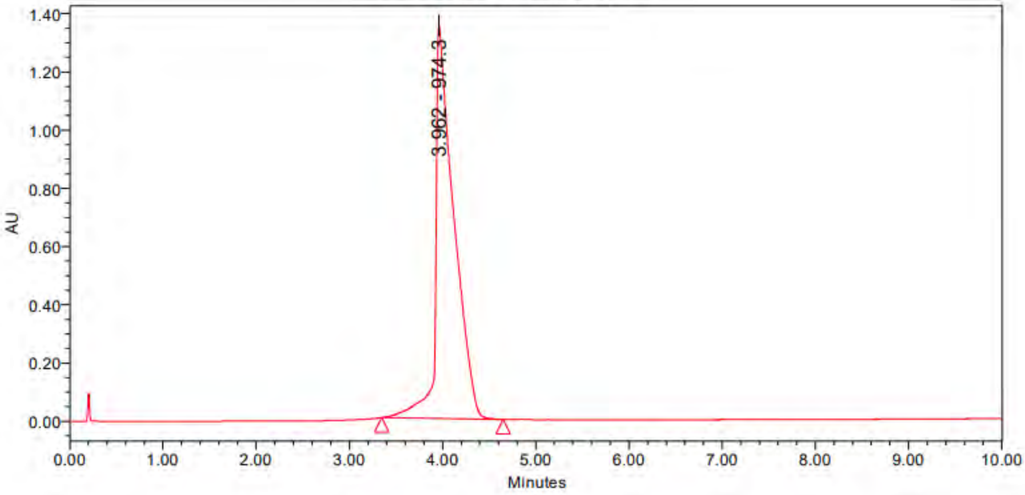

Peak Results

| | Name | RT | Area | Height | Amount | Units |
|---|---|---|---|---|---|---|
| 1 | | 3.962 | 18386448 | 1349888 | | |

PDA Result Table

| | Name | RT | Purity1 Angle | Purity1 Threshold | Match1 Spect. Name | Match1 Angle | Match1 Threshold |
|---|---|---|---|---|---|---|---|
| 1 | | 3.962 | | | | | |

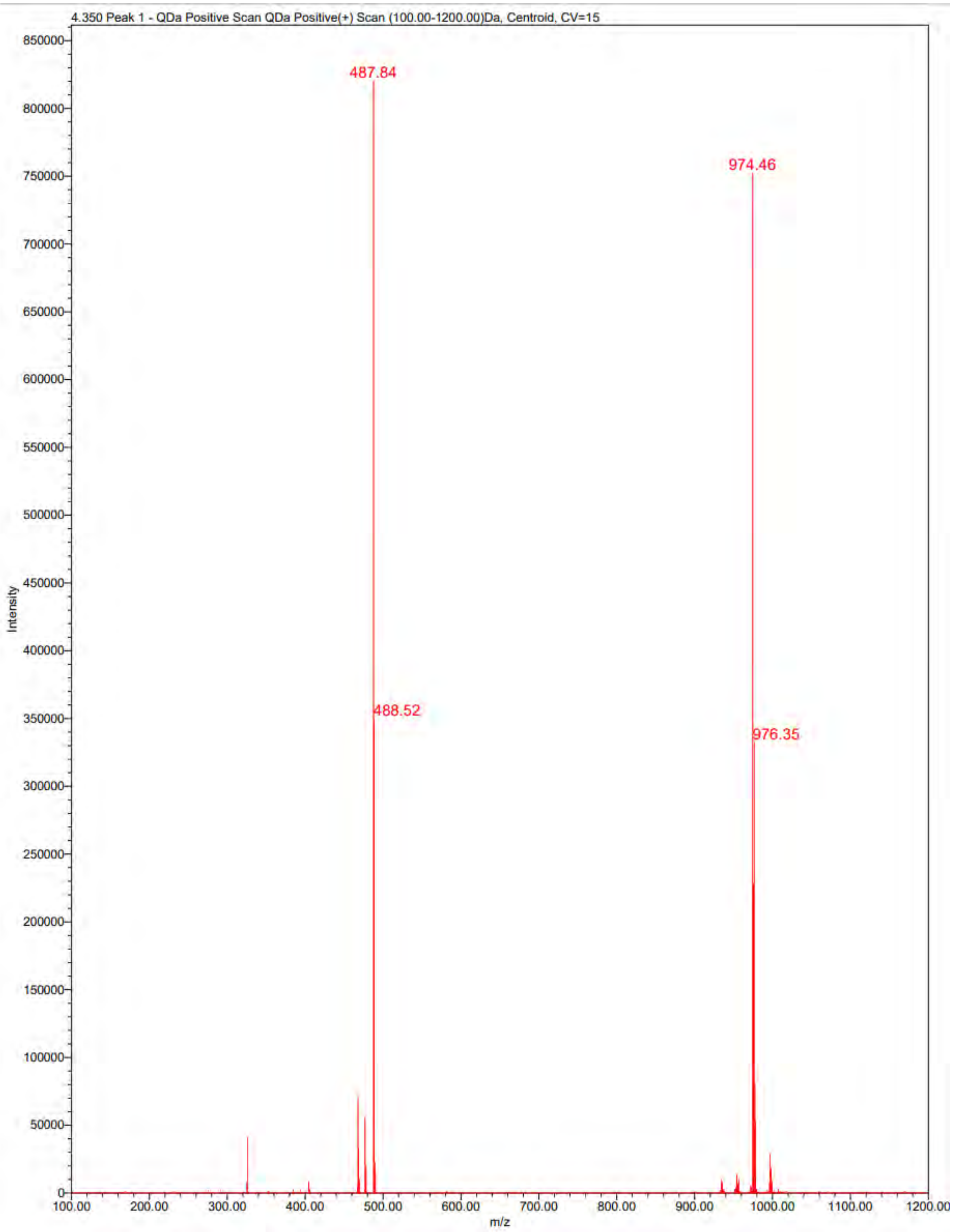

4.350 Peak 1 - QDa Positive Scan QDa Positive(+) Scan (100.00-1200.00)Da, Centroid, CV=15
487.84
974.46
488.52
976.35
Intensity
m/z

**The $^{1}$H NMR and UPLC-MS traces of Compound 1-7, and Neg-1, Neg-2, p300i, ARi, JZY3287, JZY3286, JZY3288, and JZY4055.**

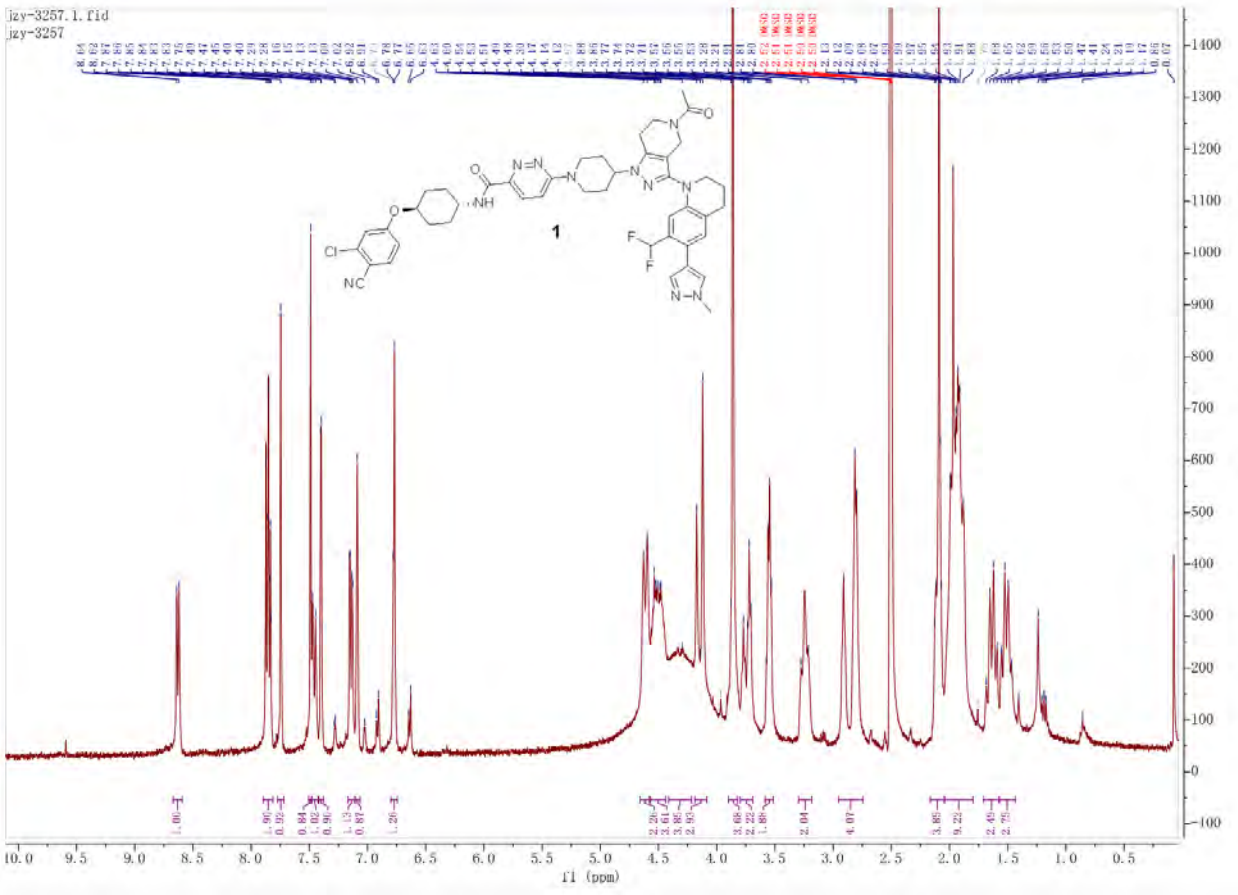

Empower 3

### Untitled

#### SAMPLE INFORMATION

| | | | |
|---|---|---|---|
| Sample Name: | JZY3257 | Acquired By: | System |
| Sample Type: | Unknown | Sample Set Name: | YQY |
| Vial: | 2:B,1 | Acq. Method Set: | 10to100Bin10min_5minDelay |
| Injection #: | 1 | Processing Method: | LC PQ ApexTrack |
| Injection Volume: | 3.00 ul | Channel Name: | 254.0nm |
| Run Time: | 10.0 Minutes | Proc. Chnl. Descr.: | PDA Spectrum PDA 254.0 nm |
| Date Acquired: | 7/26/2026 3:39:33 AM EDT | | |
| Date Processed: | 7/26/2026 3:53:57 PM EDT | | |

##### Auto-Scaled Chromatogram

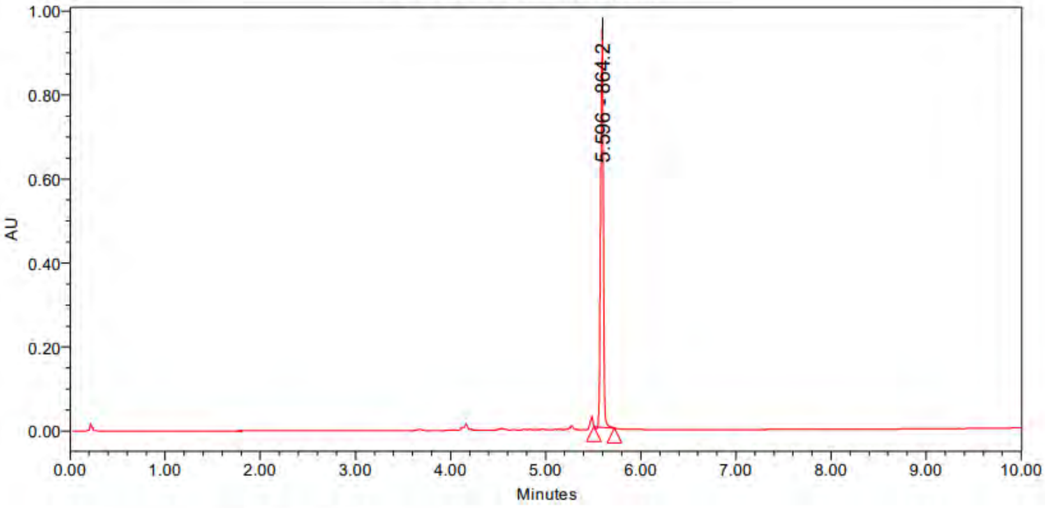

Peak Results

| | Name | RT | Area | Height | Amount | Units |
|---|---|---|---|---|---|---|
| 1 | | 5.596 | 1984387 | 952569 | | |

PDA Result Table

| | Name | RT | Purity1 Angle | Purity1 Threshold | Match1 Spect. Name | Match1 Angle | Match1 Threshold |
|---|---|---|---|---|---|---|---|
| 1 | | 5.596 | | | | | |

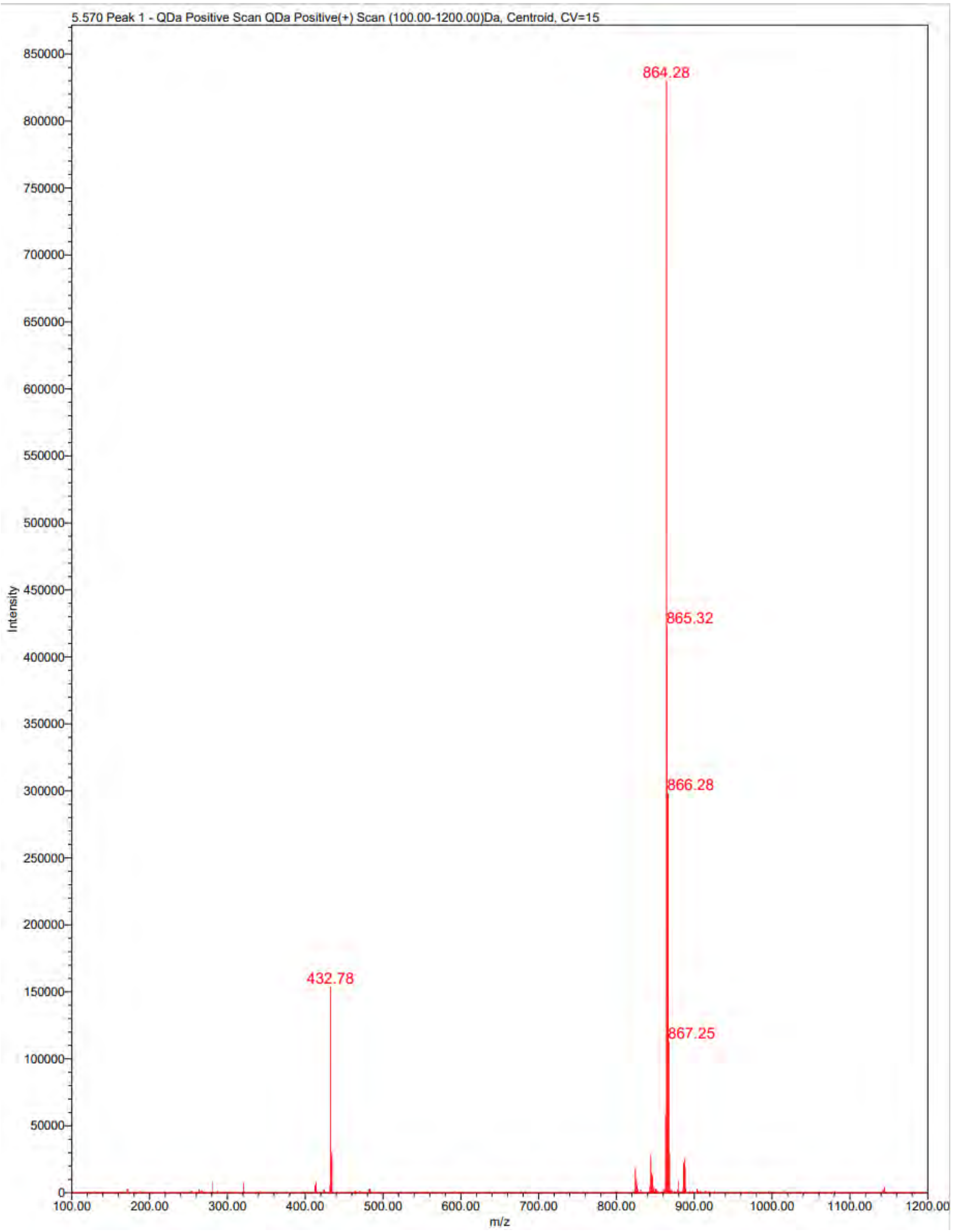
5.570 Peak 1 - QDa Positive Scan QDa Positive(+) Scan (100.00-1200.00)Da, Centroid, CV=15
Intensity
m/z
864.28
865.32
866.28
867.25
432.78

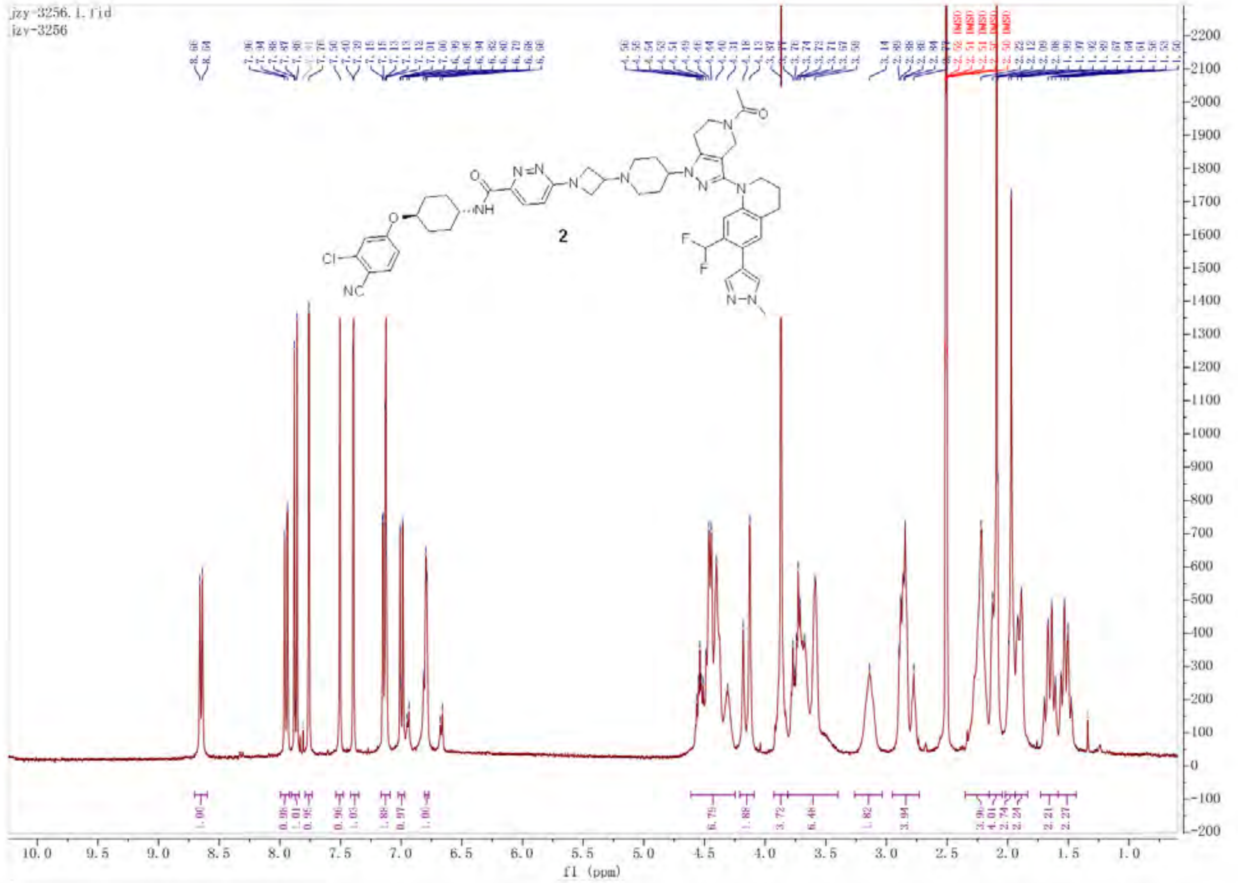

Empower 3

### Untitled

#### SAMPLE INFORMATION

| | | | |
|---|---|---|---|
| Sample Name: | JZY3256 | Acquired By: | System |
| Sample Type: | Unknown | Sample Set Name: | YQY |
| Vial: | 2:A,8 | Acq. Method Set: | 10to100Bin10min_5minDelay |
| Injection #: | 1 | Processing Method: | LC PQ ApexTrack |
| Injection Volume: | 3.00 ul | Channel Name: | 254.0nm |
| Run Time: | 10.0 Minutes | Proc. Chnl. Descr.: | PDA Spectrum PDA 254.0 nm |
| Date Acquired: | 7/26/2026 3:23:56 AM EDT | | |
| Date Processed: | 7/26/2026 3:55:11 PM EDT | | |

##### Auto-Scaled Chromatogram

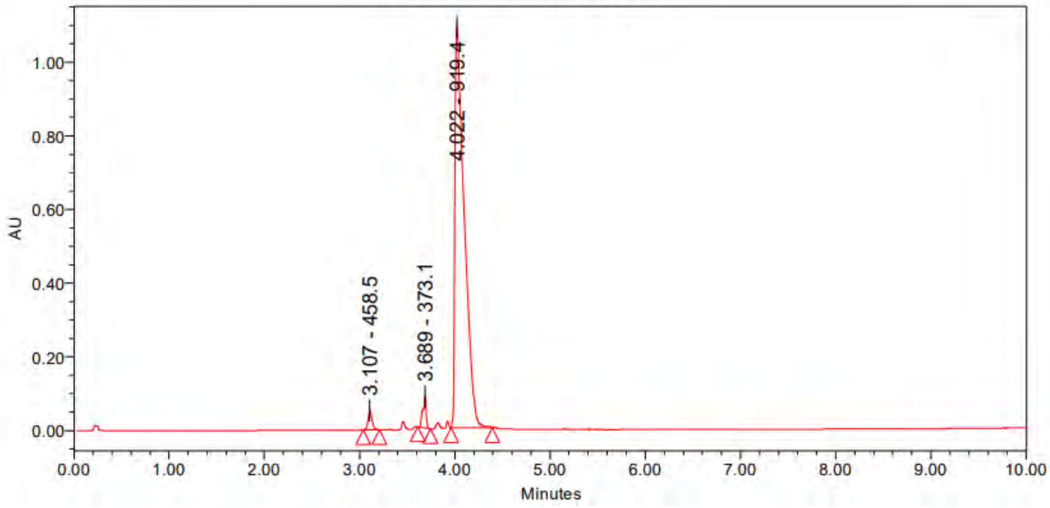

Peak Results

| | Name | RT | Area | Height | Amount | Units |
|---|---|---|---|---|---|---|
| 1 | | 3.107 | 146329 | 53226 | | |
| 2 | | 3.689 | 214774 | 85373 | | |
| 3 | | 4.022 | 7472782 | 1089154 | | |

PDA Result Table

| | Name | RT | Purity1 Angle | Purity1 Threshold | Match1 Spect. Name | Match1 Angle | Match1 Threshold |
|---|---|---|---|---|---|---|---|
| 1 | | 3.107 | | | | | |
| 2 | | 3.689 | | | | | |
| 3 | | 4.022 | | | | | |

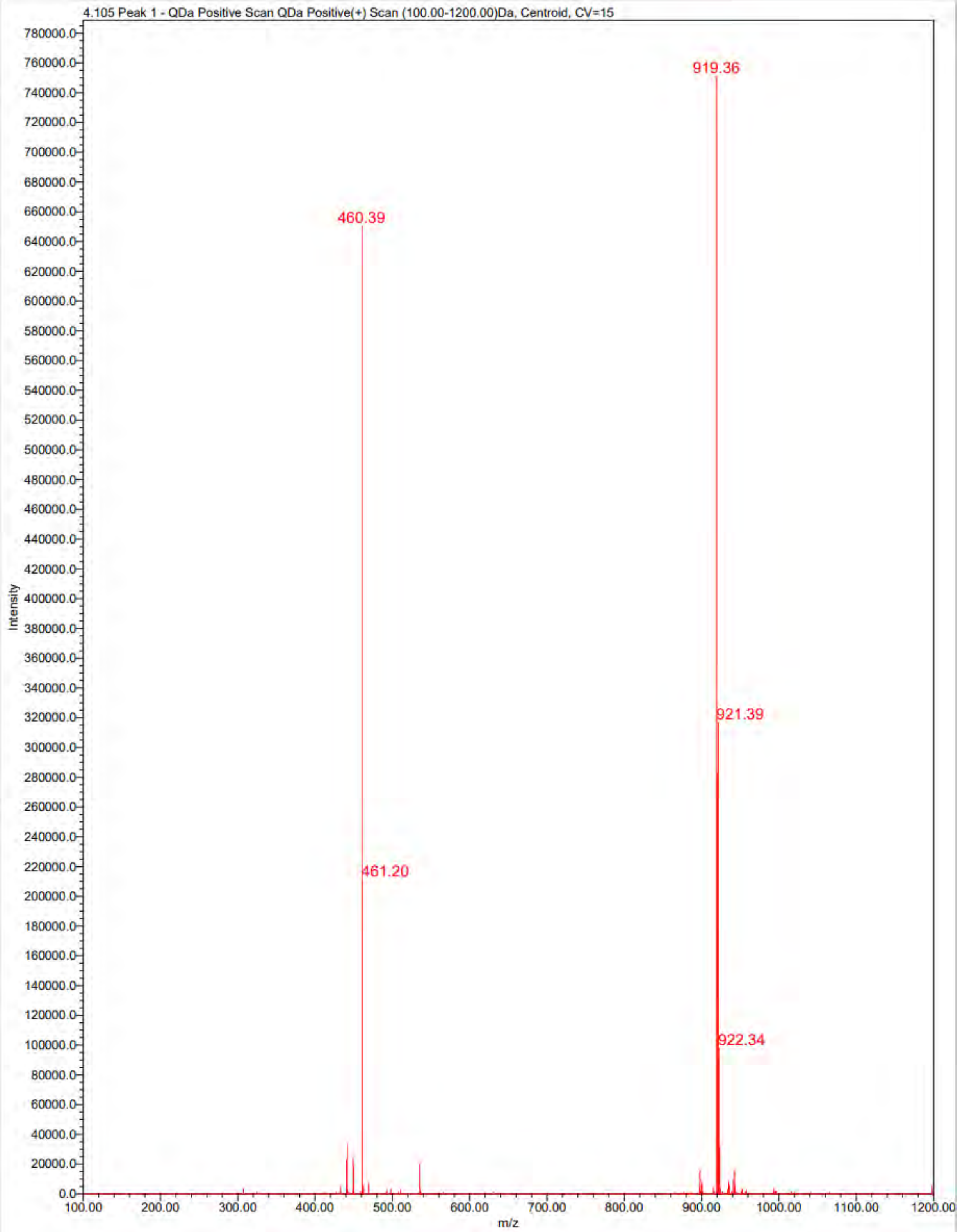

4.105 Peak 1 - QDa Positive Scan QDa Positive(+) Scan (100.00-1200.00)Da, Centroid, CV=15
Intensity
m/z
919.36
460.39
921.39
461.20
922.34

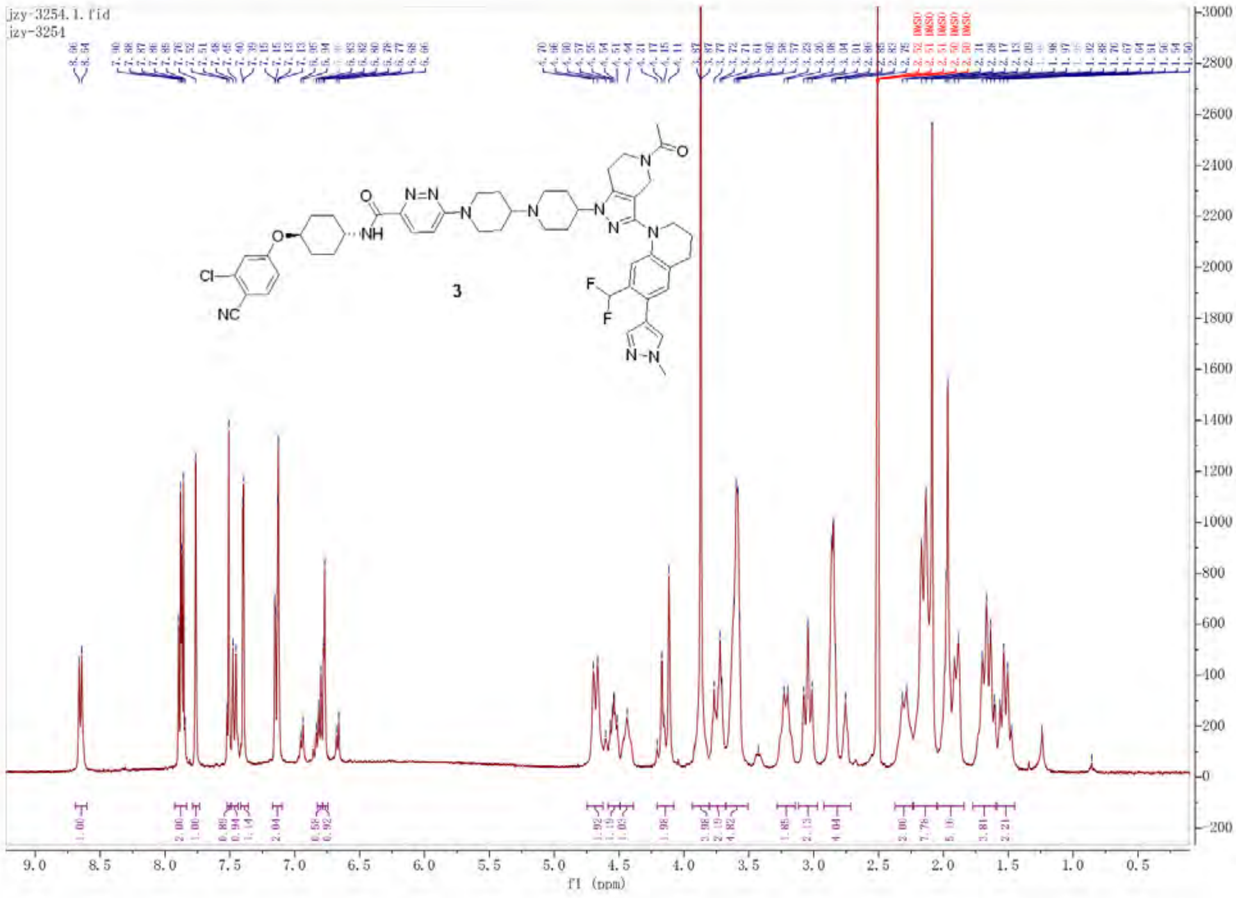

jzy-3254.1.fid
jzy-3254
3
f1 (ppm)

Untitled

#### SAMPLE INFORMATION

| | | | |
|---|---|---|---|
| Sample Name: | JZY3254 | Acquired By: | System |
| Sample Type: | Unknown | Sample Set Name: | 0 |
| Vial: | 1:D,4 | Acq. Method Set: | 10to100Bin10min_5minDelay |
| Injection #: | 1 | Processing Method: | LC PQ ApexTrack |
| Injection Volume: | 3.00 ul | Channel Name: | 254.0nm |
| Run Time: | 10.0 Minutes | Proc. Chnl. Descr.: | PDA Spectrum PDA 254.0 nm |
| Date Acquired: | 7/26/2026 3:50:09 PM EDT | | |
| Date Processed: | 7/26/2026 4:05:30 PM EDT | | |

**Auto-Scaled Chromatogram**

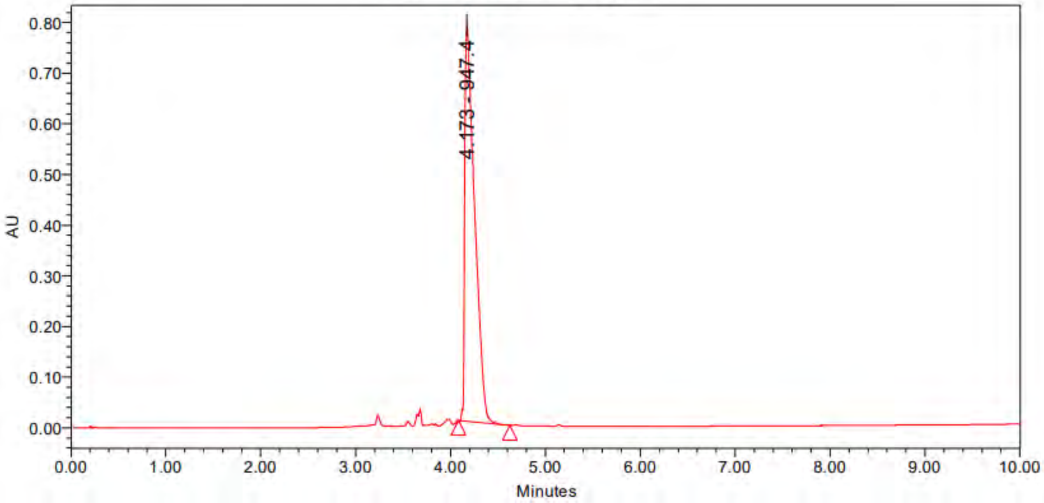

Peak Results

| | Name | RT | Area | Height | Amount | Units |
|---|---|---|---|---|---|---|
| 1 | | 4.173 | 5645918 | 781301 | | |

PDA Result Table

| | Name | RT | Purity1 Angle | Purity1 Threshold | Match1 Spect. Name | Match1 Angle | Match1 Threshold |
|---|---|---|---|---|---|---|---|
| 1 | | 4.173 | | | | | |

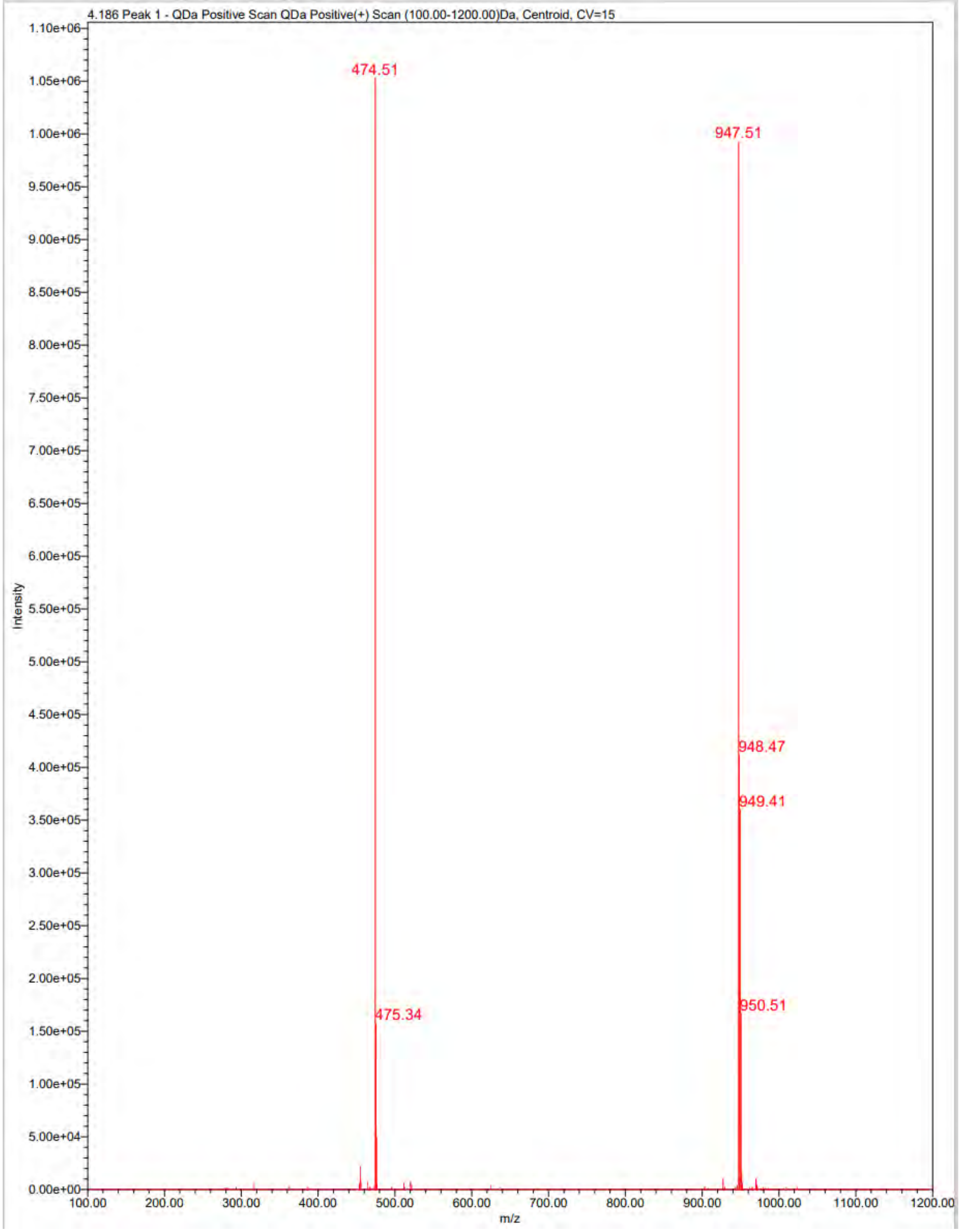

4.186 Peak 1 - QDa Positive Scan QDa Positive(+) Scan (100.00-1200.00)Da, Centroid, CV=15
474.51
947.51
948.47
949.41
475.34
950.51
Intensity
m/z
1.10e+06
1.05e+06
1.00e+06
9.50e+05
9.00e+05
8.50e+05
8.00e+05
7.50e+05
7.00e+05
6.50e+05
6.00e+05
5.50e+05
5.00e+05
4.50e+05
4.00e+05
3.50e+05
3.00e+05
2.50e+05
2.00e+05
1.50e+05
1.00e+05
5.00e+04
0.00e+00
100.00
200.00
300.00
400.00
500.00
600.00
700.00
800.00
900.00
1000.00
1100.00
1200.00

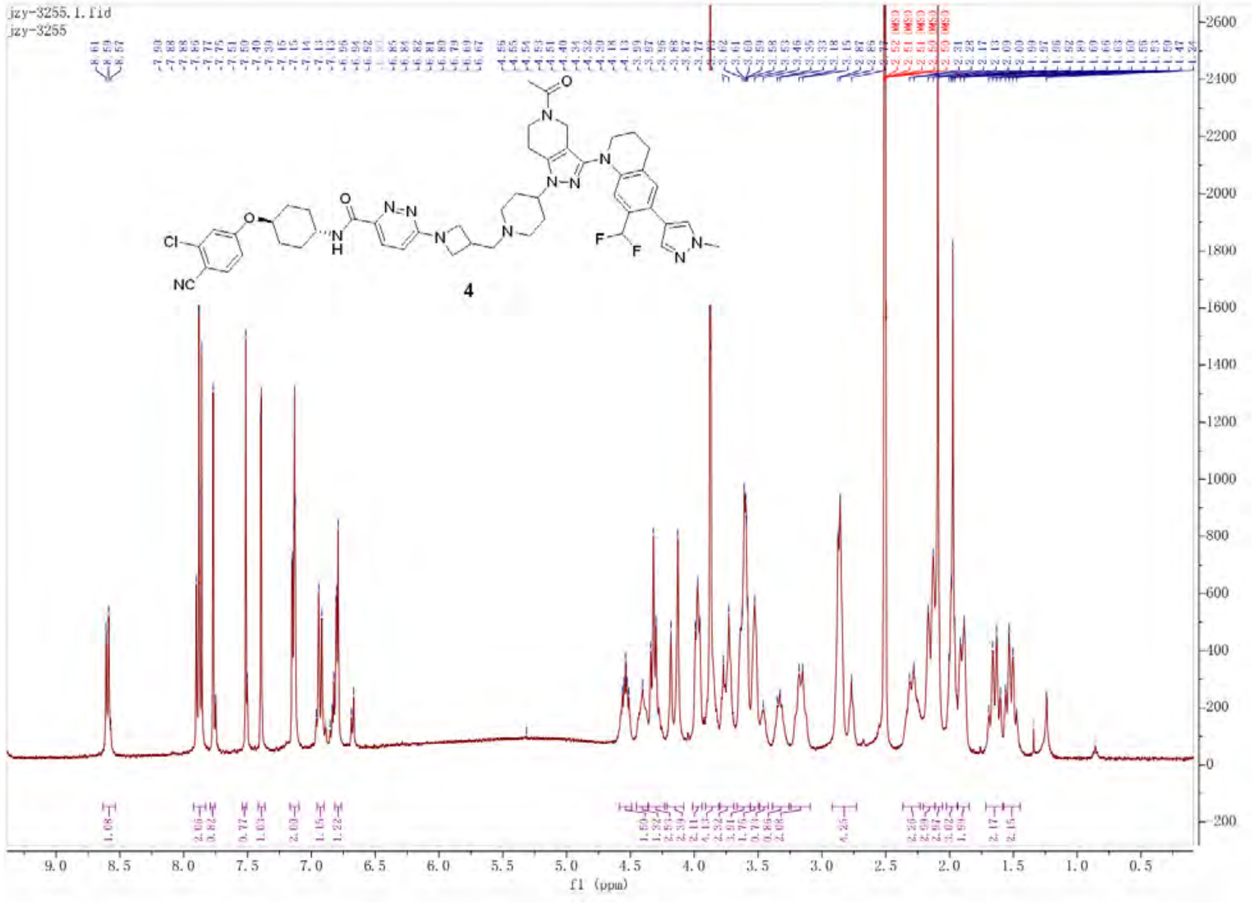

jzy-3255.1.fid
jzy-3255
4
f1 (ppm)

Empower 3

### Untitled

#### SAMPLE INFORMATION

| | | | |
|---|---|---|---|
| Sample Name: | JZY3255 | Acquired By: | System |
| Sample Type: | Unknown | Sample Set Name: | YQY |
| Vial: | 2:B,3 | Acq. Method Set: | 10to100Bin10min_5minDelay |
| Injection #: | 1 | Processing Method: | LC PQ ApexTrack |
| Injection Volume: | 3.00 ul | Channel Name: | 254.0nm |
| Run Time: | 10.0 Minutes | Proc. Chnl. Descr.: | PDA Spectrum PDA 254.0 nm |
| Date Acquired: | 7/26/2026 4:10:43 AM EDT | | |
| Date Processed: | 7/26/2026 12:07:37 PM EDT | | |

##### Auto-Scaled Chromatogram

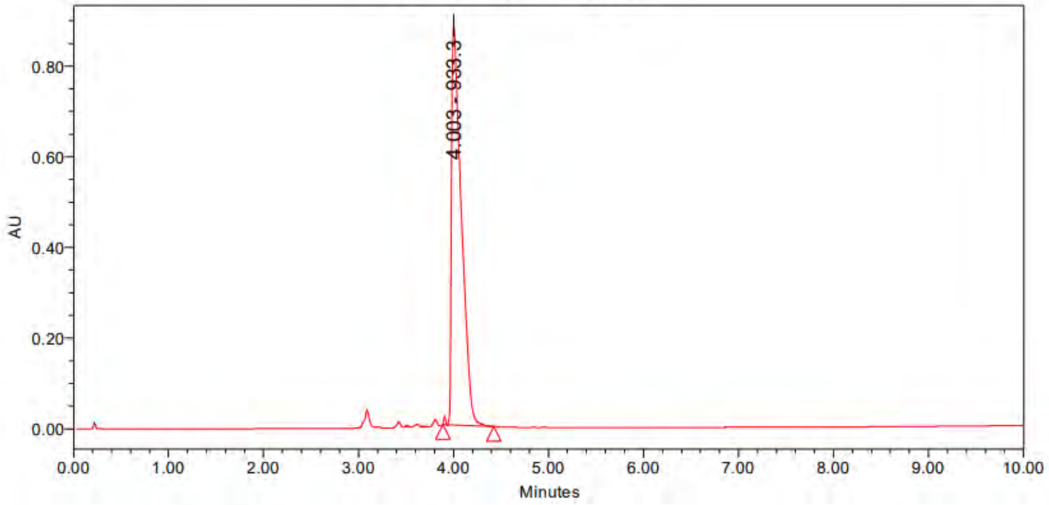

Peak Results

| | Name | RT | Area | Height | Amount | Units |
|---|---|---|---|---|---|---|
| 1 | | 4.003 | 6343332 | 881035 | | |

PDA Result Table

| | Name | RT | Purity1 Angle | Purity1 Threshold | Match1 Spect. Name | Match1 Angle | Match1 Threshold |
|---|---|---|---|---|---|---|---|
| 1 | | 4.003 | | | | | |

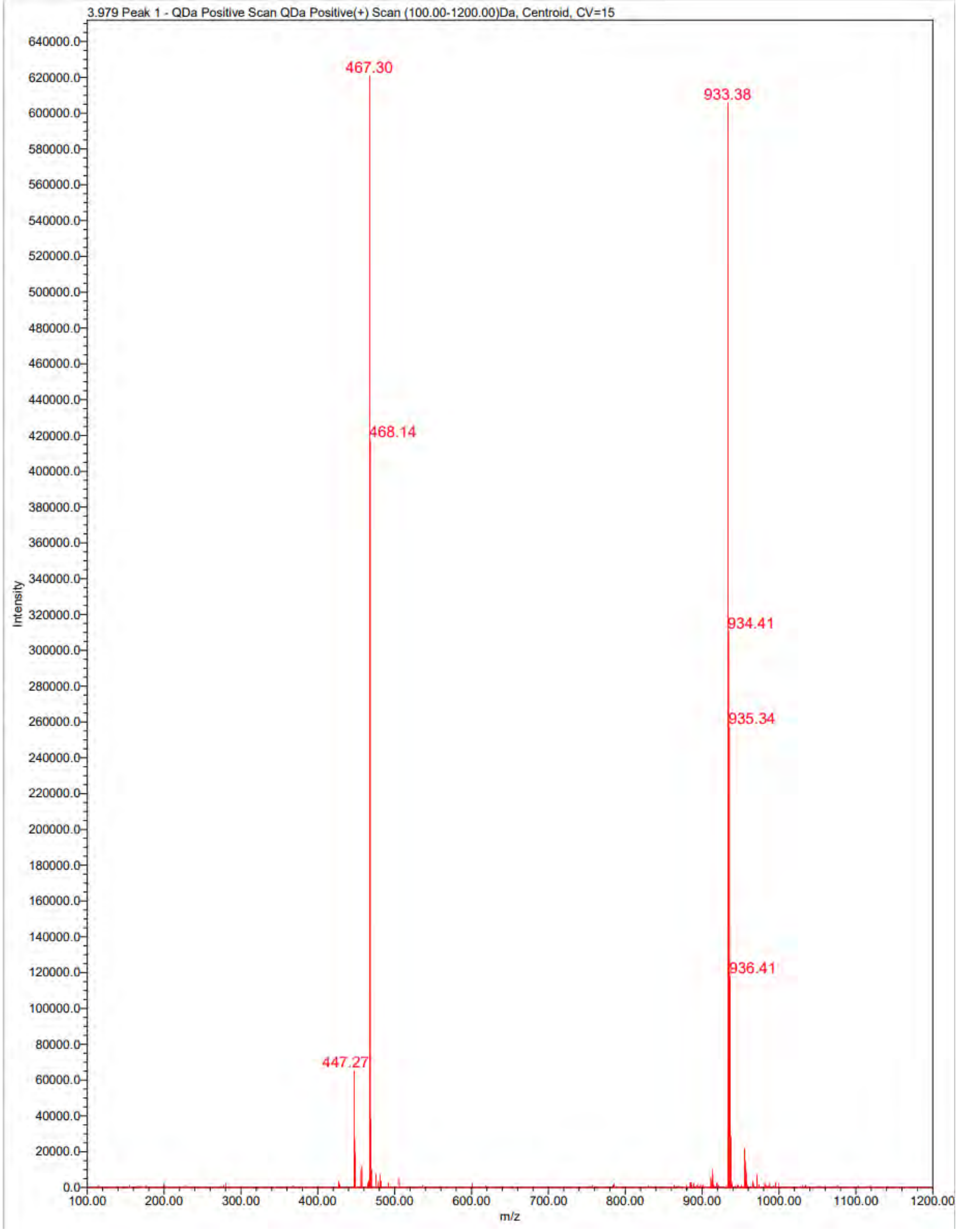
3.979 Peak 1 - QDa Positive Scan QDa Positive(+) Scan (100.00-1200.00)Da, Centroid, CV=15
467.30
933.38
468.14
934.41
935.34
936.41
447.27
Intensity
m/z
0.0
20000.0
40000.0
60000.0
80000.0
100000.0
120000.0
140000.0
160000.0
180000.0
200000.0
220000.0
240000.0
260000.0
280000.0
300000.0
320000.0
340000.0
360000.0
380000.0
400000.0
420000.0
440000.0
460000.0
480000.0
500000.0
520000.0
540000.0
560000.0
580000.0
600000.0
620000.0
640000.0
100.00
200.00
300.00
400.00
500.00
600.00
700.00
800.00
900.00
1000.00
1100.00
1200.00

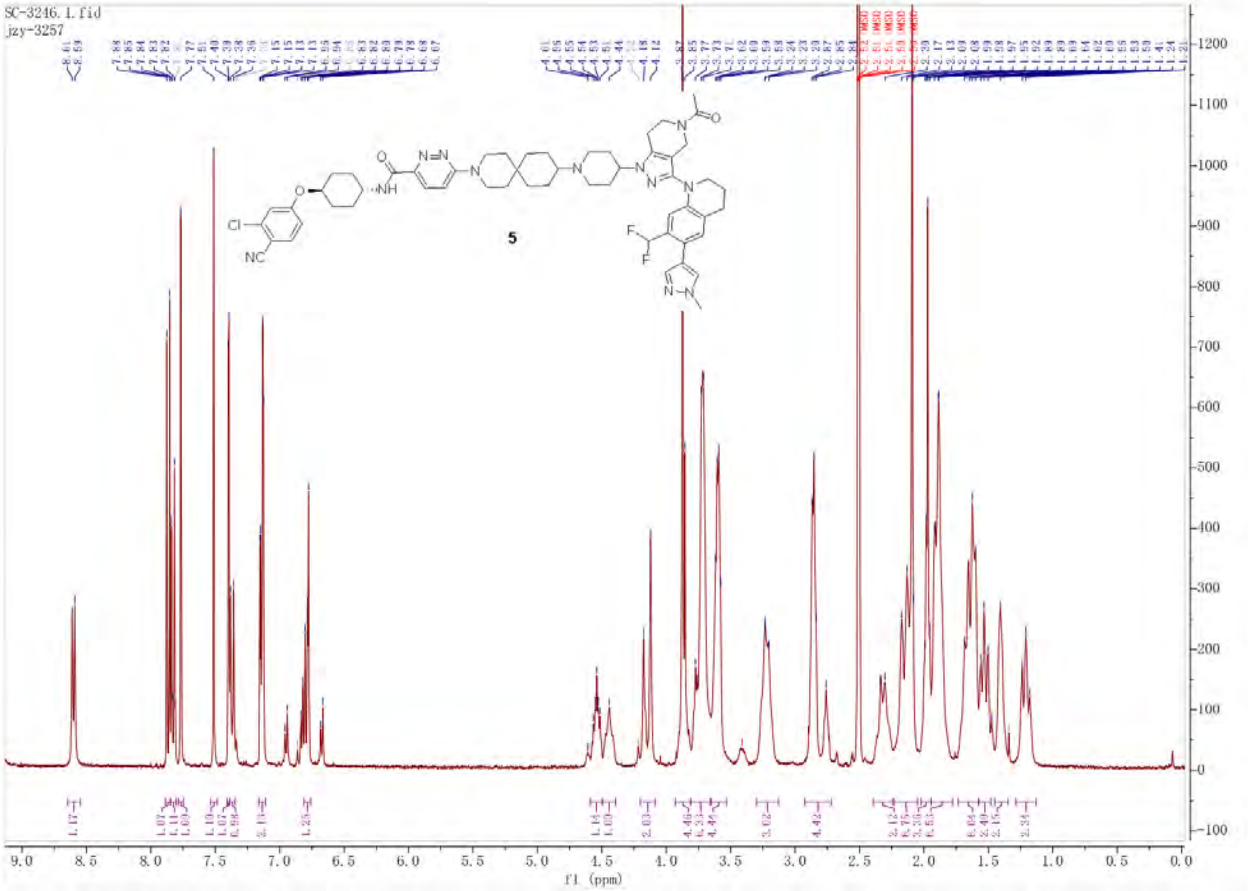
SC-3246.1.fid
jzy-3257
5
f1 (ppm)

Empower 3

Untitled

SAMPLE INFORMATION

| | | | |
|---|---|---|---|
| Sample Name: | SC-3246 | Acquired By: | System |
| Sample Type: | Unknown | Sample Set Name: | 0 |
| Vial: | 1:D,1 | Acq. Method Set: | 10to100Bin10min_5minDelay |
| Injection #: | 1 | Processing Method: | LC PQ ApexTrack |
| Injection Volume: | 3.00 ul | Channel Name: | 254.0nm |
| Run Time: | 10.0 Minutes | Proc. Chnl. Descr.: | PDA Spectrum PDA 254.0 nm |
| Date Acquired: | 7/26/2026 3:03:24 PM EDT | | |
| Date Processed: | 7/26/2026 4:31:50 PM EDT | | |

Auto-Scaled Chromatogram

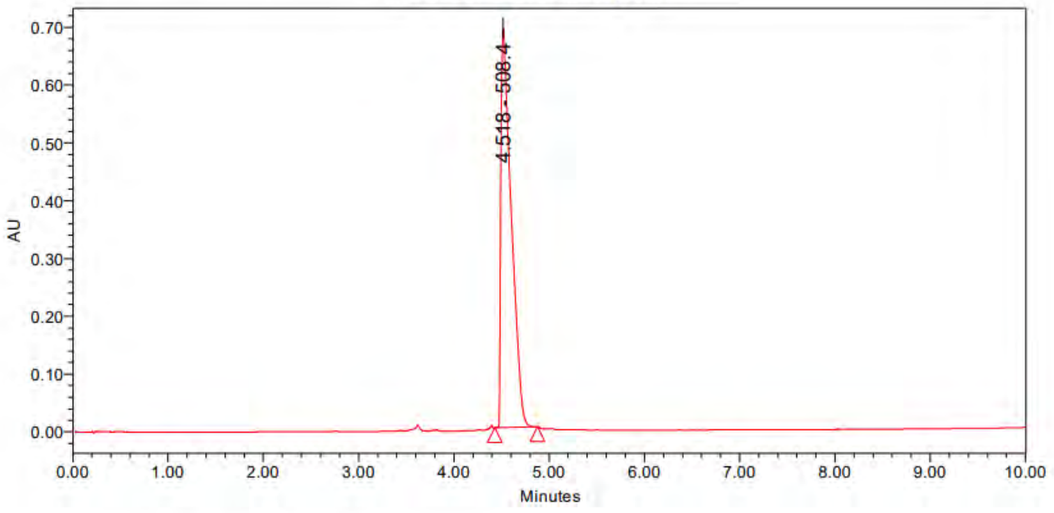

Peak Results

| | Name | RT | Area | Height | Amount | Units |
|---|---|---|---|---|---|---|
| 1 | | 4.518 | 5222713 | 690465 | | |

PDA Result Table

| | Name | RT | Purity1 Angle | Purity1 Threshold | Match1 Spect. Name | Match1 Angle | Match1 Threshold |
|---|---|---|---|---|---|---|---|
| 1 | | 4.518 | | | | | |

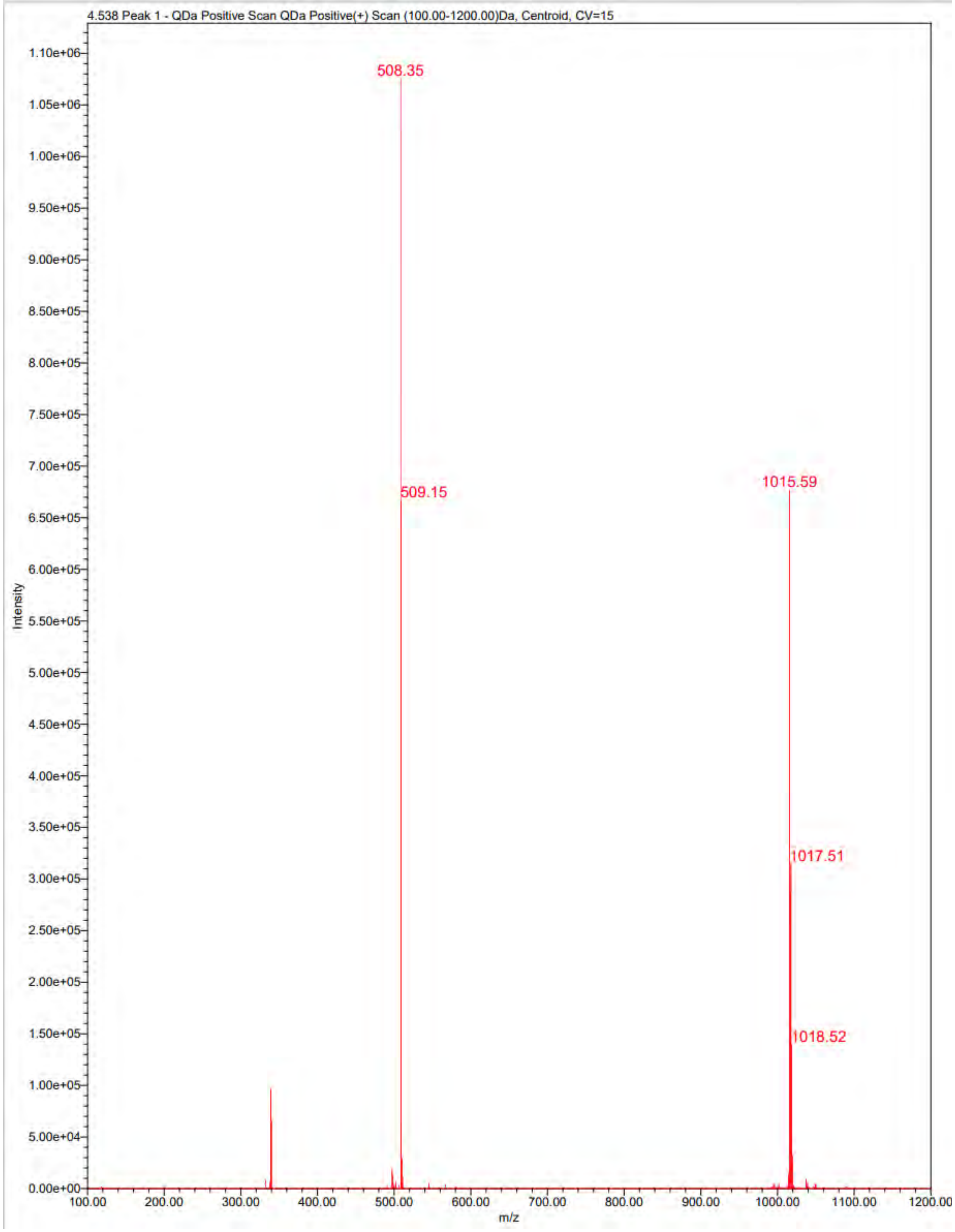
4.538 Peak 1 - QDa Positive Scan QDa Positive(+) Scan (100.00-1200.00)Da, Centroid, CV=15
508.35
509.15
1015.59
1017.51
1018.52
Intensity
m/z
1.10e+06
1.05e+06
1.00e+06
9.50e+05
9.00e+05
8.50e+05
8.00e+05
7.50e+05
7.00e+05
6.50e+05
6.00e+05
5.50e+05
5.00e+05
4.50e+05
4.00e+05
3.50e+05
3.00e+05
2.50e+05
2.00e+05
1.50e+05
1.00e+05
5.00e+04
0.00e+00
100.00
200.00
300.00
400.00
500.00
600.00
700.00
800.00
900.00
1000.00
1100.00
1200.00

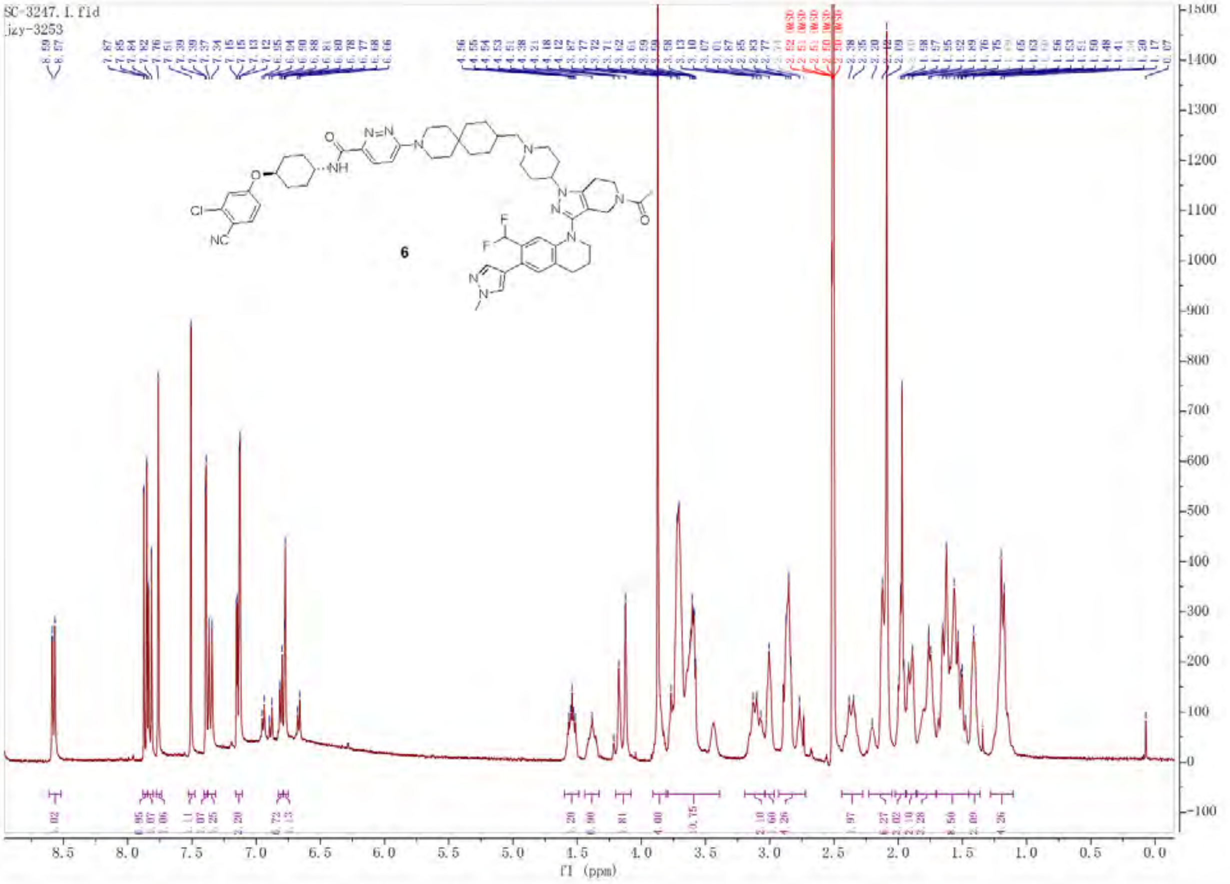

SC-3247.1.fid
jzy-3253
6
f1 (ppm)

Empower 3

### Untitled

#### SAMPLE INFORMATION

| | | | |
|---|---|---|---|
| Sample Name: | SC1 | Acquired By: | System |
| Sample Type: | Unknown | Sample Set Name: | z |
| Vial: | 1:D,7 | Acq. Method Set: | 10to100Bin10min_5minDelay |
| Injection #: | 1 | Processing Method: | LC PQ ApexTrack |
| Injection Volume: | 6.00 ul | Channel Name: | 254.0nm |
| Run Time: | 10.0 Minutes | Proc. Chnl. Descr.: | PDA Spectrum PDA 254.0 nm |
| Date Acquired: | 11/24/2025 2:09:03 PM EST | | |
| Date Processed: | 11/24/2025 2:37:28 PM EST | | |

**Auto-Scaled Chromatogram**

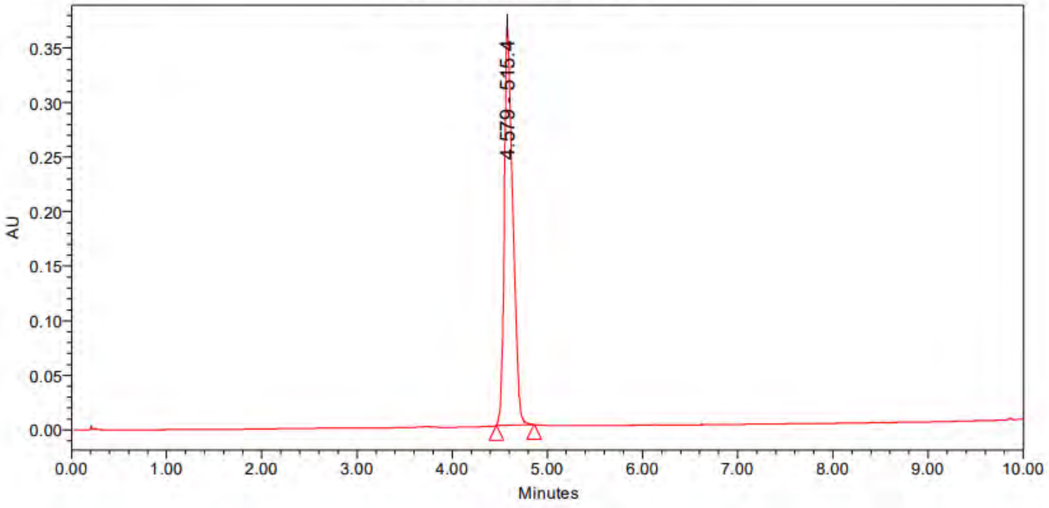

Peak Results

| | Name | RT | Area | Height | Amount | Units |
|---|---|---|---|---|---|---|
| 1 | | 4.579 | 2272959 | 366706 | | |

PDA Result Table

| | Name | RT | Purity1 Angle | Purity1 Threshold | Match1 Spect. Name | Match1 Angle | Match1 Threshold |
|---|---|---|---|---|---|---|---|
| 1 | | 4.579 | | | | | |

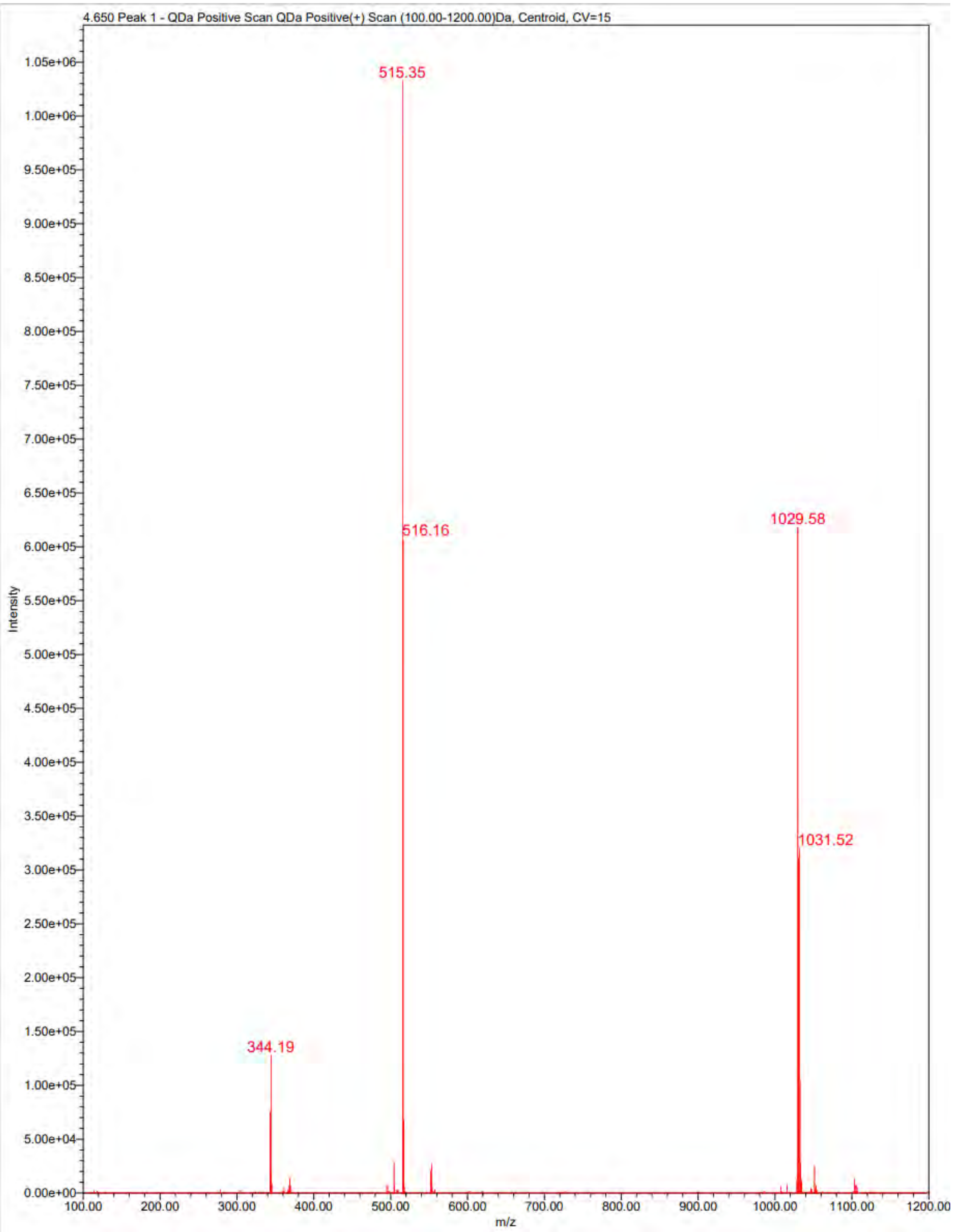
4.650 Peak 1 - QDa Positive Scan QDa Positive(+) Scan (100.00-1200.00)Da, Centroid, CV=15
515.35
516.16
1029.58
1031.52
344.19
Intensity
m/z

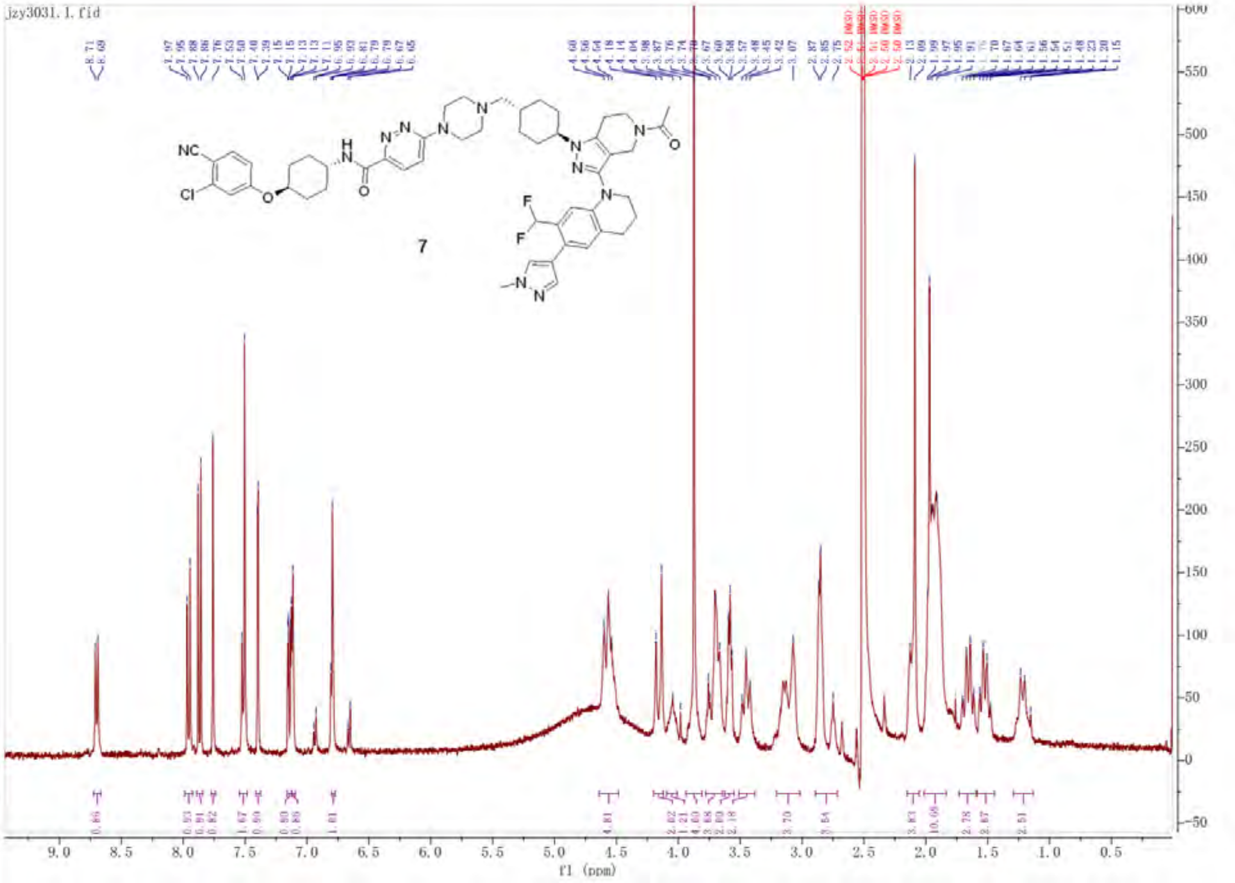
jzy3031.1.fid
NC
Cl
F
F
7
f1 (ppm)

Empower 3

### Untitled

#### SAMPLE INFORMATION

| | | | |
|---|---|---|---|
| Sample Name: | JZY3031 | Acquired By: | System |
| Sample Type: | Unknown | Sample Set Name: | YQY |
| Vial: | 2:A,2 | Acq. Method Set: | 10to100Bin10min_5minDelay |
| Injection #: | 1 | Processing Method: | LC PQ ApexTrack |
| Injection Volume: | 3.00 ul | Channel Name: | 254.0nm@1 |
| Run Time: | 10.0 Minutes | Proc. Chnl. Descr.: | PDA Spectrum PDA 254.0 nm |
| Date Acquired: | 7/26/2026 1:34:48 AM EDT | | |
| Date Processed: | 7/26/2026 11:59:38 AM EDT | | |

##### Auto-Scaled Chromatogram

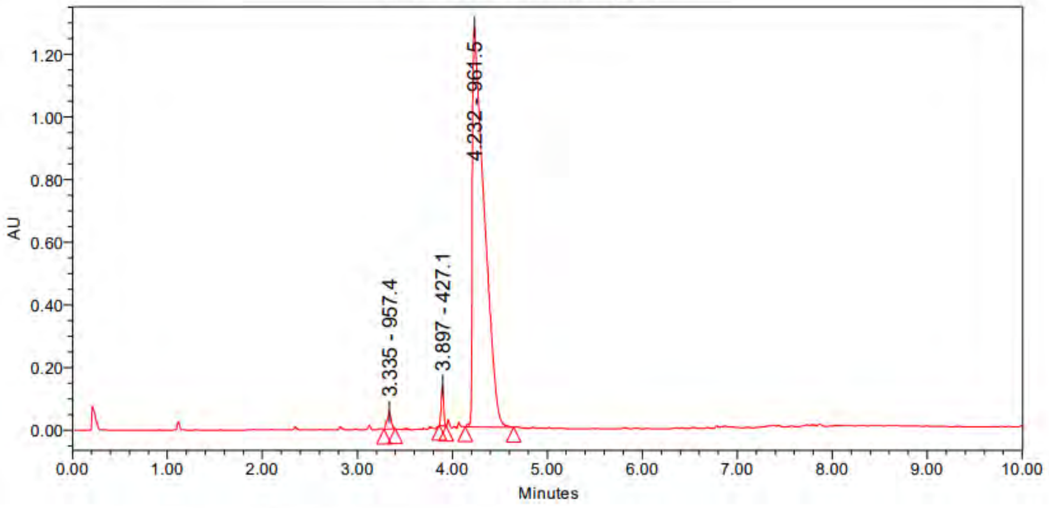

Peak Results

| | Name | RT | Area | Height | Amount | Units |
|---|---|---|---|---|---|---|
| 1 | | 3.335 | 167840 | 54402 | | |
| 2 | | 3.897 | 225921 | 128472 | | |
| 3 | | 4.232 | 11033798 | 1276873 | | |

PDA Result Table

| | Name | RT | Purity1 Angle | Purity1 Threshold | Match1 Spect. Name | Match1 Angle | Match1 Threshold |
|---|---|---|---|---|---|---|---|
| 1 | | 3.335 | | | | | |
| 2 | | 3.897 | | | | | |
| 3 | | 4.232 | | | | | |

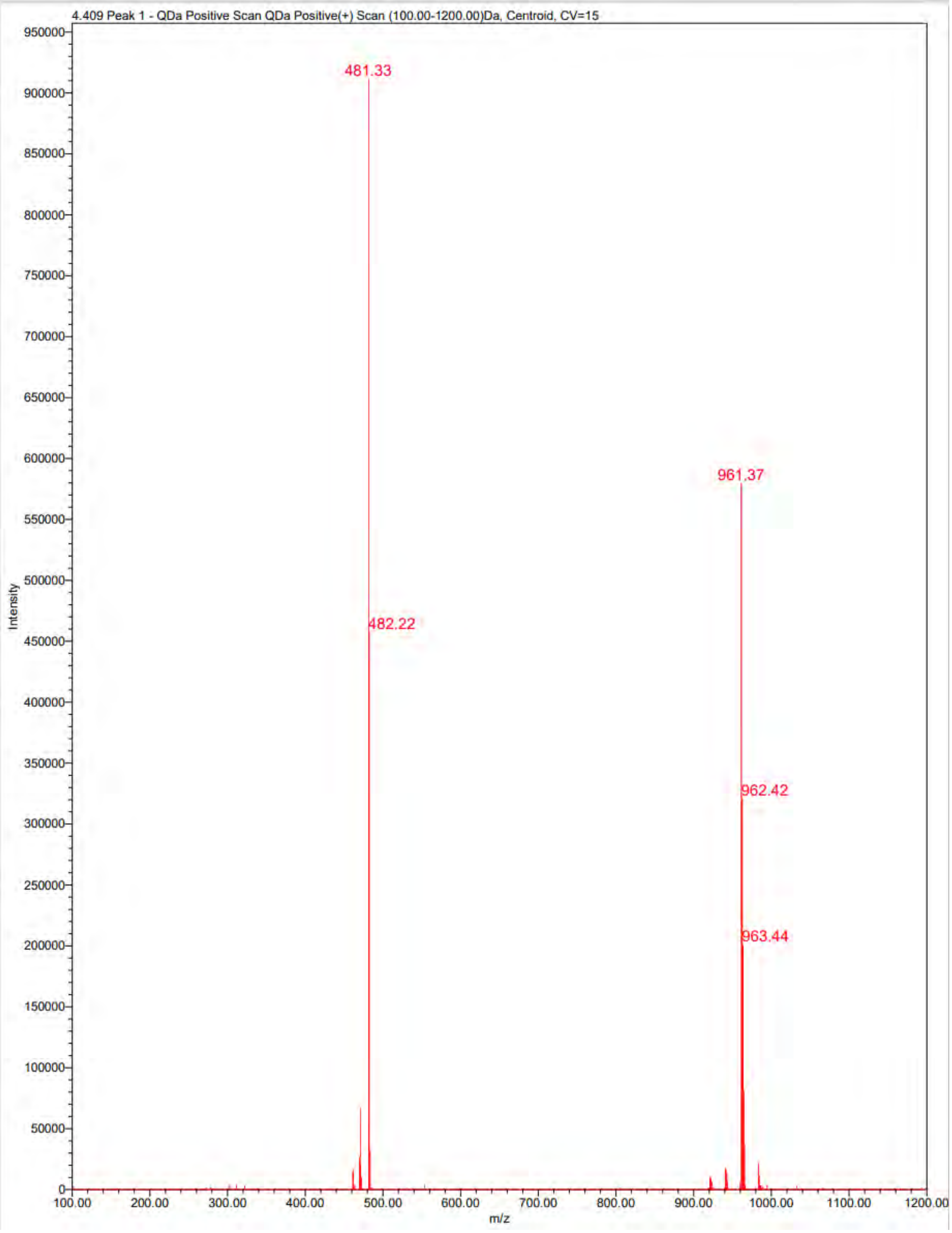

4.409 Peak 1 - QDa Positive Scan QDa Positive(+) Scan (100.00-1200.00)Da, Centroid, CV=15
481.33
482.22
961.37
962.42
963.44
Intensity
m/z

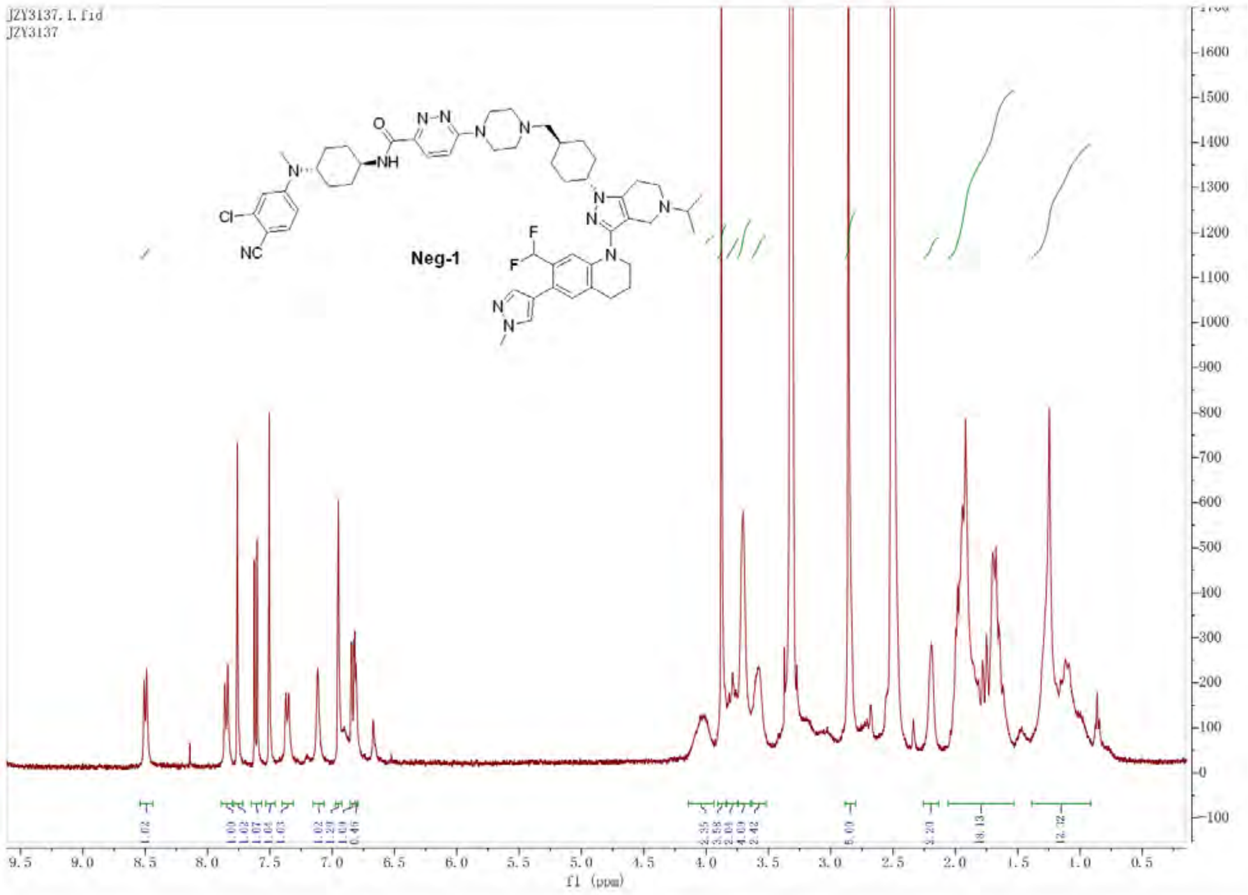

JZY3137.1.fid
JZY3137
Neg-1
f1 (ppm)

Untitled

#### SAMPLE INFORMATION

| | | | |
|---|---|---|---|
| Sample Name: | JZY3137 | Acquired By: | System |
| Sample Type: | Unknown | Sample Set Name: | YQY |
| Vial: | 2:A,4 | Acq. Method Set: | 10to100Bin10min_5minDelay |
| Injection #: | 1 | Processing Method: | LC PQ ApexTrack |
| Injection Volume: | 3.00 ul | Channel Name: | 254.0nm@2 |
| Run Time: | 10.0 Minutes | Proc. Chnl. Descr.: | PDA Spectrum PDA 254.0 nm |
| Date Acquired: | 7/26/2026 2:21:36 AM EDT | | |
| Date Processed: | 7/26/2026 12:02:54 PM EDT | | |

##### Auto-Scaled Chromatogram

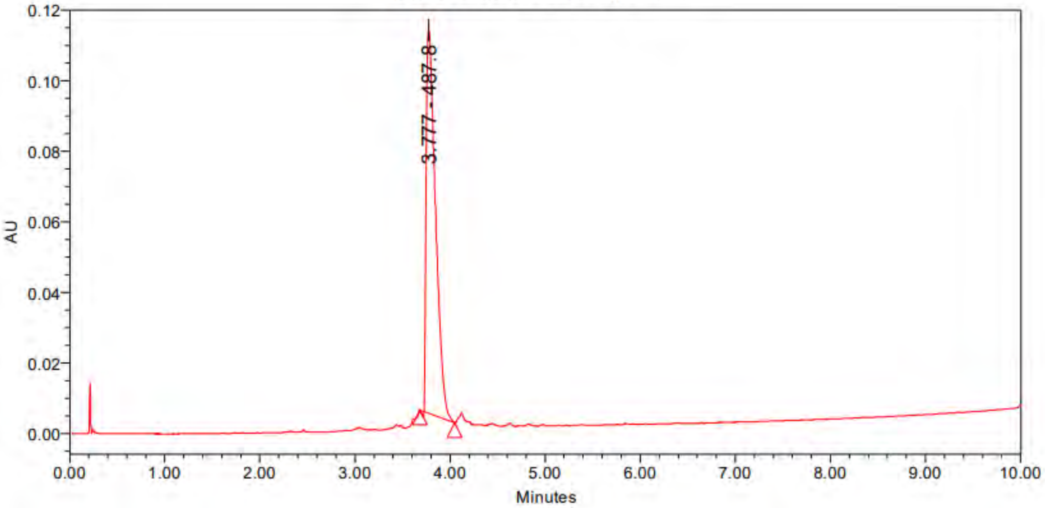

Peak Results

| | Name | RT | Area | Height | Amount | Units |
|---|---|---|---|---|---|---|
| 1 | | 3.777 | 740649 | 108655 | | |

PDA Result Table

| | Name | RT | Purity1 Angle | Purity1 Threshold | Match1 Spect. Name | Match1 Angle | Match1 Threshold |
|---|---|---|---|---|---|---|---|
| 1 | | 3.777 | | | | | |

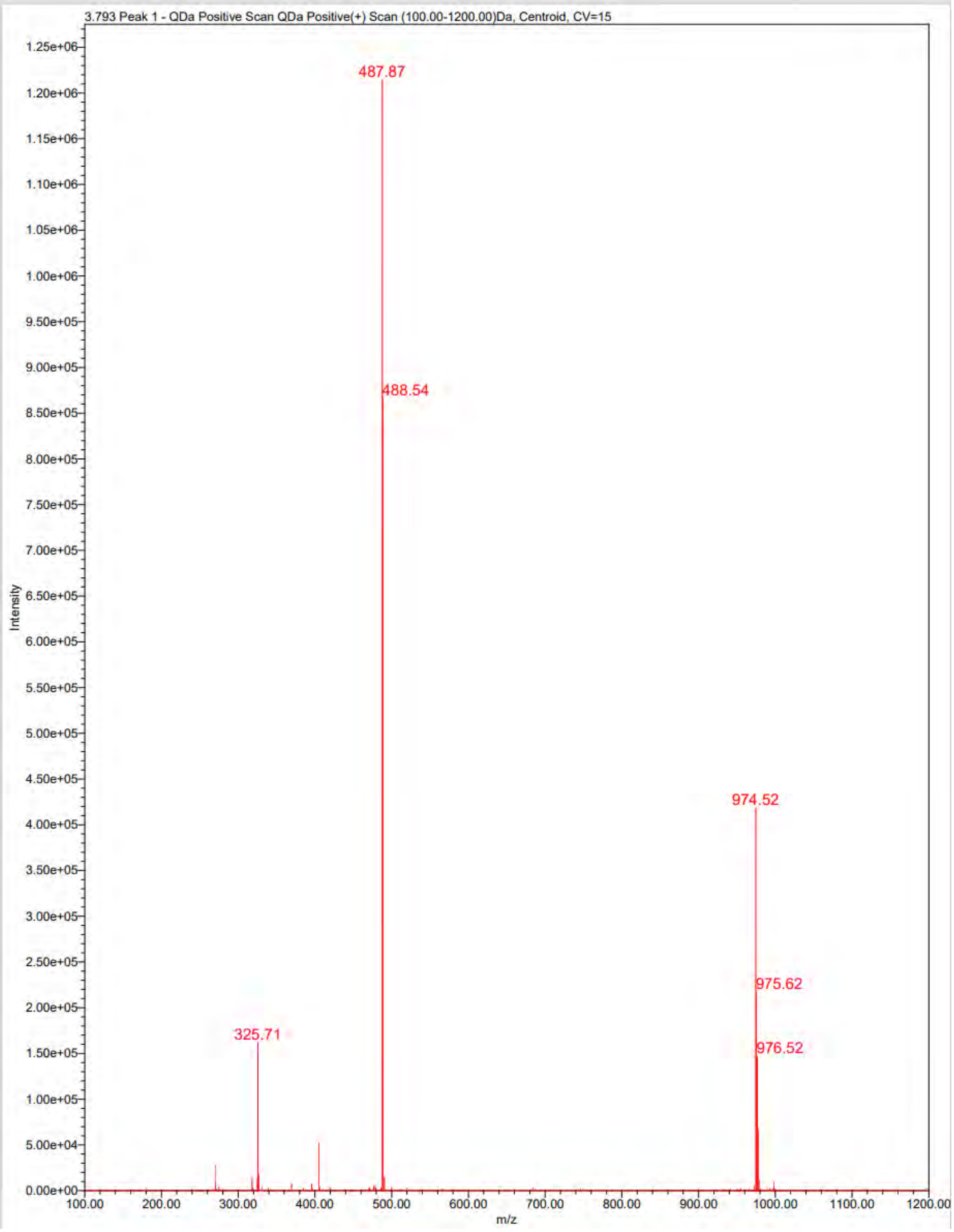

3.793 Peak 1 - QDa Positive Scan QDa Positive(+) Scan (100.00-1200.00)Da, Centroid, CV=15
487.87
488.54
974.52
975.62
976.52
325.71
Intensity
m/z
1.25e+06
1.20e+06
1.15e+06
1.10e+06
1.05e+06
1.00e+06
9.50e+05
9.00e+05
8.50e+05
8.00e+05
7.50e+05
7.00e+05
6.50e+05
6.00e+05
5.50e+05
5.00e+05
4.50e+05
4.00e+05
3.50e+05
3.00e+05
2.50e+05
2.00e+05
1.50e+05
1.00e+05
5.00e+04
0.00e+00
100.00
200.00
300.00
400.00
500.00
600.00
700.00
800.00
900.00
1000.00
1100.00
1200.00

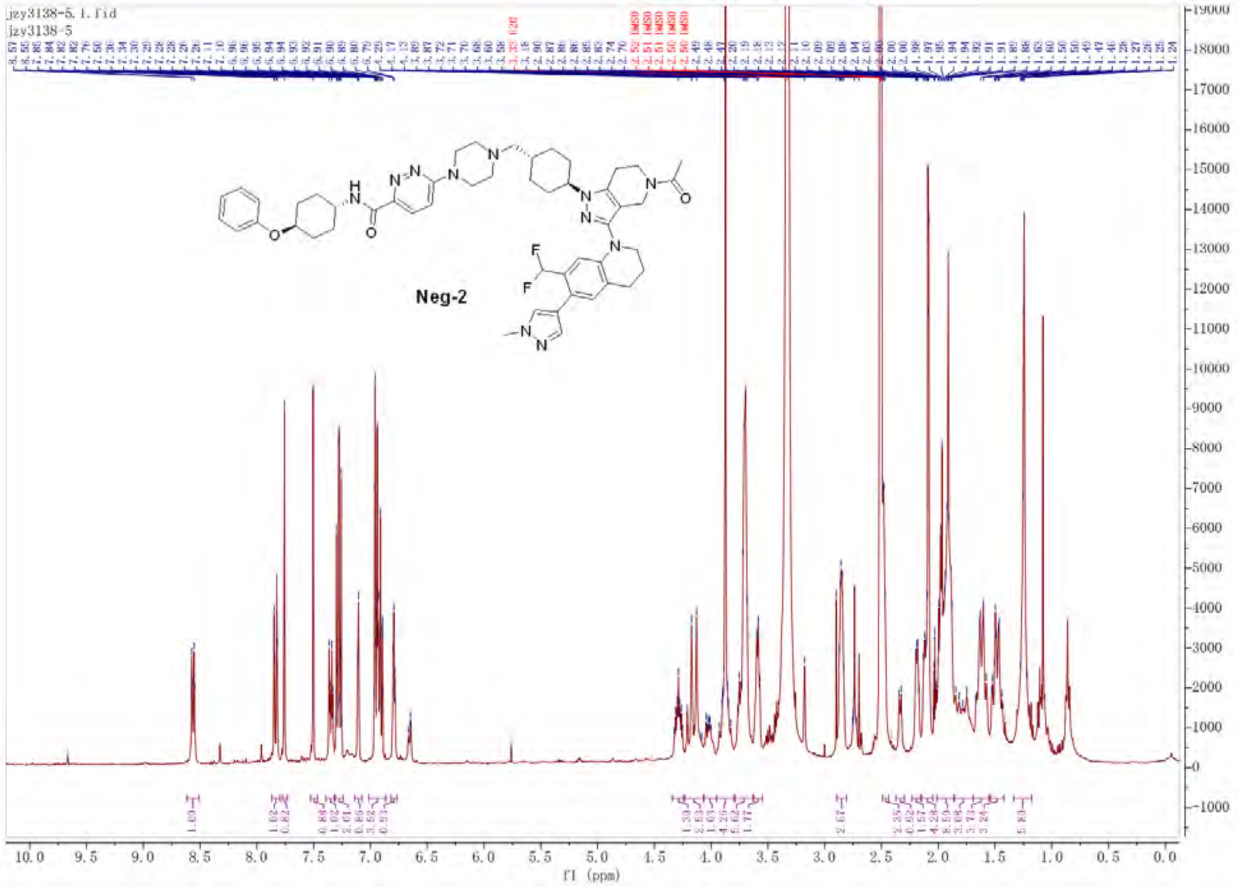
jzy3138-5.1.fid
jzy3138-5
Neg-2
f1 (ppm)

Empower 3

### Untitled

#### SAMPLE INFORMATION

| | | | |
|---|---|---|---|
| Sample Name: | JZY3138 | Acquired By: | System |
| Sample Type: | Unknown | Sample Set Name: | 0 |
| Vial: | 1:D,3 | Acq. Method Set: | 10to100Bin10min_5minDelay |
| Injection #: | 1 | Processing Method: | LC PQ ApexTrack |
| Injection Volume: | 3.00 ul | Channel Name: | 254.0nm |
| Run Time: | 10.0 Minutes | Proc. Chnl. Descr.: | PDA Spectrum PDA 254.0 nm |
| Date Acquired: | 7/26/2026 3:34:34 PM EDT | | |
| Date Processed: | 7/26/2026 3:51:25 PM EDT | | |

**Auto-Scaled Chromatogram**

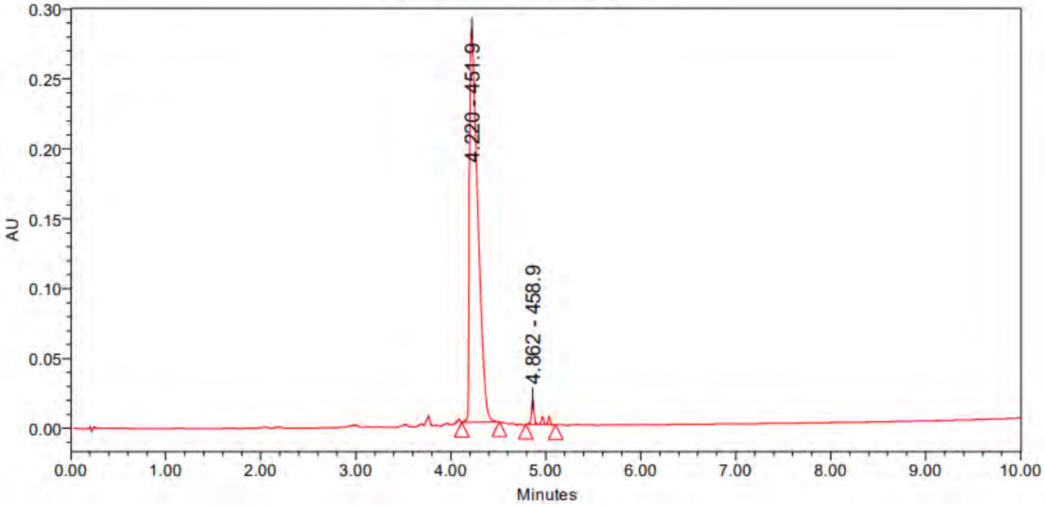

Peak Results

| | Name | RT | Area | Height | Amount | Units |
|---|---|---|---|---|---|---|
| 1 | | 4.220 | 1716851 | 281727 | | |
| 2 | | 4.862 | 53281 | 18688 | | |

PDA Result Table

| | Name | RT | Purity1 Angle | Purity1 Threshold | Match1 Spect. Name | Match1 Angle | Match1 Threshold |
|---|---|---|---|---|---|---|---|
| 1 | | 4.220 | | | | | |
| 2 | | 4.862 | | | | | |

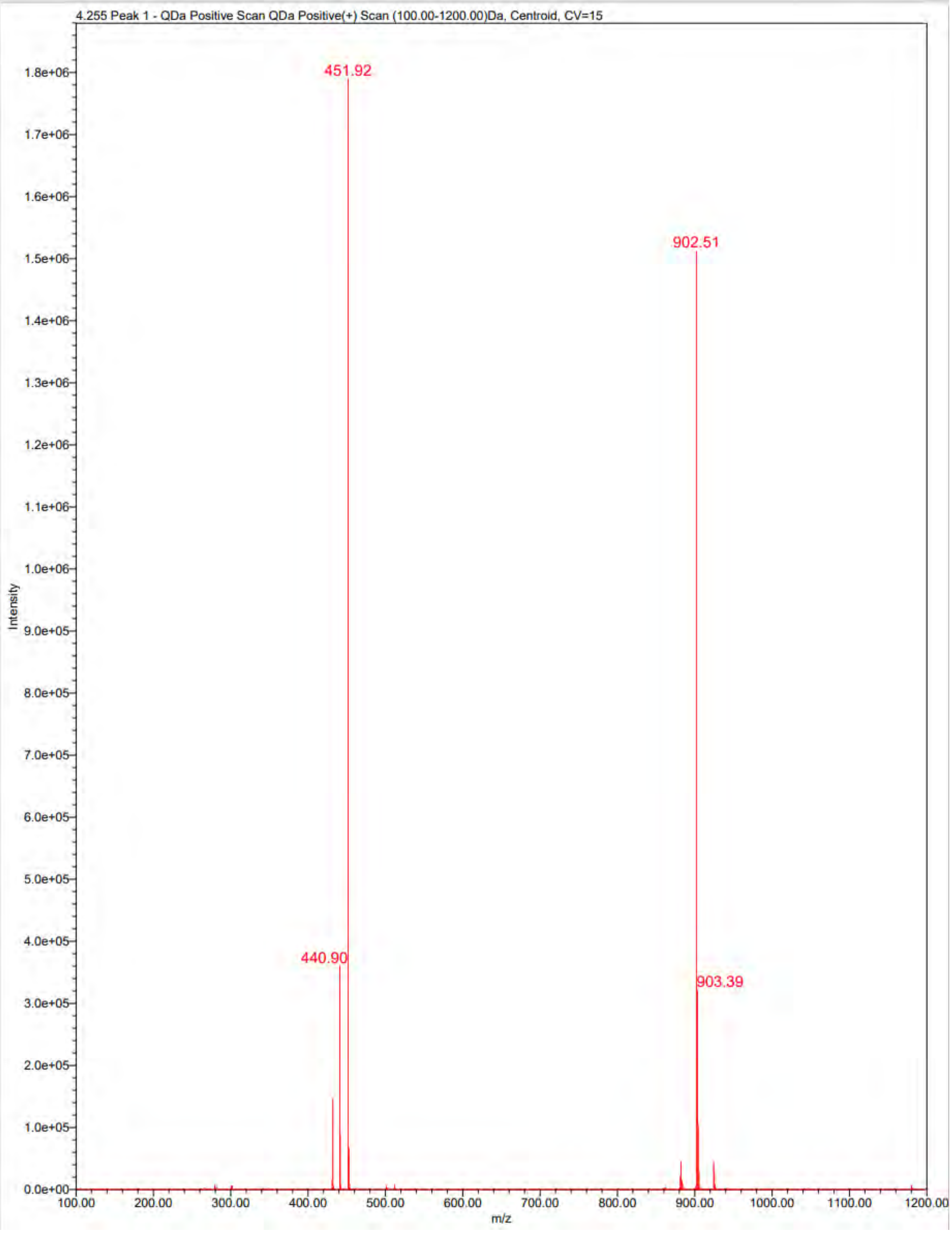

4.255 Peak 1 - QDa Positive Scan QDa Positive(+) Scan (100.00-1200.00)Da, Centroid, CV=15
451.92
902.51
440.90
903.39
Intensity
m/z
1.8e+06
1.7e+06
1.6e+06
1.5e+06
1.4e+06
1.3e+06
1.2e+06
1.1e+06
1.0e+06
9.0e+05
8.0e+05
7.0e+05
6.0e+05
5.0e+05
4.0e+05
3.0e+05
2.0e+05
1.0e+05
0.0e+00
100.00
200.00
300.00
400.00
500.00
600.00
700.00
800.00
900.00
1000.00
1100.00
1200.00

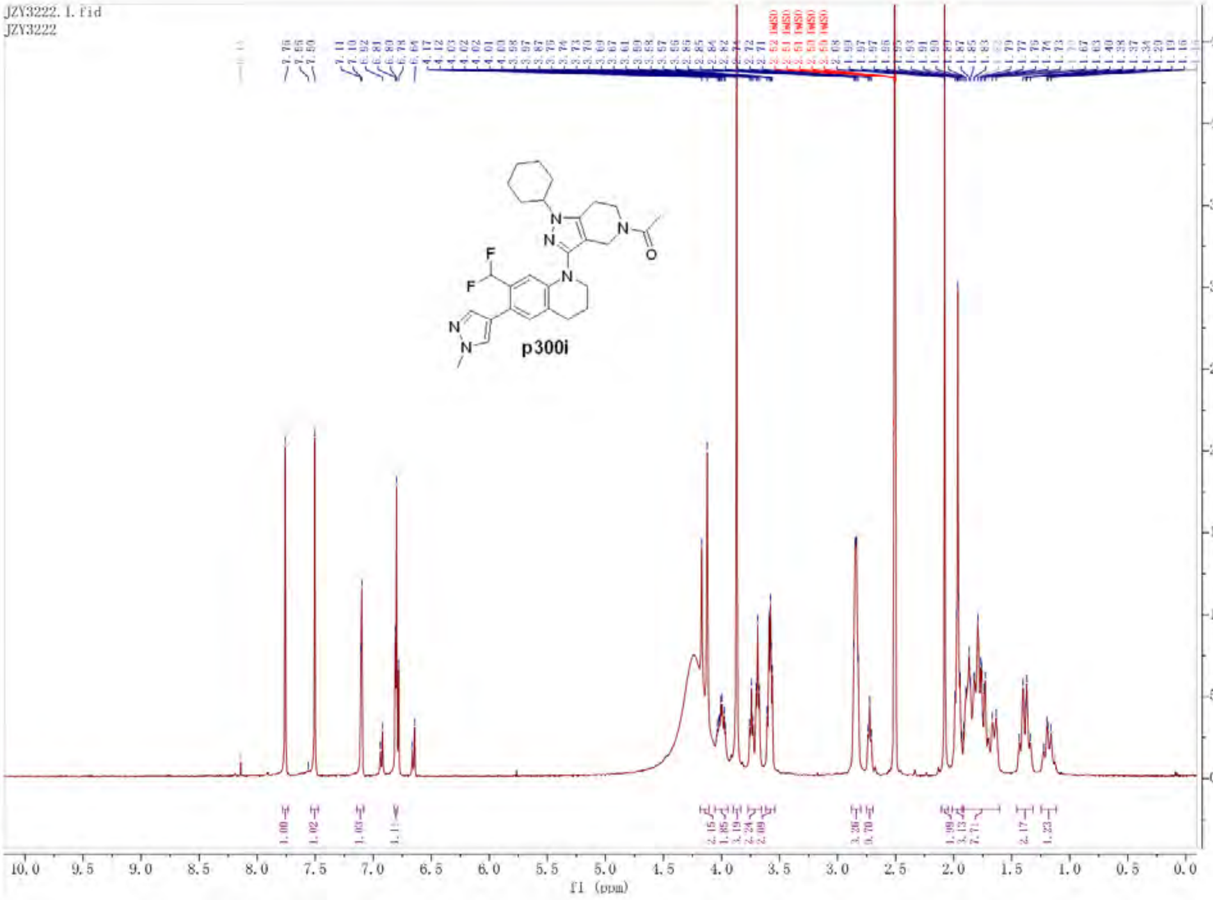
JZY3222.1.fid
JZY3222
p300i
f1 (ppm)

Empower 3

### Untitled

#### SAMPLE INFORMATION

| | | | |
|---|---|---|---|
| Sample Name: | JZY3222 | Acquired By: | System |
| Sample Type: | Unknown | Sample Set Name: | 0 |
| Vial: | 1:C,8 | Acq. Method Set: | 10to100Bin10min_5minDelay |
| Injection #: | 1 | Processing Method: | LC PQ ApexTrack |
| Injection Volume: | 3.00 ul | Channel Name: | 254.0nm |
| Run Time: | 10.0 Minutes | Proc. Chnl. Descr.: | PDA Spectrum PDA 254.0 nm |
| Date Acquired: | 7/26/2026 2:47:50 PM EDT | | |
| Date Processed: | 7/26/2026 3:46:19 PM EDT | | |

##### Auto-Scaled Chromatogram

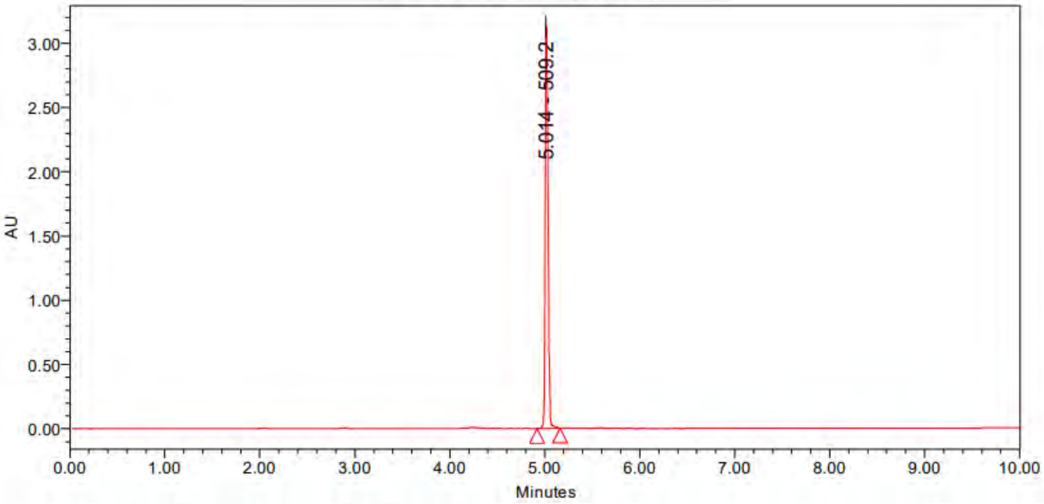

Peak Results

| | Name | RT | Area | Height | Amount | Units |
|---|---|---|---|---|---|---|
| 1 | | 5.014 | 6438227 | 3134093 | | |

PDA Result Table

| | Name | RT | Purity1 Angle | Purity1 Threshold | Match1 Spect. Name | Match1 Angle | Match1 Threshold |
|---|---|---|---|---|---|---|---|
| 1 | | 5.014 | | | | | |

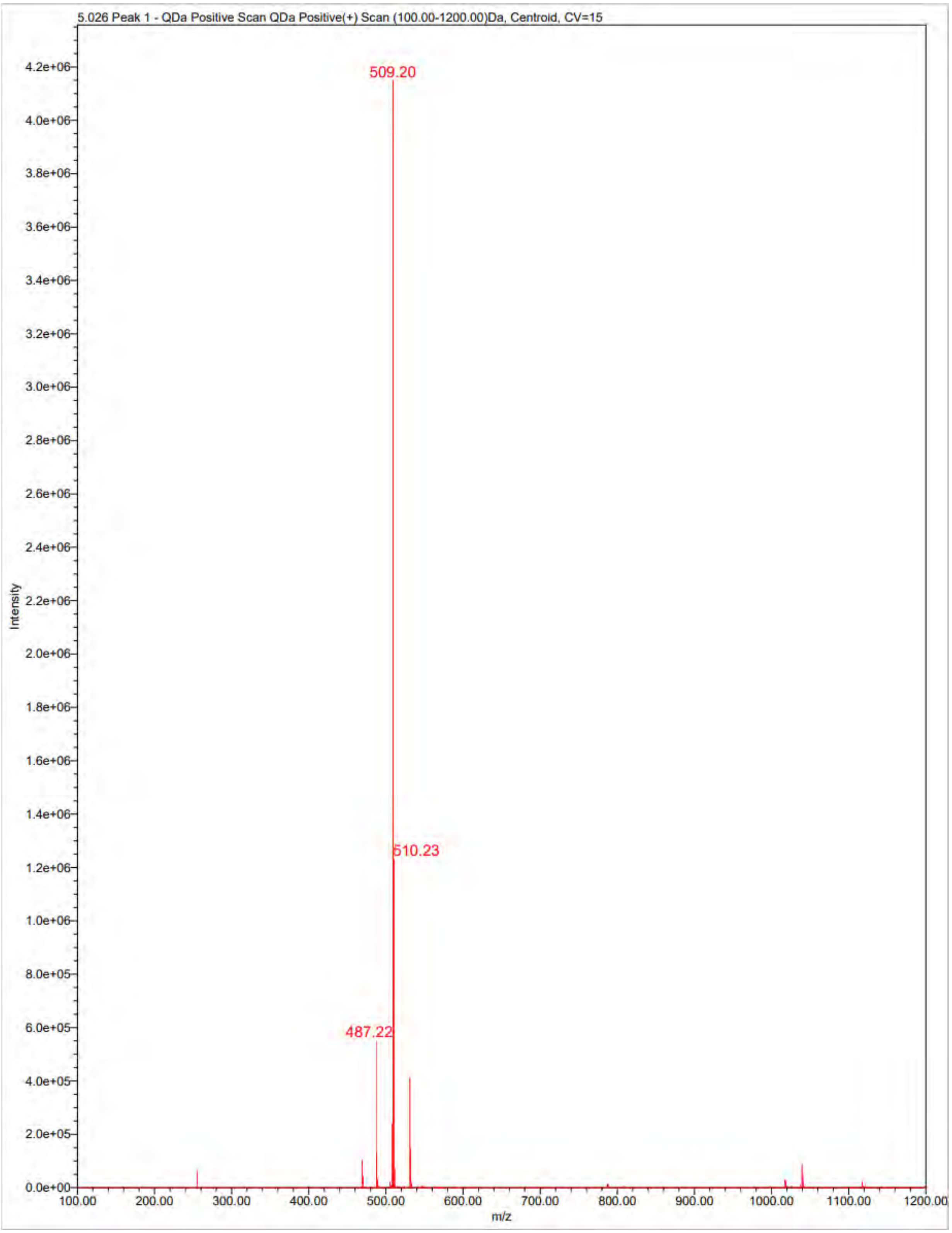
5.026 Peak 1 - QDa Positive Scan QDa Positive(+) Scan (100.00-1200.00)Da, Centroid, CV=15
509.20
510.23
487.22
Intensity
m/z
4.2e+06
4.0e+06
3.8e+06
3.6e+06
3.4e+06
3.2e+06
3.0e+06
2.8e+06
2.6e+06
2.4e+06
2.2e+06
2.0e+06
1.8e+06
1.6e+06
1.4e+06
1.2e+06
1.0e+06
8.0e+05
6.0e+05
4.0e+05
2.0e+05
0.0e+00
100.00
200.00
300.00
400.00
500.00
600.00
700.00
800.00
900.00
1000.00
1100.00
1200.00

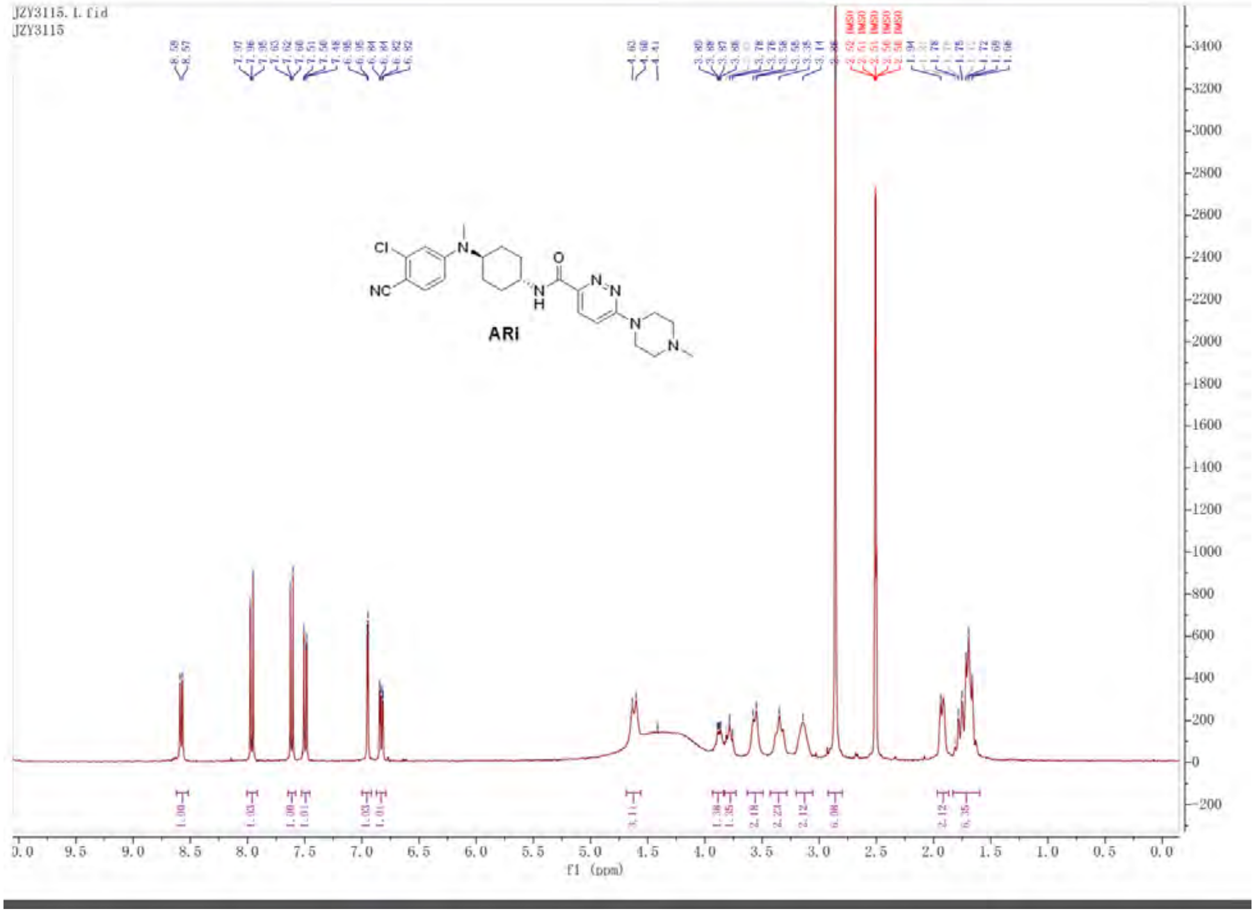

JZY3115.1.fid
JZY3115
ARi
f1 (ppm)

Untitled

#### SAMPLE INFORMATION

| | | | |
|---|---|---|---|
| Sample Name: | jzy3115 | Acquired By: | System |
| Sample Type: | Unknown | Sample Set Name: | 0 |
| Vial: | 1:C,3 | Acq. Method Set: | 10to100Bin10min_5minDelay |
| Injection #: | 1 | Processing Method: | LC PQ ApexTrack |
| Injection Volume: | 3.00 ul | Channel Name: | 254.0nm |
| Run Time: | 10.0 Minutes | Proc. Chnl. Descr.: | PDA Spectrum PDA 254.0 nm |
| Date Acquired: | 7/26/2026 1:29:52 PM EDT | | |
| Date Processed: | 7/26/2026 1:46:47 PM EDT | | |

**Auto-Scaled Chromatogram**

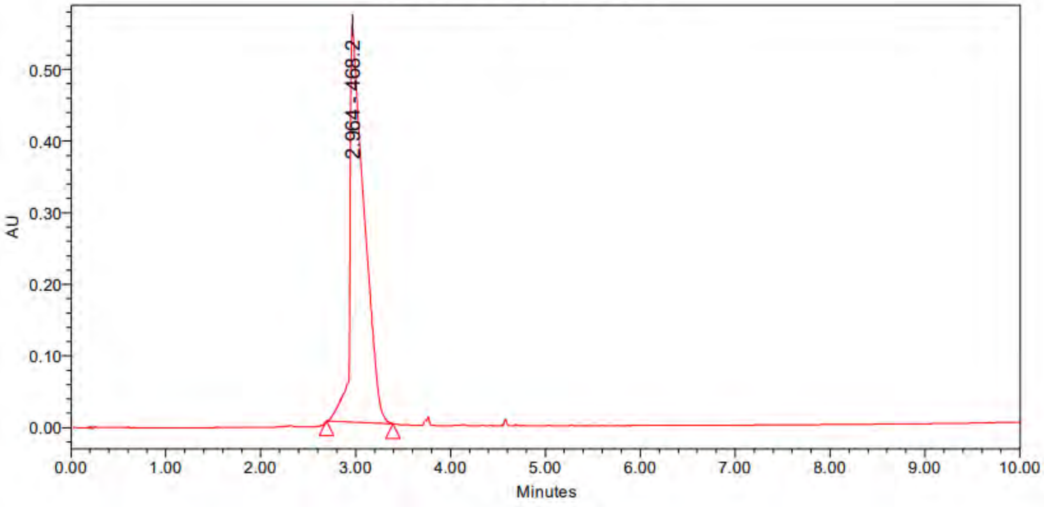

Peak Results

| | Name | RT | Area | Height | Amount | Units |
|---|---|---|---|---|---|---|
| 1 | | 2.964 | 5851768 | 554210 | | |

PDA Result Table

| | Name | RT | Purity1 Angle | Purity1 Threshold | Match1 Spect. Name | Match1 Angle | Match1 Threshold |
|---|---|---|---|---|---|---|---|
| 1 | | 2.964 | | | | | |

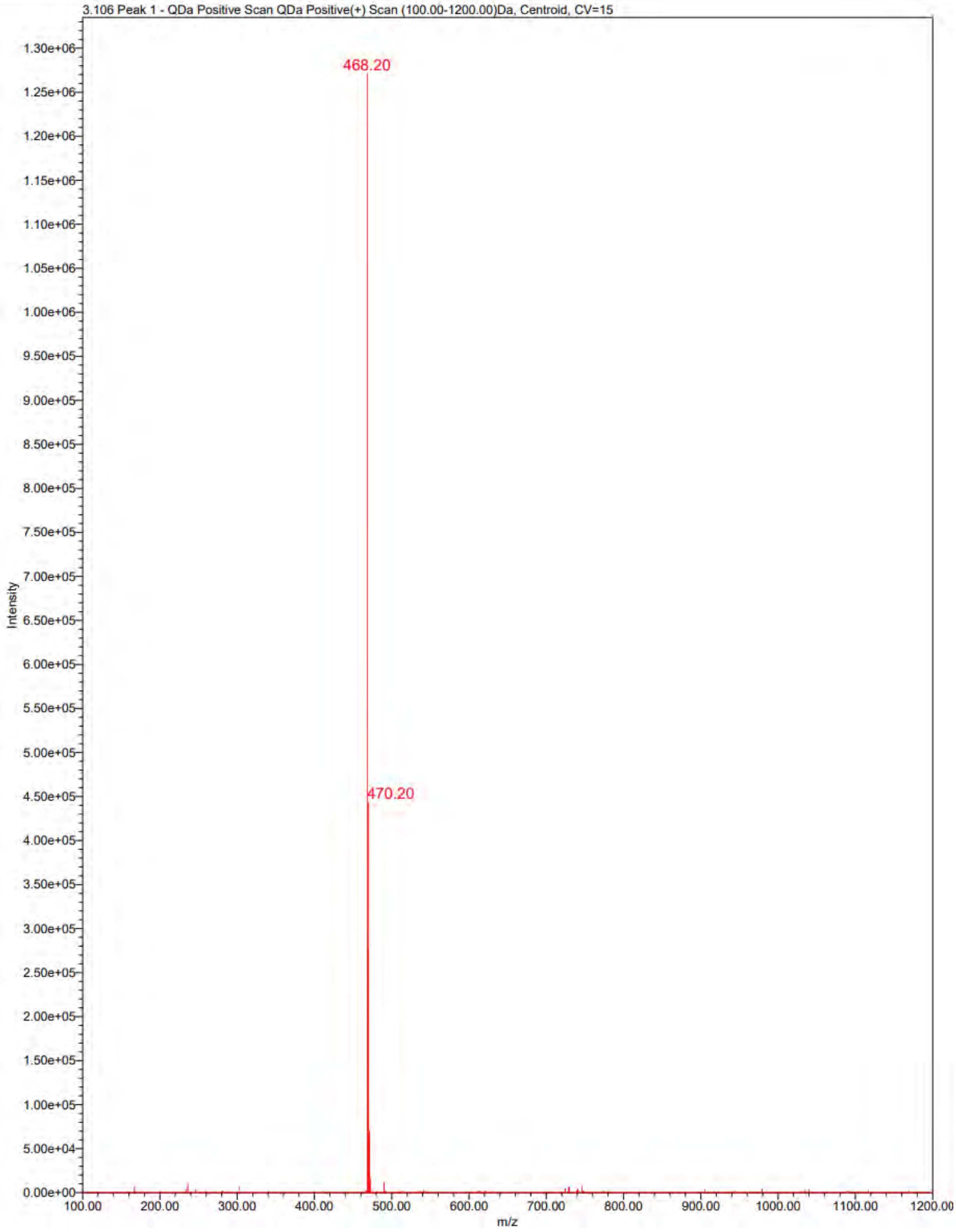
3.106 Peak 1 - QDa Positive Scan QDa Positive(+) Scan (100.00-1200.00)Da, Centroid, CV=15
468.20
470.20
Intensity
m/z

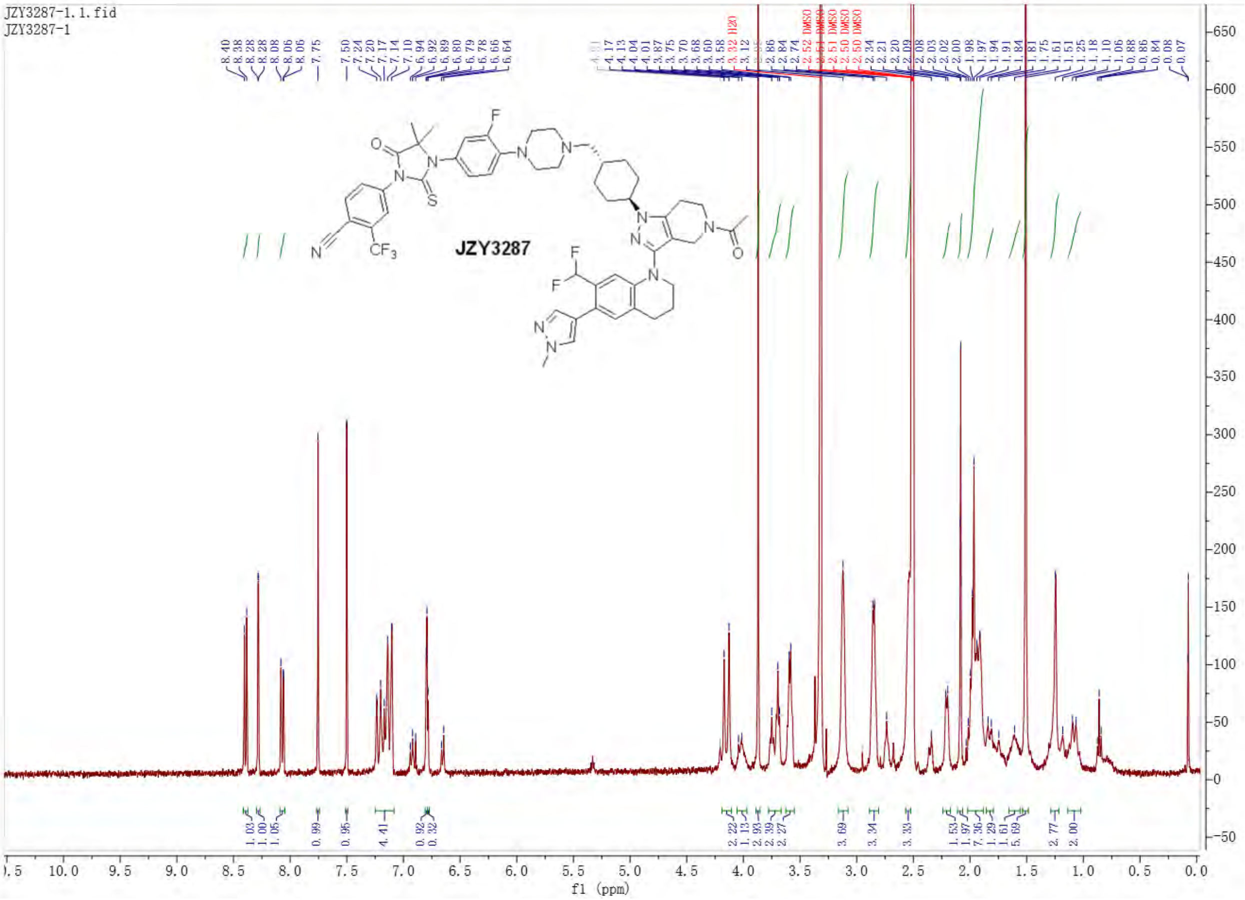

JZY3287-1.1.fid
JZY3287-1
JZY3287
f1 (ppm)

Untitled

#### SAMPLE INFORMATION

| | | | |
|---|---|---|---|
| Sample Name: | JZY3287 | Acquired By: | System |
| Sample Type: | Unknown | Sample Set Name: | YQY |
| Vial: | 2:B,2 | Acq. Method Set: | 10to100Bin10min_5minDelay |
| Injection #: | 1 | Processing Method: | LC PQ ApexTrack |
| Injection Volume: | 3.00 ul | Channel Name: | 254.0nm |
| Run Time: | 10.0 Minutes | Proc. Chnl. Descr.: | PDA Spectrum PDA 254.0 nm |
| Date Acquired: | 7/26/2026 3:55:09 AM EDT | | |
| Date Processed: | 7/26/2026 12:07:13 PM EDT | | |

##### Auto-Scaled Chromatogram

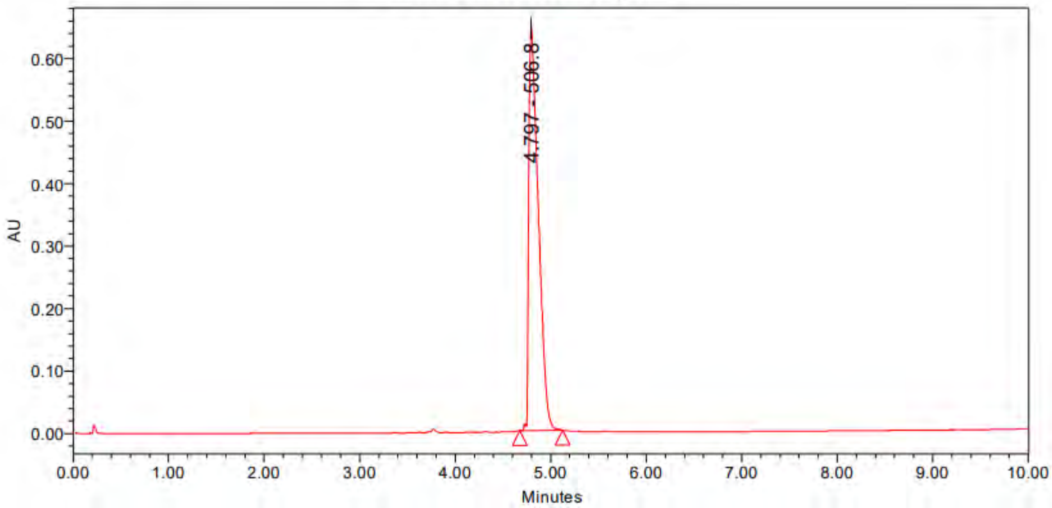

Peak Results

| | Name | RT | Area | Height | Amount | Units |
|---|---|---|---|---|---|---|
| 1 | | 4.797 | 4431921 | 644348 | | |

PDA Result Table

| | Name | RT | Purity1 Angle | Purity1 Threshold | Match1 Spect. Name | Match1 Angle | Match1 Threshold |
|---|---|---|---|---|---|---|---|
| 1 | | 4.797 | | | | | |

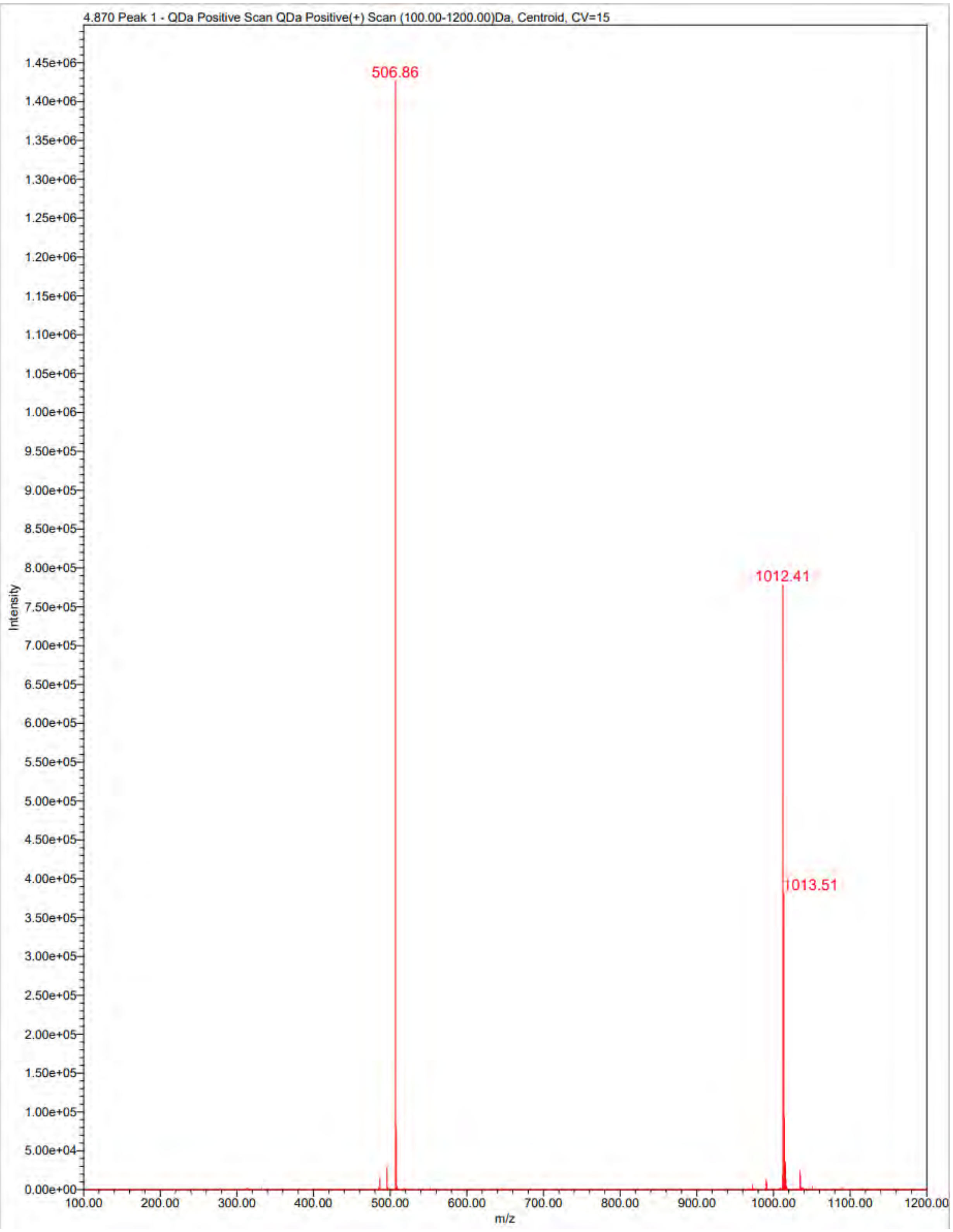

4.870 Peak 1 - QDa Positive Scan QDa Positive(+) Scan (100.00-1200.00)Da, Centroid, CV=15
506.86
1012.41
1013.51
Intensity
m/z

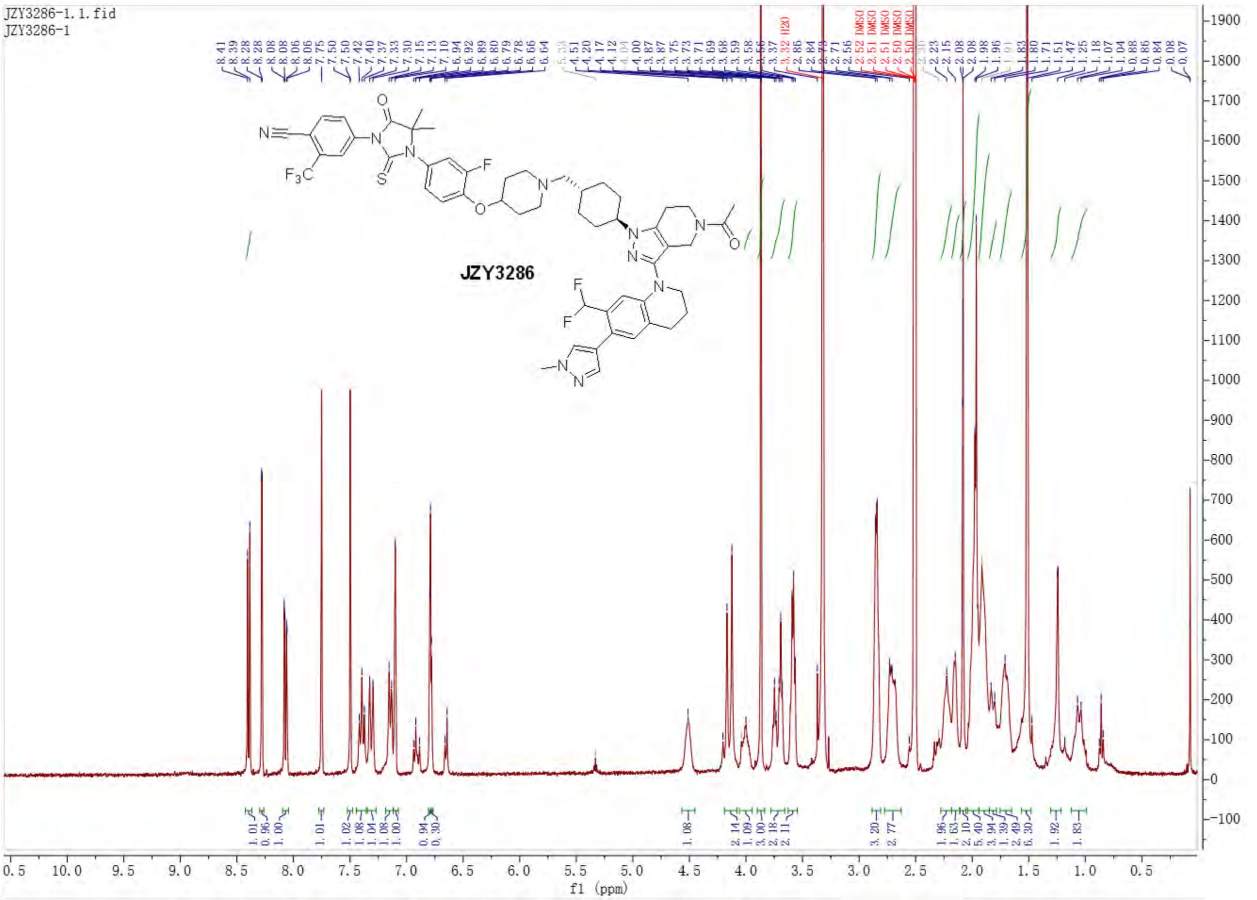
JZY3286-1.1.fid
JZY3286-1
JZY3286
f1 (ppm)

Untitled

#### SAMPLE INFORMATION

| | | | |
|---|---|---|---|
| Sample Name: | JZY3286 | Acquired By: | System |
| Sample Type: | Unknown | Sample Set Name: | YQY |
| Vial: | 2:A,5 | Acq. Method Set: | 10to100Bin10min_5minDelay |
| Injection #: | 1 | Processing Method: | LC PQ ApexTrack |
| Injection Volume: | 3.00 ul | Channel Name: | 254.0nm |
| Run Time: | 10.0 Minutes | Proc. Chnl. Descr.: | PDA Spectrum PDA 254.0 nm |
| Date Acquired: | 7/26/2026 2:37:11 AM EDT | | |
| Date Processed: | 7/26/2026 12:03:16 PM EDT | | |

**Auto-Scaled Chromatogram**

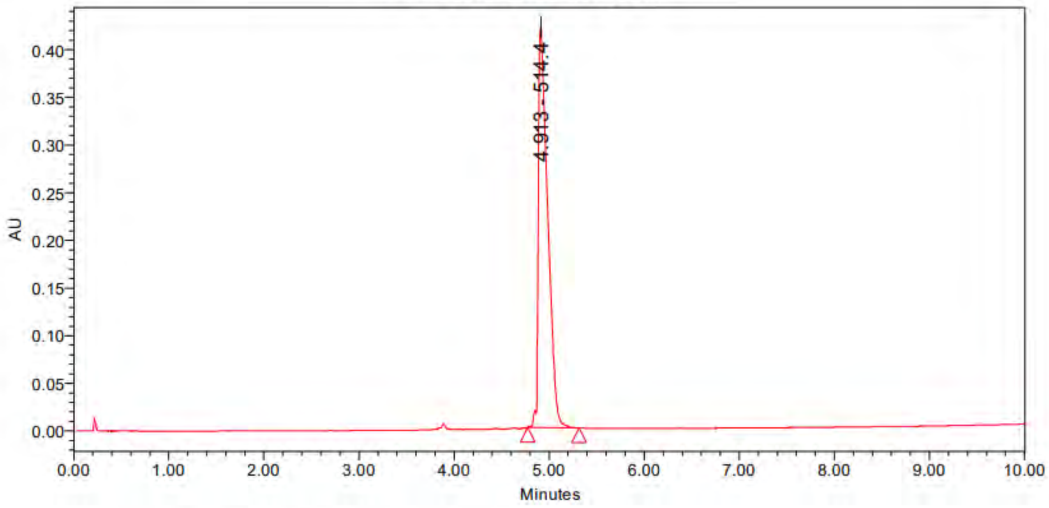

Peak Results

| | Name | RT | Area | Height | Amount | Units |
|---|---|---|---|---|---|---|
| 1 | | 4.913 | 2882350 | 419432 | | |

PDA Result Table

| | Name | RT | Purity1 Angle | Purity1 Threshold | Match1 Spect. Name | Match1 Angle | Match1 Threshold |
|---|---|---|---|---|---|---|---|
| 1 | | 4.913 | | | | | |

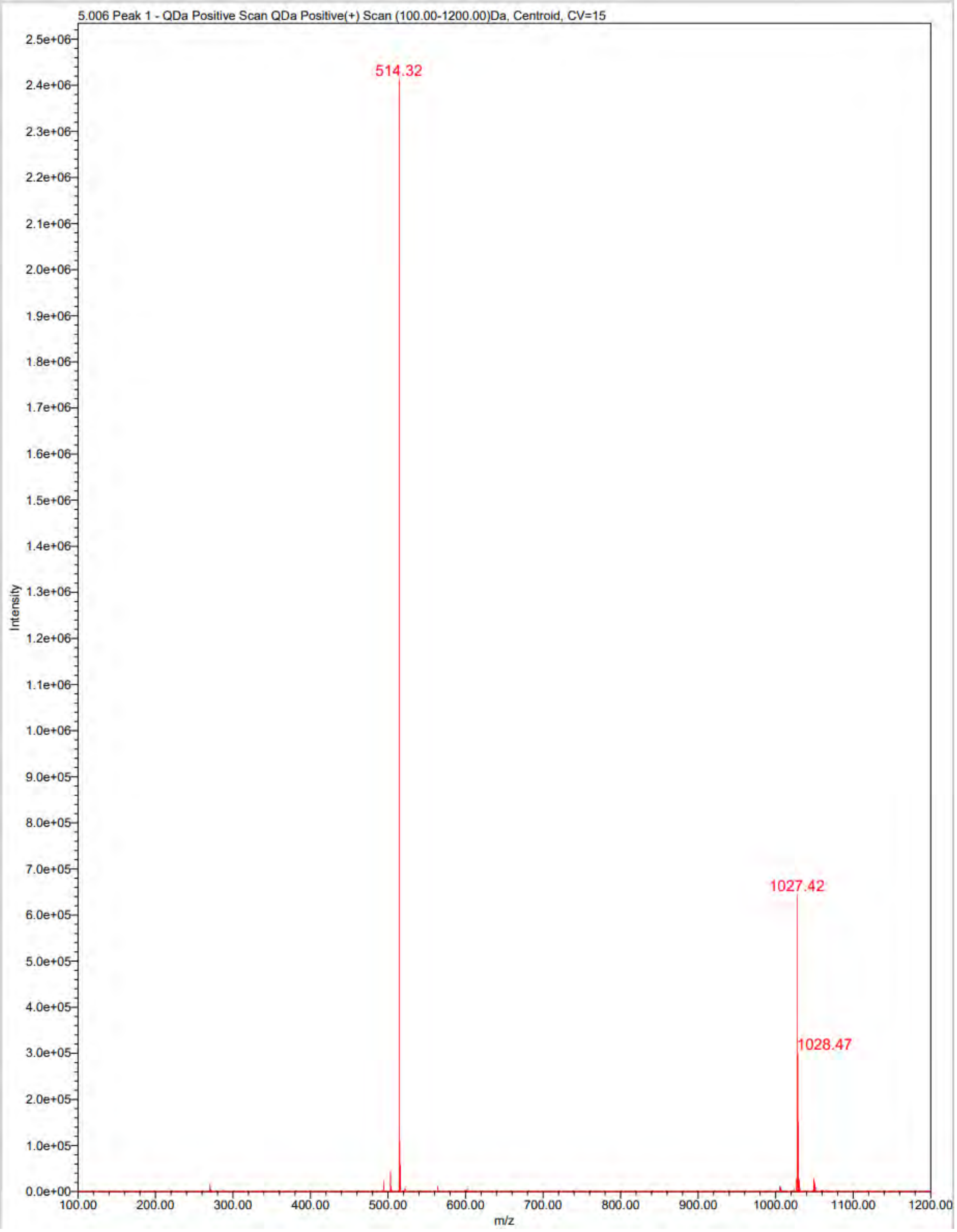

5.006 Peak 1 - QDa Positive Scan QDa Positive(+) Scan (100.00-1200.00)Da, Centroid, CV=15
514.32
1027.42
1028.47
Intensity
m/z

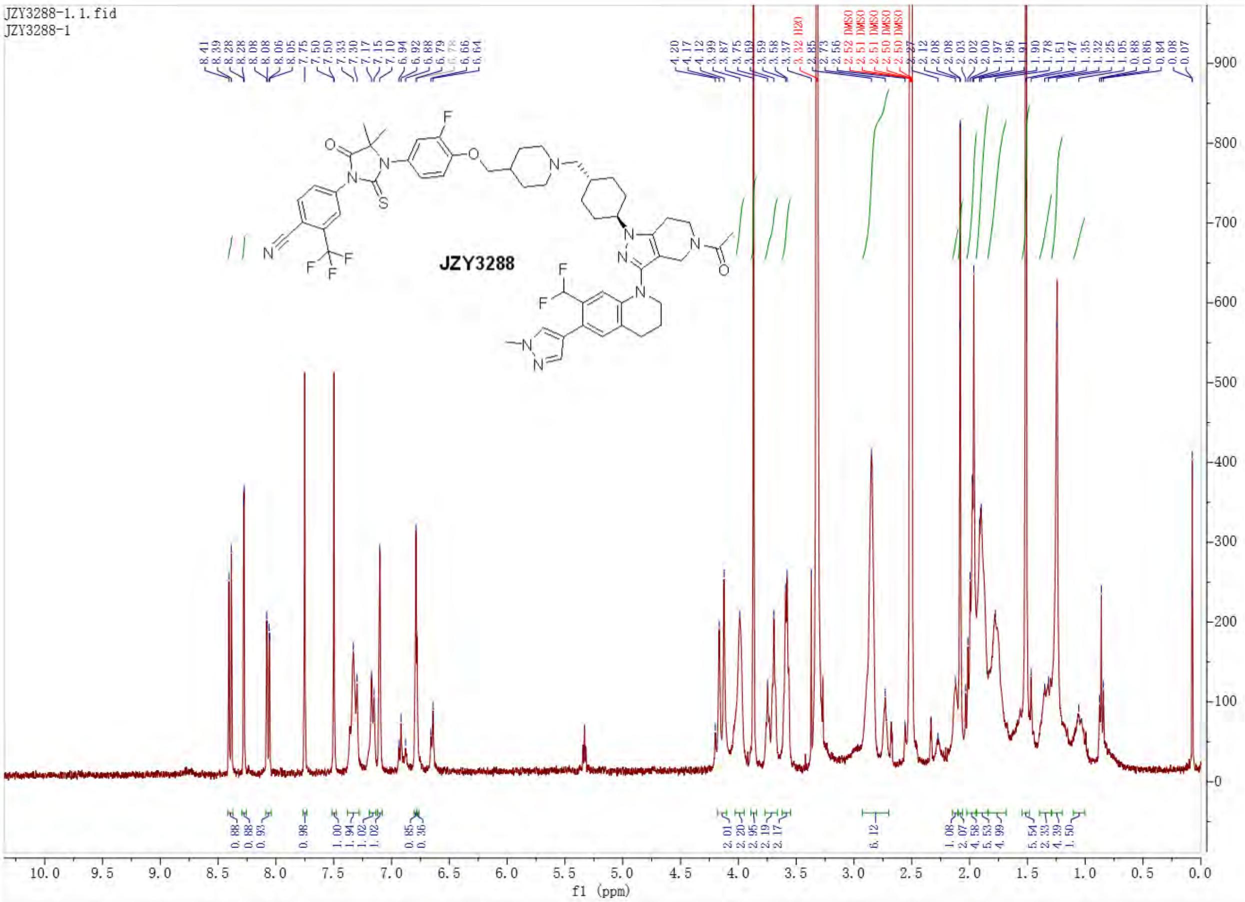

JZY3288-1.1.fid
JZY3288-1
JZY3288
f1 (ppm)

### Untitled

#### SAMPLE INFORMATION

| | | | |
|---|---|---|---|
| Sample Name: | JZY3288 | Acquired By: | System |
| Sample Type: | Unknown | Sample Set Name: | YQY |
| Vial: | 2:A,6 | Acq. Method Set: | 10to100Bin10min_5minDelay |
| Injection #: | 1 | Processing Method: | LC PQ ApexTrack |
| Injection Volume: | 3.00 ul | Channel Name: | 254.0nm@1 |
| Run Time: | 10.0 Minutes | Proc. Chnl. Descr.: | PDA Spectrum PDA 254.0 nm |
| Date Acquired: | 7/26/2026 2:52:46 AM EDT | | |
| Date Processed: | 7/26/2026 12:04:04 PM EDT | | |

**Auto-Scaled Chromatogram**

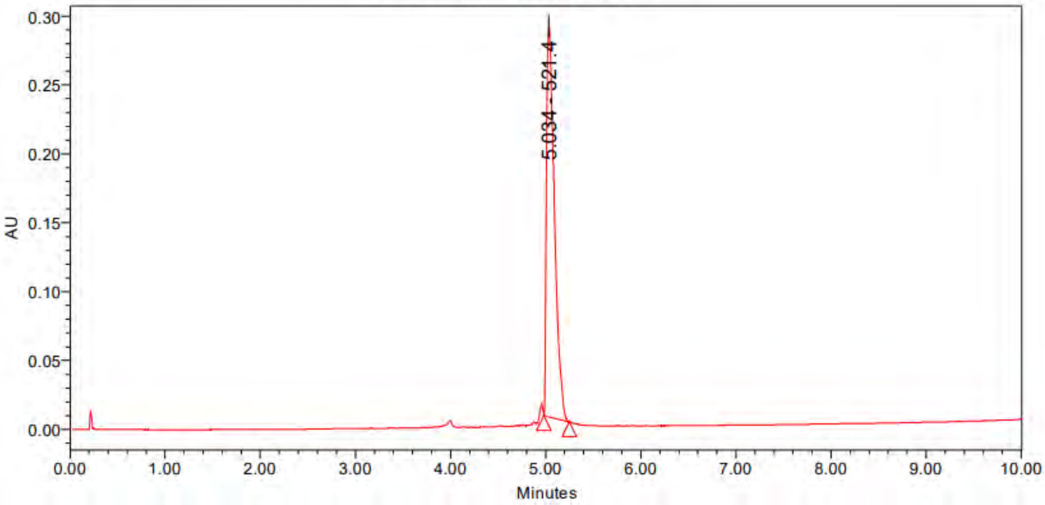

Peak Results

| | Name | RT | Area | Height | Amount | Units |
|---|---|---|---|---|---|---|
| 1 | | 5.034 | 1663947 | 283741 | | |

PDA Result Table

| | Name | RT | Purity1 Angle | Purity1 Threshold | Match1 Spect. Name | Match1 Angle | Match1 Threshold |
|---|---|---|---|---|---|---|---|
| 1 | | 5.034 | | | | | |

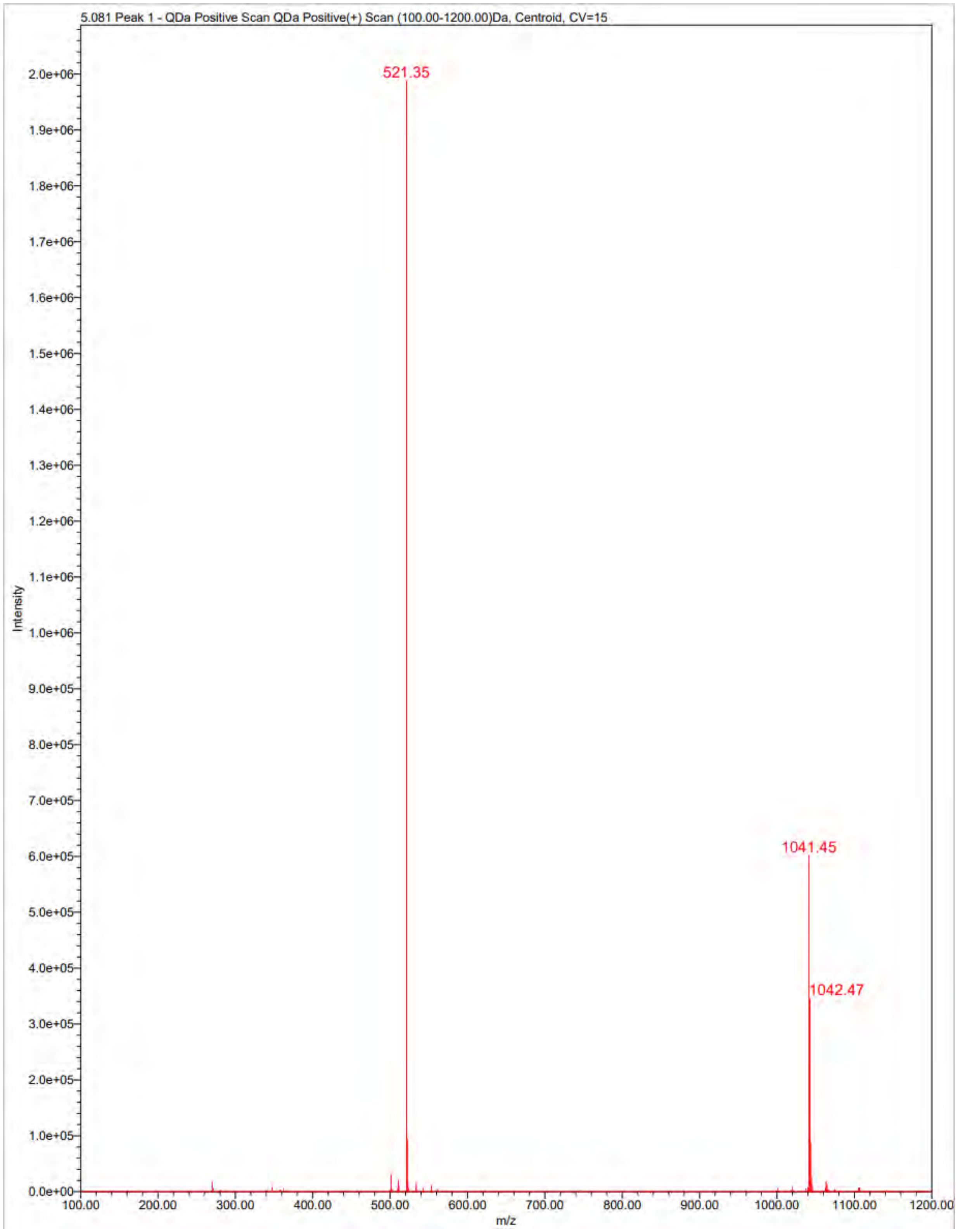
5.081 Peak 1 - QDa Positive Scan QDa Positive(+) Scan (100.00-1200.00)Da, Centroid, CV=15
521.35
1041.45
1042.47
Intensity
m/z
2.0e+06
1.9e+06
1.8e+06
1.7e+06
1.6e+06
1.5e+06
1.4e+06
1.3e+06
1.2e+06
1.1e+06
1.0e+06
9.0e+05
8.0e+05
7.0e+05
6.0e+05
5.0e+05
4.0e+05
3.0e+05
2.0e+05
1.0e+05
0.0e+00
100.00
200.00
300.00
400.00
500.00
600.00
700.00
800.00
900.00
1000.00
1100.00
1200.00

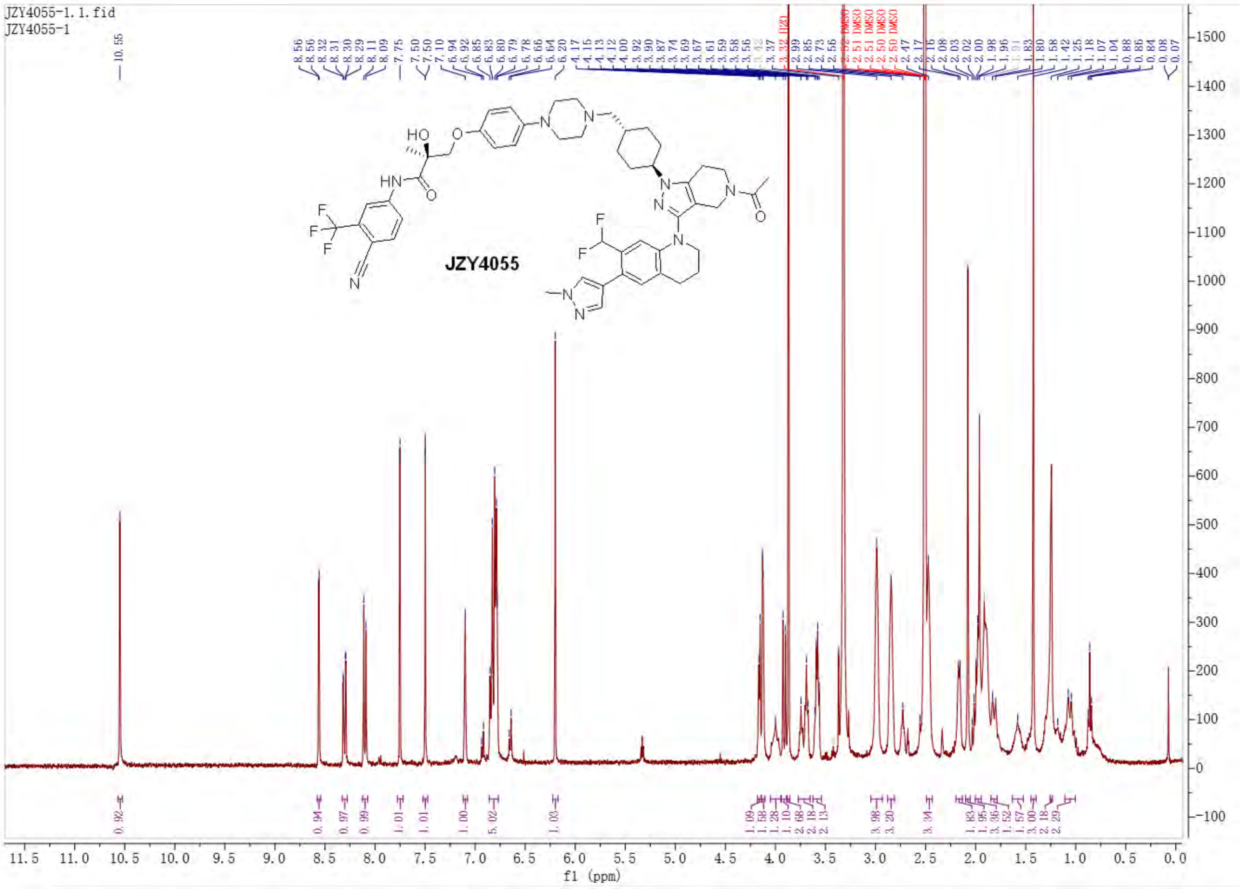

JZY4055-1.1.fid
JZY4055-1
JZY4055
f1 (ppm)

Empower 3

### Untitled

#### SAMPLE INFORMATION

| | | | |
|---|---|---|---|
| Sample Name: | JZY4055 | Acquired By: | System |
| Sample Type: | Unknown | Sample Set Name: | YQY |
| Vial: | 2:A,7 | Acq. Method Set: | 10to100Bin10min_5minDelay |
| Injection #: | 1 | Processing Method: | LC PQ ApexTrack |
| Injection Volume: | 3.00 ul | Channel Name: | 254.0nm |
| Run Time: | 10.0 Minutes | Proc. Chnl. Descr.: | PDA Spectrum PDA 254.0 nm |
| Date Acquired: | 7/26/2026 3:08:22 AM EDT | | |
| Date Processed: | 7/26/2026 12:04:30 PM EDT | | |

**Auto-Scaled Chromatogram**

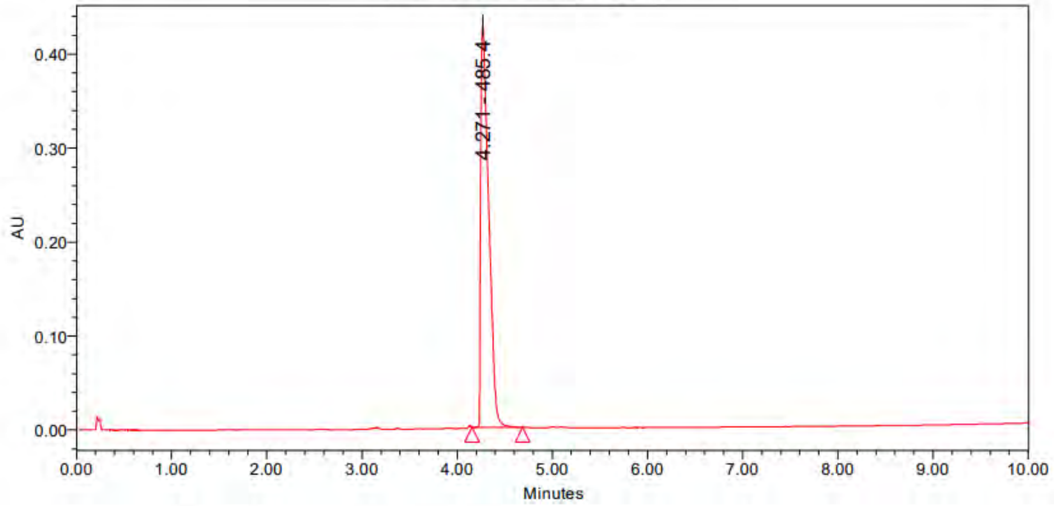

Peak Results

| | Name | RT | Area | Height | Amount | Units |
|---|---|---|---|---|---|---|
| 1 | | 4.271 | 2451117 | 427405 | | |

PDA Result Table

| | Name | RT | Purity1 Angle | Purity1 Threshold | Match1 Spect. Name | Match1 Angle | Match1 Threshold |
|---|---|---|---|---|---|---|---|
| 1 | | 4.271 | | | | | |

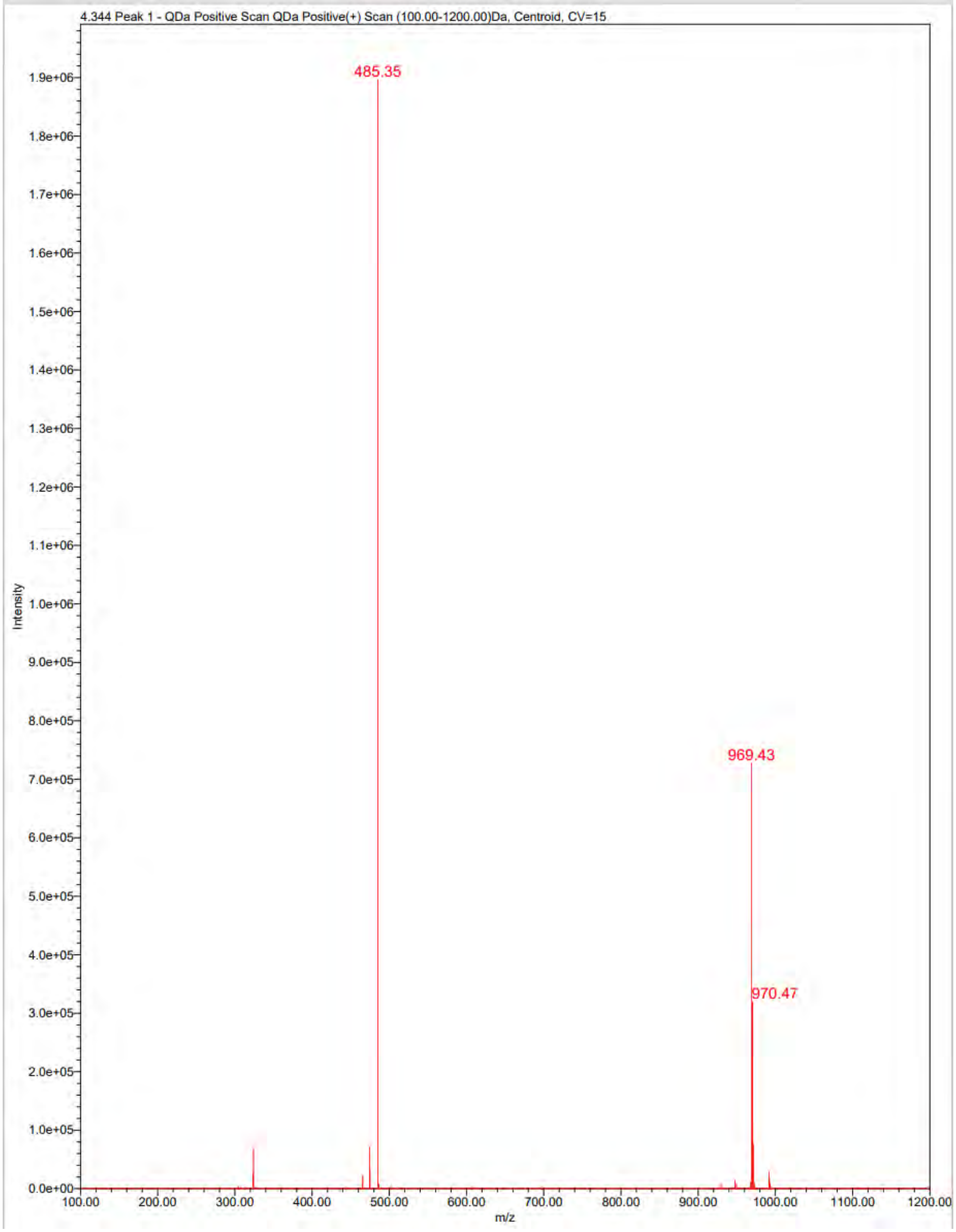
4.344 Peak 1 - QDa Positive Scan QDa Positive(+) Scan (100.00-1200.00)Da, Centroid, CV=15
485.35
969.43
970.47
Intensity
m/z
1.9e+06
1.8e+06
1.7e+06
1.6e+06
1.5e+06
1.4e+06
1.3e+06
1.2e+06
1.1e+06
1.0e+06
9.0e+05
8.0e+05
7.0e+05
6.0e+05
5.0e+05
4.0e+05
3.0e+05
2.0e+05
1.0e+05
0.0e+00
100.00
200.00
300.00
400.00
500.00
600.00
700.00
800.00
900.00
1000.00
1100.00
1200.00

### Pharmacokinetics Study

The pharmacokinetic study was performed in Shanghai Medicilon Inc in male mice, provided by Vital River Laboratory Animal Technology Co., Ltd. Mice were dosed with tested compounds solution formulation (formulation: 5% DMSO, 10% Solutol, 85% normal saline, 2 mg/kg for intravenous dose, 5 mg/kg for oral dose). Blood samples were collected at 0.083, 0.25, 0.5, 1, 2, 4, 8 and 24 h in intravenous administration, 0.25, 0.5, 1, 2, 4, 6, 8, and 24 h in oral administration. The blood samples were collected from sets of three mice for each route of administration for each compound at each time point in labeled microcentrifuge tubes containing heparin sodium as an anticoagulant. Plasma samples were separated by centrifugation (2−8 ℃, 6800 g for 6 min) within 1 h and stored below −80 ℃ until bioanalysis. All samples were processed for analysis by precipitation using acetonitrile and analyzed with a partially validated LC/MS/MS method. Pharmacokinetic parameters were calculated using the noncompartmental analysis tool of WinNonlin Enterprise software.
