## Supplemental Figures 1 - 9 (Extended Data) for "Rewiring Oncogenic Transcriptional Complexes with Domain-ALTerAtion Chimeras (DALTACs) in Prostate Cancer"

### Extended Data Figure 1

a

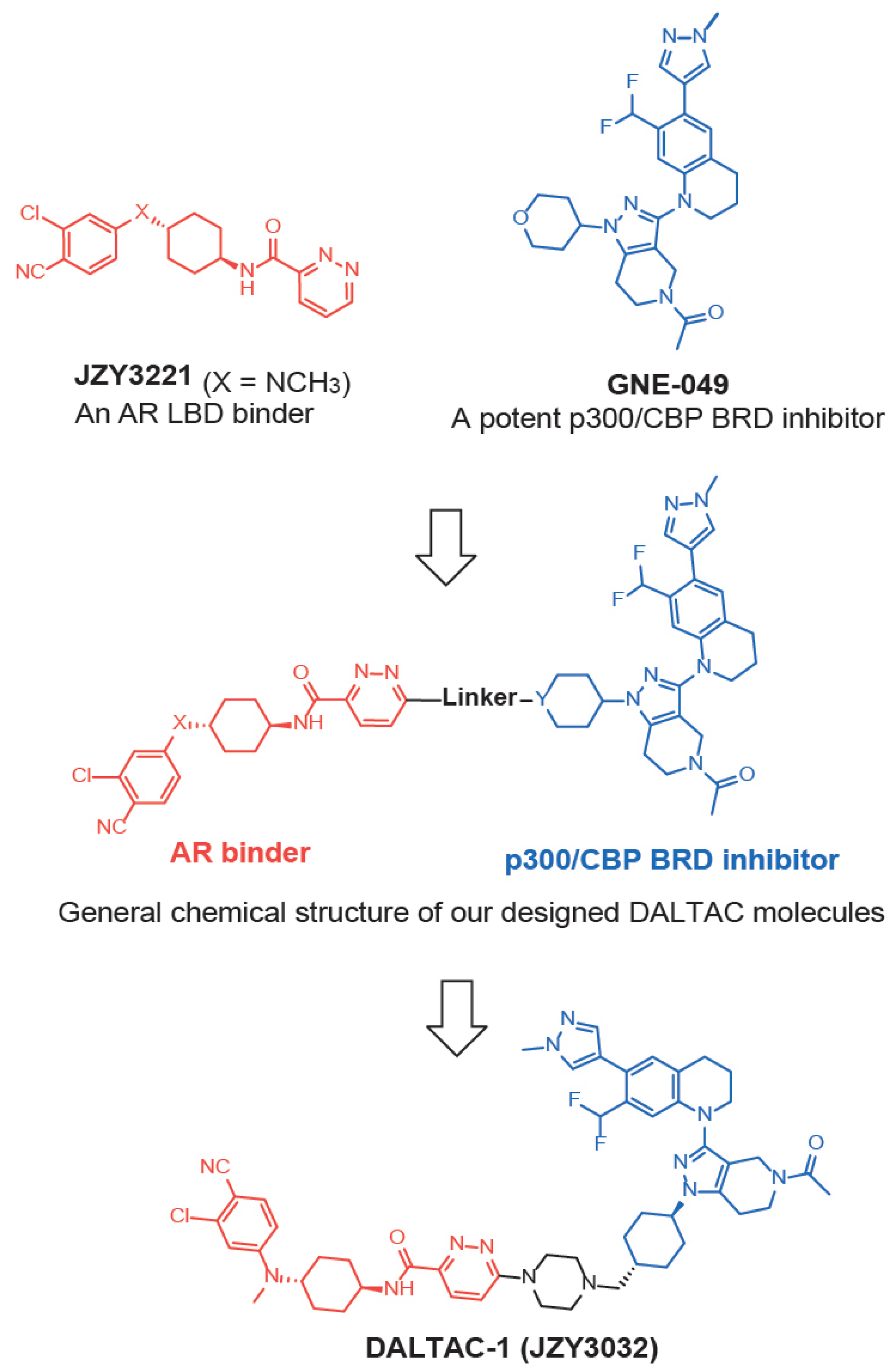


b

| Compounds | 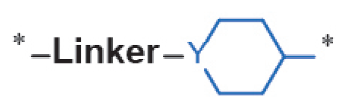 | Y | X | ternary complex formation | | cell growth inhibition in LNCaP cells | | cell growth inhibition in 22Rv1 cells | | inhibition of ARE-GFP |
|---|---|---|---|---|---|---|---|---|---|---|
| | | | | $EC_{50}$ (nM) | max. induction fold at 300 nM | $IC_{50}$ (nM) | $I_{max}$% at 3 μM | $IC_{50}$ (nM) | $I_{max}$% at 3 μM | inhibition% at 10 nM |
| GNE-049 | - | - | - | - | - | 455.4 | 52 | > 1μM | 40 | 21 |
| JZY3221 | - | - | $NCH_3$ | - | - | > 1μM | 46 | > 1μM | 10 | n/a |
| GNE-049 +JZY3221 | - | - | - | n/a | 1.33 | 588.1 | 60 | > 1μM | 25 | 13 |
| 1 (JZY3257) | 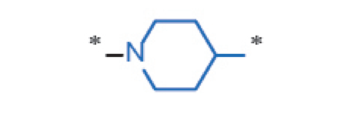 | N | O | n/a | 1.55 | 106.3 | 67 | 463 | 55 | 36.1 |
| 2 (JZY3256) | 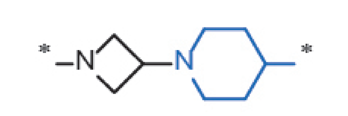 | N | O | n/a | 1.55 | 33.2 | 69 | 16.4 | 70 | 53.9 |
| 3 (JZY3254) | 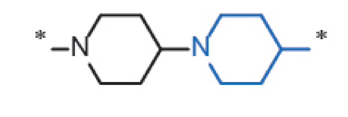 | N | O | n/a | 2.61 | 7.1 | 56 | 18.1 | 63 | 47.8 |
| 4 (JZY3255) | 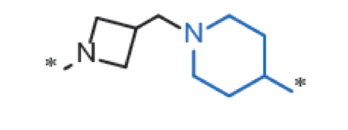 | N | O | n/a | 1.23 | 6.9 | 60 | 19.6 | 55 | 54.3 |
| 5 (SC3246) | 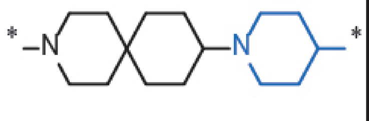 | N | O | 1.71 | 3.55 | 0.03 | 55 | 13.4 | 65 | 55.8 |
| 6 (SC3247) | 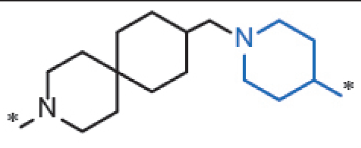 | N | O | 4.66 | 4.25 | 0.12 | 65 | 15.6 | 67 | 50.7 |
| 7 (JZY3031) | 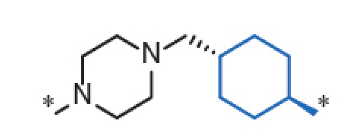 | CH | O | 0.16 | 4.84 | 1.9 | 72 | 2.6 | 70 | 58.7 |
| DALTAC-1 (JZY3032) | 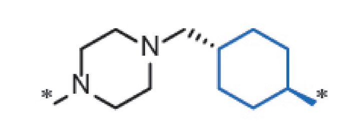 | CH | $NCH_3$ | 0.22 | 5.69 | 0.26 | 66 | 1.0 | 69 | 64.8 |

c

| PK parameters | compound 7 (JZY3031) | | DALTAC-1 (JZY3032) | |
|---|---|---|---|---|
| | i.v. (2.0 mg/kg) | p.o. (5.0 mg/kg) | i.v. (2.0 mg/kg) | p.o. (5.0 mg/kg) |
| $T_{1/2}$ (h) | 3.2 | 2.5 | 5.0 | 5.1 |
| $C_{max}$(ng/mL) | 5582 | 268.6 | 5906 | 284 |
| *AUC*(0-24) (h*ng/mL) | 4930 | 798.4 | 6394 | 2952 |
| $V_{ss}$ (L/kg) | 1.0 | - | 1.3 | - |
| *CL*(mL/min/kg) | 6.7 | - | 5.2 | - |
| *F* (%) | - | 6.5 | - | 18.5 |

d

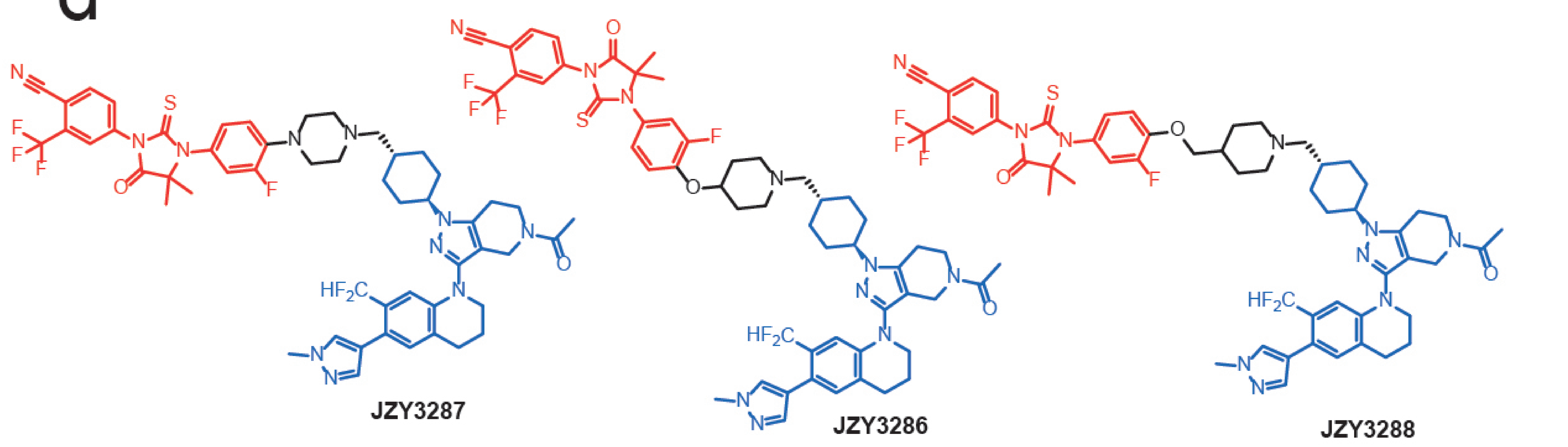


e

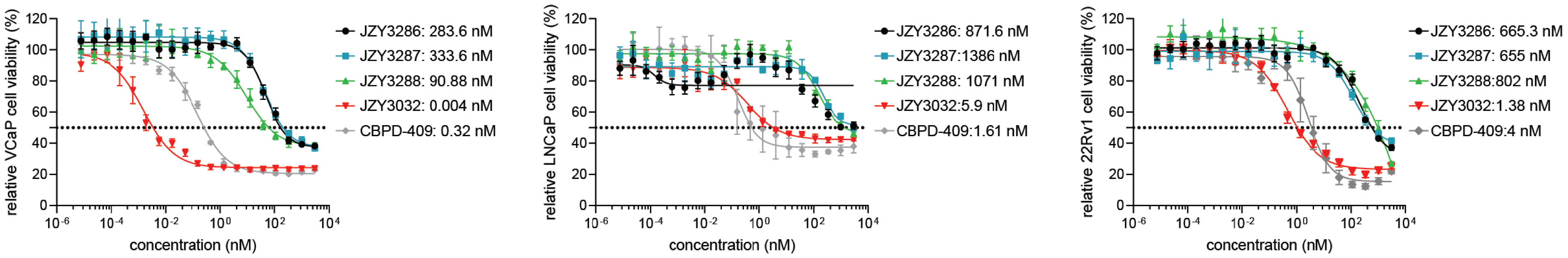


f

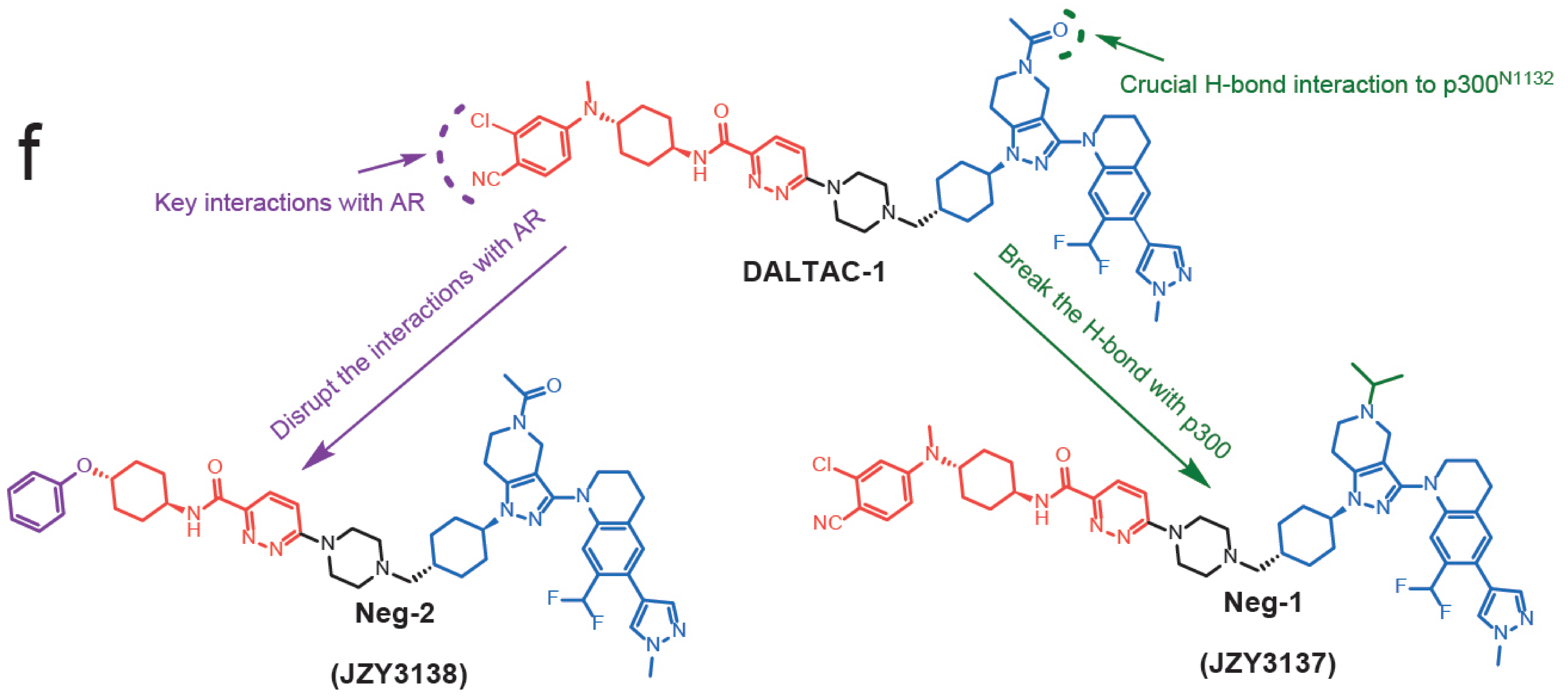


g

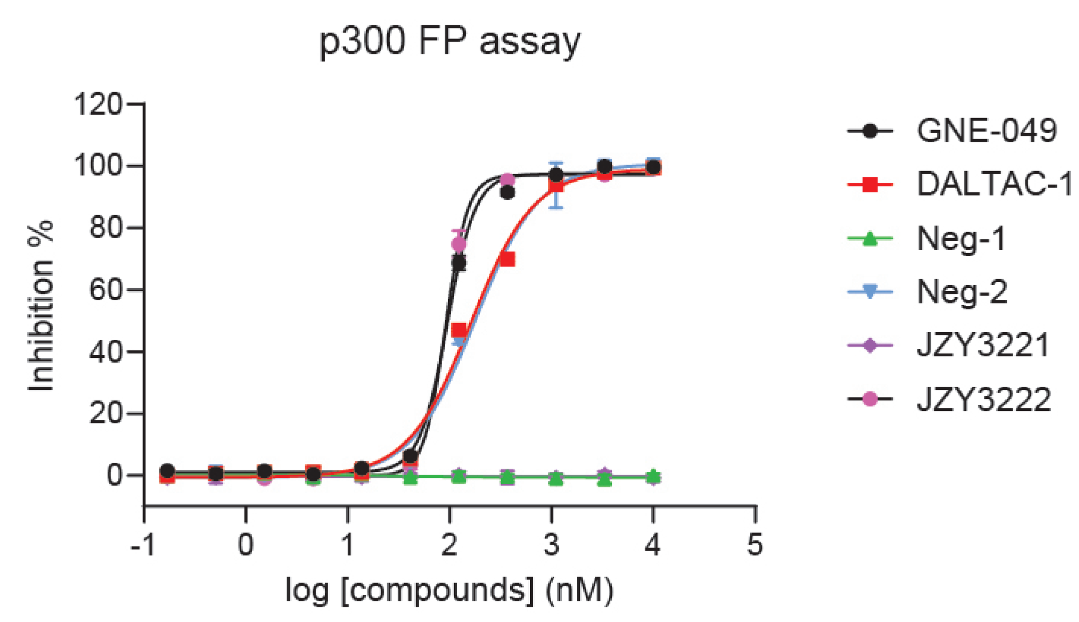


h

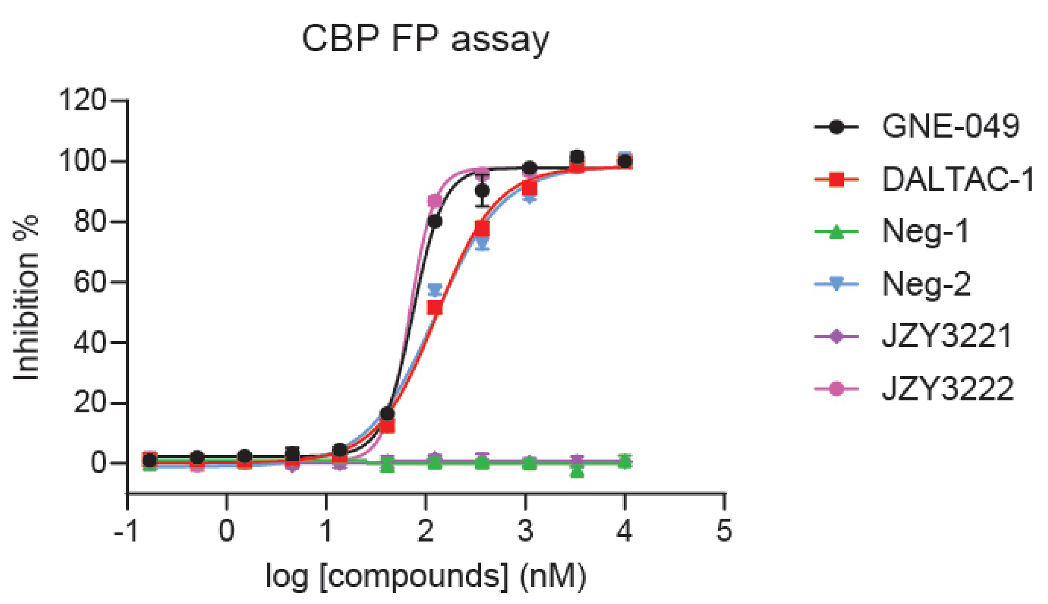


i

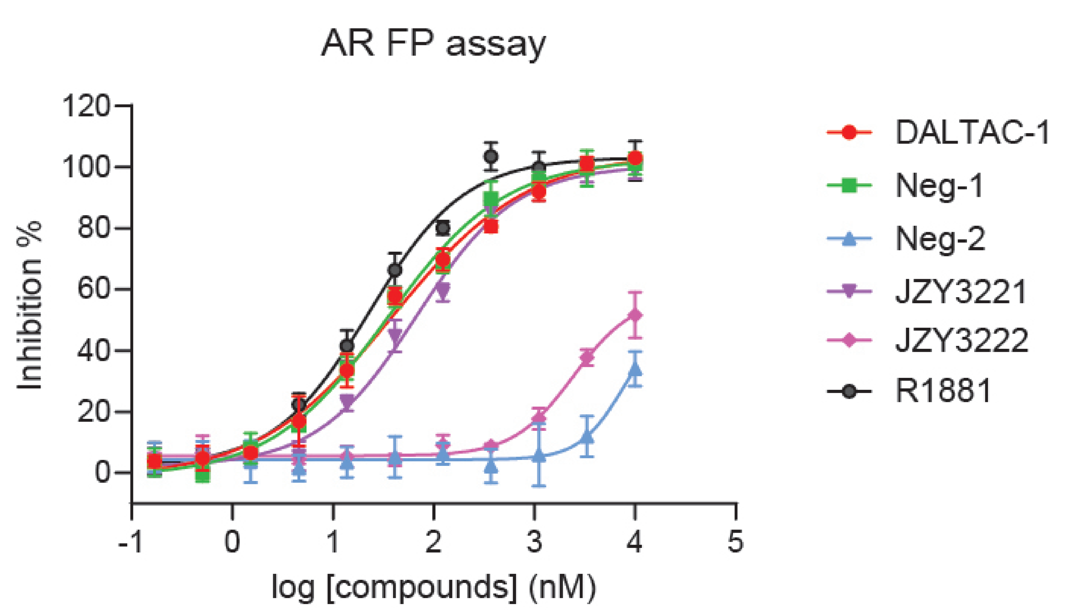


j

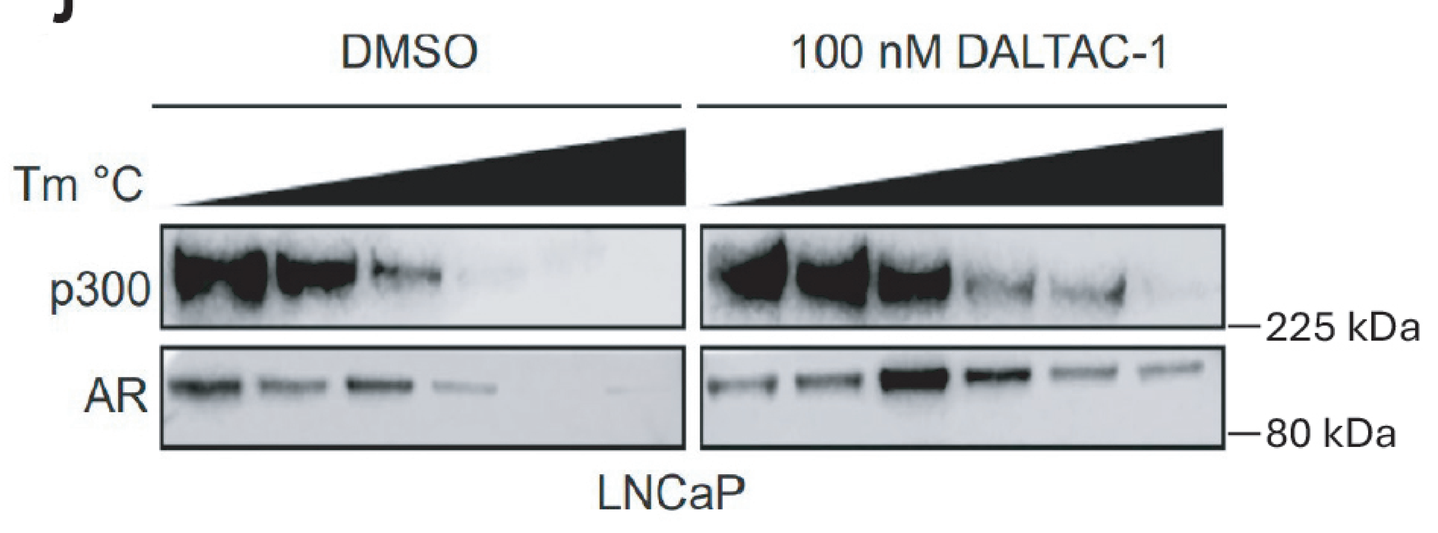

Extended Data Figure 2

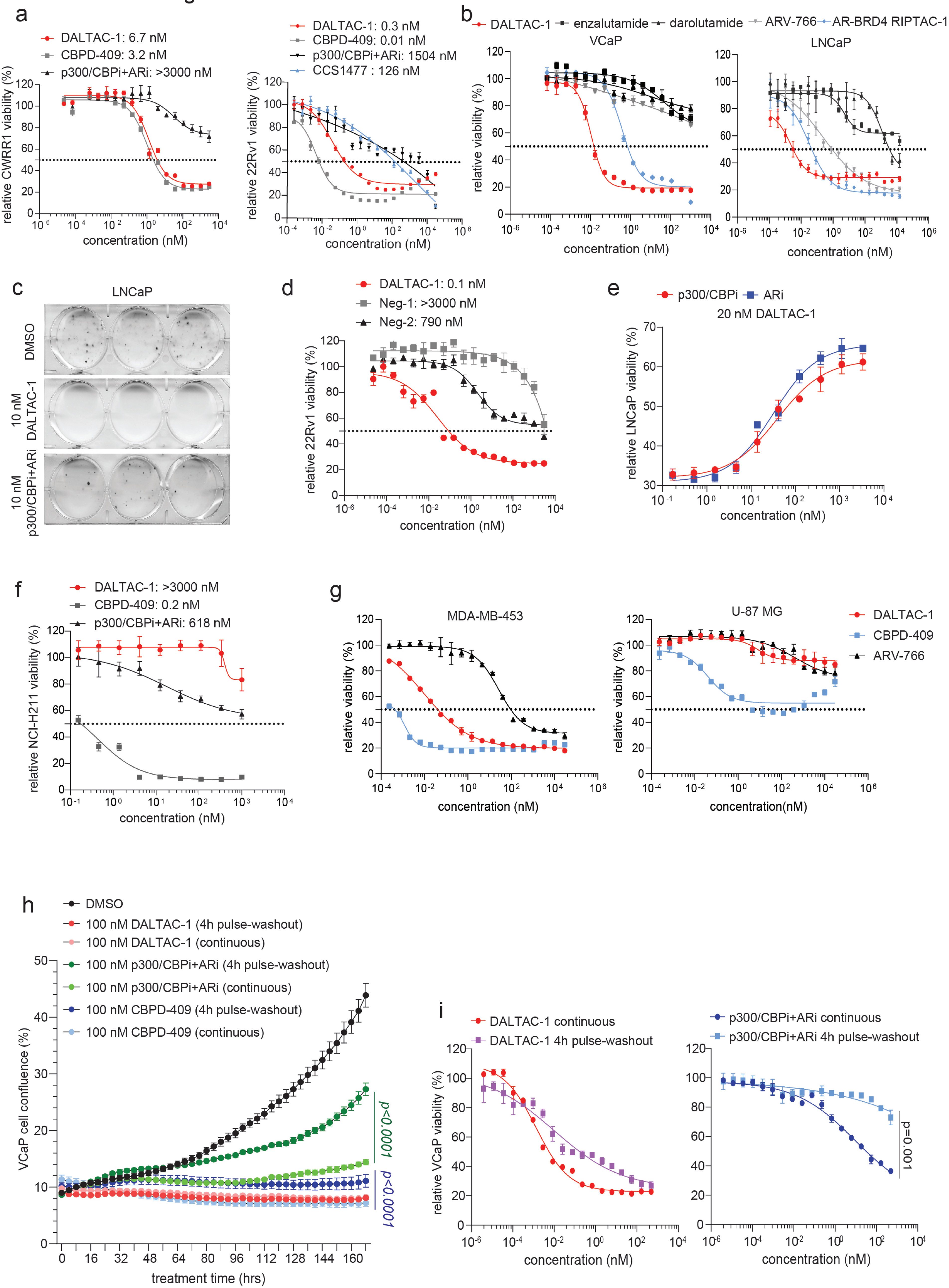

### Extended Data Figure 3

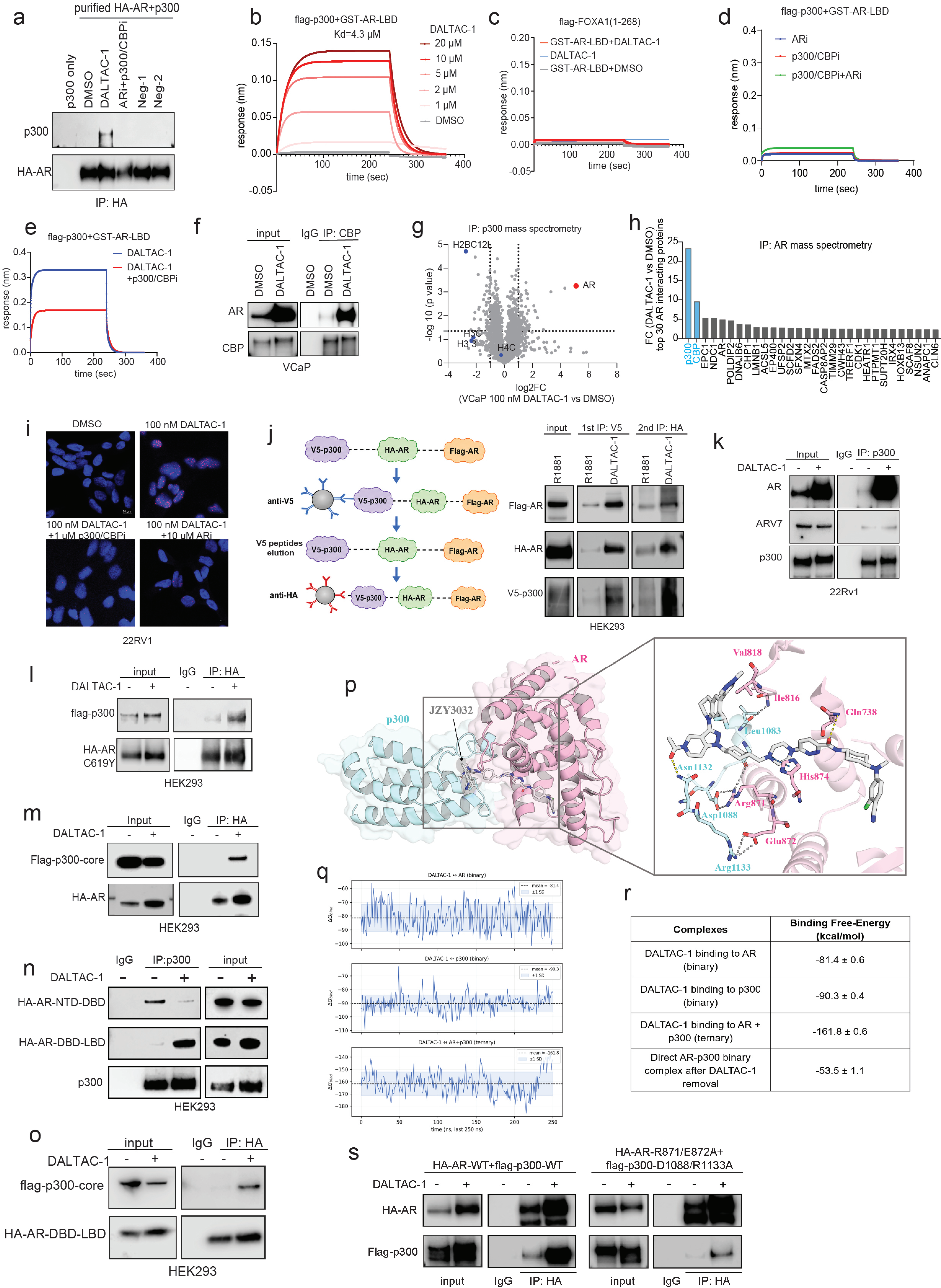


| Complexes | Binding Free-Energy (kcal/mol) |
|---|---|
| DALTAC-1 binding to AR (binary) | -81.4 ± 0.6 |
| DALTAC-1 binding to p300 (binary) | -90.3 ± 0.4 |
| DALTAC-1 binding to AR + p300 (ternary) | -161.8 ± 0.6 |
| Direct AR-p300 binary complex after DALTAC-1 removal | -53.5 ± 1.1 |

### Extended Data Figure 4

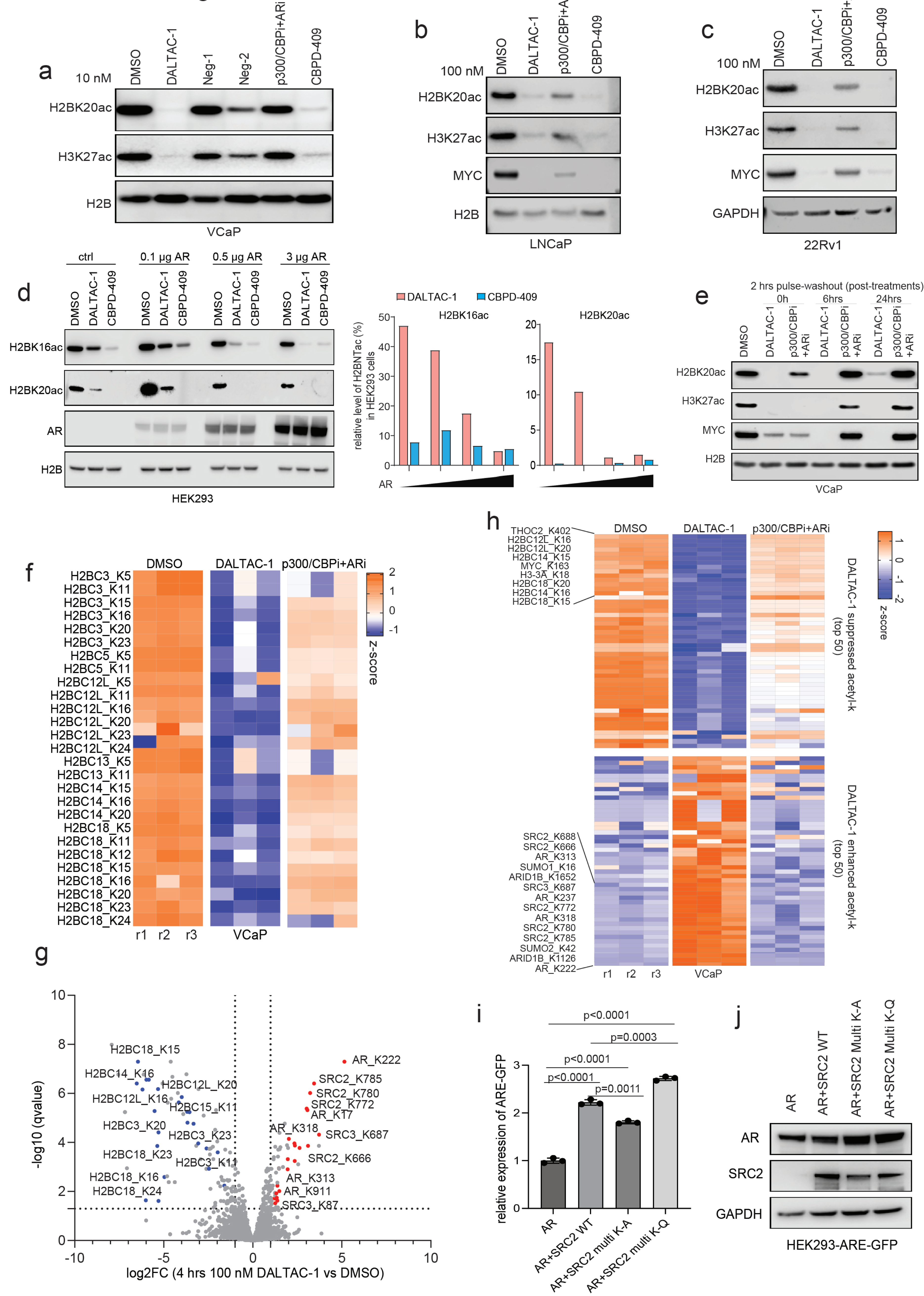

### Extended Data Figure 5

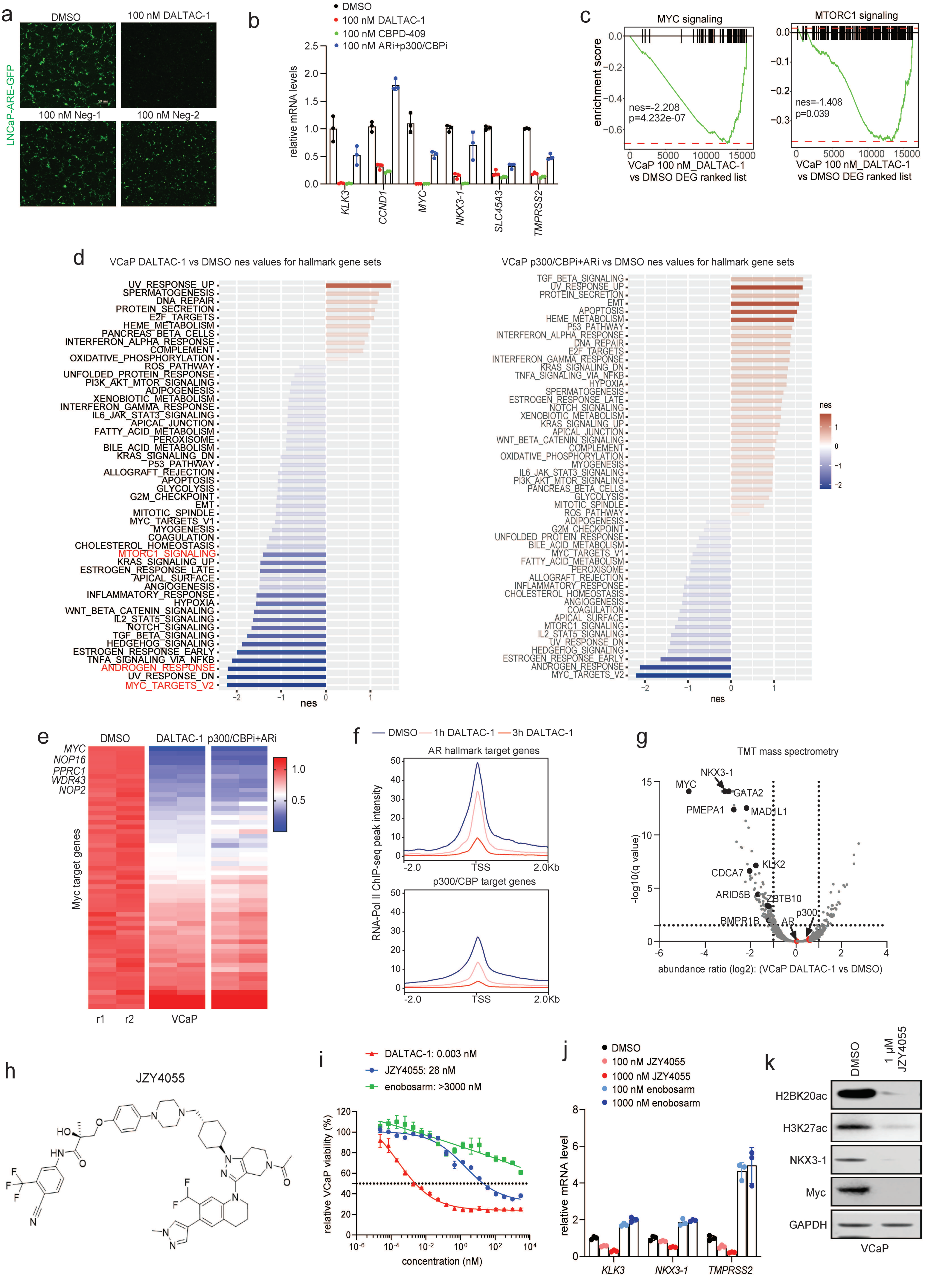

### Extended Data Figure 6

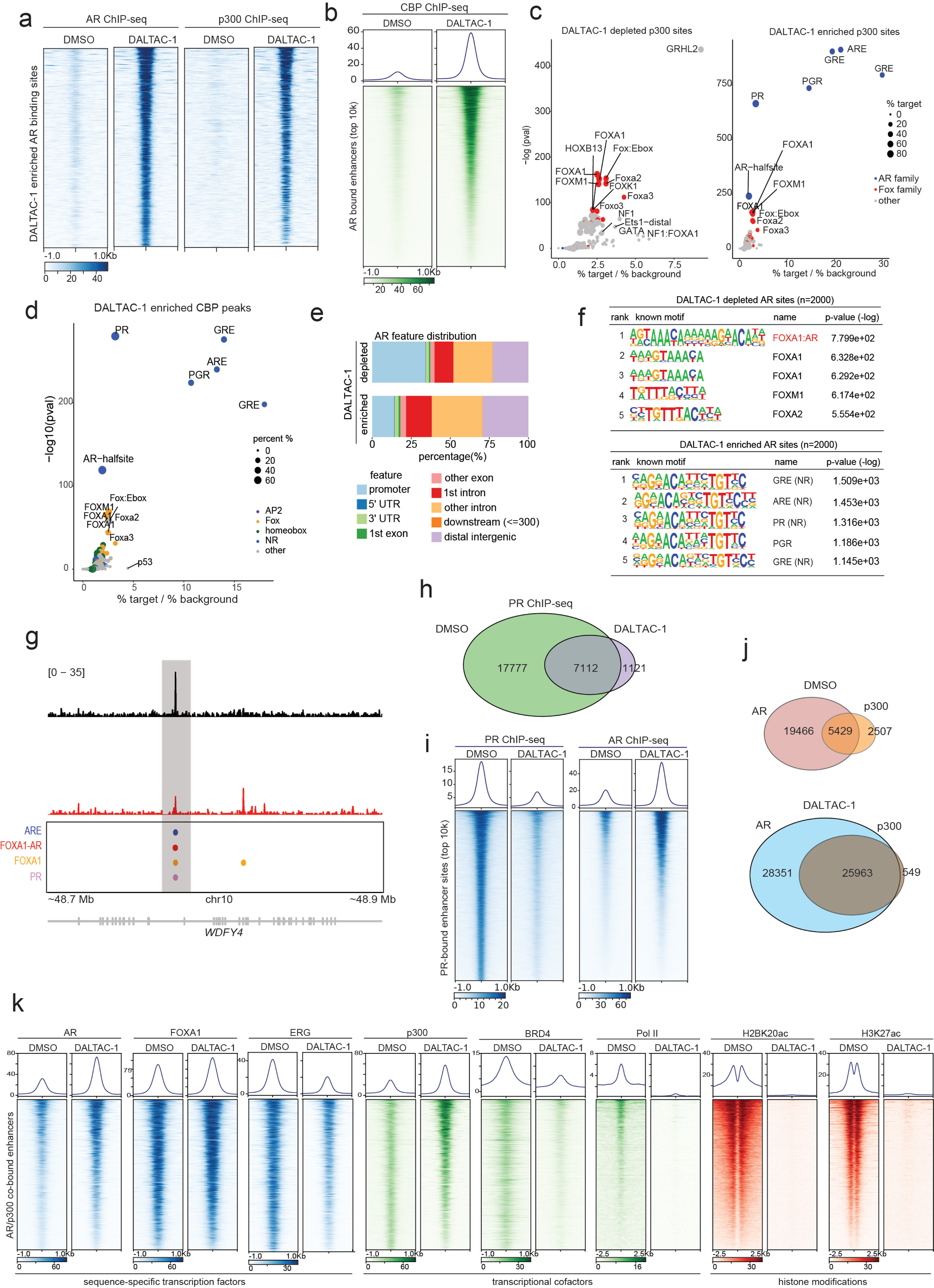

Extended Data Figure 7

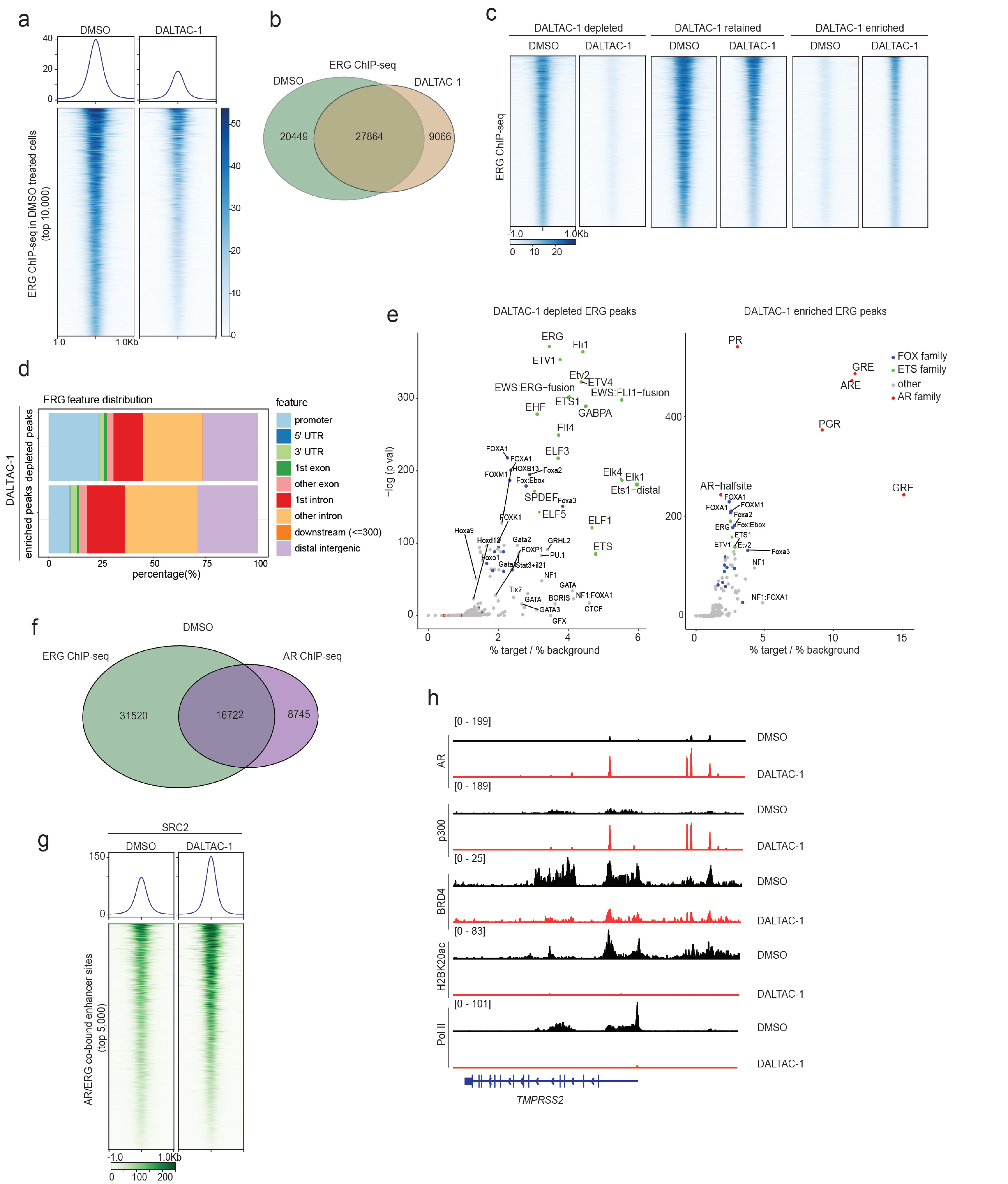

### Extended Data Figure 8

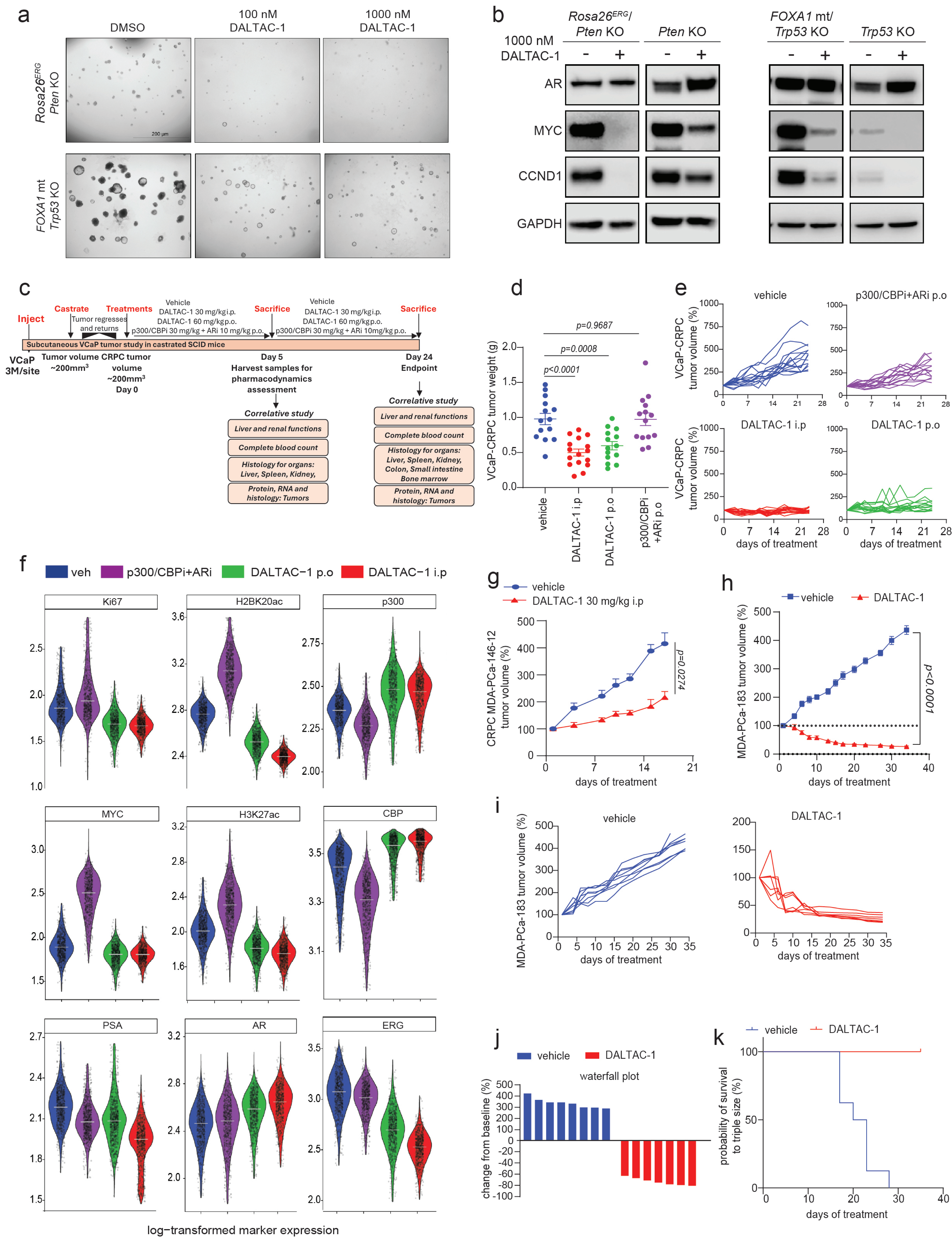

### Extended Data Figure 9

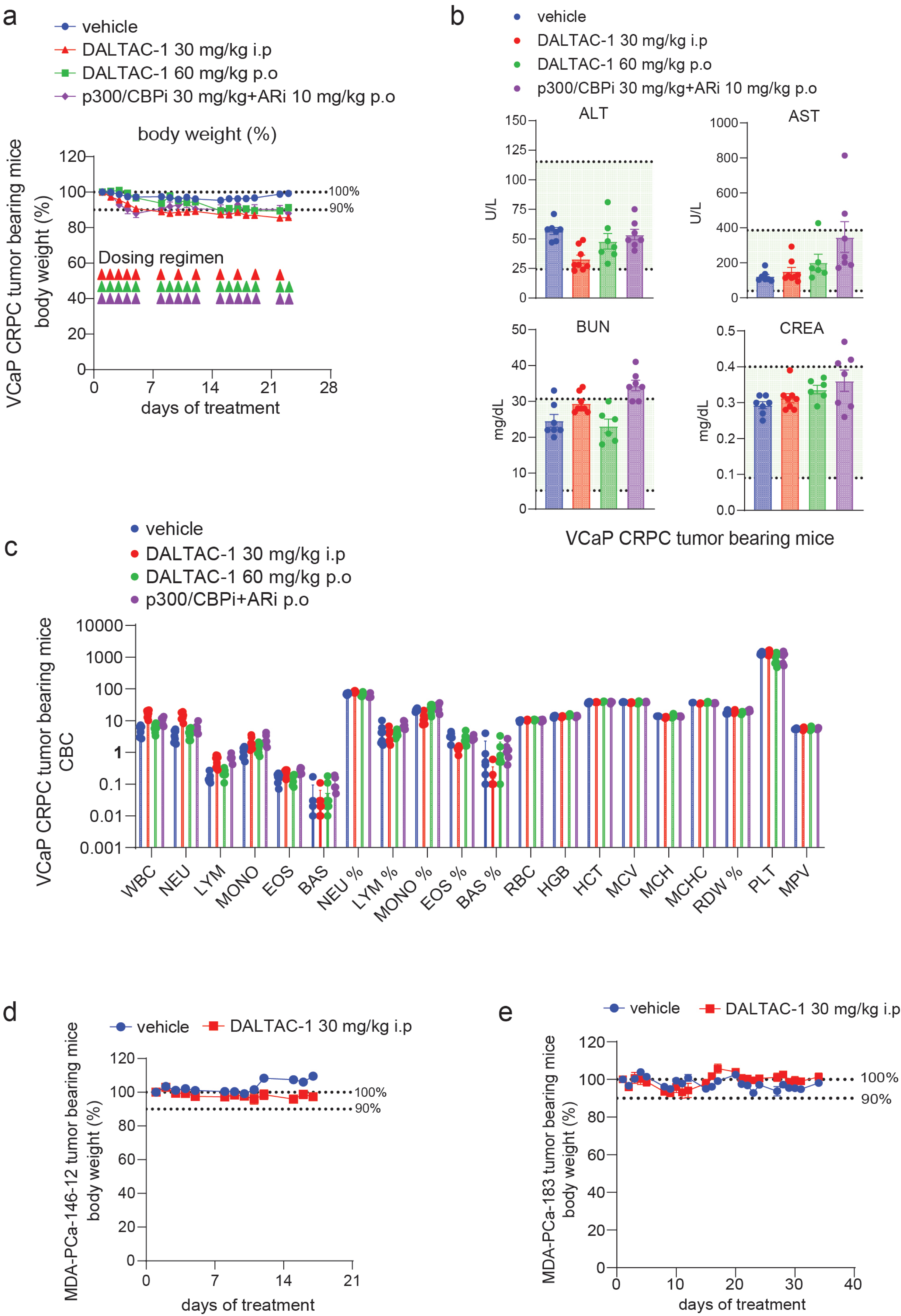
